# AMRgen: an R package for antimicrobial resistance genotype-phenotype analysis

**DOI:** 10.64898/2026.05.01.722195

**Authors:** KE Holt, S Argimón, DL Chaput, N Couto, ZA Dyson, E Foster-Nyarko, RN Goodman, J Hawkey, GM Knight, D Nagy, AB Prasad, L Sánchez-Busó, KK Tsang, MS Berends

## Abstract

Microbial whole-genome sequence data is now generated at scale, including to support antimicrobial resistance (AMR) surveillance and understand resistance mechanisms, yet analytical infrastructure for systematically linking AMR genotypes to measured phenotypes remains fragmented. Here we present *AMRgen*, an *R* package to support systematic AMR genotype-phenotype analysis. *AMRgen* imports and harmonises genotypic data from common bioinformatics tools, alongside phenotypic data from automated antimicrobial susceptibility testing instruments and public repositories. It supports common analyses linking data to reference distributions, modelling associations, quantifying concordance, and producing publication-ready visualisations including UpSet plots that jointly display genotypic marker combination frequencies and associated phenotypic distributions. We demonstrate *AMRgen*’s utility using publicly available surveillance data for World Health Organization priority AMR pathogens, *Neisseria gonorrhoeae, Klebsiella pneumoniae, Escherichia coli* and *Salmonella enterica. AMRgen*, available free and open-source at https://AMRgen.org, provides a reproducible end-to-end foundation for genotype-phenotype research in AMR genomics, clinical microbiology, and public health surveillance.

## Introduction

The expansion of whole-genome sequencing (WGS) into routine pathogen genomics has generated an unprecedented volume of microbial genomic data, a key use for which is the analysis of antimicrobial resistance (AMR) genotype profiles for clinical or public health surveillance^1^. A rapidly growing body of studies now links specific resistance determinants, including acquired genes, point mutations, and mobile genetic elements, to phenotypic outcomes such as minimum inhibitory concentrations (MICs) and clinical susceptibility categories, across a widening range of bacterial species and drug classes^2–5^. This work has advanced our understanding of resistance mechanisms, informed the development of interpretive systems that translate genotypes into predicted phenotypes, and provided the empirical basis for genotype-guided diagnostics and resistance catalogues^6–8^. Yet despite this momentum, the analytical infrastructure for systematically comparing genomic predictions with measured phenotypes remains fragmented: studies typically rely on bespoke, project-specific code, which is time-consuming and limits reproducibility and cross-study comparability.

A core reason for this methodological fragmentation is that AMR genotype-phenotype relationships are rarely straightforward. Resistance phenotypes can arise from complex combinations of determinants (e.g., enzyme production combined with porin loss and efflux up-regulation^9^), allelic heterogeneity with variable effect sizes^10,11^, and background-dependent epistasis^12^, while measured phenotypes are themselves subject to MIC censoring, platform effects, and interpretive differences between clinical breakpoints and/or epidemiological cut-offs (ECOFFs)^13^ for which there are varying standards (e.g. Clinical & Laboratory Standards Institute (CLSI) or European Committee on Antimicrobial Susceptibility Testing (EUCAST)) that change over time. As a result, analysts are frequently confronted with discordant patterns, either genotype-positive but phenotype-susceptible (major error) and/or genotype-negative but phenotype-resistant (very major error), that may reflect incomplete reference databases, novel or uncharacterised mechanisms, expression-level variation, or differences in interpretive frameworks^2^. The need for integrated analytical tools that quantify concordance, capture uncertainty, and operate reproducibly across datasets has been repeatedly articulated^14,15^, yet no broadly adopted, open-source framework currently provides end-to-end support for this workflow. In parallel, the curation and interpretation of bacterial AMR gene and mutation catalogues have become a major determinant of analytic validity; for example, AMRFinderPlus couples curated protein families and taxon-aware mutation detection to improve precision in resistome calling, positioning the tool as a bridge between AMR genotypes and phenotypes (while explicitly warning against interpreting the presence of genotypes as a prediction of phenotype due to the complexities noted above)^16^.

Within *R*, the peer-reviewed *AMR* package^17^ has become a widely adopted foundation for reproducible AMR analytics on phenotypic data: harmonising microorganism and antimicrobial identifiers, validating antimicrobial susceptibility testing (AST) results, and supporting guideline-based interpretation using ECOFFs and clinical breakpoints (including both CLSI and EUCAST) to produce standardised epidemiological summaries. Yet, it does not ingest or analyse genotypic resistance determinants such as acquired genes, point mutations, or sequencing-derived resistome profiles. As a result, genotype and phenotype data are typically processed in separate workflows, and integrated analyses rely on bespoke code, limiting reproducibility and comparability across studies.

Here, we developed the *AMRgen* R package to address this methodological bottleneck by extending the AMR data science ecosystem toward explicit genotype-phenotype integration. The design premise is that robust integration requires: (1) harmonised import of heterogeneous upstream sources (automated AST instruments, laboratory information management systems (LIMS), public repositories, and varied genotyping pipelines), (2) integration of matched genotype and phenotype data across isolates, and (3) data analysis and visualisations that support both descriptive exploration and inferential modelling of genotypic marker sets. In *AMRgen*, these objectives are operationalised by producing standardised intermediate objects (including genotype tables, phenotype tables, and binary geno-pheno matrices) that enable systematic quantification of concordance/discordance, comparison of phenotypic distributions across marker combinations, and estimation of marker effects while accommodating sparsity and separation common in AMR datasets. This explicitly supports the epidemiological reality that resistance often emerges from combinations of determinants rather than single markers, for which appropriate visualisation is essential. UpSet plots provide a matrix-based representation of set intersections^18^, enabling clear display of overlapping gene or mutation combinations together with their frequencies, and are particularly well suited to situations where multiple resistance determinants co-occur. Their use has increased in genomic epidemiology as an alternative to traditional Venn diagrams^11,19,20^, especially when the number of sets exceeds three or four. However, generating publication-ready UpSet plots that are tightly integrated with phenotype summaries and stratified resistance outcomes are technically cumbersome and require extensive data reshaping and custom annotation. *AMRgen* streamlines this process by directly generating UpSet-style visualisations from standardised geno-pheno matrices, linking marker combinations to phenotypic categories and quantitative measures, and thereby lowering the technical barrier for rigorous combination-aware analysis within reproducible AMR workflows.

*AMRgen* also aims to strengthen interpretive context by linking phenotype distributions to widely used epidemiological concepts such MIC reference distributions and ECOFFs, which are central to distinguishing wild-type from non-wild-type populations and to interpreting borderline phenotypes^21,22^. EUCAST’s MIC/zone distribution resources and methodological discussions highlight how distributional thinking complements breakpoint categories and improves representativeness across diverse data sources^2^. By enabling integrated genotype-phenotype summaries that can be interpreted alongside such distributional reference frameworks, *AMRgen* aims to support more defensible mechanistic hypotheses and more transparent reporting of genotype-phenotype relationships.

In this methodological paper, we describe *AMRgen*’s architecture and workflows aimed at diverse users including microbial epidemiologists, computational microbiologists, and bioinformaticians, who require reproducible genotype-phenotype integration at scale. We outline how *AMRgen* extends the *AMR* package’s phenotype-centric standardisation to the genotype-phenotype interface by supporting harmonised data ingestion, interaction-relevant modelling, and automated plotting, thereby enabling analysts to move from parallel reporting towards integrated inference that better reflects the biological and measurement complexities of AMR. Our overarching aim is to provide a robust and reproducible framework that reduces the manual effort traditionally required to curate, align, and analyse combined phenotypic and genotypic AMR data.

### Implementation

The core goal of the *AMRgen* package is to analyse relationships between AMR genotypes and phenotypes. To achieve this *AMRgen* includes functions to import and inspect AMR genotype and phenotype data from a variety of common bioinformatics and AST platforms, and parse these into standardised genotype and phenotype tables, as well as functions to compare and analyse genotype and phenotype data for an antimicrobial agent of interest, taking the standardised genotype and phenotype tables as input (**Figure 1**). Analysis functions focus on (i) calculating positive predictive value (PPV) for genetic markers as predictors of resistance phenotypes, alone or in combination, (ii) fitting logistic regression models for prediction of phenotypes from genetic markers, (iii) summarising the distribution of assay measures (MIC and disk diffusion zones) associated with different markers and combinations, and (iv) evaluating concordance between phenotypes and genotypic predictions. The analysis functions return not only summary results tables but also relevant data visualisations including bar plots, assay distributions, and UpSet plots to explore the combinatorial effect of genotypes (see examples in **Figures 2-6**). Helper functions are also provided to download and format genotype and phenotype data from public sequence archives (National Center for Biotechnology Information (NCBI)^16^ and the European Bioinformatics Institute (EBI)^23^), to export phenotype data in formats suitable for upload to these archives, and to download reference MIC and disk zone distributions from EUCAST. *AMRgen* utilises the *AMR* package^17^ to provide consistent parsing of antibiotic and organism names (via the ‘ab’ and ‘mo’ classes), interpretation of MIC and disk zone data against EUCAST or CLSI breakpoints (‘as.sir()’ functions), and proper handling of MIC, disk zone, and S/I/R data (‘mic’, ‘disk’ and ‘sir’ classes). Plotting functions utilise default palettes that are colourblind friendly, including the *AMR* package’s SIR palette, and tested using the ColorblindSimulator^24^. Tidyverse packages^25^ are utilised internally for data wrangling, and all plots are produced using *ggplot2* and *patchwork* so that all plotting objects output by *AMRgen* are further customisable using these packages.

**Figure 1:**
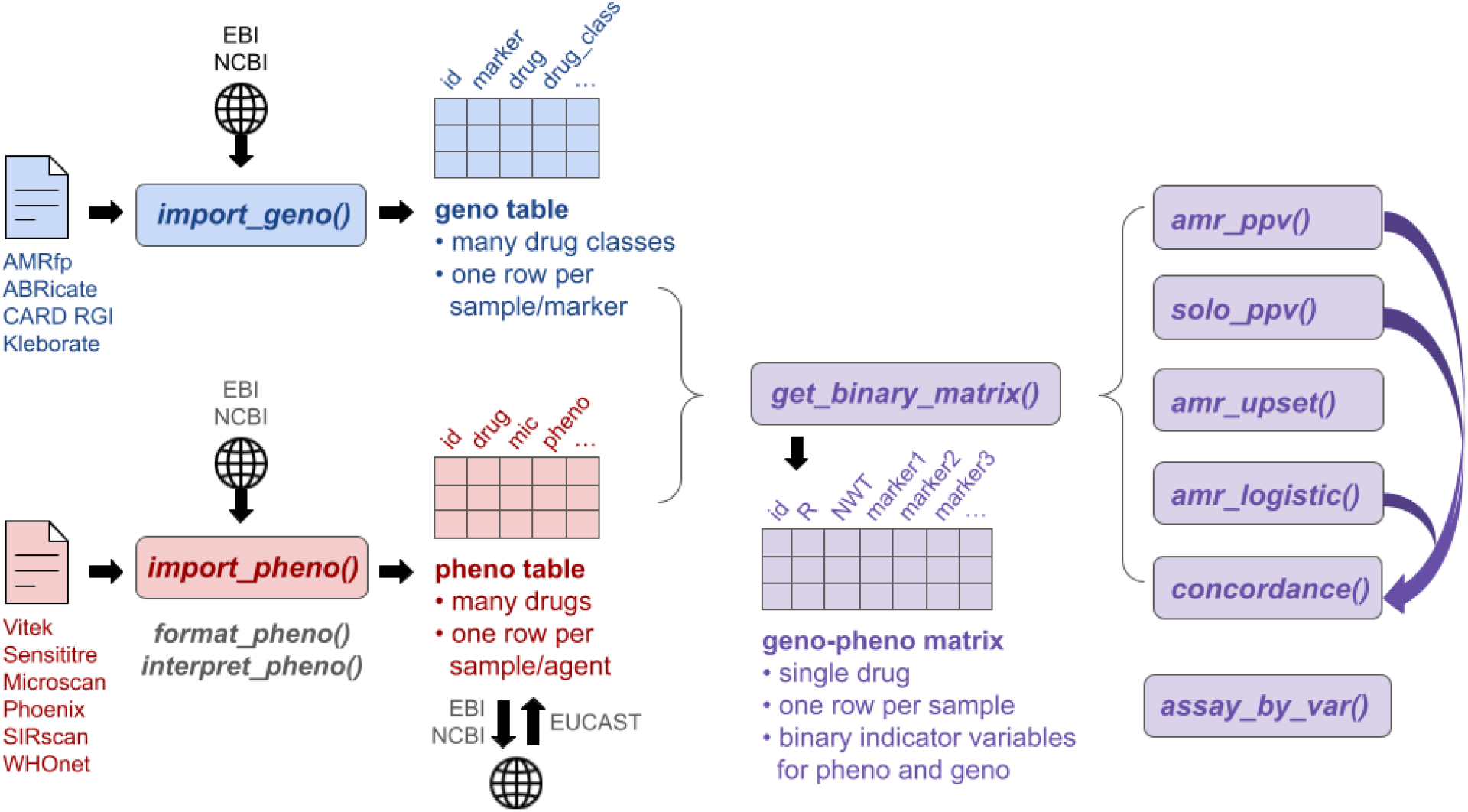
Key inputs, data objects and functions of the *AMRgen* package. Inputs (file icons), functions (bold italics labels) and data objects (grids with bold text labels) related solely to genotype data are shown in blue; phenotypes in red; and combined genotype-phenotype data in purple. Functions import_geno() and import_pheno() are dispatchers to import user data from files or objects in different standard formats (listed below the file icons); phenotype data (MIC and/or disk diffusion zones) can optionally be re-interpreted against latest breakpoints, during import or using interpret_pheno(). Functions are also provided to download and import genotype and phenotype data from public databases at NCBI or EBI, and to export phenotype data for submission to these databases; reference distributions for MIC or disk zone diameters can be automatically retrieved from EUCAST, for comparison with user data. Function get_binary_matrix() combines phenotype data for a single drug with presence/absence data for associated genotype markers. Functions amr_ppv(), solo_ppv(), amr_upset(), amr_logistic(), concordance() use this genotype-phenotype matrix to analyse and/or visualise genotype-phenotype association; these functions can take a precomputed genotype-phenotype matrix as input, or compute this matrix from separate genotype and phenotype table inputs. The assay_by_var() function provides flexible visualisation of MIC/disk assay values as histograms or boxplots, stratified by genotype or other variables. AMRfp = AMRFinderPlus, NWT = Non-wildtype, R = Resistant, PPV = Positive Predictive Value.

**Figure 2:**
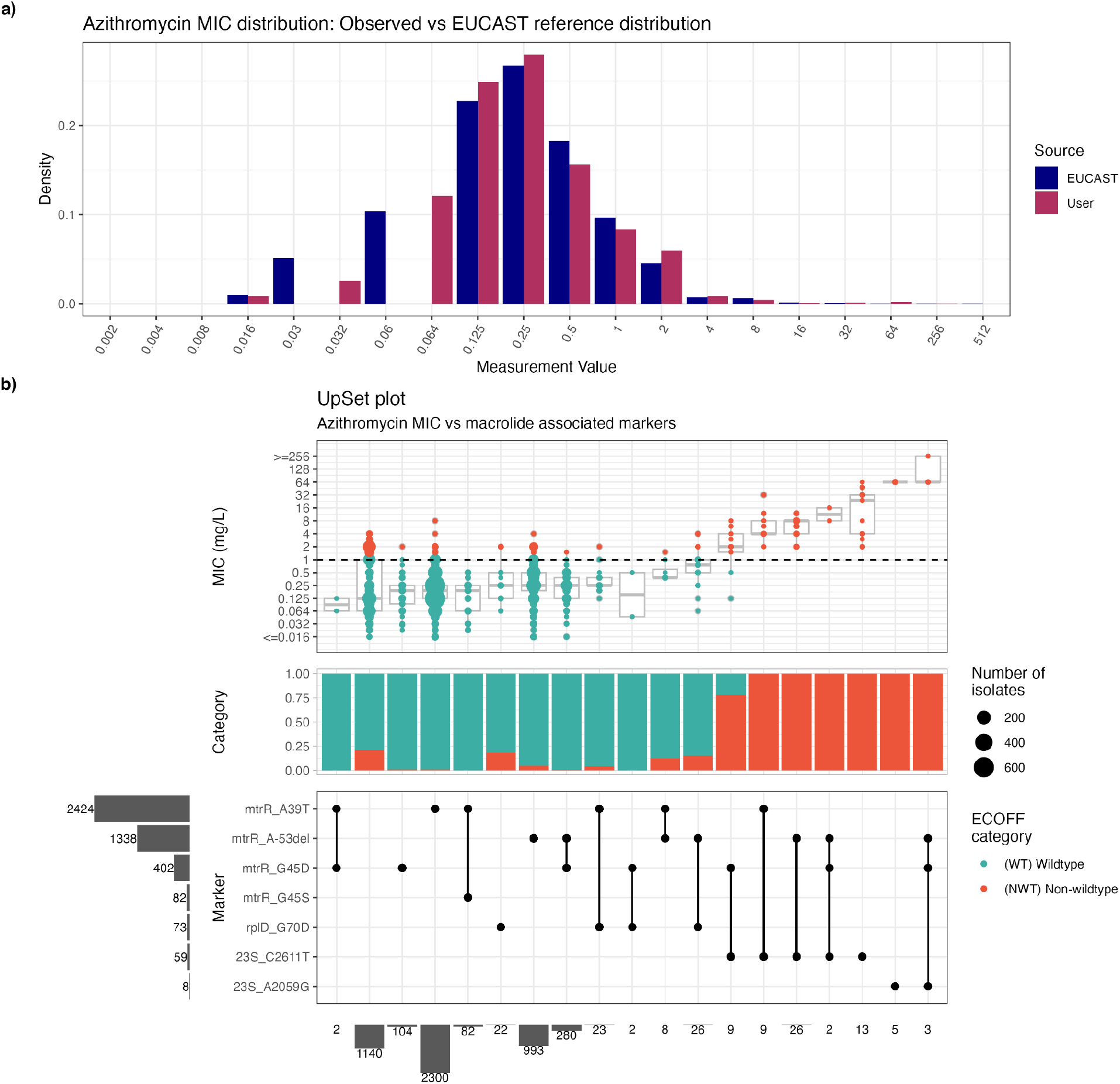
Decreased susceptibility to azithromycin in 5,055 *N. gonorrhoeae* isolates from the European Gonococcal Antimicrobial Surveillance Program (Euro-GASP). a) Distribution of minimum inhibitory concentrations (MIC) from this dataset (“User”) compared with the reference distribution (“EUCAST”), generated using the compare_mic_with_eucast() function. b) UpSet plot generated using the amr_upset() function, representing the distribution of MIC data (top panel) for each observed combination of genotypic markers (‘Marker’ grid panel) identified by AMRFinderPlus v4.0.23 (database v2025-03-25.1). The dashed horizontal line represents the epidemiological cut-off (ECOFF), automatically determined by the function. The intermediate ‘Category’ panel shows the proportion of samples with a phenotype categorised as wildtype (WT) or nonwildtype (NWT), for each combination of markers (i.e. mutations), as a stacked bar plot. Additional bar plots (in grey) show the size of each set: individual markers to the left of the marker grid, and combinations of markers below the marker grid.

### AMRgen data structures

Genotype data is stored in a long format data table, with each row representing a genetic marker detected in an input sample isolate. A genotype table may contain data from multiple samples, of any organism (one or many species), and would typically be created by importing a single multi-sample file of results from a genotyping tool such as *AMRFinderPlus* (see import functions below). Essential columns required by *AMRgen* analysis functions are sample name, genotypic marker, and drug class (individual drug is optional). Additional fields may be included (e.g. details of the genotype calls made by the bioinformatics software that generated them, or sample information such as species or source); such fields may be necessary for filtering to subsets of samples relevant to a specific analysis (e.g. those of a specific organism), but are not required by *AMRgen* functions.

Phenotype data is stored in a long format table, with each row representing an AST result for an input sample. A phenotype table may contain data from multiple samples, of any organism (one or many species), and would typically be created by importing a single multi-sample file of assay results exported from an AST instrument or LIMS, or sourced from a public database (see import functions below). Essential columns required by AMRgen analysis functions are sample name, drug (class ‘AMR::ab’, i.e. from *AMR* package functionality), and phenotype (S/I/R phenotype call, class ‘AMR::sir’). Where MIC or disk diffusion assay measurements are available, these can be included in columns of class ‘AMR::mic’ and ‘AMR::disk’ respectively (one or both may be provided). The phenotype column may be derived from these assay measurements using the *AMRgen* function ‘interpret_pheno()’, which uses the ‘AMR::as.sir()’ function to interpret each measurement against the relevant breakpoint (note that species and drug must also be provided, either as a single value applied to the whole table, or per row via additional data columns). Multiple phenotype columns may be included, representing interpretation against different breakpoints (e.g. interpret_pheno() stores its interpretations in columns labelled ‘pheno_eucast’, ‘pheno_clsi’, and/or ‘ecoff’). Some *AMRgen* functions explicitly accommodate two phenotype columns, one containing S/I/R calls against clinical breakpoints, and one containing WT/NWT calls against epidemiological cutoffs (ECOFF) which can be generated with ‘interpret_pheno()’ or ‘AMR::as.sir()’. Additional fields may be retained in the phenotype table (e.g. details of the assay used, sample sources, or species); such fields may be necessary for filtering to subsets of samples relevant to a specific analysis, but are not required by *AMRgen* functions (with the exception of interpret_pheno() which requires species information in order to identify relevant breakpoints, as described above). For example, the AST import/export functions (see below) store information on assay method (e.g. MIC or disk diffusion), AST platform (e.g. Vitek, Sensititre, Microscan), testing standard (e.g. CLSI, EUCAST), and dataset source (e.g. BioProject name, study name, PubMed ID) using standard field names ‘method’, ‘platform’, ‘guideline’, and ‘source’, respectively.

Combined genotype and phenotype data are stored in a wide format table, with each row representing a unique input sample. Columns indicate phenotypes for a single drug (R, I and/or NWT), and genotypes for markers relevant to the drug or drug class, encoded as binary variables (encoded as 1=presence or 0=absence for each phenotype or genotype). The table can be generated from an input genotype table and an input phenotype table using the function get_binary_matrix(), which must specify a single drug whose phenotypes are to be extracted and included. By default, all genotypic markers associated with this drug and its class will be extracted from the genotype table, however the user may override this and provide specific drug or class labels with which to filter the genotype markers. Additional columns indicating AST assay measurements (MICs and/or disk zones) or phenotype calls (against breakpoints and/or ECOFF) are required for some *AMRgen* functions, and can be optionally included when running get_binary_matrix(). An additional column indicating the combination of markers found in each sample, can be added to the table using the function get_combo_matrix().

### Analysing genotype/phenotype relationships

Functions are provided to calculate PPV for a specific drug, from a set of associated genotypic markers. These functions can simultaneously calculate PPV for different phenotype variables (all related to the same drug), defined as resistance (R vs S/I), non-susceptibility (R/I vs S), and/or nonwildtype phenotype (NWT vs WT, as determined by comparison to ECOFF) for the specified drug. These functions utilise a genotype-phenotype binary matrix (either supplied as input, or calculated from separate genotype and phenotype input tables), and output a table summarising the PPV for each phenotype variable, for each marker and marker combination observed amongst the input samples. PPV is calculated as a simple proportion (PPV_i,R_ = *x*_i,R_ / *n*_i_, PPV_i,NWT_ = *x*_i,NWT_ / *n*_i_, where *n*_i_ is the number of samples with the marker/combination, *x*_i,R_ and *x*_i,NWT_ are the number of such samples that are resistant or NWT). The PPV functions also output the numerator, denominator, and upper and lower values of the 95% confidence interval (CI) of these proportions. If assay measures are provided in the input data, these are also summarised in terms of median, geometric mean, and interquartile range associated with each marker/combination. The PPV functions also output plots summarising the phenotypes associated with each marker/combination in up to three plotting panels, chosen from (i) the point estimate and 95% CI of PPV for each phenotype, (ii) the categorical breakdown of phenotype values (S/I/R calls, as stacked bar plot), and (iii) the distribution of assay values (MICs or disk diffusion zone values, as boxplot) (e.g. **Figure 4a**). The primary function amr_ppv() includes data for all individual markers and marker combinations (optionally filtered to a minimum frequency threshold). The solo_ppv() function returns data only for markers found ‘solo’ in a genome without any other markers in the current binary matrix (which should include all markers associated with the target drug and/or drug class, as set by the user when generating this matrix). This is considered a strong indicator of the impact of an individual genetic marker on phenotype, although in some organisms the number of observations can be low due to genetic linkage between mobile AMR genes which results in some markers being very rarely observed in the absence of others^20^.

In the PPV plots, each row indicates a unique marker or combination of markers. Combinations can be labelled either as a list of marker names, or indicated in a grid of intersections in the form of an UpSet plot^18^. A complementary function amr_upset() is available to create an UpSet plot laid out horizontally, where each column indicates a marker combination and the plotting panels show the distribution of assay values and categorical breakdown of phenotype (e.g. **Figures 2b, 3a**). This orientation provides a more typical way of visualising the impact on phenotype of combinations of resistance markers, with the phenotypic assay values presented on the y-axis as a function of the presence of genotypic markers/combinations indicated on the x-axis^19,20^. However, the function utilises the same underlying calculations and plotting components as ppv(), and returns the same summary statistics.

The amr_logistic() function fits logistic regression models for resistance (R vs S/I) and/or nonwildtype phenotype (NWT vs WT, as determined by comparison to ECOFF), for a single drug and a set of associated markers. The function utilises a genotype-phenotype binary matrix (either supplied as input, or calculated from separate genotype and phenotype input tables as described above), and fits one model for each phenotype as outcome, with genotypic markers as independent covariates. By default only those markers present in at least 10 genomes are included in each model, as regression often fails when very rare markers are included; however this threshold can be adjusted by the user. To reduce the impact of separation which is quite likely with sparse marker data, the default is to fit models using Firth’s bias-reduced regression implemented in the *logistf* package; however users may instead opt to use the base *R* ‘glm()’ function. The amr_logistic() function outputs (i) the fitted models, (ii) tables summarising the effects estimated for each marker (estimate, 95% CI, and p-value), and (iii) a plot representing the results for each marker (e.g. **Figures 3b, 4b**).

**Figure 3:**
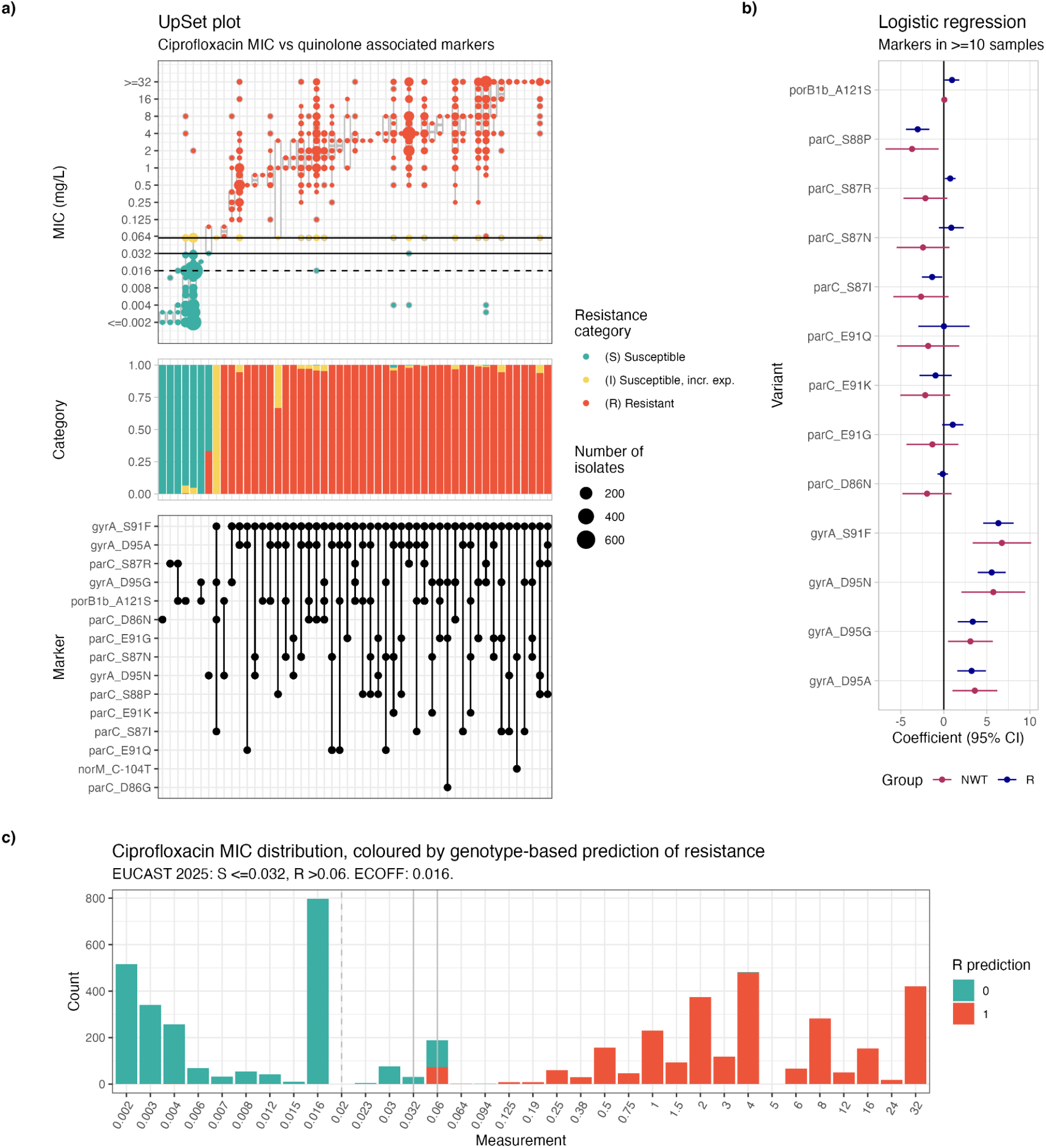
Resistance to ciprofloxacin in 5,360 *N. gonorrhoeae* isolates from the European Gonococcal Antimicrobial Surveillance Program (Euro-GASP). **(a)** Distribution of minimum inhibitory concentration (MIC) data (top panel) for each observed combination of genetic markers (grid panel) identified by AMRFinderPlus, visualised using amr_upset(). Horizontal lines represent EUCAST clinical breakpoints and the epidemiological cut-off (ECOFF, dashed line). **(b)** Coefficients and 95% confidence intervals (CI) representing the contribution of each marker to a Firth’s logistic regression model (one for resistance, R and one for nonwildtype, NWT as phenotypic outcome), and each individual marker as covariate using amr_logistic(). Only markers present in >=10 samples are included. **(c)** MIC distribution coloured by resistance prediction based on the logistic regression model, predictions were generated using the concordance() function and plotted using assay_by_var(). Vertical lines represent the EUCAST clinical breakpoints (solid) and ECOFF (dashed).

**Figure 4:**
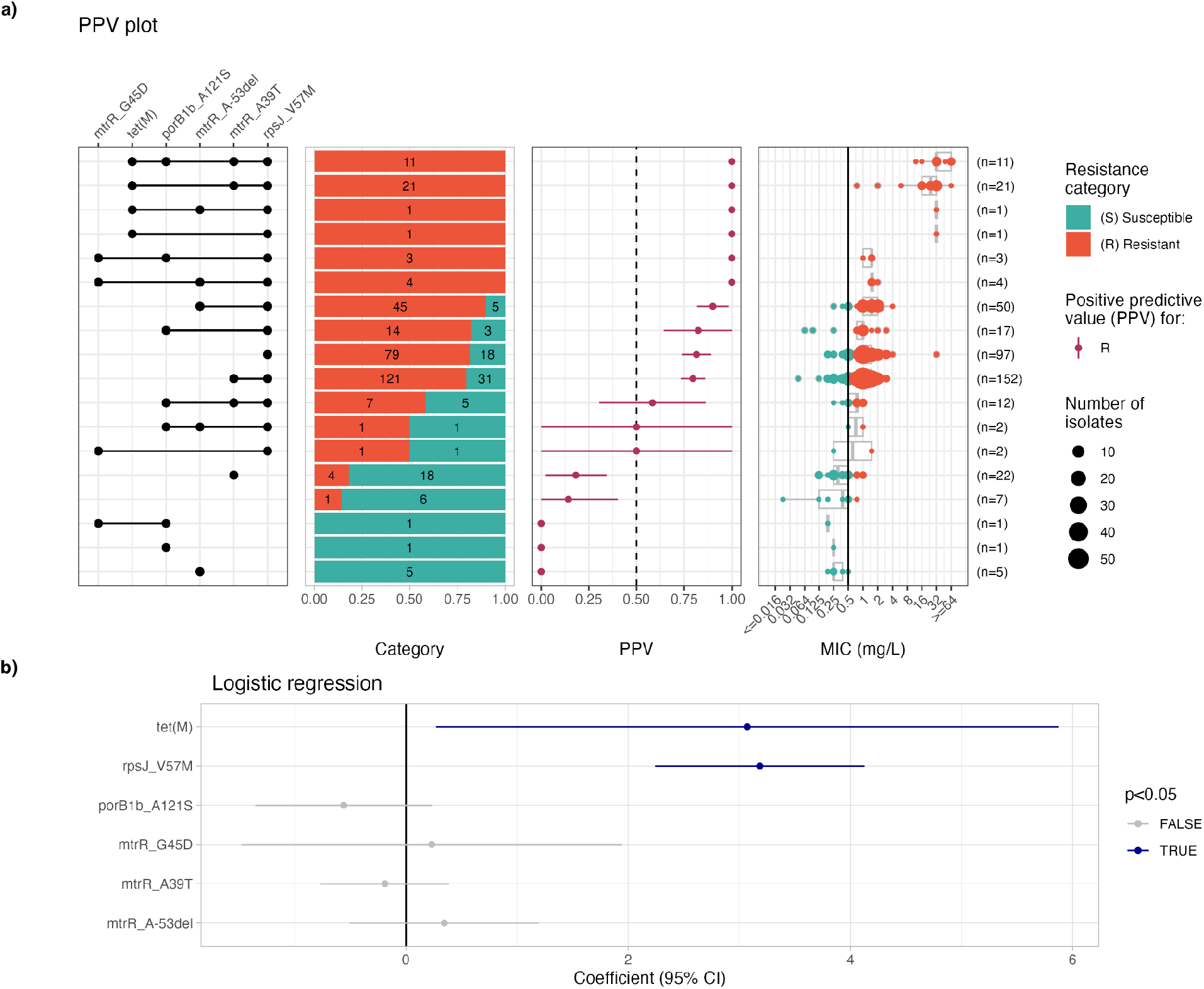
Resistance to tetracycline in 409 *N. gonorrhoeae* isolates from Eastern Spain. **(a)** Positive predictive values (PPV) for each observed combination of markers, plotted together with the distribution of minimum inhibitory concentrations (MIC) observed for each marker combination, generated using the ppv() function. In the central PPV panel, the vertical dashed line indicates PPV=0.50, to help distinguish PPV values >0.5 thus indicating a positive association with resistance (whereby isolates carrying the marker/s are more likely than not to be resistant). In the MIC panel, the vertical line indicates the resistance breakpoint (MIC >0.5 mg/L). **(b)** Results from the amr_logistic() function showing the coefficients and 95% confidence intervals (CI) for each individual marker in a Firth’s logistic regression model built using the resistance category as phenotypic outcome, and each individual marker as an independent covariate.

The concordance() function provides quantitative assessment of the concordance of phenotypes (resistance and/or NWT) for a specific drug with associated genotype-based predictions (which can be defined in various ways). The function utilises the standard genotype-phenotype binary matrix (either supplied as input, or calculated from separate genotype and phenotype input tables), and defines genotypic prediction variables by applying a genotypic prediction rule to the marker presence/absence data. Options for genotypic prediction rules include (i) presence of any marker, (ii) presence of all markers, (iii) presence of a minimum number of markers, (iv) logistic regression. Users may also supply a list of markers to include or exclude in the calculation of genotypic predictions; or thresholds for solo PPV or logistic regression p-values to filter markers prior to calculating predictions. This provides flexibility to explore concordance of observed phenotypes, with genotypic predictions obtained using a variety of different rules for inferring phenotypes from genotypes (see vignette: “Assessing geno-pheno concordance” in **Supplementary Information**). The output includes a list of included genotypic markers, the phenotype prediction for each sample, and a matrix of quantitative metrics. Standard metrics (sensitivity, specificity, PPV, negative predictive value, accuracy, kappa, F-measure) are calculated using the *yardstick* package^26^. AMR-specific error rates are calculated according to ISO 20776-2: (i) major error (ME), defined as the proportion of susceptible isolates incorrectly predicted resistant from genotype (equal to 1-specificity); and (ii) very major error (VME), defined as the proportion of truly resistant isolates not predicted as such from genotype (equal to 1-sensitivity).

### Visualisation

In addition to the plots generated by amr_ppv() and amr_upset() described above, *AMRgen* includes three generic plotting functions, to support visualising distributions of assay data (MIC/disk) or quantitative metrics associated with specific genotype markers. The function assay_by_var() can be used to plot the distribution of MIC or disk diffusion zone values, in either histogram or boxplot format, with samples coloured by one variable and faceted by another (see examples in **Figures 3c, 5a, 6a, 6b**). Taking a phenotype table as input, this can be used flexibly, for example to explore input phenotype data by checking the accuracy of S/I/R phenotype calls vs assay values (colouring by S/I/R call); or to explore the distribution of assay data aggregated from different sources (stratifying by assay method and colouring by source or laboratory). The function can also be used to explore how genotypes are distributed on the assay distribution, e.g. colouring samples by the presence of a particular genotype, or by the number of known markers detected (e.g. **Figure 6a, 6b**); or to see how categorical S/I/R calls predicted from genotypes using the concordance() function map to the assay distribution (**Figure 3c**). Optionally, breakpoints can be annotated on the plot; these may be supplied by the user, or retrieved from the *AMR* package by specifying a species and antibiotic to query. A set of functions are also provided to retrieve reference distributions from EUCAST, and plot these against user data (e.g. **Figure 2a**). Finally, the function plot_estimates() is provided to visualise quantitative metrics of association between genotype markers or combinations, and phenotypes, including e.g. PPV, or effects measured from logistic regression or other models (e.g. **Figures 3b, 4b**).

### Data import and export

User data can be imported into *AMRgen*’s standard genotype and phenotype table formats using the import_geno() and import_pheno() wrapper functions, which take as input either a file path or an existing R data frame. The import_pheno() functions support importing phenotype data obtained from different sources, including data exported from automated AST instruments (Vitek, Sensititre, Microscan, Phoenix, SIRscan), or data in formats used by WHOnet, NCBI or EBI databases. The function format_pheno() is also provided to help process data from non-standard formats. The import_pheno() functions can optionally re-interpret assay measurements against breakpoints and/or ECOFFs from CLSI and/or EUCAST using the *AMR* package. The import_geno() function provides direct support for importing results generated by the genotyping software *AMRFinderPlus*^16^, *CARD Resistance Gene Identifier*^6^, and *ABRicate* (using NCBI or *ResFinder*^27^ databases).

*AMRgen* includes functions to download public genotype or phenotype data directly from NCBI Pathogen Detection (https://www.ncbi.nlm.nih.gov/pathogens/) or the EBI AMR Portal (https://www.ebi.ac.uk/amr). Data can be retrieved from NCBI using either Google Cloud BigQuery (via the *bigrquery* package^28^, which requires an authorised Google Cloud account) or the NCBI EUtils API (via the *rentrez* package^29^, which does not require authorisation but is slower and retrieves AST data only). Data is retrieved from EBI by downloading parquet files from their FTP site using RCurl, which are then read using the *arrow* package^30^. Notably, the EBI AST database includes data from multiple other databases and individual research articles, retrieved and aggregated by the CABBAGE project^23^. Download functions include the option to reformat to standard AMRgen genotype or phenotype tables, and to re-interpret phenotype data against breakpoints and/or ECOFFs. Functions are also provided to write out AST data from the phenotype table format into files suitable for submission to the NCBI or EBI public sequence archives, to facilitate sharing of AST data associated with public whole genome sequence data linked by BioSample.

Functions are also supplied to retrieve reference distributions for MIC or disk diffusion zone data from the EUCAST website, https://mic.eucast.org/, using the *rvest* package^31^. Users must specify a single antibiotic, and optionally a specific organism (otherwise distributions for all organisms with available data will be retrieved). Helper functions are also provided to calculate and compare the distribution of user data with the reference distribution, to plot the reference distribution, or to plot a comparison of user and reference distributions.

### Example Use Cases

The *AMRgen* package includes example datasets, and Vignettes using these to demonstrate how to undertake genotype-phenotype analyses with *AMRgen* functions, covering several World Health Organization (WHO) priority AMR pathogens^32,33^. Here we use example datasets for *Neisseria gonorrhoeae* and *Klebsiella pneumoniae*, to illustrate how *AMRgen* can be used to investigate the genetic basis for clinically important resistance phenotypes identified through coordinated regional surveillance. We further highlight some examples from the package Vignettes, illustrating how additional genetic and/or source information can be incorporated into *AMRgen* visualisations to facilitate exploring real-world scenarios with WHO priority pathogens *Escherichia coli* and *Salmonella enterica*. All figures can be reproduced using the package Vignettes (also included here as **Supplementary Information**).

### Neisseria gonorrhoeae

Gonorrhoea is the second most commonly notified sexually transmitted infection in Europe^34^. Treatment is complicated by resistance to multiple antimicrobials^35^, and *N. gonorrhoeae* is classified by the WHO as a high priority AMR pathogen^33^. The European Gonococcal Antimicrobial Surveillance Programme (Euro-GASP), co-ordinated by the European Centre for Disease Prevention and Control (ECDC), has since 2013 included whole genome sequencing of a subset of surveillance isolates, enabling monitoring not only of resistance prevalence trends but the genetic mechanisms underlying these^11^. Here we use publicly available data from Euro-GASP^11,36,37^ and a recent regional surveillance study in Spain^38^, to demonstrate how AMRgen can be used to understand the contribution of known genetic variants to phenotypic resistance. AMR genotypes were identified from whole genome assemblies using *AMRFinderPlus*^16^ (v4.0.23, database version 2025-03-25.1), and the results imported into *R* using *AMRgen*’s import_amrfp() function. Phenotype data (MIC measurements) were sourced from the corresponding publications^11,36–38^. These data are included in the *AMRgen* package as example data, and the specific commands to repeat these analyses and generate the figures shown here are included in the *AMRgen* package vignette: “Example using large-scale regional/national surveillance data” (see **Supplementary Information**).

#### Use of the epidemiological cut-off (ECOFF) to study decreased susceptibility to azithromycin

Azithromycin was historically used in dual therapy for the treatment of gonorrhoea but has been progressively withdrawn from clinical guidelines due to the global expansion of resistant lineages^11,36,37^. As no EUCAST clinical breakpoint exists for azithromycin in *N. gonorrhoeae*, we demonstrate the use of *AMRgen* using the EUCAST epidemiological cut-off (ECOFF > 1 mg/L^39^) as the binary phenotypic threshold to distinguish wildtype (WT) from nonwildtype (NWT) isolates. The MIC distribution of azithromycin across the 5,055 Euro-GASP isolates with available phenotypic data was compared against the EUCAST reference distribution using *AMRgen*’s compare_mic_with_eucast() function (**Figure 2a**). This comparison allows users to contextualise their study population relative to the reference wildtype distribution published by EUCAST^39^, which is especially valuable when no clinical breakpoint is defined^2,21^.

The distribution of MIC values across observed marker combinations (**Figure 2b**) was explored using the amr_upset() function on the genotype-phenotype binary matrix built with get_binary_matrix(). By passing the species name to the UpSet function, the EUCAST ECOFF is automatically detected and represented. Of the seven genotypic markers detected by AMRFinderPlus and associated with macrolides (the drug class to which azithromycin belongs), the UpSet plot revealed that only mutations in the 23S rDNA gene^40,41^ (2059A>G or 2611C>T) were associated with MIC increases sufficient to cross the NWT threshold, whereas mutations in the *mtrR* gene or its promoter^10^, or the RplD G70D substitution^42^, did not reach this threshold individually or in combination (see **Figure 2b**). This was confirmed by using amr_logistic() to fit a logistic regression model using all seven markers, which showed statistically significant associations for 23S rDNA mutations only (see full results in Vignette). Concordance between the genotypic prediction (predicted using the logistic regression model) and the ECOFF classification was assessed using the concordance() function. The marker set achieved only a 17.6% sensitivity (very major error, VME = 82.3%), indicating that the majority of NWT isolates are not explained by the set of markers under study. This discrepancy can be attributed to mosaic variants in the genes encoding the MtrCDE efflux pump^43^, including MtrD mutations associated with decreased azithromycin susceptibility^44^ that have not yet been incorporated into the AMRFinderPlus database, highlighting the importance of the choice of genotyping software and databases and the need for ongoing expansion and curation of these resources^45^.

#### Ciprofloxacin genotype-phenotype analysis using logistic regression and concordance-based prediction

Ciprofloxacin resistance is highly prevalent (∼60-70%) in the global and European populations of *N. gonorrhoeae*^37,38,46^. Resistance is driven by known genetic determinants, primarily mutations in the *gyrA* and *parC* genes^47^. For the combined Euro-GASP dataset, containing MIC data for 5,360 strains, we demonstrate the full *AMRgen* analytical workflow for an antibiotic with an established EUCAST clinical breakpoint (S <= 0.032 mg/L; R > 0.06 mg/L), combining UpSet plot visualisation, logistic regression, and concordance analysis. For this antibiotic, the UpSet plot obtained with amr_upset() shows that the vast majority of isolates above the clinical breakpoint for resistance carry the S91F mutation in the *gyrA* gene (**Figure 3a**), the principal driver of ciprofloxacin resistance in *N. gonorrhoeae*. An ECOFF is also available for this antibiotic (ECOFF > 0.016 mg/L), and this is used for downstream analyses in parallel with the resistance breakpoint. A logistic regression model fit using amr_logistic() (using all markers identified in ≥ 10 isolates) confirmed that *gyrA* mutations are positively associated with both resistant (R) and nonwildtype (NWT) phenotype categorisations (**Figure 3b**). Analysis of concordance between logistic regression-based genotypic predictions and observed resistance phenotypes showed 99.85% sensitivity, 96.55% specificity, and a PPV of 97.09%, with a ME of 3.45% and no VME. Genotype-based resistance predictions were extracted from the results of the concordance() function and used to colour the distribution of ciprofloxacin MICs in the dataset (**Figure 3c**), demonstrating excellent agreement between genetic markers and phenotypic AST data for ciprofloxacin in this species.

#### Tetracycline resistance: implications for novel prevention strategies

We used *AMRgen* to characterise the genetic determinants of tetracycline resistance in a local population of *N. gonorrhoeae* strains collected in Eastern Spain between 2021-2024^38^. This use case is particularly relevant in the context of doxycycline (a tetracycline agent) post-exposure prophylaxis (doxy-PEP), a strategy for preventing sexually-transmitted infections whose long-term effectiveness for preventing gonorrhoea infections may be compromised by the selection of pre-existing resistant lineages^48^. PPVs were obtained for solo markers and marker combinations using the ppv() function, which can also output the distribution of available phenotypic data for each set (**Figure 4a**). As expected, tetracycline resistance is primarily explained by two genetic determinants: the chromosomal RpsJ V57M mutation and the plasmid-borne *tet(M)* gene. Prevalence analysis of the study population revealed that 91.2% of the population carried the RpsJ V57M substitution, while 8.3% carried *tet(M)*. A logistic regression analysis performed with the amr_logistic() function confirmed the independent significant contributions of both markers to tetracycline resistance (**Figure 4b**). The concordance() function was run with the results of the logistic regression as prediction rule. The marker set achieved 98.72% sensitivity and 82.84% PPV, but only 28.09% specificity, resulting in a high major error rate (ME = 71.91%). This pattern reflects the high prevalence of RpsJ V57M in isolates with elevated MICs that remain below the clinical breakpoint for resistance, meaning that many phenotypically susceptible isolates are genotypically predicted as resistant. Together, these findings suggest that doxy-PEP is likely to select for pre-existing resistant lineages, given that chromosomal resistance determinants are widespread and the *tet(M)* gene can be readily acquired by circulating susceptible lineages.

### Klebsiella pneumoniae

*K. pneumoniae* is the fastest growing cause of AMR healthcare-associated infections in Europe^49^, and carbapenem-resistant *K. pneumoniae* is classified by the WHO as a critical priority^33^. The European Survey of Carbapenemase-Producing Enterobacteriaceae (EuSCAPE) was undertaken to systematically determine the incidence and epidemiology of carbapenem-non-susceptible *K. pneumoniae* across the region^50^. This surveillance programme included whole genome sequencing of representative isolates of carbapenem susceptible and non-susceptible isolates, to identify the genetic mechanisms underlying resistance, and measurement of MIC to the carbapenem, meropenem^51^. Here we use publicly available data on 1,490 isolates from EuSCAPE to demonstrate how *AMRgen* can be used to understand the contribution of carbapenemases^52^ and outer-membrane porin mutations^9,53^ to phenotypic carbapenem resistance. AMR genotypes were identified from whole genome assemblies using Kleborate, which screens for both acquired carbapenemase genes and porin mutations (truncation of OmpK35, and resistance-associated mutations or truncation of OmpK36)^54^, and the results were imported using *AMRgen*’s import_kleborate() function. Phenotype data (meropenem MIC measurements) were sourced from the EuSCAPE publication^51^, imported and interpreted against 2025 EUCAST breakpoints using import_pheno(). These genotype and phenotype data are included in the *AMRgen* package as examples, and the specific commands to repeat these analyses and generate the figures shown here are included as a package vignette: “Exploring gene vs mutation combinations” (see **Supplementary Information**).

Carbapenemase genes were detected in 579 isolates, OmpK35 and/or OmpK36 mutations were detected in 579 isolates, and 386 isolates had both. In isolates lacking any of these markers, the median meropenem MIC was 0.064 mg/L, which as expected is below the ECOFF (0.125 mg/L) and thus in the wildtype part of the reference distribution. A standard UpSet plot (see vignette in **Supplementary Information**) is sufficient to see that most individual markers were associated with an increase in MIC, but only a subset of marker combinations were associated with median MIC above the clinical breakpoint (>8 mg/L). However it is quite difficult to tease out specific patterns from the UpSet visualisation due to the number of markers and combinations, and none of the options for ordering these (i.e. by their frequency, name, marker count, resistance PPV, median MIC) make it easy to see the interactions between carbapenemase genes vs porin mutations. To do this, we can instead use the flexible assay_by_var() function in *AMRgen* to visualise the MIC distribution as boxplots, grouped by porin mutation and faceted to show one panel per carbapenemase (**Figure 5a**). To further reduce complexity and increase clarity of the visualisation, we also grouped carbapenemases into their respective gene families (*bla*_IMP_, *bla*_KPC_, *bla*_NDM_, *bla*_OXA_, and *bla*_VIM_) and categorised porin mutations as either loss/truncation or β-strand loop 3 insertions. Note the median and geometric mean MIC values for each marker combination are also output by the assay_by_var() function, and the resulting table is visualised in **Figure 5b**.

**Figure 5:**
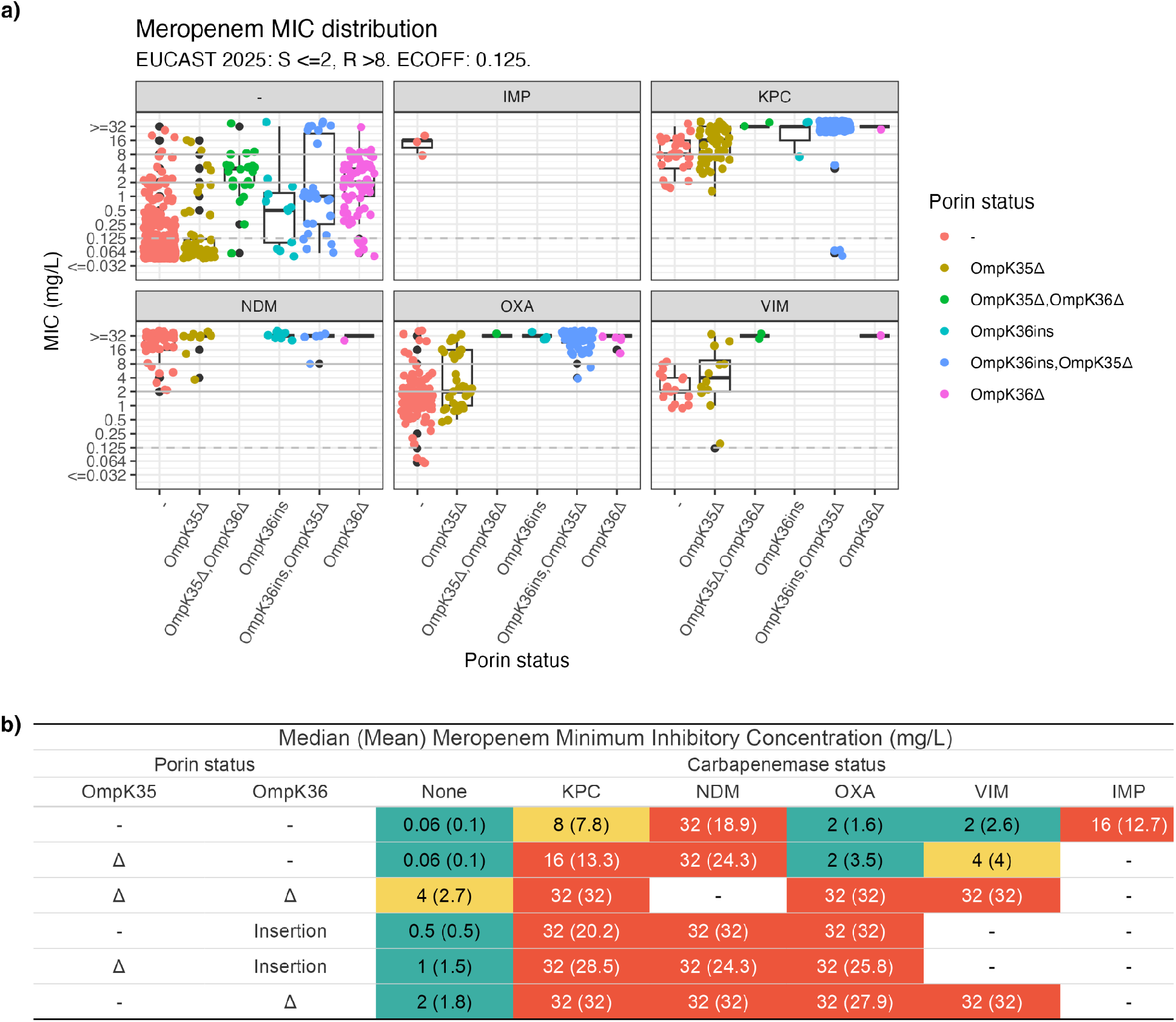
Meropenem resistance patterns in *Klebsiella pneumoniae* isolates (n = 1,490) and their associated carbapenem resistance determinants identified in the European Survey of Carbapenemase-Producing Enterobacteriaceae (EuSCAPE). **(a)** Distribution of meropenem minimum inhibitory concentration (MIC, mg/L) stratified by carbapenemase and porin status, generated using *AMRgen’*s assay_by_var() function. Each panel presents MIC data for the subset of isolates carrying the carbapenemase gene printed at the top of the panel (‘-’ indicates no carbapenemase detected). Within each panel, MIC distributions are shown for subsets of isolates grouped by the presence or absence (-) of mutations detected in the outer membrane porins OmpK35 or OmpK36. OmpK35Δ/36Δ indicate a missing or truncated porin gene, and OmpK36ins indicates an OmpK36 β-strand loop 3 (L3) insertion of amino acids GD, TD, or D. Isolates with multiple carbapenemases (n=11) or only CTX-M-33 carbapenemase (n=1) were excluded. Horizontal lines represent EUCAST S (≤2 mg/L) and R (>8 mg/L) clinical breakpoints (solid lines) and the epidemiological cut-off (ECOFF 0.125mg/L, dashed line). **(b)** Median (and geometric mean) of meropenem MIC values, for each combination of carbapenemase and porin mutation shown in the boxplots in panel **(a)**, output by the same assay_by_var() call that generated the boxplots. Cells in the table are coloured by the median meropenem MIC value based on the EUCAST clinical breakpoint interpretation (red indicates R, yellow I, green S).

Exploring the data in this way using *AMRgen* helps to illuminate a few key points regarding the effect of different carbapenemases, porin mutations, and their interactions, on meropenem MIC in *K. pneumoniae*. The upper-left panel makes it easier to see the effects of porin mutations on meropenem MIC in the absence of carbapenemase: no effect for OmpK35Δ (median MIC 0.064 mg/L), and increasing MIC impacts for OmpK36 insertions (median MIC 0.5 mg/L), OmpK36 insertion + OmpK35Δ (median MIC 1 mg/L), OmpK36Δ (median MIC 2 mg/L), and OmpK35Δ + OmpK36Δ (median MIC 4 mg/L) (**Figure 5a**). The other panels highlight that some carbapenemases are associated with much more significant increases in MIC, in the absence of porin mutations (red data points), particularly *bla*_NDM_, *bla*_IMP_ which were associated with MIC above the breakpoint (32 mg/L and 16 mg/L, respectively, see **Figure 5b**). These visualisations also show that OmpK35 loss has much less impact on MIC compared with OmpK36 loss or mutation, whichever carbapenemase is present. Notably, we can see that OmpK36 loss or mutation is associated with clinical resistance (MIC ≥32 mg/L) when any of these carbapenemases are present, whereas isolates with disrupted OmpK35 show much more variable MIC distributions that are driven by the specific carbapenemase present.

Borderline meropenem MICs, between the S (≤2 mg/L) and R (>8 mg/L) breakpoints, can be challenging to interpret in the clinical laboratory. According to EUCAST guidelines such values should be interpreted as “I, Susceptible, Increased exposure” meaning there is a high likelihood of therapeutic success when meropenem exposure is increased (e.g. by adjusting the dosing regimen or by its concentration at the site of infection). However these data clarify that such values are expected when *bla*_OXA_ or *bla*_VIM_ are present, and that for such strains the MIC could easily shift above the breakpoint with the evolution of one or two porin mutations in response to meropenem exposure (in particular with OmpK36 loss or truncation, which can in principle arise through a wide range of genetic mechanisms). Notably, EUCAST sets an alternative breakpoint of R >2 mg/L for meningitis, which would generally avoid using meropenem whenever any carbapenemase is present; CLSI sets the general meropenem breakpoint at this same value (R ≥4 mg/L) and advises to test for specific carbapenemases in addition to MIC to further guide therapy decisions^55^. Importantly, the *AMRgen* package enables both visualisations and quantitative analyses of these complex AMR genotype-phenotype relationships, facilitating a clearer understanding of the diverse combinatorial mechanisms underlying meropenem MIC patterns.

### Other examples

#### Exploring the phenotypic impact of gene deletion variants

Infections caused by extended-spectrum beta-lactamase (ESBL)-producing Enterobacterales are a critical global health threat, often leaving clinicians with few treatment options beyond last-resort carbapenems. In Malawi, first-line treatment for sepsis shifted from chloramphenicol to ceftriaxone in 2004, this was followed by a notable re-emergence of chloramphenicol susceptibility as its clinical use declined^56^. A recent study revealed that in *Escherichia coli*, this re-emergence is frequently driven by the stable degradation of resistance genes rather than their total loss from the population. Specifically, insertion sequence-mediated truncations in the plasmid-borne chloramphenicol resistance gene, *catB3*, were associated with reversion to a chloramphenicol-susceptible phenotype. Here we demonstrate how AMRgen can be used to explore the phenotypic effects of deletion variants in acquired AMR genes, using public *E. coli* data. We used a matched phenotype-genotype dataset from the Developing an Antimicrobial Strategy for Sepsis In Malawi (DASSIM) study^57^, and retrieved publicly available genotype and phenotype data from NCBI. These data are included in the *AMRgen* package as example datasets, and the specific commands to repeat these analyses and generate the figures shown here are included in the *AMRgen* package Vignette: “Analysing the impact of deletion variants on susceptibility”. **Figure 6a** shows the distribution of MIC values for isolates carrying the complete *catB3* gene (100% coverage to the *catB3* reference gene, as reported by AMRFinderPlus) and the truncated variant (≤70% coverage), generated using the *AMRgen* function assay_by_var(). The plot reveals the majority of isolates carrying a truncated version of the *catB3* gene are susceptible, with MICs below the CLSI breakpoint (≤8 µg/mL; note there are no EUCAST breakpoints). Experimental evolution reported by Graf *et al*^58^ suggests that these silencing events are highly stable under antibiotic pressure, reinforcing the potential for chloramphenicol to be reintroduced as a targeted reserve agent for ESBL Enterobacterales infections in low-resource settings.

**Figure 6:**
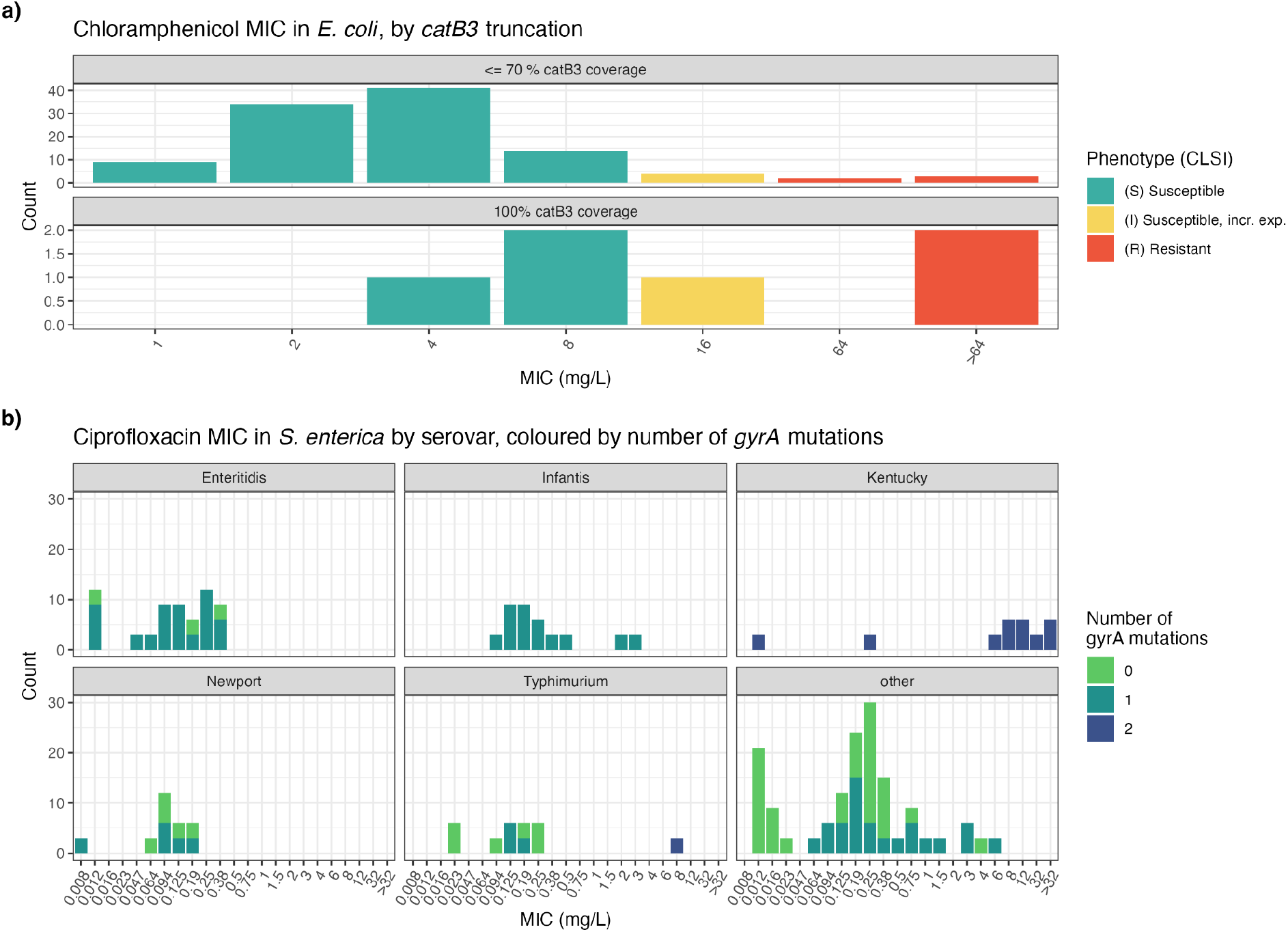
Examples exploring relationships between MIC and other variables using AMRgen plotting functions. **(a)** Exploring impacts of *catB3* gene truncations on chloramphenicol susceptibility phenotypes in *E. coli*, using public data. Panels shows the distribution of chloramphenicol MICs for isolates in which AMRFinderPlus reported partial (≤70% coverage) vs complete (100% coverage) hits to the *catB3* reference gene, created using the assay_by_var() function. Bars are coloured to indicate the S/I/R interpretation based on CLSI breakpoints (note that EUCAST does not define breakpoints for chloramphenicol in Enterobacterales). **(b)** Exploring variation in ciprofloxacin susceptibility in *Salmonella enterica*, by serovar and *gyrA* mutation count. Panels show the distribution of ciprofloxacin MICs for isolates of different serovars, coloured by the count of *gyrA* mutations, created using the assay_by_var() function.

#### Exploring the variation in ciprofloxacin resistance markers among *Salmonella* serovars

Globally, *Salmonella* is a leading cause of diarrhoeal disease, with fluoroquinolone-resistant non-typhoidal *Salmonella* listed as a high priority pathogen by the WHO^32,33^. Salmonellae are commensals and pathogens of a wide range of domestic and wild animals, and they can survive for extended periods in dry and aquatic environments. There are over 2,500 serovars of *Salmonella enterica* subspecies *enterica*, and these vary in their host specificity, disease severity, and AMR patterns. Here, we show how *AMRgen* can help explore variation in susceptibility to ciprofloxacin (the most widely prescribed, and susceptibility tested, fluoroquinolone agent) among non-typhoidal *Salmonella* serovars, using a small user-generated data set that is included in the *AMRgen* package. This anonymised data set, exported from a LIMS as a combined genotypic/phenotypic table with extra metadata columns for serovar and source, was first converted to *AMRgen*-compatible tables using a combination of *dplyr* functions and *AMRgen*’s format_pheno() function. By retaining the additional metadata columns, we can use the assay_by_var() function to examine the relationships between ciprofloxacin MIC, number of *gyrA* mutations, and serovar (**Figure 6b**). This highlights that isolates with zero or one *gyrA* mutations had overlapping MICs, and only those with two mutations had substantially increased MIC. Faceting by serovar further highlights that in this dataset, all *S*. Kentucky isolates had two *gyrA* mutations, all *S*. Infantis isolates had one *gyrA* mutation, and the other serovars had variable numbers of mutations. Specific commands to repeat these analyses and generate the figure shown here are in the AMRgen package Vignette: “Example with custom stratification by isolate source” (see **Supplementary Information**).

## Conclusions

*AMRgen* fills a key methodological gap, providing a reproducible and extensible foundation for AMR genotype-phenotype investigation within the *R* computing environment. Interoperable with a wide range of instruments and data platforms, and building on the foundations of the widely-used *AMR* package, this free and open-source software has broad applications across the research, clinical and public health domains.

## Data availability

Data used in the examples presented here are included as data objects in the AMRgen package, available at https://github.com/AMRverse/AMRgen.

## Code availability

The AMRgen package source code is available at https://github.com/AMRverse/AMRgen and is distributed under a GNU General Public License v3.0. The version described here is v0.1.0.9000 (DOI: 10.5281/zenodo.19947408). AMRgen documentation is available within the package, and at the AMRgen website https://amrgen.org which includes Vignettes showing how to reproduce all examples included in this manuscript.

## Acknowledgments

This work was supported by the Wellcome Trust [Grant number 226432/Z/22/Z to KEH]. DN was supported by the UK Health Security Agency (PhD funding competition). ZAD is funded by the LSHTM ITD Fellowship. LSB is funded by the Miguel Servet program (Instituto de Salud Carlos III, Spain, ref. CP23/00084), co-funded by the European Union through Fondo Social Europeo, and is part of the CSIC’s Global Health Platform (PTI Salud Global). GMK was supported by Medical Research Council UK (MR/W026643/1). JH was supported by an Emerging Leadership Fellowship from the National Health and Medical Research Council (NHMRC) of Australia (2034741). RNG is supported by funding from the Medical Research Council (MRC), Biotechnology and Biological Sciences Research Council (BBSRC) and Natural Environmental Research Council (NERC), which are all Councils of UK Research and Innovation (Grant number MR/W030578/1) under the umbrella of the JPIAMR - Joint Programming Initiative on Antimicrobial Resistance via the STRESST project, and from UKRI through the Strength in Places Fund (grant no. SIPF 36348), as part of the infection innovation Consortium (iiCON).

This work was supported in part by the National Center for Biotechnology Information of the National Library of Medicine (NLM) and the National Institute of Allergy and Infectious Diseases (NIAID), National Institutes of Health (NIH), and the Human Foods Program (HFP), Food and Drug Administration (FDA). The contributions of the NIH author are considered Works of the United States Government. The findings and conclusions presented in this paper are those of the author(s) and do not necessarily reflect the views of the NIH or the U.S. Department of Health and Human Services. For the purpose of open access, the author has applied a CC BY public copyright licence to any Author Accepted Manuscript version arising from this submission. We thank attendees of the hackathon held to kick-start development of the AMRgen package in January 2025 for their valuable discussion and early exploration of the idea. We also thank the teams who generated and made public the datasets used here as examples and included in the package (EURO-GASP, EuSCAPE, DASSIM).

## Author contributions

Conceptualization: KEH, NC, JH, MSB. Software and Validation: KEH, SA, CD, NC, ZAD, EFN, RNG, JH, GMK, DN, ABP, LSB, KKT, MSB. Writing-Original draft preparation: KEH, DLC, RNG, LSB, KKT, MSB. Writing-Review and editing: all authors.

