## Supplementary Information for "AMRgen: an R package for antimicrobial resistance genotype-phenotype analysis"

### R package Vignettes

The AMRgen R package (https://AMRgen.org) includes a set of vignettes to illustrate package functionality. These are available within the package, and in interactive form on the AMRgen website (https://amrgen.org/articles/).

This PDF file includes static versions of the vignettes (v1-beta).

**Index of Vignettes**


##### 1. Analysing Geno-Pheno Data

*Overview of available functions, and how they can be used to analyse genotypic and phenotypic data together.*

##### 2. Downloading Geno-Pheno Data

*Downloading genotype and phenotype data from public databases at EBI or NCBI.*

##### 3. Assessing Geno-Pheno Concordance

*How to assess concordance between genotypic predictions of phenotype (defined using logistic regression models or other interpretive rules), with observed phenotypes.*

##### 4. Large-scale surveillance data for *Neisseria gonorrhoeae*

##### 5. Example with multiple *Salmonella enterica* serovars

*Illustrating different ways to explore geno-pheno data by source, genotypic marker count, etc.*

##### 6. Analysing clindamycin resistance in *Staphylococcus aureus*

*Illustrating how to dig further into AMRFinderPlus genotype calls and explore how different types of hits relate differently to phenotype*

##### 7. Exploring *catB3* deletion variants and impact on chloramphenicol susceptibility in *Escherichia coli*

*Illustrating how to explore the impact of gene deletion variants on phenotypes*

##### 8. Analysing meropenem resistance in *Klebsiella pneumoniae*

*Exploring combined impacts of acquired genes vs mutations, and comparing genotype calls from Kleborate, CARD RGI, AMRFinderPlus*

### Analysing Geno-Pheno Data

### Introduction

`AMRgen` is a comprehensive R package designed to integrate antimicrobial resistance genotype and phenotype data. It provides tools to:

- Import AMR genotype data (e.g. from AMRFinderPlus, hAMRonization)
- Import AST phenotype data (e.g. public data from NCBI or EBI, or your own data in formats like Vitek or WHOnet)
- Conduct genotype-phenotype analyses to explore the impact of genotypic markers on phenotype, including via logistic regression, solo marker analysis, and upset plots
- Fetch MIC or disk zone reference distributions from EUCAST

This vignette walks through a basic workflow using example datasets included in the `AMRgen` package, and explains how to wrangle your own data files into the right formats to use the same workflow.

Start by loading the package:

```
library(AMRgen)
library(dplyr)
#>
#> Attaching package: 'dplyr'
#> The following objects are masked from 'package:stats':
#>
#>     filter, lag
#> The following objects are masked from 'package:base':
#>
#>     intersect, setdiff, setequal, union
```

#### 1. Genotype table

##### 1a. Importing genotype data to AMRgen's standard table format

The `import_amrfp()` function lets you load genotype data from AMRFinderPlus output files, and process it to generate an object with the key columns needed to work with the `AMRgen` package.

```
# Example AMRFinderPlus genotyping output (from AllTheBacteria project)
ecoli_geno_raw
#> # A tibble: 45,228 × 28
#>    Name      `Protein identifier` `Contig id`  Start   Stop Strand `Gene symbol`
#>    <chr>     <lgl>                <chr>        <dbl>  <dbl> <chr>  <chr>
#>  1 SAMN0317… NA                   SAMN031776…  74721  75851 -      blaEC
#>  2 SAMN0317… NA                   SAMN031776… 166214 169315 +      acrF
#>  3 SAMN0317… NA                   SAMN031776…  20678  22033 -      glpT_E448K
#>  4 SAMN0317… NA                   SAMN031776…    758   1969 -      floR
#>  5 SAMN0317… NA                   SAMN031776…   4440   5666 +      mdtM
#>  6 SAMN0317… NA                   SAMN031776…   3941   4798 +      blaTEM-1
#>  7 SAMN0317… NA                   SAMN031776…    142    954 +      sul2
```

```
#>  8 SAMN0317… NA                     SAMN031776…   1018   1818 +      aph(3'')-Ib
#>  9 SAMN0317… NA                     SAMN031776…   1821   2654 +      aph(6)-Id
#> 10 SAMN0317… NA                     SAMN031776…    788   1957 +      tet(A)
#> # ℹ 45,218 more rows
#> # ℹ 21 more variables: `Sequence name` <chr>, Scope <chr>,
#> #   `Element type` <chr>, `Element subtype` <chr>, Class <chr>, Subclass <chr>,
#> #   Method <chr>, `Target length` <dbl>, `Reference sequence length` <dbl>,
#> #   `% Coverage of reference sequence` <dbl>,
#> #   `% Identity to reference sequence` <dbl>, `Alignment length` <dbl>,
#> #   `Accession of closest sequence` <chr>, `Name of closest sequence` <chr>, …

# Load AMRFinderPlus output
ecoli_geno <- import_amrfp(
  input_table = ecoli_geno_raw, # (replace 'ecoli_geno_raw' with the filepath for any
        AMRFinderPlus output)
  sample_col = "Name",
  # you can optionally specify the below key column names if they differ in your dataframe
        to standard AMRFinderPlus outputs
  element_symbol_col = "Gene symbol", # or "Element symbol"
  element_type_col = "Element type", # or "Type"
  element_subtype_col = "Element subtype",
  method_col = "Method",
  node_col = "Hierarchy node",
  subclass_col = "Subclass",
  class_col = "Class"
)

# Check the format of the processed genotype table
head(ecoli_geno)
#> # A tibble: 6 × 37
#>   id          marker      gene  mutation drug  drug_class `variation type` node
#>   <chr>       <chr>       <chr> <chr>    <ab>  <chr>      <chr>            <chr>
#> 1 SAMN03177615 blaEC      blaEC <NA>     NA    Beta-lact… Gene presence d… blaEC
#> 2 SAMN03177615 acrF       acrF  <NA>     NA    Efflux     Gene presence d… acrF
#> 3 SAMN03177615 glpT_E448K glpT  Glu448L… FOS:… Phosphoni… Protein variant… glpT
#> 4 SAMN03177615 floR       floR  <NA>     CHL:… Phenicols  Gene presence d… floR
#> 5 SAMN03177615 floR       floR  <NA>     FLR:… Phenicols  Gene presence d… floR
#> 6 SAMN03177615 mdtM       mdtM  <NA>     NA    Efflux     Gene presence d… mdtM
#> # ℹ 29 more variables: marker.label <chr>, `Protein identifier` <lgl>,
#> #   `Contig id` <chr>, Start <dbl>, Stop <dbl>, Strand <chr>,
#> #   `Gene symbol` <chr>, `Sequence name` <chr>, Scope <chr>,
#> #   `Element type` <chr>, `Element subtype` <chr>, Class <chr>, Subclass <chr>,
#> #   Method <chr>, `Target length` <dbl>, `Reference sequence length` <dbl>,
#> #   `% Coverage of reference sequence` <dbl>,
#> #   `% Identity to reference sequence` <dbl>, `Alignment length` <dbl>, …
```

The genotype table has one row for each genetic marker detected in an input genome, i.e. one per strain/marker combination. This means your output table may end up with more columns than your input table, as markers conferring resistance to multiple drug classes will be expanded into several rows.

If your genotype data is not in AMRFinderPlus format, you can wrangle other input data files into the necessary format. If only your column names differ to standard AMRFinderPlus inputs, you can specify these using `element_symbol_col`, `element_type_col`, `element_subtype_col`, `method_col`, `node_col`, `subclass_col` and `class_col`.

The essential columns for a genotype table to work with downstream `AMRgen` functions are:

- `id`: character string giving the sample name, used to link to sample names in the phenotype file (this column can have a different name, in which case you'll need to make sure it is the first column in the dataframe OR pass its name to the functions using `geno_sample_col`)
- `marker`: character string giving the name of the genetic marker detected
- `drug_class`: character string giving the antibiotic class associated with this marker

NOTE: You should consider whether you have genomes with no AMR markers detected by genotyping, and how to make sure these are include in your analyses. E.g. AMRFinderPlus will output one row per genome/marker combination, but if you have a genome with no markers detected, there will be no row at all for that genome in the concatenated output file. If your species has core genes included in AMRFinderPlus this probably won't be a problem as you would expect some calls for every genome (e.g. AMRFinderPlus will report blaSHV, oqxA, oqxB, fosA in all Klebsiella pneumoniae genomes, so all input genomes will appear in the concatenated output file). An easy solution is to run a check to make sure that all genome names in your input dataset are represented in the genotype table, and if any are missing add empty rows for these using e.g. `tibble(Name=missing_samples) %>% bind_rows(genotype_table)`.

#### 1b. Summarising a genotype table

You can summarise the content of a genotype table using the inbuilt `summarise_geno()` function.

```
ecoli_geno_summary <- summarise_geno(ecoli_geno)

# Number of unique samples, markers, genes, drugs, classes, and variation types
ecoli_geno_summary$uniques
#> # A tibble: 6 × 2
#>   column         n_unique
#>   <chr>             <int>
#> 1 id                 5258
#> 2 marker              244
#> 3 drug                 35
#> 4 drug_class           26
#> 5 gene                196
#> 6 variation type        5

# Unique counts of samples, markers, genes, drugs, and classes - per variation type
ecoli_geno_summary$per_type
#> # A tibble: 5 × 6
#>   `variation type`                 id marker  drug drug_class  gene
#>   <chr>                         <int>  <int> <int>      <int> <int>
#> 1 Gene presence detected         5258    164    22         17   164
#> 2 Inactivating mutation detected  615     42    15         14    42
#> 3 Nucleotide variant detected      57      2     3          3     1
#> 4 Promoter variant detected        93      4     1          1     1
#> 5 Protein variant detected       4920     65    18         16    21
```

The `summarise_geno()` function also returns a list of drugs and classes represented in the table, and the associated number of unique markers, unique samples, and total hits for each drug/class. Ordering by sample count, we see the most common class is efflux… there are only 3 efflux-associated markers but there are 11,828 hits to these across 5,258 samples (i.e. all samples have at least one). Next most common are markers associated with beta-lactams… there are 22 different markers with 6,379 hits across 4,989 of our 5,258 samples.

```
ecoli_geno_summary$drugs
#> # A tibble: 44 × 6
#>    drug                            drug_name   drug_class markers samples  hits
#>    <ab>                            <chr>       <chr>        <int>   <int> <int>
#>  1 AMC: Amoxicillin/clavulanic acid Amoxicilli… Aminopeni…       2      57    57
#>  2 AMK: Amikacin                   Amikacin    Aminoglyc…       6     176   180
#>  3 AMP: Ampicillin                 Ampicillin  Aminopeni…       6     749   749
#>  4 APR: Apramycin                  Apramycin   Aminoglyc…       1      98    98
#>  5 ATM: Aztreonam                  Aztreonam   Monobacta…       2      39    39
#>  6 AZM: Azithromycin               Azithromyc… Macrolides       4     472   478
#>  7 BLM: Bleomycin                  Bleomycin   Glycopept…       2      40    40
#>  8 CHL: Chloramphenicol            Chloramphe… Phenicols       15    1121  1181
#>  9 CLI: Clindamycin                Clindamycin Lincosami…       1      26    26
#> 10 CLR: Clarithromycin             Clarithrom… Macrolides       1       1     1
#> # ℹ 34 more rows

# Order by sample count per drug/class
ecoli_geno_summary$drugs %>% arrange(-samples)
#> # A tibble: 44 × 6
#>    drug                 drug_name       drug_class         markers samples  hits
#>    <ab>                 <chr>           <chr>                <int>   <int> <int>
#>  1 NA                   <NA>            Efflux                   3    5258 11828
#>  2 NA                   <NA>            Beta-lactams            22    4989  6379
#>  3 FOS: Fosfomycin      Fosfomycin      Phosphonics              9    4861  6411
#>  4 COL: Colistin        Colistin        Polymyxins              11    3415  3436
#>  5 NA                   <NA>            Tetracyclines           13    2634  2929
#>  6 NA                   <NA>            Quinolones              45    1822  4497
#>  7 STR1: Streptomycin   Streptomycin    Aminoglycosides         13    1669  3291
#>  8 SSS: Sulfonamide     Sulfonamide     Sulfonamides             5    1500  1876
#>  9 CHL: Chloramphenicol Chloramphenicol Phenicols               15    1121  1181
#> 10 NA                   <NA>            Cephalosporins (3…      32    1065  1285
#> # ℹ 34 more rows
```

Ordering by marker count, we see the classes with the most different markers associated are quinolones (45 unique markers detected across 1,822 unique samples), third generation cephalosporins (32 unique markers detected across 1,065 unique samples) and beta-lactams (22 unique markers detected across 4,989 unique samples).

```
ecoli_geno_summary$drugs %>% arrange(-markers)
#> # A tibble: 44 × 6
#>    drug                 drug_name       drug_class         markers samples  hits
#>    <ab>                 <chr>           <chr>                <int>   <int> <int>
#>  1 NA                   <NA>            Quinolones              45    1822  4497
#>  2 NA                   <NA>            Cephalosporins (3…      32    1065  1285
#>  3 NA                   <NA>            Beta-lactams            22    4989  6379
#>  4 NA                   <NA>            Trimethoprims           16     824   877
#>  5 CHL: Chloramphenicol Chloramphenicol Phenicols               15    1121  1181
#>  6 GEN: Gentamicin      Gentamicin      Aminoglycosides         14     687   704
#>  7 STR1: Streptomycin   Streptomycin    Aminoglycosides         13    1669  3291
#>  8 NA                   <NA>            Carbapenems             13      62    64
#>  9 NA                   <NA>            Tetracyclines           13    2634  2929
#> 10 COL: Colistin        Colistin        Polymyxins              11    3415  3436
#> # ℹ 34 more rows
```

The summarise_geno() function also returns a list of markers represented in the table, annotated with the associated drugs/classes and variation types. The column 'n' indicates the count of hits detected per marker. Ordering by this column, we see the most common markers are acrF, blaEC and glpT_E448K.

```
ecoli_geno_summary$markers %>% arrange(-n)
#> # A tibble: 349 × 6
#>    marker     drug           drug_name drug_class   `variation type`       n
#>    <chr>      <ab>           <chr>     <chr>        <chr>              <int>
#>  1 acrF       NA             <NA>      Efflux       Gene presence dete…  5002
#>  2 blaEC      NA             <NA>      Beta-lactams Gene presence dete…  4749
#>  3 glpT_E448K FOS: Fosfomycin Fosfomycin Phosphonics Protein variant de…  4731
#>  4 mdtM       NA             <NA>      Efflux       Gene presence dete…  3675
#>  5 emrD       NA             <NA>      Efflux       Gene presence dete…  2914
#>  6 pmrB_E123D COL: Colistin  Colistin  Polymyxins   Protein variant de…  1873
#>  7 pmrB_Y358N COL: Colistin  Colistin  Polymyxins   Protein variant de…  1531
#>  8 blaTEM-1   NA             <NA>      Beta-lactams Gene presence dete…  1279
#>  9 uhpT_E350Q FOS: Fosfomycin Fosfomycin Phosphonics Protein variant de…  1145
#> 10 tet(A)     NA             <NA>      Tetracyclines Gene presence dete…  1087
#> # ℹ 339 more rows
```

Filtering specifically for markers associated with quinolones, we can find out more about the 45 markers for this class that were found in the dataset. Summarising by variation type, we see there are 12 markers indicating detection of an acquired gene, 2 indicating inactivating mutaions, and 31 indicating protein mutations. Sorting by marker frequency, we can see the full list of 45 unique markers and that the most common is a protein variant in gyrA, gyrA_S83L. Filtering to "Gene presence detected" we can that the most common acquired gene was aac(6')-Ib-cr5.

```
# Count the different types of variants found
ecoli_geno_summary$markers %>%
  filter(drug_class == "Quinolones") %>%
  count(`variation type`)
#> # A tibble: 3 × 2
#>   `variation type`                 n
#>   <chr>                        <int>
#> 1 Gene presence detected          12
#> 2 Inactivating mutation detected   2
#> 3 Protein variant detected        31

# Sort by marker frequency to see the most common markers
ecoli_geno_summary$markers %>%
  filter(drug_class == "Quinolones") %>%
  arrange(-n)
#> # A tibble: 45 × 6
#>    marker         drug drug_name drug_class `variation type`             n
#>    <chr>          <ab> <chr>     <chr>      <chr>                    <int>
#>  1 gyrA_S83L      NA   <NA>      Quinolones Protein variant detected   855
#>  2 marR_S3N       NA   <NA>      Quinolones Protein variant detected   726
#>  3 parC_S80I      NA   <NA>      Quinolones Protein variant detected   639
#>  4 gyrA_D87N      NA   <NA>      Quinolones Protein variant detected   622
#>  5 parE_I529L     NA   <NA>      Quinolones Protein variant detected   442
#>  6 parC_E84V      NA   <NA>      Quinolones Protein variant detected   294
#>  7 aac(6')-Ib-cr5 NA   <NA>      Quinolones Gene presence detected     153
#>  8 parE_D475E     NA   <NA>      Quinolones Protein variant detected   147
```

```
#>  9 parE_L416F     NA    <NA>      Quinolones Protein variant detected   134
#> 10 parE_S458A     NA    <NA>      Quinolones Protein variant detected   111
#> # ℹ 35 more rows

# Filter to acquired genes and sort by frequency, to see the most common acquired genes
ecoli_geno_summary$markers %>%
  filter(drug_class == "Quinolones" & `variation type` == "Gene presence detected") %>%
  arrange(-n)
#> # A tibble: 12 × 6
#>    marker         drug drug_name drug_class `variation type`           n
#>    <chr>          <ab> <chr>     <chr>      <chr>                  <int>
#>  1 aac(6')-Ib-cr5 NA   <NA>      Quinolones Gene presence detected   153
#>  2 qnrS1          NA   <NA>      Quinolones Gene presence detected    61
#>  3 qnrB19         NA   <NA>      Quinolones Gene presence detected    36
#>  4 qnrB4          NA   <NA>      Quinolones Gene presence detected    11
#>  5 qnrB1          NA   <NA>      Quinolones Gene presence detected     3
#>  6 qnrB2          NA   <NA>      Quinolones Gene presence detected     3
#>  7 qepA1          NA   <NA>      Quinolones Gene presence detected     2
#>  8 qnrA1          NA   <NA>      Quinolones Gene presence detected     2
#>  9 qnrB6          NA   <NA>      Quinolones Gene presence detected     2
#> 10 qnrS2          NA   <NA>      Quinolones Gene presence detected     2
#> 11 qnrB           NA   <NA>      Quinolones Gene presence detected     1
#> 12 qnrB7          NA   <NA>      Quinolones Gene presence detected     1
```

### 2. Phenotype table

#### 2a. Importing phenotype data to AMRgen's standard table format

The `import_pheno()` function imports phenotype data from NCBI or other standard formats.

```
# Example E. coli phenotype data from NCBI
# This one has already been imported and phenotypes interpreted from assay data
ecoli_pheno
#> # A tibble: 4,168 × 11
#>    id           drug         mic  disk pheno_clsi ecoff guideline method platform
#>    <chr>        <ab>       <mic> <dsk> <sir>      <sir> <chr>     <chr>  <chr>
#>  1 SAMN11638310 CIP: Ci… 256.00    NA R          NWT   CLSI      broth… <NA>
#>  2 SAMN05729964 CIP: Ci…  64.00    NA R          NWT   CLSI      Etest  Etest
#>  3 SAMN10620111 CIP: Ci… >=4.00    NA R          NWT   CLSI      broth… <NA>
#>  4 SAMN10620168 CIP: Ci… >=4.00    NA R          NWT   CLSI      broth… <NA>
#>  5 SAMN10620104 CIP: Ci… <=0.25    NA S          NI    CLSI      broth… <NA>
#>  6 SAMN10620102 CIP: Ci… >=4.00    NA R          NWT   CLSI      broth… <NA>
#>  7 SAMN10620129 CIP: Ci… >=4.00    NA R          NWT   CLSI      broth… <NA>
#>  8 SAMN10620121 CIP: Ci… >=4.00    NA R          NWT   CLSI      broth… <NA>
#>  9 SAMN10620086 CIP: Ci… >=4.00    NA R          NWT   CLSI      broth… <NA>
#> 10 SAMN04122821 CIP: Ci…   1.00    NA R          NWT   CLSI      broth… Vitek
#> # ℹ 4,158 more rows
#> # ℹ 2 more variables: pheno_provided <sir>, spp_pheno <mo>

head(ecoli_pheno)
#> # A tibble: 6 × 11
#>   id           drug          mic  disk pheno_clsi ecoff guideline method platform
```

```
#>   <chr>        <ab>       <mic> <dsk> <sir>      <sir> <chr>     <chr>  <chr>
#> 1 SAMN11638310 CIP: Cip… 256.00    NA R          NWT   CLSI      broth… <NA>
#> 2 SAMN05729964 CIP: Cip…  64.00    NA R          NWT   CLSI      Etest  Etest
#> 3 SAMN10620111 CIP: Cip… >=4.00    NA R          NWT   CLSI      broth… <NA>
#> 4 SAMN10620168 CIP: Cip… >=4.00    NA R          NWT   CLSI      broth… <NA>
#> 5 SAMN10620104 CIP: Cip… <=0.25    NA S          NI    CLSI      broth… <NA>
#> 6 SAMN10620102 CIP: Cip… >=4.00    NA R          NWT   CLSI      broth… <NA>
#> # ℹ 2 more variables: pheno_provided <sir>, spp_pheno <mo>
```

```
# You can make your own from different file formats, and interpret against breakpoints,
        using:
#    import_pheno("filepath/NCBI_pheno.tsv", format="ncbi", interpret_clsi=T)
#    import_pheno("filepath/Vitek_pheno.tsv", format="vitek", interpret_eucast=T)
```

Data can be imported from various standard formats using the `import_pheno` function, and re-interpreted using latest breakpoints and/or ECOFF. Use `?import_pheno` to see the available formats and other options.

If your assay data is not in a standard format, you can wrangle other input data files into the necessary format, manually and/or with the help of the `format_pheno` function.

```
?import_pheno
```

```
?format_pheno
```

The phenotype table is long form, with one row for each assay measurement, i.e. one per strain/drug combination.

The essential columns for a phenotype table to work with `AMRgen` functions are:

- `id`: character string giving the sample name, used to link to sample names in the genotype file (this column can have a different name, in which case you'll need to make sure it is the first column in the dataframe OR pass its name to the functions using `pheno_sample_col`)
- `spp_pheno`: species in the form of an AMR package `mo` class (can be created from a column with species name as string, using `AMR::as.mo(species_string)`)
- `drug`: antibiotic name in the form of an AMR package `ab` class (can be created from a column with antibiotic name as string, using `AMR::as.ab(antibiotic_string)`)
- a phenotype column, e.g. the import functions output fields `pheno_eucast`, `pheno_clsi`, `pheno_provided`, `ecoff`: S/I/R phenotype calls in the form of an AMR package `sir` class (can be created from a column with phenotype values as string, using `AMR::as(sir_string)`, or generated by interpreting MIC or disk assay data using `AMR::as.sir`)

If you want to do analyses with raw assay data (e.g. upset plots) you will need that data in one or both of:

- `mic`: MIC in the form of an AMR package `mic` class (can be created from a column with assay values as string, using `AMR::as.mic(mic_string)`)
- `disk`: disk diffusion zone diameter in the form of an AMR package `disk` class (can be created from a column with assay values as string, using `AMR::as.disk(disk_string)`)

The import functions also standardise names for the following common fields:

- `method`: The laboratory testing method (e.g., "MIC", "disk diffusion", "Etest", "agar dilution")
- `platform`: The laboratory testing platform/instrument if relevant (e.g., "Vitek", "Phoenix", "Sensititre").

- `guideline`: The testing standard recorded in the input file as being used to make the provided phenotype interpretations (e.g. "CLSI", "EUCAST")
- `source`: An identifier for the dataset from which each data point was sourced (e.g. study or hospital name, pubmed ID, bioproject accession).

#### 2b. Summarising a phenotype table

You can summarise the content of a phenotype table using the inbuilt `summarise_pheno()` function.

```
ecoli_pheno_summary <- summarise_pheno(ecoli_pheno, pheno_cols = c("pheno_clsi",
        "pheno_provided", "ecoff"))

# Number of samples, drugs, species, and methods included in phenotype table
ecoli_pheno_summary$uniques
#> # A tibble: 6 × 2
#>   column    n_unique
#>   <chr>        <int>
#> 1 id            4164
#> 2 drug             1
#> 3 spp_pheno        1
#> 4 method           2
#> 5 platform         8
#> 6 guideline        1
```

The `summarise_pheno()` function returns a list of drugs and species represented in the table, and the associated number of samples with MIC measures, disk measures, both, or neither (S/I/R calls only).

```
# Number of samples with measurements from MIC vs disk vs both or neither, per bug-drug
        combination
ecoli_pheno_summary$drugs
#> # A tibble: 1 × 4
#>   drug                 drug_name     spp_pheno          mic
#>   <ab>                 <chr>         <chr>            <int>
#> 1 CIP: Ciprofloxacin Ciprofloxacin Escherichia coli  4168

# Number of samples with measurements from different methods, platforms, and guidelines
ecoli_pheno_summary$details
#> # A tibble: 8 × 7
#>   drug                 drug_name     spp_pheno     method platform guideline   mic
#>   <ab>                 <chr>         <chr>         <chr>  <chr>    <chr>     <int>
#> 1 CIP: Ciprofloxacin Ciprofloxacin Escherichia … Etest  Etest    CLSI          1
#> 2 CIP: Ciprofloxacin Ciprofloxacin Escherichia … broth… Microsc… CLSI          2
#> 3 CIP: Ciprofloxacin Ciprofloxacin Escherichia … broth… Phoenix  CLSI        483
#> 4 CIP: Ciprofloxacin Ciprofloxacin Escherichia … broth… Phoenix… CLSI          1
#> 5 CIP: Ciprofloxacin Ciprofloxacin Escherichia … broth… Sensiti… CLSI         59
#> 6 CIP: Ciprofloxacin Ciprofloxacin Escherichia … broth… Sensiti… CLSI       2708
#> 7 CIP: Ciprofloxacin Ciprofloxacin Escherichia … broth… Vitek    CLSI        502
#> 8 CIP: Ciprofloxacin Ciprofloxacin Escherichia … broth… <NA>     CLSI        412
```

The `summarise_pheno()` can also summarise, for each categorical phenotype column, the number in each category (S/I/R for interpretation against breakpoints, or NWT/WT for interpretation against ECOFF). This is helpful to explore whether your data set has sufficient numbers of R vs S, or NWT vs NWT, to be informative for downstream analyses of genotypes.

```
ecoli_pheno_summary$pheno_counts_list
#> $pheno_clsi
#> # A tibble: 1 × 6
#>   drug               drug_name     spp_pheno            S     I     R
#>   <ab>               <chr>         <chr>            <int> <int> <int>
#> 1 CIP: Ciprofloxacin Ciprofloxacin Escherichia coli  3011    63  1094
#>
#> $pheno_provided
#> # A tibble: 1 × 7
#>   drug               drug_name     spp_pheno            S     R    NI  `NA`
#>   <ab>               <chr>         <chr>            <int> <int> <int> <int>
#> 1 CIP: Ciprofloxacin Ciprofloxacin Escherichia coli  3113   970    37    48
#>
#> $ecoff
#> # A tibble: 1 × 6
#>   drug               drug_name     spp_pheno           NI    WT   NWT
#>   <ab>               <chr>         <chr>            <int> <int> <int>
#> 1 CIP: Ciprofloxacin Ciprofloxacin Escherichia coli   170  2768  1230
```

#### 3. Plot phenotype data distribution

It is always a good idea to check the distribution of raw AST data that we have to work with. The function `assay_by_var()` can be used to plot the distribution of MIC or disk measurements, coloured by a variable.

```
# Example E. coli AST data from NCBI

# Plot MIC distribution, coloured by CLSI S/I/R call
assay_by_var(pheno_table = ecoli_pheno, pheno_drug = "Ciprofloxacin", measure = "mic",
        colour_by = "pheno_clsi")
```

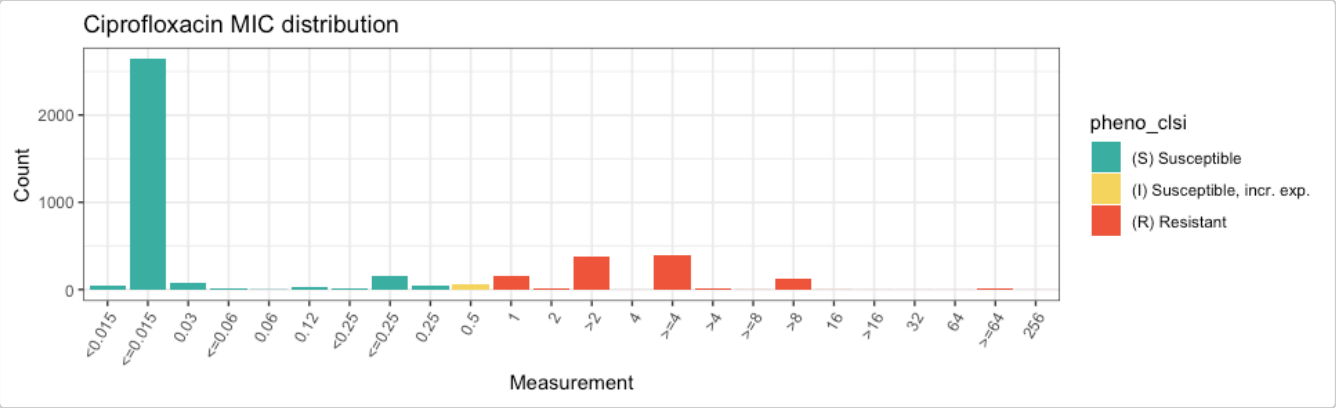


It's a good idea to make sure that the `SIR` field in the input data file has been interpreted correctly against the breakpoints. The AMRgen function `checkBreakpoints()` can be used to help look up breakpoints in the `AMR` package. Or, if you provide the function `assay_by_var()` with a species and guideline, it can look up the breakpoints and ECOFF and annotate these directly on the plot.

```
# Look up breakpoints recorded in the AMR package
checkBreakpoints(species = "E. coli", guide = "CLSI 2025", antibiotic = "Ciprofloxacin",
        assay = "MIC")
#>   MIC breakpoints determined using AMR package: S <= 0.25 and R > 1
#> $breakpoint_S
```

```
#> [1] 0.25
#>
#> $breakpoint_R
#> [1] 1
#>
#> $bp_standard
#> [1] "-"

# Specify species and guideline, to annotate with CLSI breakpoints
assay_by_var(pheno_table = ecoli_pheno, pheno_drug = "Ciprofloxacin", measure = "mic",
        colour_by = "pheno_clsi", species = "E. coli", guideline = "CLSI 2025")
#>   MIC breakpoints determined using AMR package: S <= 0.25 and R > 1
```

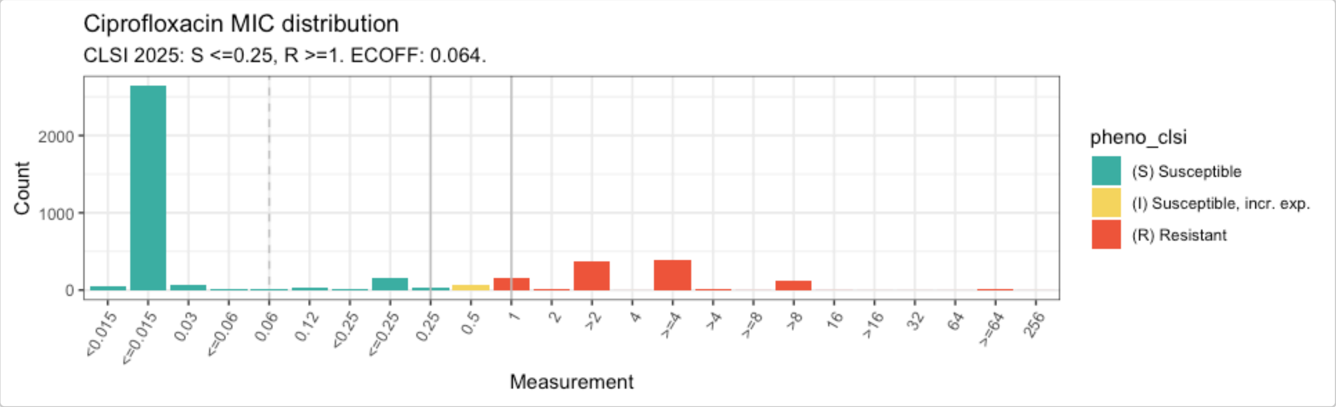


When aggregating AST data from different methods and sources, it is a good idea to check the distributions broken down by method or source. This can be done easily by passing the `assay_by_var()` function a variable name to facet by, which means a separate distribution will be plotted for each value of that variable (e.g. each type of 'method' in our AST test data). Note that this public data from NCBI includes non-standard values in the platform (e.g., `"Sensititre"` / `"Sensititer"`) in the platform.

```
# specify facet_by="method" to generate facet plots by assay method
mic_by_platform <- assay_by_var(pheno_table = ecoli_pheno, pheno_drug = "Ciprofloxacin",
        measure = "mic", colour_by = "pheno_clsi", species = "E. coli", guideline = "CLSI
        2025", facet_by = "method")
#>   MIC breakpoints determined using AMR package: S <= 0.25 and R > 1

mic_by_platform
```

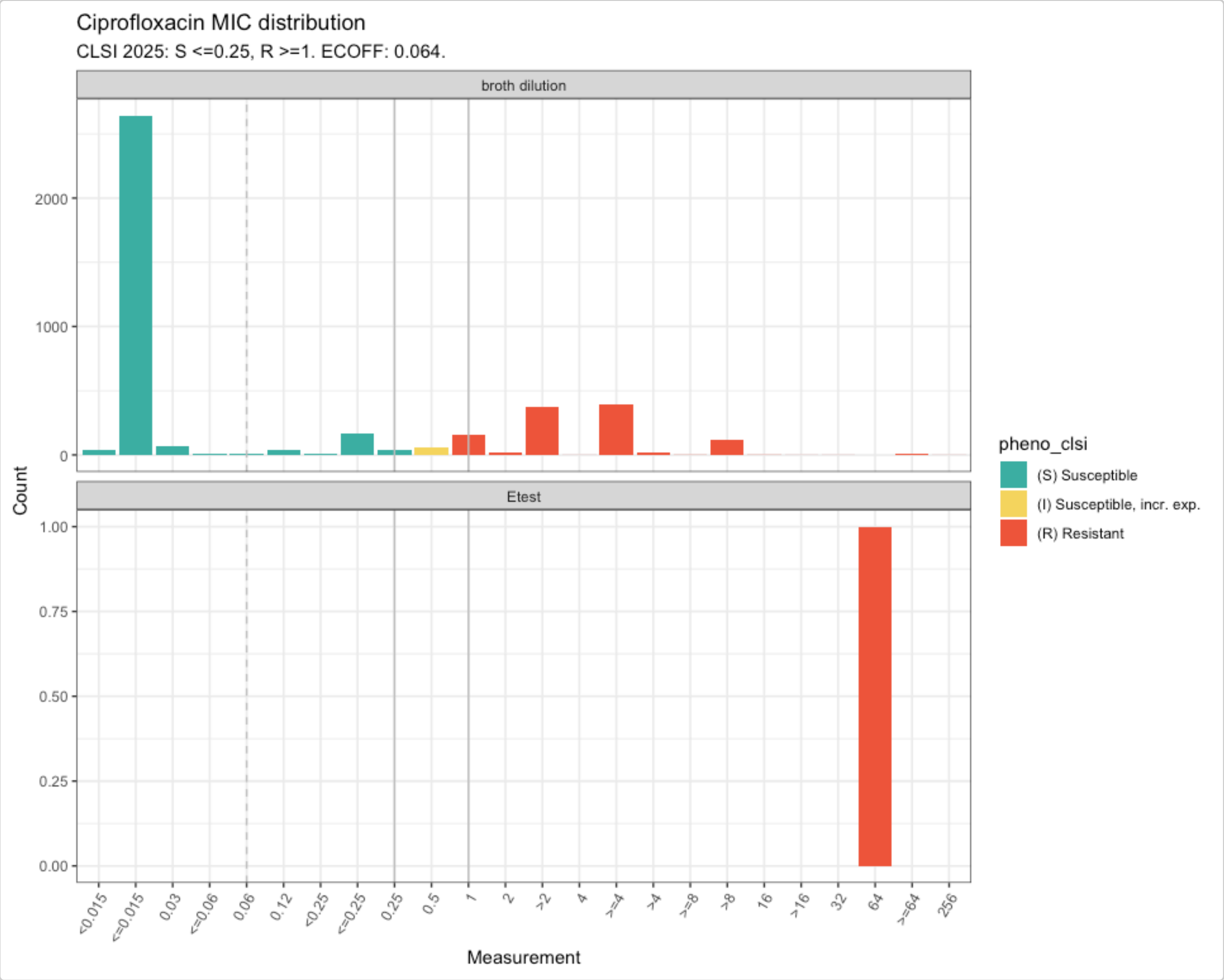


#### 4. Download reference assay distributions and compare to your data

It can also be helpful to check how your MIC or disk zone distribution compares to the reference distributions, to get a sense of whether your assays were calibrated correctly or if there may be some issues with a given dataset. AMRgen has functions to download the latest reference distributions from EUCAST (mic.eucast.org), and plot them on their own or with your data overlaid.

```
# get MIC distribution for ciprofloxacin, for all organisms
cip_mic_data <- get_eucast_mic_distribution("cipro")

# specify microorganism to only get results for that pathogen
ecoli_cip_mic_data <- get_eucast_mic_distribution("cipro", "E. coli")

# get disk diffusion data instead
ecoli_cip_disk_data <- get_eucast_disk_distribution("cipro", "E. coli")


# Ciprofloxacin MIC reference distribution for E. coli
ecoli_cip_mic_data
#> # A tibble: 19 × 2
#>          mic count
#>        <mic> <int>
#>  1   0.002    14
```

```
#>   2   0.004   189
#>   3   0.008  3952
#>   4   0.016  7238
#>   5   0.030  1355
#>   6   0.060   356
#>   7   0.125   401
#>   8   0.250   521
#>   9   0.500   171
#> 10   1.000    94
#> 11   2.000    47
#> 12   4.000   119
#> 13   8.000   246
#> 14  16.000   229
#> 15  32.000   564
#> 16  64.000   166
#> 17 128.000    85
#> 18 256.000    59
#> 19 512.000     7
```

```
# Compare reference distribution to example E. coli data
ecoli_cip <- ecoli_pheno$mic[ecoli_pheno$drug == "CIP"]

ecoli_cip_vs_ref <- compare_mic_with_eucast(ecoli_cip, ab = "cipro", mo = "E. coli")


ecoli_cip_vs_ref
#> # A tibble: 32 × 3
#>    value    user eucast
#>  * <fct>   <int>  <int>
#>  1 0.002       0     14
#>  2 0.004       0    189
#>  3 0.008       0   3952
#>  4 <0.015     41      0
#>  5 <=0.015  2642      0
#>  6 0.016       0   7238
#>  7 0.03       69   1355
#>  8 <=0.06     11      0
#>  9 0.06        5    356
#> 10 0.12       34      0
#> # ℹ 22 more rows
#> Use ggplot2::autoplot() on this output to visualise.

ggplot2::autoplot(ecoli_cip_vs_ref)
```

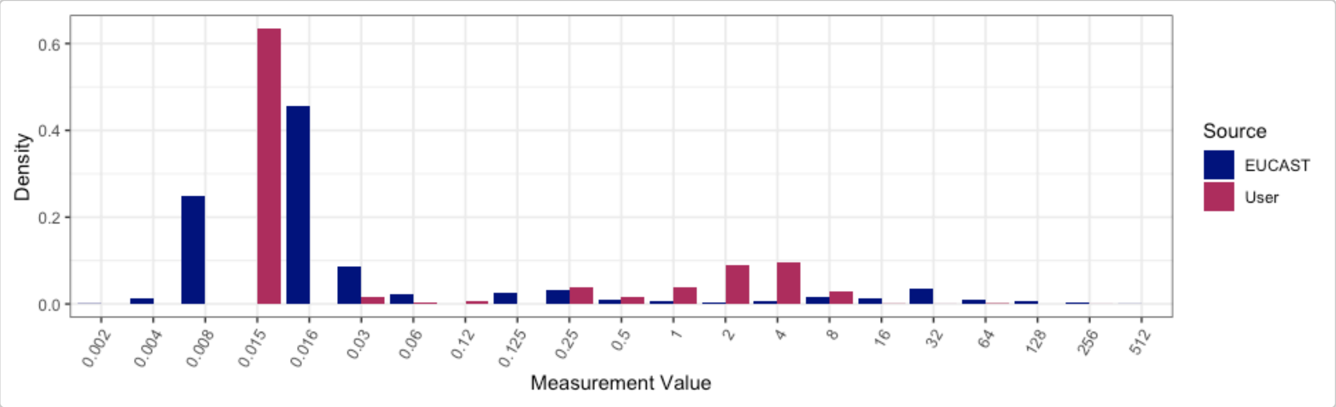


#### 5. Summarise the intersection of a genotype table and a phenotype table

You can summarise the intersecting content of a genotype table and a phenotype table using the inbuilt `summarise_geno_pheno()` function.

```
ecoli_geno_pheno <- summarise_geno_pheno(ecoli_geno,
  ecoli_pheno,
  pheno_cols = c("pheno_clsi", "ecoff")
)

# Total number of samples that appear in both tables, i.e. that have both
# genotype and phenotype data available
ecoli_geno_pheno$overlapping_samples
#> [1] 3629

# Table of drugs encountered in the phenotype table, indicating the
# number of samples that have phenotype data for this drug and also
# appear in the genotype table
ecoli_geno_pheno$drugs_with_pheno
#> # A tibble: 2 × 6
#>   drug                  n drug_class       drug_name     spp_pheno           mic
#>   <ab>              <int> <chr>            <chr>         <chr>             <int>
#> 1 CIP: Ciprofloxacin  3629 Fluoroquinolones Ciprofloxacin Escherichia coli  4168
#> 2 CIP: Ciprofloxacin  3629 Quinolones       Ciprofloxacin Escherichia coli  4168

# List of tables, one for each phenotype column in the input, indicating
# the number in each category (WT/NWT, S/I/R), amongst samples that also
# appear in the genotype table.
ecoli_geno_pheno$pheno_counts_list
#> $ecoff
#> # A tibble: 1 × 6
#>   drug               drug_name     spp_pheno           NI    WT   NWT
#>   <ab>               <chr>         <chr>            <int> <int> <int>
#> 1 CIP: Ciprofloxacin Ciprofloxacin Escherichia coli   170  2768  1230
#>
#> $pheno_clsi
#> # A tibble: 1 × 6
#>   drug               drug_name     spp_pheno            S     I     R
#>   <ab>               <chr>         <chr>            <int> <int> <int>
#> 1 CIP: Ciprofloxacin Ciprofloxacin Escherichia coli  3011    63  1094

# Number of markers encountered for each drug/class in the genotype table,
```

```r
# amongst samples that have phenotype data for the relevant drug/class
ecoli_geno_pheno$geno_hits
#> # A tibble: 1 × 6
#>   drug drug_name drug_class markers samples  hits
#>   <ab> <chr>     <chr>        <int>   <int> <int>
#> 1 NA   <NA>      Quinolones      44    1039  3618

# Frequency of each marker in the genotype table, amongst samples that have
# phenotype data for the relevant drug/class
ecoli_geno_pheno$geno_markers
#> # A tibble: 44 × 6
#>    marker         drug drug_name drug_class `variation type`                n
#>    <chr>          <ab> <chr>     <chr>      <chr>                       <int>
#>  1 aac(6')-Ib-cr  NA   <NA>      Quinolones Inactivating mutation detected     1
#>  2 aac(6')-Ib-cr5 NA   <NA>      Quinolones Gene presence detected           150
#>  3 acrR_R45C      NA   <NA>      Quinolones Protein variant detected           1
#>  4 gyrA_D87G      NA   <NA>      Quinolones Protein variant detected           3
#>  5 gyrA_D87N      NA   <NA>      Quinolones Protein variant detected         611
#>  6 gyrA_D87Y      NA   <NA>      Quinolones Protein variant detected          16
#>  7 gyrA_S83A      NA   <NA>      Quinolones Protein variant detected           3
#>  8 gyrA_S83L      NA   <NA>      Quinolones Protein variant detected         782
#>  9 gyrA_S83W      NA   <NA>      Quinolones Protein variant detected           1
#> 10 marR_R77C      NA   <NA>      Quinolones Protein variant detected           1
#> # ℹ 34 more rows
```

#### 6. Combine genotype and phenotype data for a given drug

The genotype and phenotype tables can include data related to many different drugs, but we need to analyse things one drug at a time. The function `get_binary_matrix()` can be used to extract phenotype data for a specified drug, and genotype data for markers associated with a specified drug class. It returns a single dataframe with one row per strain, for the subset of strains that appear in both the genotype and phenotype input tables. Each row indicates, for one strain, both the phenotypes (with SIR column, any assay columns if desired, and boolean 1/0 coding of R and NWT status) and the genotypes (one column per marker, with boolean 1/0 coding of marker presence/absence).

This binary matrix can be used as the starting a lot of downstream analyses, discussed below.

```r
# Get matrix combining phenotype data for ciprofloxacin, binary calls for R/NWT phenotype,
#    and genotype presence/absence data for all markers associated with the relevant drug
#    class (which are labelled "Quinolones" in AMRFinderPlus).
cip_bin <- get_binary_matrix(
  ecoli_geno,
  ecoli_pheno,
  pheno_drug = "Ciprofloxacin",
  geno_class = "Quinolones",
  sir_col = "pheno_clsi",
  keep_assay_values = TRUE,
  keep_assay_values_from = "mic"
)
#>  Defining NWT in binary matrix using ecoff column provided: ecoff

# check format
head(cip_bin)
```

```
#> # A tibble: 6 × 50
#>   id      pheno ecoff     mic     R   NWT gyrA_S83L gyrA_D87Y gyrA_D87N parC_S80I
#>   <chr>   <sir> <sir>   <mic> <dbl> <dbl>     <dbl>     <dbl>     <dbl>     <dbl>
#> 1 SAMN0… S     WT    <=0.015     0     0         0         0         0         0
#> 2 SAMN0… S     WT    <=0.015     0     0         0         0         0         0
#> 3 SAMN0… S     WT    <=0.015     0     0         0         0         0         0
#> 4 SAMN0… S     NWT     0.250     0     1         1         0         0         0
#> 5 SAMN0… S     NWT     0.120     0     1         0         1         0         0
#> 6 SAMN0… S     WT    <=0.015     0     0         0         0         0         0
#> # ℹ 40 more variables: parE_S458A <dbl>, parC_S80R <dbl>, parE_L416F <dbl>,
#> #   qnrB6 <dbl>, gyrA_D87G <dbl>, parC_S57T <dbl>, parC_E84A <dbl>,
#> #   soxS_A12S <dbl>, qnrB2 <dbl>, qnrS2 <dbl>, parC_E84K <dbl>,
#> #   parC_A56T <dbl>, qnrB19 <dbl>, `aac(6')-Ib-cr5` <dbl>, parC_E84V <dbl>,
#> #   parE_I529L <dbl>, parE_S458T <dbl>, parE_E460D <dbl>, parC_E84G <dbl>,
#> #   qnrS1 <dbl>, marR_S3N <dbl>, `aac(6')-Ib-cr` <dbl>, soxR_R20H <dbl>,
#> #   qnrB1 <dbl>, parE_I355T <dbl>, soxR_G121D <dbl>, qnrB4 <dbl>, qepA <dbl>, …

# list colnames, to see full list of quinolone markers included
colnames(cip_bin)
#>  [1] "id"          "pheno"       "ecoff"       "mic"
#>  [5] "R"           "NWT"         "gyrA_S83L"   "gyrA_D87Y"
#>  [9] "gyrA_D87N"   "parC_S80I"   "parE_S458A"  "parC_S80R"
#> [13] "parE_L416F"  "qnrB6"       "gyrA_D87G"   "parC_S57T"
#> [17] "parC_E84A"   "soxS_A12S"   "qnrB2"       "qnrS2"
#> [21] "parC_E84K"   "parC_A56T"   "qnrB19"      "aac(6')-Ib-cr5"
#> [25] "parC_E84V"   "parE_I529L"  "parE_S458T"  "parE_E460D"
#> [29] "parC_E84G"   "qnrS1"       "marR_S3N"    "aac(6')-Ib-cr"
#> [33] "soxR_R20H"   "qnrB1"       "parE_I355T"  "soxR_G121D"
#> [37] "qnrB4"       "qepA"        "gyrA_S83A"   "qnrA1"
#> [41] "parE_D475E"  "parC_A108V"  "qepA1"       "parE_E460K"
#> [45] "gyrA_S83W"   "marR_R77C"   "parE_L445H"  "parE_I464F"
#> [49] "qnrB"        "acrR_R45C"
```

For example, we can use it as input to `assay_by_var` to plot the assay distribution coloured by presence of a particular genetic marker

```
assay_by_var(cip_bin, measure = "mic", colour_by = "parC_S80I", pheno_drug =
        "Ciprofloxacin")
#> Warning in assay_by_var(cip_bin, measure = "mic", colour_by = "parC_S80I", : Column
        'drug' not found in phenotype table, so can't filter to the specified pheno_drug.
#> Ensure your input table is already filtered to the relevant drug.
```

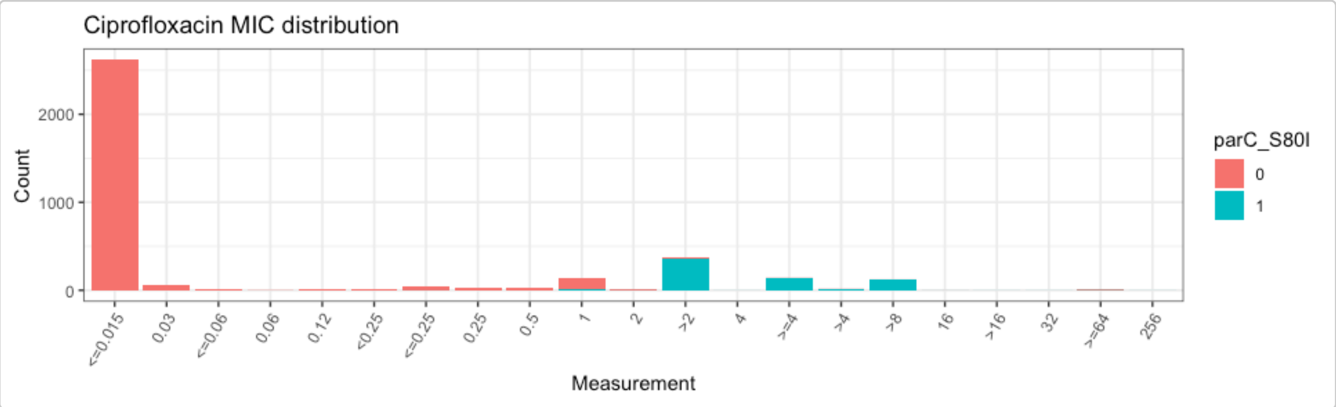

```r
# count the number of gyrA mutations per genome
gyrA_mut <- cip_bin %>%
  dplyr::mutate(gyrA_mut = rowSums(across(contains("gyrA_") & where(is.numeric)), na.rm =
        T)) %>%
  select(mic, gyrA_mut)

# plot the MIC distribution, coloured by count of gyrA mutations
mic_by_gyrA_count <- assay_by_var(gyrA_mut, measure = "mic", colour_by = "gyrA_mut",
        colour_legend_label = "No. gyrA mutations", pheno_drug = "Ciprofloxacin")
#> Warning in assay_by_var(gyrA_mut, measure = "mic", colour_by = "gyrA_mut", : Column
        'drug' not found in phenotype table, so can't filter to the specified pheno_drug.
#> Ensure your input table is already filtered to the relevant drug.

mic_by_gyrA_count
```

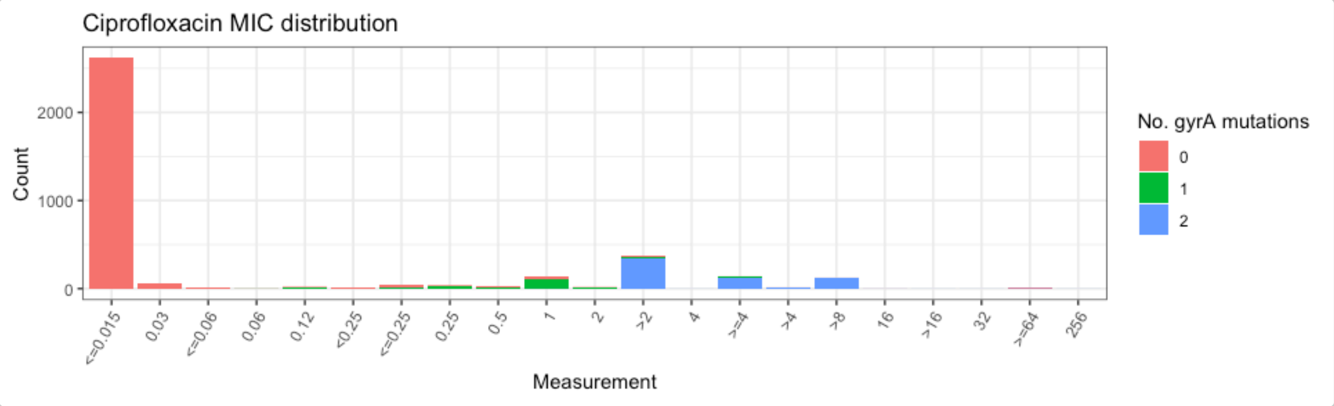


```r
# count the number of genetic determinants per genome
marker_count <- cip_bin %>%
  mutate(marker_count = rowSums(across(where(is.numeric) & !any_of(c("R", "NWT"))), na.rm
        = T)) %>%
  select(mic, marker_count)

# plot the MIC distribution, coloured by count of associated genetic markers
mic_by_marker_count <- assay_by_var(marker_count, measure = "mic", colour_by =
        "marker_count", colour_legend_label = "No. markers detected", pheno_drug =
        "Ciprofloxacin", colours = viridisLite::viridis(max(marker_count$marker_count) +
        1))
#> Warning in assay_by_var(marker_count, measure = "mic", colour_by = "marker_count", :
        Column 'drug' not found in phenotype table, so can't filter to the specified
        pheno_drug.
#> Ensure your input table is already filtered to the relevant drug.

mic_by_marker_count
```

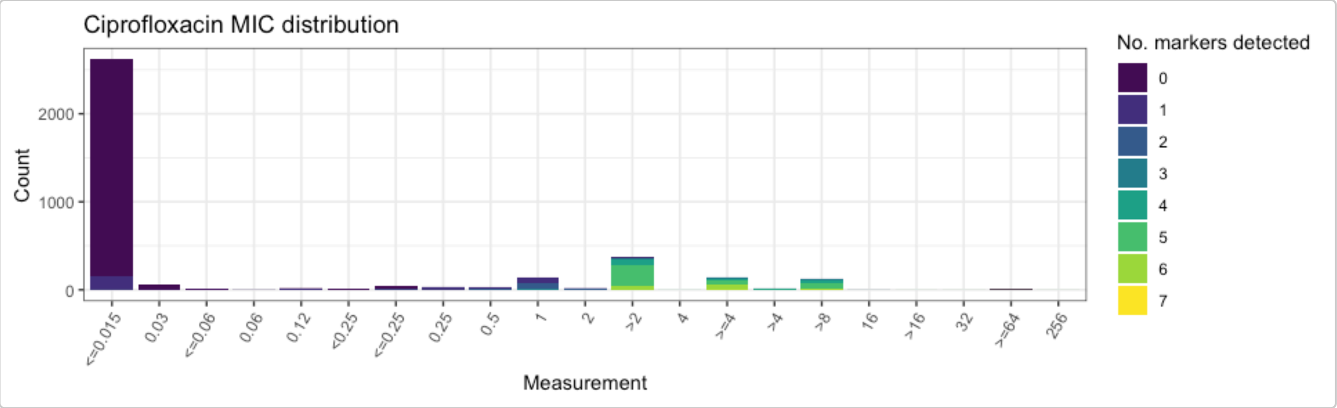


### 7. Model a binary drug phenotype using genetic marker presence/absence data

Logistic regression models can be informative to get an overview of the association between a drug resistance phenotype, and each marker thought to be associated with the relevant drug class.

The `amr_logistic()` function uses the `get_binary_matrix` function to generate binary-coded genotype and phenotype data for a specified drug and class; and fits two logistic regression models of the form `R ~ marker1 + marker2 + marker3 + ...` and `NWT ~ marker1 + marker2 + marker3 + ...`.

Note that the 'NWT' variable in the latter model can be taken either from a precomputed ECOFF-based call of WT=wildtype/NWT=nonwildtype (encoded in the input column `ecoff_col`), or computed from the S/I/R phenotype as NWT=R/I and WT=S.

The `amr_logistic()` function can fit the model using either the standard logistic regression approach implemented in the `glm()` function, or Firth's bias-reduced penalised-likelihood logistic regression implemented in the `logistf` package. The default is to use Firth's regression, as standard logistic regression can fail if there are too few observations in some subgroups, which happens quite often with this kind of data. To use `glm()` instead, set `glm=TRUE`.

The function also filters out markers with too few observations in the combined genotype/phenotype dataset. The default minimum is 10 but this can be changed using the `maf` parameter (maf stands for 'minor allele frequency'). If you are having trouble fitting models, it may be because too many markers and combinations have very few observations, and you might try increasing the `maf` value to ensure that rare markers are excluded prior to model fitting.

Using this modelling approach, a negative association with a single marker and phenotype call of R and NWT is a strong indication that marker does not contribute to resistance. Note however that a positive association between a marker and R or NWT does not necessarily imply the marker is independently contributing to the resistance phenotype, as there may be non-independence between markers that is not adequately adjusted for by the model.

The function returns 4 objects:

- `modelR`, `modelNWT`: data frames summarising each model, with beta coefficient, lower and upper values of 95% confidence intervals, and p-value for each marker (generated from the raw model output using `logistf_details()` or `glm_details()` as relevant)
- `plot`: a ggplot2 object generated from the `modelR` and `modelNWT` objects using the `compare_estimates()` function
- `bin_mat`: the binary matrix used as input to the regression models

```
# Manually run Firth's logistic regression model using the binary matrix produced above
dataR <- cip_bin[, setdiff(names(cip_bin), c("id", "pheno", "ecoff", "mic", "NWT"))]
dataR <- dataR[, colSums(dataR, na.rm = TRUE) > 5]
modelR <- logistf::logistf(R ~ ., data = dataR, pl = FALSE)
```

```
#> Warning in logistf::logistf(R ~ ., data = dataR, pl = FALSE): logistf.fit:
#> Maximum number of iterations for full model exceeded. Try to increase the
#> number of iterations or alter step size by passing 'logistf.control(maxit=...,
#> maxstep=...)' to parameter control

summary(modelR)
#> logistf::logistf(formula = R ~ ., data = dataR, pl = FALSE)
#>
#> Model fitted by Penalized ML
#> Coefficients:
#>                        coef  se(coef) lower 0.95 upper 0.95       Chisq
#> (Intercept)      -5.3175097 0.2633180 -5.8336034 -4.8014159         Inf
#> gyrA_S83L         5.1068379 0.3391112  4.4421922  5.7714836         Inf
#> gyrA_D87Y         0.6085750 1.9592283 -3.2314420  4.4485920  0.09648461
#> gyrA_D87N         1.0349517 1.2899342 -1.4932729  3.5631763  0.64373192
#> parC_S80I         3.5331411 1.2607159  1.0621833  6.0040989  7.85393846
#> parE_S458A       -0.6705019 1.4210624 -3.4557329  2.1147292  0.22262490
#> parC_S80R         0.9079621 0.9428897 -0.9400679  2.7559920  0.92728574
#> parE_L416F        1.0858606 1.4754461 -1.8059607  3.9776818  0.54162839
#> parC_S57T         1.4187529 1.4556325 -1.4342344  4.2717401  0.94997028
#> soxS_A12S         1.5838463 1.4862202 -1.3290918  4.4967844  1.13568986
#> parC_A56T         2.6703270 1.5168050 -0.3025562  5.6432103  3.09934110
#> qnrB19            5.2773513 0.4540398  4.3874495  6.1672530         Inf
#> `aac(6')-Ib-cr5`  4.2828221 1.3434569  1.6496951  6.9159492 10.16278221
#> parC_E84V        -0.6804449 1.7890326 -4.1868843  2.8259946  0.14466032
#> parE_I529L        2.2022178 0.4576747  1.3051918  3.0992437 23.15297055
#> parE_S458T       -2.7640228 1.9475381 -6.5811274  1.0530817  2.01424047
#> parE_E460D       -1.5249129 1.8857582 -5.2209311  2.1711053  0.65391016
#> parC_E84G         1.2081572 1.5876105 -1.9035023  4.3198167  0.57910718
#> qnrS1             5.5126904 0.4383455  4.6535490  6.3718319         Inf
#> marR_S3N          3.1530001 0.5135941  2.1463742  4.1596261 37.68841990
#> parE_I355T        1.9462857 0.8716735  0.2378370  3.6547344  4.98546287
#> soxR_G121D       -2.5712233 1.6085720 -5.7239664  0.5815198  2.55504535
#> qnrB4             6.9269063 1.5713547  3.8471075 10.0067050 19.43256528
#> parE_D475E       -0.7063419 1.4133734 -3.4765029  2.0638192  0.24975608
#>                             p method
#> (Intercept)      0.000000e+00      1
#> gyrA_S83L        0.000000e+00      1
#> gyrA_D87Y        7.560897e-01      1
#> gyrA_D87N        4.223626e-01      1
#> parC_S80I        5.071012e-03      1
#> parE_S458A       6.370471e-01      1
#> parC_S80R        3.355692e-01      1
#> parE_L416F       4.617586e-01      1
#> parC_S57T        3.297269e-01      1
#> soxS_A12S        2.865649e-01      1
#> parC_A56T        7.832399e-02      1
#> qnrB19           0.000000e+00      1
#> `aac(6')-Ib-cr5` 1.433042e-03      1
#> parC_E84V        7.036913e-01      1
#> parE_I529L       1.496119e-06      1
#> parE_S458T       1.558292e-01      1
#> parE_E460D       4.187182e-01      1
#> parC_E84G        4.466625e-01      1
#> qnrS1            0.000000e+00      1
```

```
#> marR_S3N         8.299580e-10      1
#> parE_I355T       2.556115e-02      1
#> soxR_G121D       1.099427e-01      1
#> qnrB4            1.042148e-05      1
#> parE_D475E       6.172469e-01      1
#>
#> Method: 1-Wald, 2-Profile penalized log-likelihood, 3-None
#>
#> Likelihood ratio test=3338.78 on 23 df, p=0, n=3629
#> Wald test = 514.8685 on 23 df, p = 0

# Extract model summary details using `logistf_details()`
modelR_summary <- logistf_details(modelR)

modelR_summary
#> # A tibble: 24 × 5
#>    marker         est ci.lower ci.upper    pval
#>  * <chr>        <dbl>    <dbl>    <dbl>   <dbl>
#>  1 (Intercept) -5.32    -5.83     -4.80 0
#>  2 gyrA_S83L    5.11     4.44      5.77 0
#>  3 gyrA_D87Y    0.609   -3.23      4.45 0.756
#>  4 gyrA_D87N    1.03    -1.49      3.56 0.422
#>  5 parC_S80I    3.53     1.06      6.00 0.00507
#>  6 parE_S458A  -0.671   -3.46      2.11 0.637
#>  7 parC_S80R    0.908   -0.940     2.76 0.336
#>  8 parE_L416F   1.09    -1.81      3.98 0.462
#>  9 parC_S57T    1.42    -1.43      4.27 0.330
#> 10 soxS_A12S    1.58    -1.33      4.50 0.287
#> # ℹ 14 more rows
#> Use ggplot2::autoplot() on this output to visualise

# Plot the point estimates and 95% confidence intervals of the model
plot_estimates(modelR_summary)
```

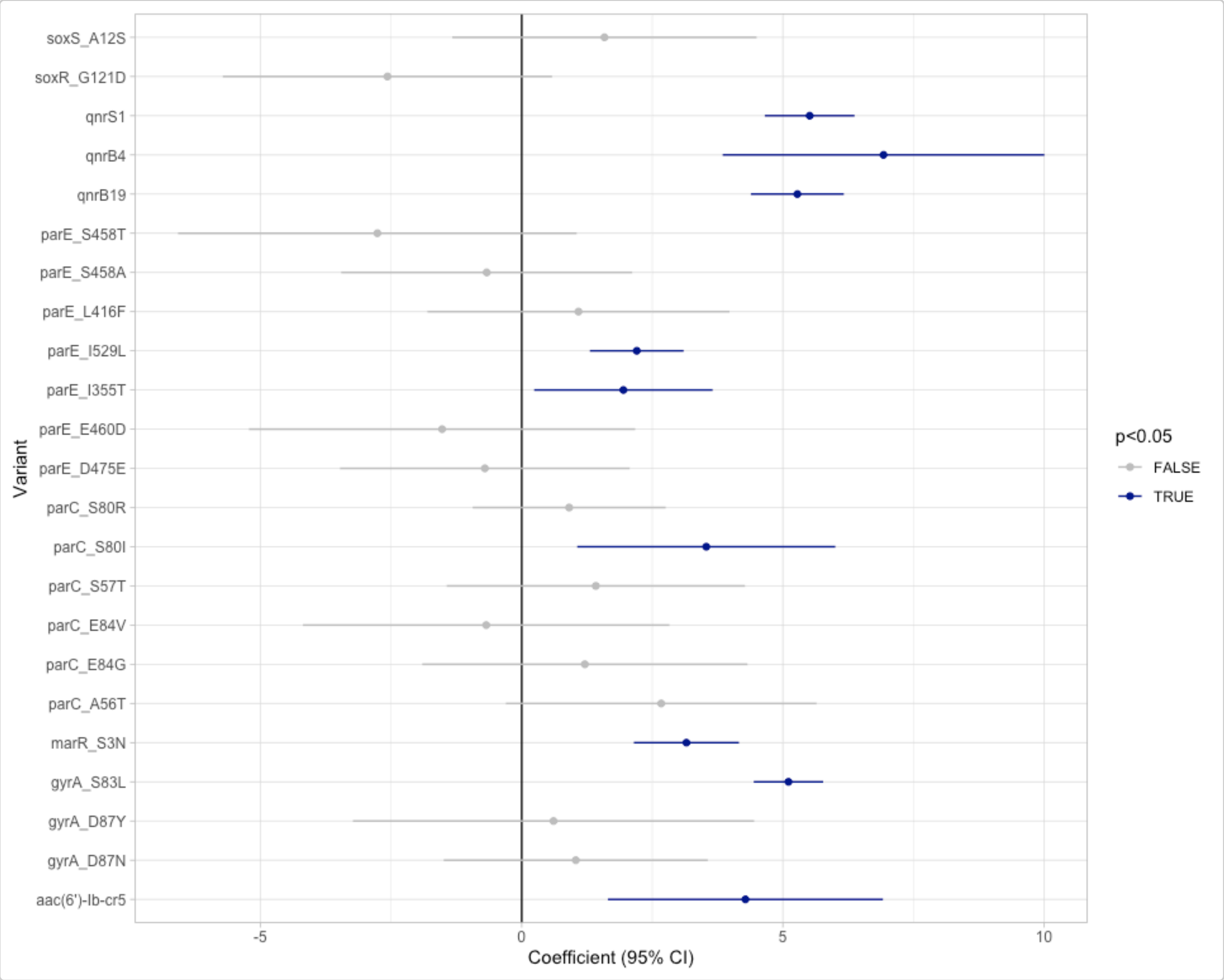


```
# Alternatively, use the amr_logistic() function to model R and NWT and plot the results
        together
models <- amr_logistic(
  geno_table = ecoli_geno,
  pheno_table = ecoli_pheno,
  sir_col = "pheno_clsi",
  pheno_drug = "Ciprofloxacin",
  geno_class = c("Quinolones"),
  maf = 10
)
#> Generating geno-pheno binary matrix
#>  Defining NWT in binary matrix using ecoff column provided: ecoff
#> ...Fitting logistic regression model to R using logistf
#>    Filtered data contains 3629 samples (793 => 1, 2836 => 0) and 19 variables.
#> Warning in logistf::logistf(R ~ ., data = to_fit, pl = FALSE): logistf.fit:
#> Maximum number of iterations for full model exceeded. Try to increase the
#> number of iterations or alter step size by passing 'logistf.control(maxit=...,
#> maxstep=...)' to parameter control
#> ...Fitting logistic regression model to NWT using logistf
#>    Filtered data contains 3576 samples (875 => 1, 2701 => 0) and 19 variables.
#> Warning in logistf::logistf(NWT ~ ., data = to_fit, pl = FALSE): logistf.fit:
#> Maximum number of iterations for full model exceeded. Try to increase the
#> number of iterations or alter step size by passing 'logistf.control(maxit=...,
#> maxstep=...)' to parameter control
```

```
#> Generating plots
#> Plotting 2 models
```

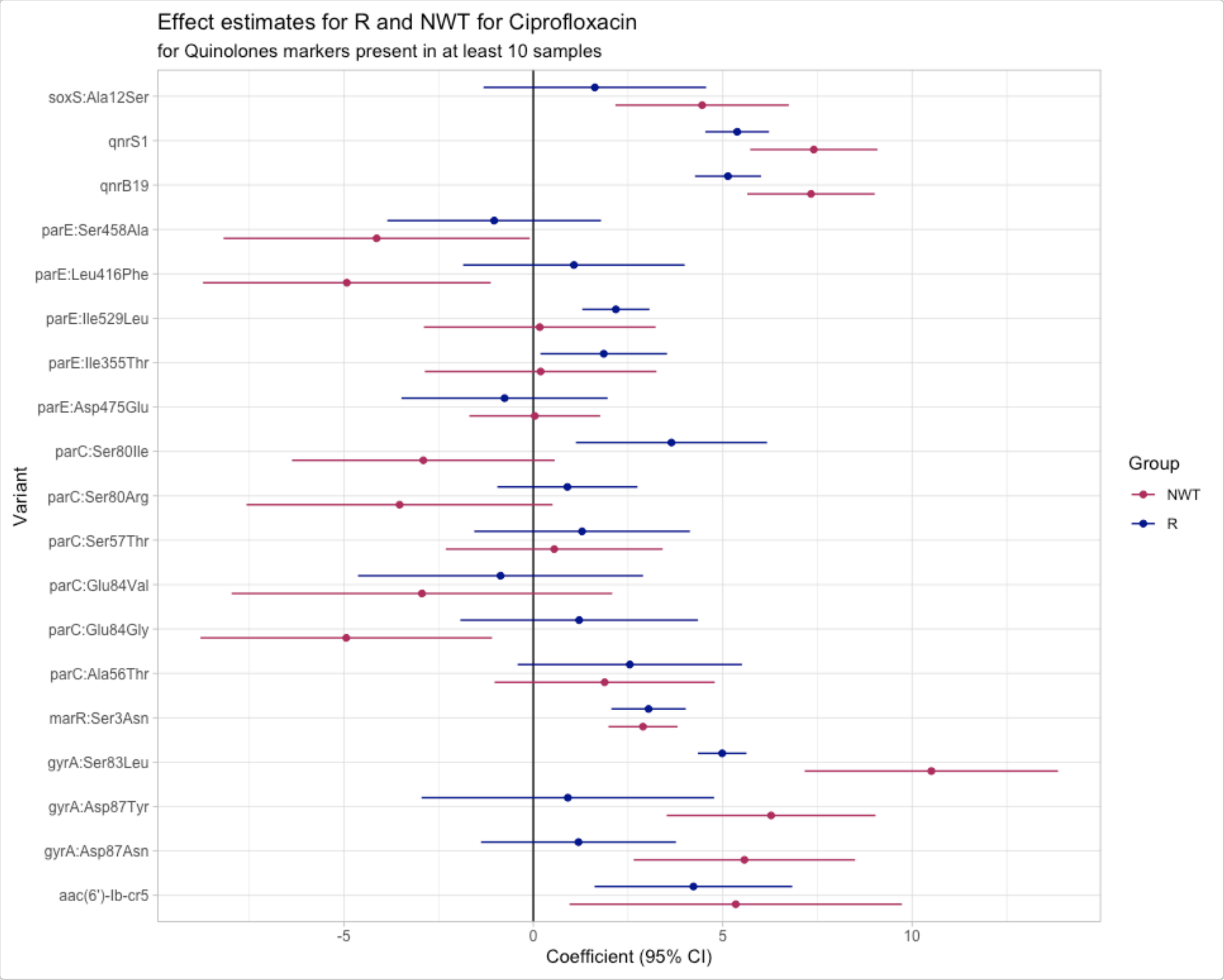


```
# Output tables
models$modelR
#> # A tibble: 20 × 5
#>    marker            est ci.lower ci.upper         pval
#>    <chr>           <dbl>    <dbl>    <dbl>        <dbl>
#>  1 (Intercept)    -5.19    -5.67     -4.70 0
#>  2 gyrA:Ser83Leu   4.99     4.34      5.63 0
#>  3 gyrA:Asp87Tyr   0.912   -2.95      4.77 0.643
#>  4 gyrA:Asp87Asn   1.19    -1.38      3.77 0.364
#>  5 parC:Ser80Ile   3.65     1.12      6.17 0.00462
#>  6 parE:Ser458Ala -1.03    -3.85      1.79 0.473
#>  7 parC:Ser80Arg   0.900   -0.949     2.75 0.340
#>  8 parE:Leu416Phe  1.07    -1.85      4.00 0.473
#>  9 parC:Ser57Thr   1.29    -1.56      4.13 0.376
#> 10 soxS:Ala12Ser   1.62    -1.31      4.56 0.279
#> 11 parC:Ala56Thr   2.55    -0.415     5.51 0.0919
#> 12 qnrB19          5.14     4.27      6.01 0
#> 13 aac(6')-Ib-cr5  4.23     1.62      6.84 0.00150
#> 14 parC:Glu84Val  -0.866   -4.63      2.90 0.652
#> 15 parE:Ile529Leu  2.18     1.29      3.07 0.00000151
#> 16 parC:Glu84Gly   1.21    -1.93      4.35 0.450
```

```
#> 17 qnrS1           5.38     4.54      6.22 0
#> 18 marR:Ser3Asn    3.04     2.06      4.02 0.00000000128
#> 19 parE:Ile355Thr  1.86     0.188     3.53 0.0292
#> 20 parE:Asp475Glu -0.759   -3.48      1.96 0.585
#> Use ggplot2::autoplot() on this output to visualise

models$modelNWT
#> # A tibble: 20 × 5
#>    marker             est ci.lower ci.upper     pval
#>    <chr>            <dbl>    <dbl>    <dbl>    <dbl>
#>  1 (Intercept)    -4.46     -4.82   -4.10   0
#>  2 gyrA:Ser83Leu  10.5       7.16   13.9    7.30e-10
#>  3 gyrA:Asp87Tyr   6.28      3.52    9.03   8.08e- 6
#>  4 gyrA:Asp87Asn   5.57      2.65    8.49   1.87e- 4
#>  5 parC:Ser80Ile  -2.90     -6.37    0.564  1.01e- 1
#>  6 parE:Ser458Ala -4.14     -8.18   -0.0956 4.48e- 2
#>  7 parC:Ser80Arg  -3.53     -7.57    0.509  8.67e- 2
#>  8 parE:Leu416Phe -4.92     -8.72   -1.12   1.11e- 2
#>  9 parC:Ser57Thr   0.552    -2.31    3.41   7.06e- 1
#> 10 soxS:Ala12Ser   4.46      2.17    6.75   1.36e- 4
#> 11 parC:Ala56Thr   1.88     -1.02    4.79   2.04e- 1
#> 12 qnrB19          7.33      5.65    9.01   0
#> 13 aac(6')-Ib-cr5  5.34      0.957   9.73   1.70e- 2
#> 14 parC:Glu84Val  -2.94     -7.96    2.08   2.51e- 1
#> 15 parE:Ile529Leu  0.170    -2.89    3.23   9.13e- 1
#> 16 parC:Glu84Gly  -4.94     -8.79   -1.09   1.19e- 2
#> 17 qnrS1           7.40      5.72    9.08   0
#> 18 marR:Ser3Asn    2.90      1.98    3.81   4.78e-10
#> 19 parE:Ile355Thr  0.193    -2.86    3.25   9.02e- 1
#> 20 parE:Asp475Glu  0.0389   -1.69    1.77   9.65e- 1
#> Use ggplot2::autoplot() on this output to visualise

# Note the matrix output is the same as cip_bin.
models$binary_matrix
#> # A tibble: 3,629 × 51
#>    id         pheno ecoff     mic  disk     R   NWT gyrA..Ser83Leu gyrA..Asp87Tyr
#>    <chr>      <sir> <sir>   <mic> <dsk> <dbl> <dbl>          <dbl>          <dbl>
#>  1 SAMN0317… S      WT    <=0.015    NA     0     0              0              0
#>  2 SAMN0317… S      WT    <=0.015    NA     0     0              0              0
#>  3 SAMN0317… S      WT    <=0.015    NA     0     0              0              0
#>  4 SAMN0317… S      NWT     0.250    NA     0     1              1              0
#>  5 SAMN0317… S      NWT     0.120    NA     0     1              0              1
#>  6 SAMN0317… S      WT    <=0.015    NA     0     0              0              0
#>  7 SAMN0317… S      WT    <=0.015    NA     0     0              0              0
#>  8 SAMN0317… R      NWT    >4.000    NA     1     1              1              0
#>  9 SAMN0317… S      NWT     0.250    NA     0     1              1              0
#> 10 SAMN0317… R      NWT    >4.000    NA     1     1              1              0
#> # ℹ 3,619 more rows
#> # ℹ 42 more variables: gyrA..Asp87Asn <dbl>, parC..Ser80Ile <dbl>,
#> #   parE..Ser458Ala <dbl>, parC..Ser80Arg <dbl>, parE..Leu416Phe <dbl>,
#> #   qnrB6 <dbl>, gyrA..Asp87Gly <dbl>, parC..Ser57Thr <dbl>,
#> #   parC..Glu84Ala <dbl>, soxS..Ala12Ser <dbl>, qnrB2 <dbl>, qnrS2 <dbl>,
#> #   parC..Glu84Lys <dbl>, parC..Ala56Thr <dbl>, qnrB19 <dbl>,
#> #   `aac(6')-Ib-cr5` <dbl>, parC..Glu84Val <dbl>, parE..Ile529Leu <dbl>, …
```

#### 8. Assess solo positive predictive value of genetic markers

The strongest evidence of the effect of an individual genetic marker on a drug phenotype is its positive predictive value (PPV) for resistance amongst strains that carry this marker 'solo' with no other markers known to be associated with resistance to the drug class. This is referred to as 'solo PPV'.

The function `solo_ppv()` takes as input our genotype and phenotype tables, and calculates solo PPV for resistance to a specific drug (included in our phenotype table) for markers associated with the specified drug class (included in our genotype table). It uses the `get_binary_matrix()` function to first calculate the binary matrix, then filters out all samples that have more than one marker.

It then calculates for each remaining marker, amongst the genomes in which that marker is found solo, the number of genomes, the number and proportion that are R or NWT, and the 95% confidence intervals for these proportions. The values are returned as a table, and also plotted so we can easily visualise the distribution of S/I/R calls and the solo PPV for R and NWT, for each solo marker.

The function returns 4 objects:

- `solo_stats`: data frame containing the numbers, proportions and confidence intervals for PPV of R and NWT categories
- `amr_binary`: the (wide format) binary matrix for all strains with geno/pheno data for the specified drug/class
- `solo_binary`: the (long format) binary matrix for only those strains in which a solo marker was found, i.e. the data used to calculate PPV
- `combined_plot`: a plot showing the distribution of S/I/R calls and the solo PPV for R and NWT, for each solo marker

```
# Run a solo PPV analysis
soloPPV_cipro <- solo_ppv(
  ecoli_geno,
  ecoli_pheno,
  sir_col = "pheno_clsi",
  pheno_drug = "Ciprofloxacin",
  geno_class = "Quinolones"
)
#> Generating geno-pheno binary matrix
#>  Defining NWT in binary matrix using ecoff column provided: ecoff
#> Warning: Removed 1 row containing missing values or values outside the scale range
#> (`geom_segment()`).
#> Warning: Removed 1 row containing missing values or values outside the scale range
#> (`geom_point()`).
```

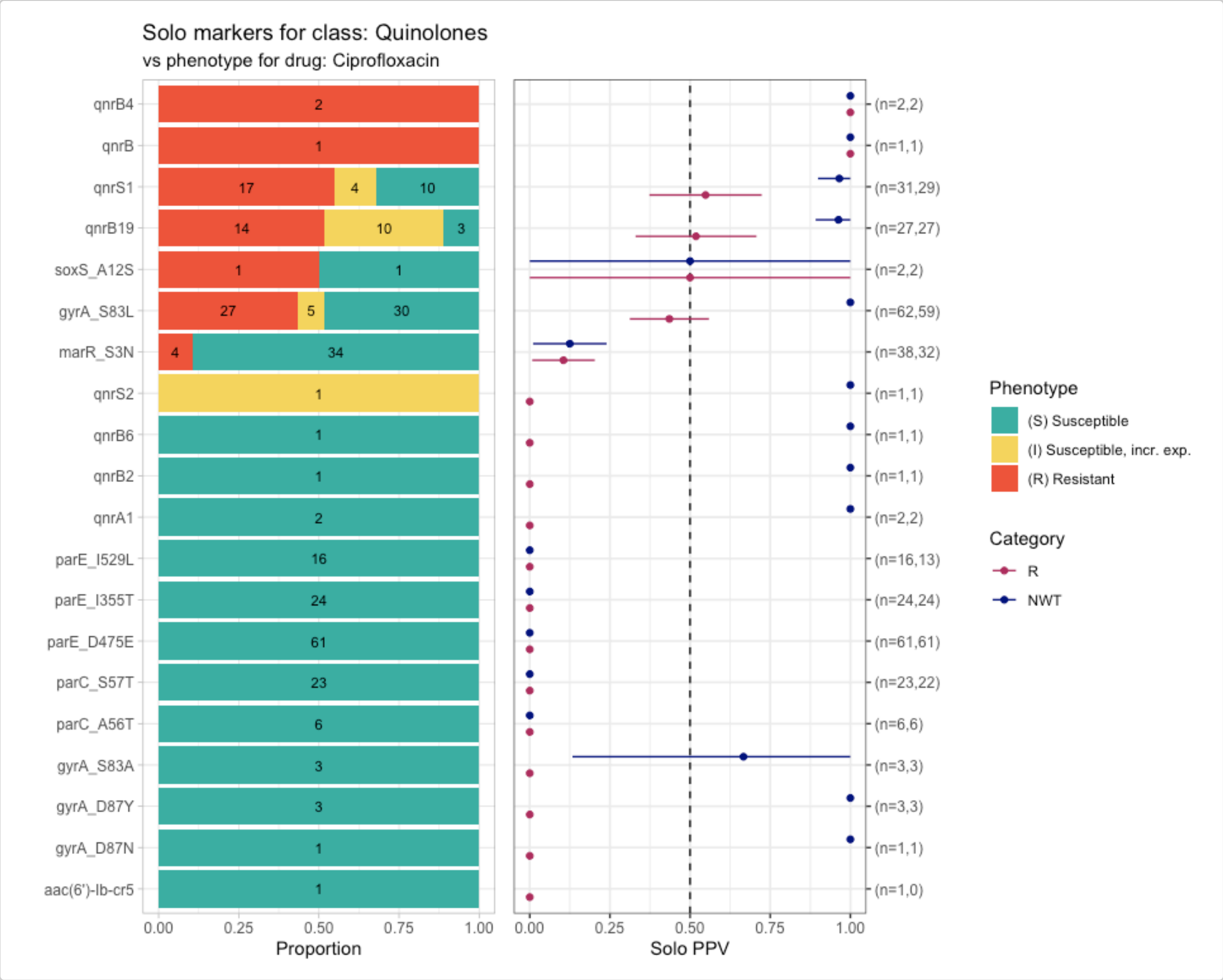


```
# Output table
soloPPV_cipro$solo_stats
#> # A tibble: 40 × 8
#>    marker          category     x     n   ppv    se ci.lower ci.upper
#>    <chr>           <chr>    <dbl> <int> <dbl> <dbl>    <dbl>    <dbl>
#>  1 aac(6')-Ib-cr5 R            0     1     0     0        0        0
#>  2 gyrA_D87N      R            0     1     0     0        0        0
#>  3 gyrA_D87Y      R            0     3     0     0        0        0
#>  4 gyrA_S83A      R            0     3     0     0        0        0
#>  5 parC_A56T      R            0     6     0     0        0        0
#>  6 parC_S57T      R            0    23     0     0        0        0
#>  7 parE_D475E     R            0    61     0     0        0        0
#>  8 parE_I355T     R            0    24     0     0        0        0
#>  9 parE_I529L     R            0    16     0     0        0        0
#> 10 qnrA1          R            0     2     0     0        0        0
#> # ℹ 30 more rows

# Interim matrices with data used to compute stats and plots
soloPPV_cipro$solo_binary
#> # A tibble: 306 × 9
#>    id           pheno ecoff   mic  disk     R   NWT marker     value
#>    <chr>        <sir> <sir> <mic> <dsk> <dbl> <dbl> <chr>      <dbl>
#>  1 SAMN03177618 S     NWT    0.25    NA     0     1 gyrA_S83L      1
#>  2 SAMN03177619 S     NWT    0.12    NA     0     1 gyrA_D87Y      1
```

```
#>  3 SAMN03177623 S     NWT    0.25    NA     0     1 gyrA_S83L     1
#>  4 SAMN03177631 S     NWT    0.25    NA     0     1 gyrA_S83L     1
#>  5 SAMN03177635 S     NWT    0.25    NA     0     1 gyrA_S83L     1
#>  6 SAMN03177637 S     NWT    0.25    NA     0     1 gyrA_S83L     1
#>  7 SAMN03177638 S     NWT    0.25    NA     0     1 qnrB6         1
#>  8 SAMN03177639 S     NWT    0.12    NA     0     1 gyrA_S83L     1
#>  9 SAMN03177643 S     NWT    0.25    NA     0     1 gyrA_S83L     1
#> 10 SAMN03177646 S     NWT    0.25    NA     0     1 gyrA_S83L     1
#> # ℹ 296 more rows
```

```
soloPPV_cipro$amr_binary
#> # A tibble: 3,629 × 51
#>    id         pheno ecoff     mic  disk     R   NWT gyrA_S83L gyrA_D87Y gyrA_D87N
#>    <chr>      <sir> <sir>   <mic> <dsk> <dbl> <dbl>     <dbl>     <dbl>     <dbl>
#>  1 SAMN0317… S     WT    <=0.015    NA     0     0         0         0         0
#>  2 SAMN0317… S     WT    <=0.015    NA     0     0         0         0         0
#>  3 SAMN0317… S     WT    <=0.015    NA     0     0         0         0         0
#>  4 SAMN0317… S     NWT     0.250    NA     0     1         1         0         0
#>  5 SAMN0317… S     NWT     0.120    NA     0     1         0         1         0
#>  6 SAMN0317… S     WT    <=0.015    NA     0     0         0         0         0
#>  7 SAMN0317… S     WT    <=0.015    NA     0     0         0         0         0
#>  8 SAMN0317… R     NWT    >4.000    NA     1     1         1         0         1
#>  9 SAMN0317… S     NWT     0.250    NA     0     1         1         0         0
#> 10 SAMN0317… R     NWT    >4.000    NA     1     1         1         0         1
#> # ℹ 3,619 more rows
#> # ℹ 41 more variables: parC_S80I <dbl>, parE_S458A <dbl>, parC_S80R <dbl>,
#> #   parE_L416F <dbl>, qnrB6 <dbl>, gyrA_D87G <dbl>, parC_S57T <dbl>,
#> #   parC_E84A <dbl>, soxS_A12S <dbl>, qnrB2 <dbl>, qnrS2 <dbl>,
#> #   parC_E84K <dbl>, parC_A56T <dbl>, qnrB19 <dbl>, `aac(6')-Ib-cr5` <dbl>,
#> #   parC_E84V <dbl>, parE_I529L <dbl>, parE_S458T <dbl>, parE_E460D <dbl>,
#> #   parC_E84G <dbl>, qnrS1 <dbl>, marR_S3N <dbl>, `aac(6')-Ib-cr` <dbl>, …
```

#### 9. Compare markers with assay data

So far we have considered only the impact of individual markers, and their association with categorical S/I/R or WT/NWT calls.

##### UpSet plots

The function `amr_upset()` takes as binary matrix table `cip_bin` summarising ciprofloxacin resistance vs quinolone markers, generated using `get_binary_matrix()`, and explores the distribution of MIC or disk diffusion assay values for all observed combinations of markers (solo or multiple markers). It visualises the data in the form of an upset plot, showing the distribution of assay values and S/I/R calls for each observed marker combination, and returns a summary of these distributions (including sample size, median and interquartile range, number and proportion classified as R).

The function returns 2 objects:

- `summary`: data frame containing summarising the data associated with each combination of markers
- `plot`: an upset plot showing the distribution of assay values, and breakdown of S/I/R calls, for each observed marker combination

```
# Compare ciprofloxacin MIC data with quinolone marker combinations,
#    using the binary matrix we constructed earlier via get_binary_matrix()
cipro_mic_upset <- amr_upset(
  cip_bin,
  min_set_size = 2,
  assay = "mic",
  order = "value"
)
#> Ordering markers by frequency
#> Scale for y is already present.
#> Adding another scale for y, which will replace the existing scale.
#> Scale for y is already present.
#> Adding another scale for y, which will replace the existing scale.
```

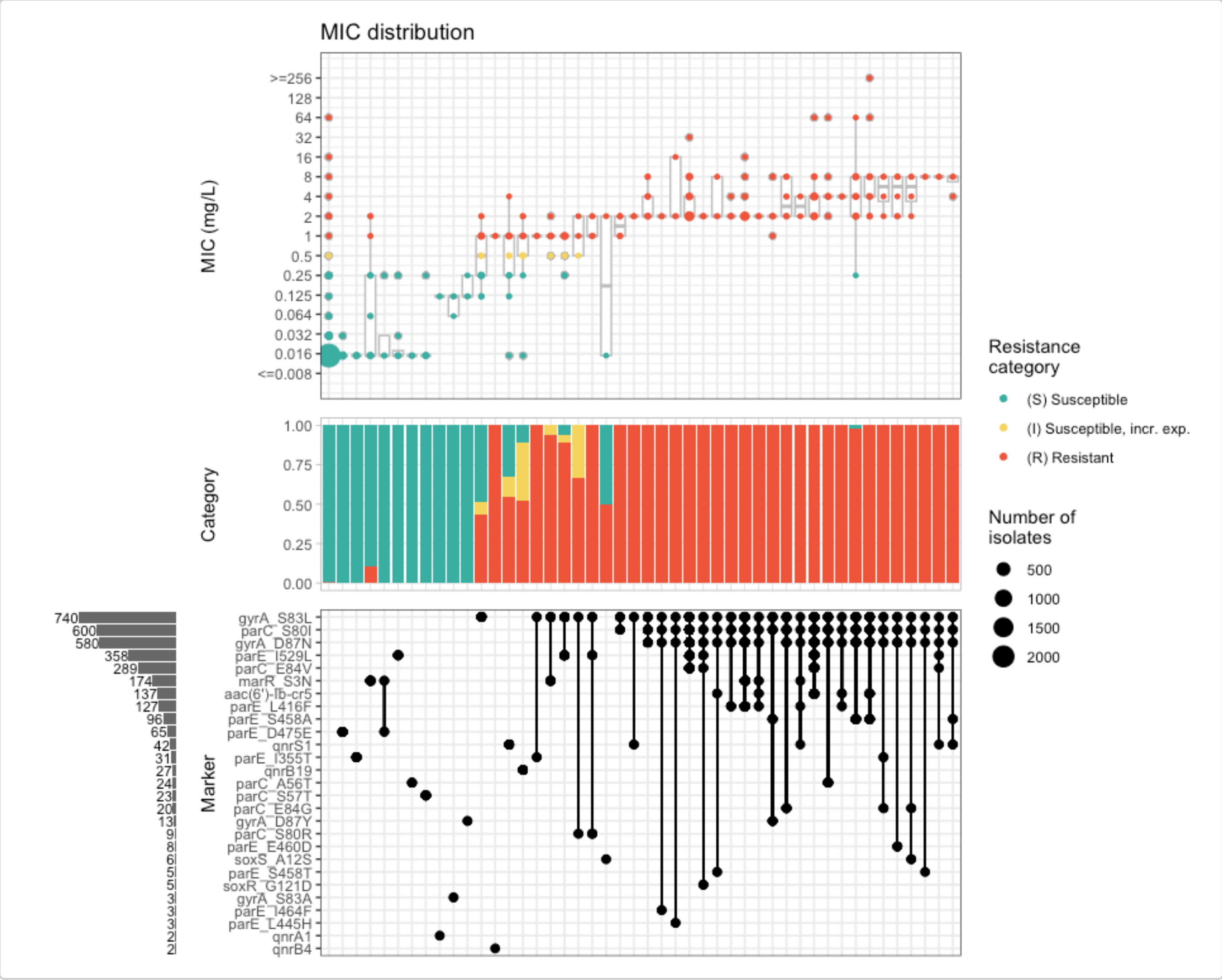


```
# Output table
cipro_mic_upset$summary
#> # A tibble: 103 × 21
#>    marker_list        marker_count     n combination_id   R.n   R.ppv R.ci_lower
#>    <chr>                     <dbl> <int> <fct>           <dbl>   <dbl>      <dbl>
#>  1 ""                            0  2590 0_0_0_0_0_0_0…     10 0.00386    0.00147
#>  2 "qnrB"                        1     1 0_0_0_0_0_0_0…      1 1          1
#>  3 "parE_E460K, gyrA…            2     1 0_0_0_0_0_0_0…      1 1          1
#>  4 "parE_D475E"                  1    61 0_0_0_0_0_0_0…      0 0          0
```

```
#>  5 "qnrA1"               1     2 0_0_0_0_0_0_0…    0 0         0
#>  6 "gyrA_S83A"           1     3 0_0_0_0_0_0_0…    0 0         0
#>  7 "qnrB4"               1     2 0_0_0_0_0_0_0…    2 1         1
#>  8 "parE_I355T"          1    24 0_0_0_0_0_0_0…    0 0         0
#>  9 "marR_S3N"            1    38 0_0_0_0_0_0_0…    4 0.105     0.00769
#> 10 "marR_S3N, parE_D…    2     4 0_0_0_0_0_0_0…    0 0         0
#> # ℹ 93 more rows
#> # ℹ 14 more variables: R.ci_upper <dbl>, R.denom <int>, NWT.n <dbl>,
#> #   NWT.ppv <dbl>, NWT.ci_lower <dbl>, NWT.ci_upper <dbl>, NWT.denom <int>,
#> #   median_excludeRangeValues <dbl>, q25_excludeRangeValues <dbl>,
#> #   q75_excludeRangeValues <dbl>, n_excludeRangeValues <int>,
#> #   median_ignoreRanges <dbl>, q25_ignoreRanges <dbl>, q75_ignoreRanges <dbl>
```

#### PPV plots

The function `amr_ppv()` uses the same underlying approach as `amr_upset()` but transposes the orientation of the data so it looks more like the `solo_ppv()` plot but for combinations of markers as well as those found solo.

Like `amr_upset` it takes as binary matrix table `cip_bin` summarising ciprofloxacin resistance vs quinolone markers, generated using `get_binary_matrix()`, and creates a multi-panel plot and summary statistics table.

The left-most panel indicates which marker/combinations are shown in each row of the plot, either as a list of marker names (default) or as an upset-style grid (set upset_grid=`TRUE` to turn this on).

The other panels available are

- category plot: stacked bar plot showing S/I/R calls (ON by default, set plot_category=`FALSE` to turn this off)
- PPV plot: forest-style plot showing point estimates for R/NWT PPV, with horizontal lines indicating 95% confidence intervals (ON by default, set plot_ppv=`FALSE` to turn this off)
- assay plot: boxplot of assay (MIC/disk) values (OFF by default, set plot_assay=`TRUE` and assay="`mic`" or assay="`disk`" to turn this on)

The function returns 2 objects:

- `summary`: data frame containing summarising the data associated with each combination of markers
- `plot`: an upset plot showing the distribution of assay values, and breakdown of S/I/R calls, for each observed marker combination

```
# Default plot
cipro_mic_ppv <- amr_ppv(
  cip_bin,
  min_set_size = 10,
  upset_grid = T
)
#> Ordering markers by frequency
#> Scale for y is already present.
#> Adding another scale for y, which will replace the existing scale.
```

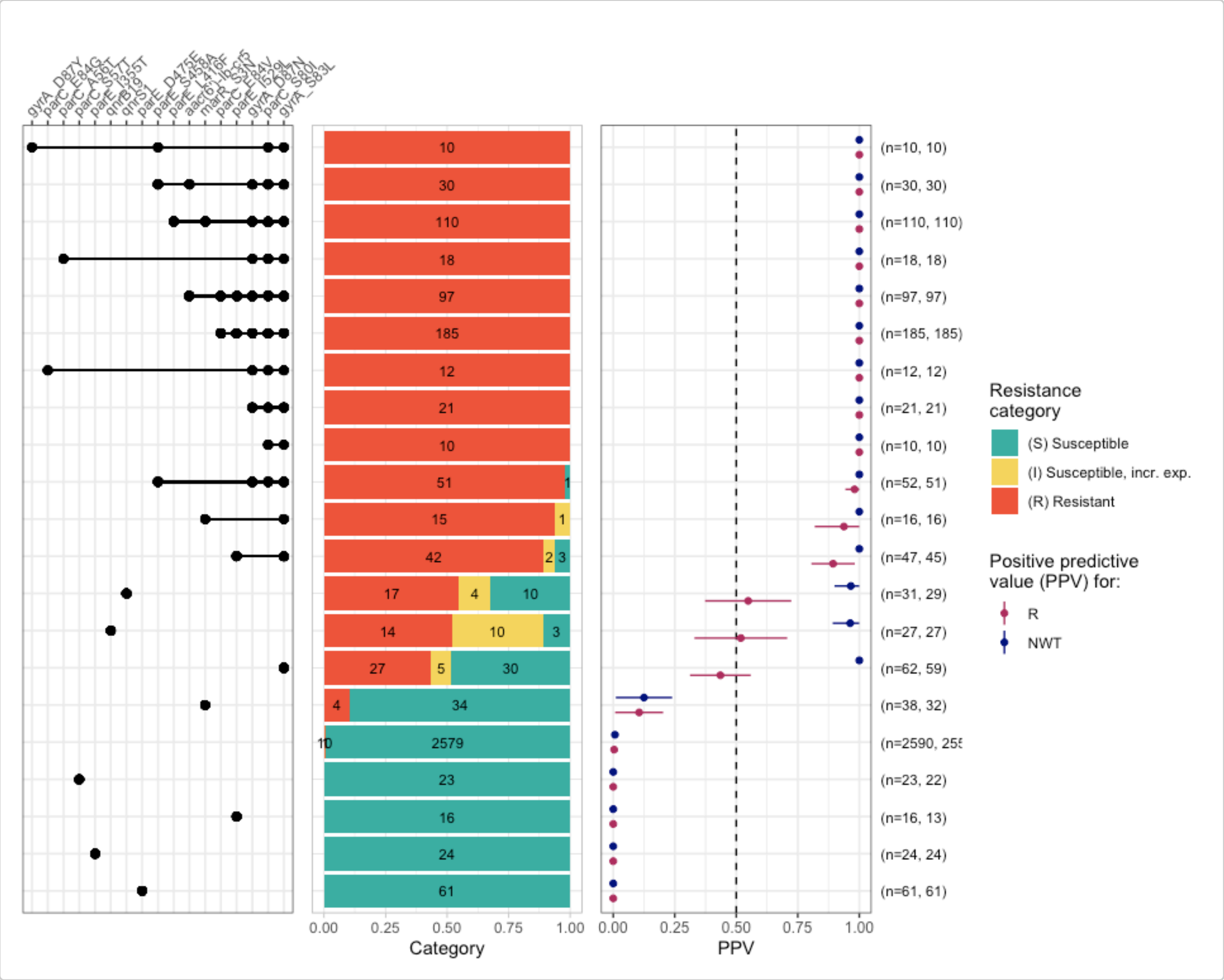


```
# add MIC, remove category plot, label rows with marker list
cipro_mic_ppv2 <- amr_ppv(
  cip_bin,
  min_set_size = 10,
  upset_grid = F,
  plot_assay = T, assay = "mic",
  plot_category = F
)
#> Ordering markers by frequency
```

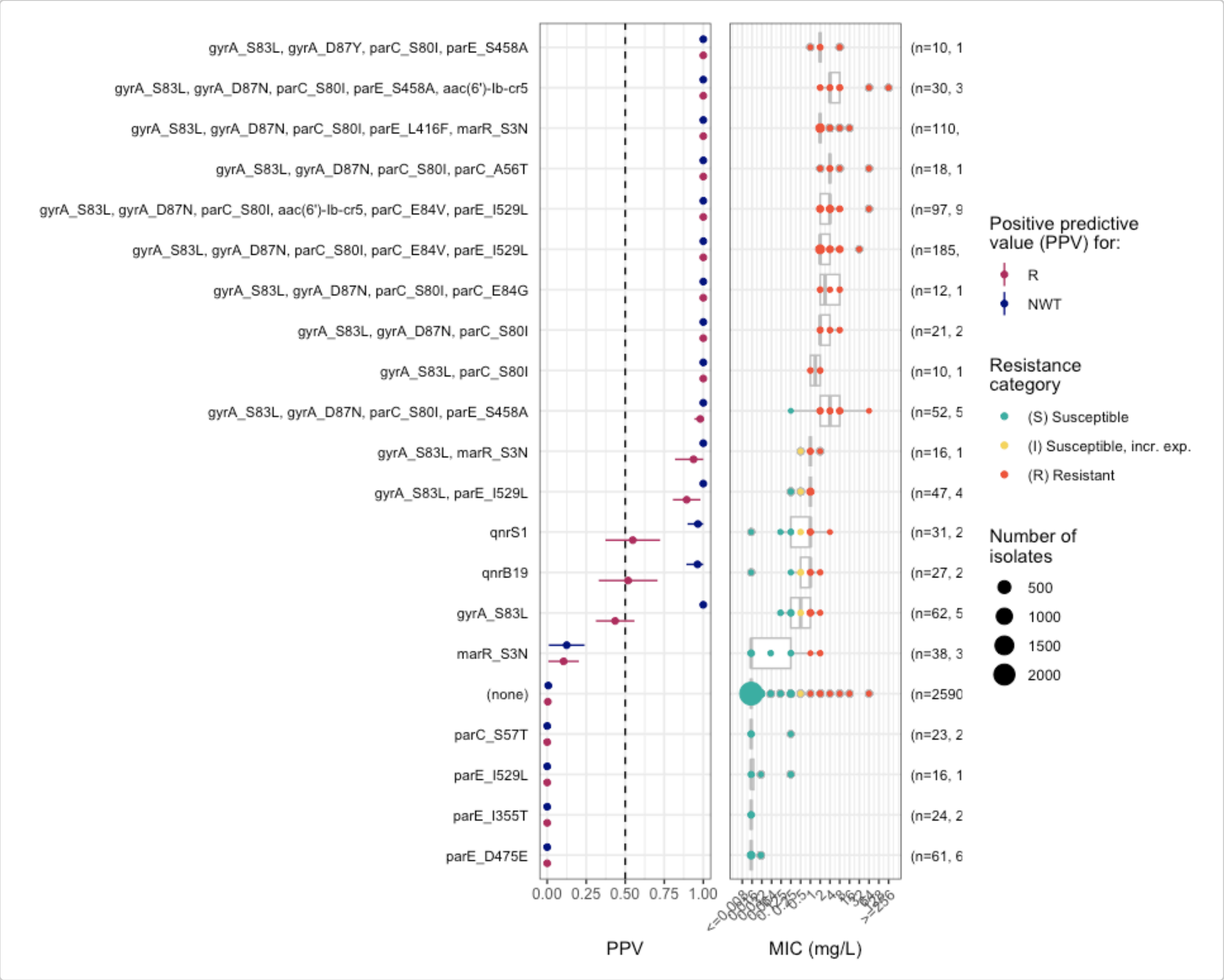


```
# Output table
cipro_mic_ppv$summary
#> # A tibble: 103 × 14
#>    marker_list        marker_count     n combination_id   R.n   R.ppv R.ci_lower
#>    <chr>                     <dbl> <int> <fct>          <dbl>   <dbl>      <dbl>
#>  1 ""                            0  2590 0_0_0_0_0_0_0…    10 0.00386    0.00147
#>  2 "qnrB"                        1     1 0_0_0_0_0_0_0…     1 1          1
#>  3 "parE_E460K, gyrA…            2     1 0_0_0_0_0_0_0…     1 1          1
#>  4 "parE_D475E"                  1    61 0_0_0_0_0_0_0…     0 0          0
#>  5 "qnrA1"                       1     2 0_0_0_0_0_0_0…     0 0          0
#>  6 "gyrA_S83A"                   1     3 0_0_0_0_0_0_0…     0 0          0
#>  7 "qnrB4"                       1     2 0_0_0_0_0_0_0…     2 1          1
#>  8 "parE_I355T"                  1    24 0_0_0_0_0_0_0…     0 0          0
#>  9 "marR_S3N"                    1    38 0_0_0_0_0_0_0…     4 0.105      0.00769
#> 10 "marR_S3N, parE_D…            2     4 0_0_0_0_0_0_0…     0 0          0
#> # ℹ 93 more rows
#> # ℹ 7 more variables: R.ci_upper <dbl>, R.denom <int>, NWT.n <dbl>,
#> #   NWT.ppv <dbl>, NWT.ci_lower <dbl>, NWT.ci_upper <dbl>, NWT.denom <int>
```

### Downloading geno-pheno data from NCBI and EBI

### Introduction

This vignette demonstrates how to download antibiotic susceptibility testing (AST) data from NCBI and EBI, and to re-interpret it using different clinical breakpoints.

Start by loading the `AMRgen` package:

```
library(AMRgen)
library(dplyr)
```

#### Option 1: Download data from NCBI

##### Option 1a: Download AST data from NCBI via rentrez

The `download_ncbi_pheno()` function lets you download antibiogram data from NCBI via their 'EUtils' API using the rentrez R pacakge. You must specify a species, and can optionally limit the download to one or more specific drugs. The function can also re-format the data into an AMRgen phenotype table, and re-interpret phenotypes against clinical breakpoints from EUCAST or CLSI.

This function is quite slow, especially for organisms with many BioSamples to search through, but it is free and requires no authentication to use. Alternatively, you may prefer to try the BigQuery functions below (Option 1b), which provide more efficient and complete access to NCBI data but require authentication via a Google Cloud account.

```
# Download Staphylococcus aureus AST data from NCBI, filtering for amikacin and
        doxycycline, and re-interpret with EUCAST breakpoints
staph_pheno_ncbi <- download_ncbi_pheno(
  species = "Staphylococcus aureus",
  pheno_drug = c("amikacin", "DOX"), # antibiotics can be listed in short or long form
  reformat = TRUE,
  interpret_eucast = TRUE
) # reformat must be true to use interpret_* argument


# check how many samples retrieved
nrow(staph_pheno_ncbi)
#> [1] 143

# check the output
head(staph_pheno_ncbi)
#> # A tibble: 6 × 19
#>   id      drug    mic  disk pheno_provided pheno_eucast guideline method platform
#>   <chr>   <ab>  <mic> <dsk> <sir>          <sir>        <chr>     <chr>  <chr>
#> 1 SAMN4… DOX:…    <=1    NA S              S            CLSI      broth… <NA>
#> 2 SAMN4… DOX:…    <=1    NA S              S            CLSI      broth… <NA>
#> 3 SAMN3… AMK:…    <=2    NA S              S            CLSI      broth… Vitek
```

```
#> 4 SAMN3… AMK:…    NA    21 S                S            CLSI      disk … Scan 500
#> 5 SAMN2… AMK:…    NA    23 S                S            CLSI      disk … Scan 500
#> 6 SAMN2… AMK:…    NA    22 S                S            CLSI      disk … Scan 500
#> # ℹ 10 more variables: source <chr>, spp_pheno <mo>,
#> #   `Resistance phenotype` <chr>, `Measurement sign` <chr>, Measurement <chr>,
#> #   `Measurement units` <chr>, Vendor <chr>,
#> #   `Laboratory typing method version or reagent` <chr>,
#> #   pheno_eucast_mic <sir>, pheno_eucast_disk <sir>


# This is the same as downloading the data then re-interpreting it separately:
staph_pheno_ncbi_raw <- download_ncbi_pheno(
  species = "Staphylococcus aureus",
  pheno_drug = c("amikacin", "DOX"),
  reformat = FALSE,
  interpret_eucast = FALSE
)


head(staph_pheno_ncbi_raw)
#> # A tibble: 6 × 13
#>   id    BioProject organism Antibiotic `Resistance phenotype` `Measurement sign`
#>   <chr> <chr>      <chr>    <chr>      <chr>                  <chr>
#> 1 SAMN… PRJNA3915… Staphyl… doxycycli… susceptible            <=
#> 2 SAMN… PRJNA3915… Staphyl… doxycycli… susceptible            <=
#> 3 SAMN… PRJNA2788… Staphyl… amikacin   susceptible            <=
#> 4 SAMN… PRJNA7548… Staphyl… amikacin   susceptible            ==
#> 5 SAMN… PRJNA7548… Staphyl… amikacin   susceptible            ==
#> 6 SAMN… PRJNA7548… Staphyl… amikacin   susceptible            ==
#> # ℹ 7 more variables: Measurement <chr>, `Measurement units` <chr>,
#> #   `Laboratory typing method` <chr>, `Laboratory typing platform` <chr>,
#> #   Vendor <chr>, `Laboratory typing method version or reagent` <chr>,
#> #   `Testing standard` <chr>

# Then reformat and re-interpret using EUCAST and CLSI breakpoints, and ECOFFs using the
        import_ncbi_pheno() function
staph_pheno_ncbi2 <- import_ncbi_biosample(
  input = staph_pheno_ncbi_raw,
  interpret_clsi = TRUE,
  interpret_eucast = TRUE,
  interpret_ecoff = TRUE
)
#> Parsing column organism as micro-organism (class 'mo')
#> Renaming column organism to standard name 'spp_pheno'
#> Parsing column Antibiotic as antibiotic (class 'ab')
#> Renaming column Antibiotic to standard name 'drug'
#> Parsing column mic as class 'mic'
#> Parsing column disk as class 'disk'
#> Parsing column pheno_provided as class 'sir'
#> Renaming column Laboratory typing method to standard name 'method'
#> Renaming column Laboratory typing platform to standard name 'platform'
#> Renaming column Testing standard to standard name 'guideline'
#> Renaming column BioProject to standard name 'source'

head(staph_pheno_ncbi2)
#> # A tibble: 6 × 25
```

```
#>   id    drug    mic  disk pheno_provided pheno_eucast pheno_clsi ecoff guideline
#>   <chr> <ab>  <mic> <dsk> <sir>          <sir>        <sir>      <sir> <chr>
#> 1 SAMN… DOX:…   <=1    NA S              S            S          NI    CLSI
#> 2 SAMN… DOX:…   <=1    NA S              S            S          NI    CLSI
#> 3 SAMN… AMK:…   <=2    NA S              S            NA         WT    CLSI
#> 4 SAMN… AMK:…    NA    21 S              S            NA         WT    CLSI
#> 5 SAMN… AMK:…    NA    23 S              S            NA         WT    CLSI
#> 6 SAMN… AMK:…    NA    22 S              S            NA         WT    CLSI
#> # ℹ 16 more variables: method <chr>, platform <chr>, source <chr>,
#> #   spp_pheno <mo>, `Resistance phenotype` <chr>, `Measurement sign` <chr>,
#> #   Measurement <chr>, `Measurement units` <chr>, Vendor <chr>,
#> #   `Laboratory typing method version or reagent` <chr>,
#> #   pheno_eucast_mic <sir>, pheno_eucast_disk <sir>, pheno_clsi_mic <sir>,
#> #   pheno_clsi_disk <sir>, ecoff_mic <sir>, ecoff_disk <sir>

# Note that when there is a mix of MIC and disk data, a separate _disk and _mic
        interpretation column, as well as an overall phenotype column, is produced for
        each interpretation.
```

This produces a long format data frame, with one row per sample and drug combination. This is compatible with downstream functions in the AMRgen package.

Consider altering the `max_records`, `batch_size` or `sleep_time` options if you want to download a lot of data or run into NCBI server issues.

#### Option 1b: Download AST and genotype data from NCBI via bigrquery

NCBI data can be accessed via Google Cloud BigQuery. This requires a Google Cloud account. For more information about using BigQuery to explore NCBI Pathogen Detection data see [https://www.ncbi.nlm.nih.gov/pathogens/docs/getting_started_bigquery/](https://www.ncbi.nlm.nih.gov/pathogens/docs/getting_started_bigquery/).

The `query_ncbi_bq_pheno()` function lets you download antibiogram data from NCBI via Google Cloud BigQuery using the bigrquery R pacakge. You must specify a species, and can optionally limit the download to one or more specific drugs. The function can also re-format the data into an AMRgen phenotype table, and re-interpret phenotypes against clinical breakpoints from EUCAST or CLSI.

This function is fast but requires authentication via a [Google Cloud account]() and may require payment. Free trial accounts can be set up, but require credit card authorisation. Google currently provides enough free tier usage for >150 different queries for genotype data per month. To use this you will also need to install the `bigrquery` package and authorize it to use your Google cloud account.

```
install.packages("bigrquery")
library(bigrquery)
bigrquery::bq_auth()
```

##### To download AST data

```
# Download Staphylococcus aureus AST data from NCBI, filtering for amikacin and
        doxycycline
# NOTE: you may need to add 'project_id="xxx"' to the command if you have not set up
        application default credentials
staph_pheno_ncbi_cloud_raw <- query_ncbi_bq_pheno(
  taxgroup = "Staphylococcus aureus",
  pheno_drug = c("amikacin", "DOX")
)
```

```r
# Import and reinterpret using CLSI breakpoints
# NOTE: you may need to add 'project_id="xxx"' to the command if you have not set up
        application default credentials
staph_pheno_ncbi_cloud <- import_ncbi_pheno(staph_pheno_ncbi_cloud_raw, interpret_clsi =
        TRUE)
#> Warning: There was 1 warning in `mutate()`.
#> ℹ In argument: `pheno_provided = as.sir(`Resistance phenotype`)`.
#> Caused by warning:
#> ! in `as.sir()`: 8 results in column pheno_provided truncated (6%) that were
#> invalid antimicrobial interpretations: "intermediate"
#> Manually setting `pheno_provided` to "I" where `Resistance phenotype` was
        "intermediate"
```

##### To download genotype data

The `query_ncbi_bq_geno()` function lets you download AMRFinderPlus genotype data, for BioSamples that have matching AST data, from NCBI via Google Cloud BigQuery using the bigrquery R pacakge. You must specify a species, and can optionally limit the download to one or more specific drug classes (see NCBI AMR Class-Subclass Reference for valid terms). The function can also re-format the data into an AMRgen genotype table.

Note that to save memory and disk space BioSamples with no AST data in NCBI will not be included in the download.

Not all NCBI genotype results are updated with each new release of AMRFinderPlus, so older genomes may have genotype results obtained with older versions of AMRFinderPlus, and newer genomes will have genotype results obtained with more recent versions.

```r
# Download Staphylococcus aureus genotype data from NCBI, filtering for variants
        associated with class 'AMINOGLYCOSIDES' or 'TETRACYCLINES'
# NOTE: you may need to add 'project_id="xxx"' to the command if you have not set up
        application default credentials
staph_geno_ncbi_cloud_raw <- query_ncbi_bq_geno(
  taxgroup = "Staphylococcus aureus",
  geno_class = c("AMINOGLYCOSIDE", "TETRACYCLINE")
) %>% filter(biosample_acc %in% staph_pheno_ncbi_cloud_raw$BioSample)


staph_geno_ncbi_cloud_raw
#> # A tibble: 119 × 9
#>    biosample_acc `Gene symbol`    Class Subclass `Element type` `Element subtype`
#>    <chr>         <chr>            <chr> <chr>    <chr>          <chr>
#>  1 SAMN07291566  mepA             TETR… TIGECYC… AMR            AMR
#>  2 SAMN04901618  ant(9)-Ia        AMIN… SPECTIN… AMR            AMR
#>  3 SAMN04901605  mepA             TETR… TIGECYC… AMR            AMR
#>  4 SAMN04901606  mepA             TETR… TIGECYC… AMR            AMR
#>  5 SAMN07291567  aac(6')-Ie/aph… AMIN… AMIKACI… AMR            AMR
#>  6 SAMN07291564  mepA             TETR… TIGECYC… AMR            AMR
#>  7 SAMN04901609  tet(M)           TETR… TETRACY… AMR            AMR
#>  8 SAMN07291562  mepA             TETR… TIGECYC… AMR            AMR
#>  9 SAMN04901615  mepA             TETR… TIGECYC… AMR            AMR
#> 10 SAMN30333622  ant(9)-Ia        AMIN… SPECTIN… AMR            AMR
#> # ℹ 109 more rows
#> # ℹ 3 more variables: Method <chr>, Hierarchy_node <chr>, scientific_name <chr>

# Import and parse results into AMRgen genotype table
```

```r
staph_geno_ncbi_cloud <- import_amrfp(staph_geno_ncbi_cloud_raw, sample_col =
        "biosample_acc")

staph_geno_ncbi_cloud
#> # A tibble: 148 × 18
#>    id           marker     gene   mutation drug  drug_class `variation type` node
#>    <chr>        <chr>      <chr>  <chr>    <ab>  <chr>      <chr>            <chr>
#>  1 SAMN07291566 mepA       mepA   <NA>     TGC:… Tetracycl… Gene presence d… mepA
#>  2 SAMN04901618 ant(9)-Ia  ant(…  <NA>     SPT:… Other      Gene presence d… ant(…
#>  3 SAMN04901605 mepA       mepA   <NA>     TGC:… Tetracycl… Gene presence d… mepA
#>  4 SAMN04901606 mepA       mepA   <NA>     TGC:… Tetracycl… Gene presence d… mepA
#>  5 SAMN07291567 aac(6')-…  aac(…  <NA>     AMK:… Aminoglyc… Gene presence d… aac(…
#>  6 SAMN07291567 aac(6')-…  aac(…  <NA>     GEN:… Aminoglyc… Gene presence d… aac(…
#>  7 SAMN07291567 aac(6')-…  aac(…  <NA>     KAN:… Aminoglyc… Gene presence d… aac(…
#>  8 SAMN07291567 aac(6')-…  aac(…  <NA>     TOB:… Aminoglyc… Gene presence d… aac(…
#>  9 SAMN07291564 mepA       mepA   <NA>     TGC:… Tetracycl… Gene presence d… mepA
#> 10 SAMN04901609 tet(M)     tet(…  <NA>     NA    Tetracycl… Gene presence d… tet(…
#> # ℹ 138 more rows
#> # ℹ 10 more variables: marker.label <chr>, `Gene symbol` <chr>, Class <chr>,
#> #   Subclass <chr>, `Element type` <chr>, `Element subtype` <chr>,
#> #   Method <chr>, Hierarchy_node <chr>, scientific_name <chr>,
#> #   subclass_to_parse <chr>
```

#### Visualise the downloaded phenotype data to check the distribution of the AST data

```r
# Select one antibiotic at a time
# Specify species and guideline, to annotate with EUCAST breakpoints

# Doxycycline
staph_dox_mic_plot <- assay_by_var(
  pheno_table = staph_pheno_ncbi,
  pheno_drug = c("DOX"),
  measure = "mic",
  colour_by = "pheno_eucast",
  species = "Staphylococcus aureus",
  guideline = "EUCAST 2024"
)
#>   MIC breakpoints determined using AMR package: S <= 1 and R > 1

staph_dox_mic_plot
```

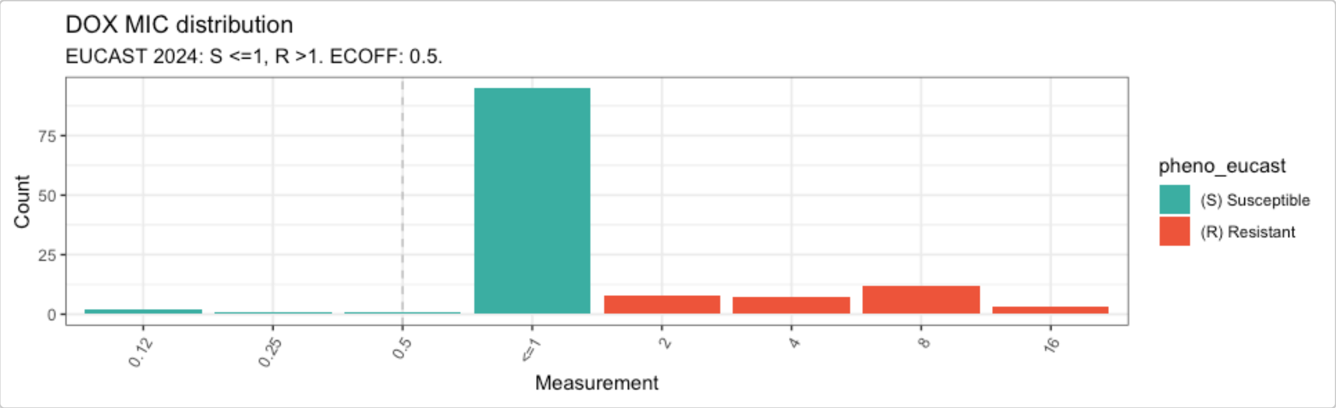

```r
# Amikacin
staph_ami_mic_plot <- assay_by_var(
  pheno_table = staph_pheno_ncbi,
  pheno_drug = c("amikacin"),
  measure = "mic",
  colour_by = "pheno_eucast",
  species = "Staphylococcus aureus",
  guideline = "EUCAST 2024"
)
#>   MIC breakpoints determined using AMR package: S <= 16 and R > 16

staph_ami_mic_plot
```

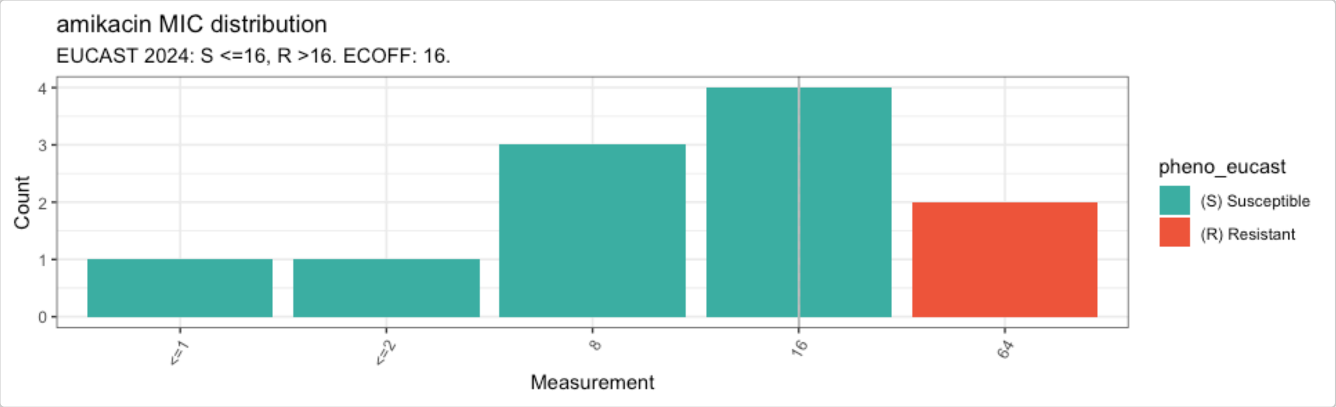


For more guidance on how to visulaise phenotypic data and combine it with genotypic data, have a look at the other `AMRgen` vignettes.

#### Option 2: Download data from EBI

The `download_ebi` function lets you retrieve phenotype or genotype data (by setting `data ="genotype'`) from the EBI AMR Portal. Genotypes are called using AMRFinderPlus but processed by EBI. You can optionally filter the downloaded file to a specified genus or species, and a specific antibiotic (for phenotype data) or NCBI class/subclass (for genotype data; check the NCBI AMR Class-Subclass Reference for valid terms). The function can also reformat and re-interpret the retrieved results with the breakpoint standards of your choosing.

Note that while NCBI and EBI AST databases have a lot of overlapping content, they are not identical and each includes biosamples that the other does not.

```r
# Download EBI phenotype data for all Staphylococcus, using the same example drugs as
        above. Reformat and re-interpret using EUCAST breakpoints
staph_pheno_ebi <- download_ebi(
  genus = "Staphylococcus",
  pheno_drug = c("amikacin", "DOX"),
  reformat = TRUE,
  interpret_eucast = TRUE,
  interpret_clsi = TRUE,
  interpret_ecoff = TRUE
) # chose which guideline to use for re-interpretation. reformat must be TRUE for re-
        interpretation


# check output
nrow(staph_pheno_ebi)
```

```
#> [1] 218

length(unique(staph_pheno_ebi$id))
#> [1] 190

head(staph_pheno_ebi)
#> # A tibble: 6 × 46
#>   id    drug    mic  disk pheno_provided pheno_eucast pheno_clsi ecoff guideline
#>   <chr> <ab>  <mic> <dsk> <sir>          <sir>        <sir>      <sir> <chr>
#> 1 SAME… AMK:…    NA    NA R              NA           NA         NA    <NA>
#> 2 SAME… AMK:…    NA    NA R              NA           NA         NA    <NA>
#> 3 SAME… AMK:…    NA    NA R              NA           NA         NA    <NA>
#> 4 SAME… AMK:…    NA    NA R              NA           NA         NA    <NA>
#> 5 SAME… AMK:…    NA    NA R              NA           NA         NA    <NA>
#> 6 SAME… AMK:…    NA    NA R              NA           NA         NA    <NA>
#> # ℹ 37 more variables: method <chr>, platform <chr>, source <chr>,
#> #   spp_pheno <mo>, SRA_accession <chr>, assembly_ID <chr>,
#> #   collection_year <int>, ISO_country_code <chr>, host <chr>, host_age <chr>,
#> #   host_sex <chr>, isolate <chr>, isolation_source <chr>,
#> #   isolation_source_category <chr>, isolation_latitude <chr>,
#> #   isolation_longitude <chr>, genus <chr>, organism <chr>,
#> #   Updated_phenotype_CLSI <chr>, Updated_phenotype_EUCAST <chr>, …
```

Genotype data can be retrieved from EBI using the same function. Currently, not all samples in the EBI AMR portal have both AST and genotype data.

Note the input assemblies used to call genotypes, and the version of AMRFinderPlus, in the EBI portal can be different from what's available for download from NCBI using the above functions.

```
# Download genotype data for Staphylococcus, filter to markers associated with
        aminoglycosides or tetracyclines, and re-format the data into an AMRgen genotype
        table. Note that not all samples with phenotype data have genotype data.
staph_geno_ebi <- download_ebi(
  data = "genotype",
  genus = "Staphylococcus",
  geno_class = c("AMINOGLYCOSIDE", "TETRACYCLINE"),
  reformat = T
)


nrow(staph_geno_ebi)
#> [1] 45945

length(unique(staph_geno_ebi$id))
#> [1] 7547

head(staph_geno_ebi)
#> # A tibble: 6 × 34
#>   id        marker gene  mutation drug  drug_class marker.label assembly_ID genus
#>   <chr>     <chr>  <chr> <chr>    <ab>  <chr>      <chr>        <chr>       <chr>
#> 1 SAMEA53… tet(3… tet(… -        NA    Tetracycl… tet(38)      ERZ25282410 Stap…
#> 2 SAMEA53… tet(K) tet(… -        NA    Tetracycl… tet(K)       ERZ25282410 Stap…
#> 3 SAMEA53… mepA   mepA  -        TGC:… Tetracycl… mepA         ERZ25282410 Stap…
#> 4 SAMEA53… tet(3… tet(… -        NA    Tetracycl… tet(38)      ERZ25282723 Stap…
#> 5 SAMEA53… mepA   mepA  -        TGC:… Tetracycl… mepA         ERZ25282723 Stap…
#> 6 SAMEA53… tet(K) tet(… -        NA    Tetracycl… tet(K)       ERZ25282853 Stap…
```

```
#> # ℹ 25 more variables: species <chr>, organism <chr>, isolate <chr>,
#> #   taxon_id <int>, region <chr>, region_start <int>, region_end <int>,
#> #   strand <chr>, `_bin` <int>, id2 <chr>, gene_symbol <chr>,
#> #   amr_element_symbol <chr>, element_type <chr>, element_subtype <chr>,
#> #   class <chr>, subclass <chr>, split_subclass <chr>, antibiotic_name <chr>,
#> #   antibiotic_ontology <chr>, antibiotic_ontology_link <chr>,
#> #   evidence_accession <chr>, evidence_type <chr>, evidence_link <chr>, …
```

#### Compare downloaded phenotypes and genotypes

The downloaded phenotype and genotype data for a specified antibiotic can then be extracted and combined using the `get_binary_matrix` function.

```
# first filter both EBI pheno and geno dataframes for Staph aureus only
# filter pheno data
staph_pheno_ebi_filtered <- staph_pheno_ebi %>%
  filter(organism == "Staphylococcus aureus")

# filter geno data
staph_geno_ebi_filtered <- staph_geno_ebi %>%
  filter(species == "Staphylococcus aureus")

# Make binary geno-pheno matrix for doxycycline phenotype (re-interpreted with EUCAST),
        and genotypes associated with the associated drug class (Tetracyclines)
tet_bin <- get_binary_matrix(
  geno_table = staph_geno_ebi_filtered,
  pheno_table = staph_pheno_ebi_filtered,
  pheno_drug = "DOX",
  geno_class = "Tetracyclines", # matches drug_class in geno_table
  sir_col = "pheno_eucast", # phenotype column in pheno_table
  keep_assay_values = TRUE,
  keep_assay_values_from = "mic"
)
#>  Some samples had multiple phenotype rows, taking the most resistant only for binary
        matrix
#>  Defining NWT in binary matrix using ecoff column provided: ecoff

nrow(tet_bin)
#> [1] 116

head(tet_bin)
#> # A tibble: 6 × 11
#>    id     pheno ecoff   mic     R   NWT  mepA `tet(38)` `tet(K)` `tet(M)` `tet(L)`
#>    <chr> <sir> <sir> <mic> <dbl> <dbl> <dbl>     <dbl>    <dbl>    <dbl>    <dbl>
#> 1 SAMN… S     NI     <=1     0    NA     1         1        0        0        0
#> 2 SAMN… S     NI     <=1     0    NA     1         1        0        0        0
#> 3 SAMN… S     NI     <=1     0    NA     1         1        0        0        0
#> 4 SAMN… S     NI     <=1     0    NA     1         1        1        0        0
#> 5 SAMN… R     NWT      8     1     1     1         1        0        1        0
#> 6 SAMN… S     NI     <=1     0    NA     1         1        0        0        0


# plot positive predictive value for each marker/combination
tet_ppv <- amr_ppv(tet_bin)
```

```
#> Removing 69 rows with no phenotype call
#> Ordering markers by frequency
```

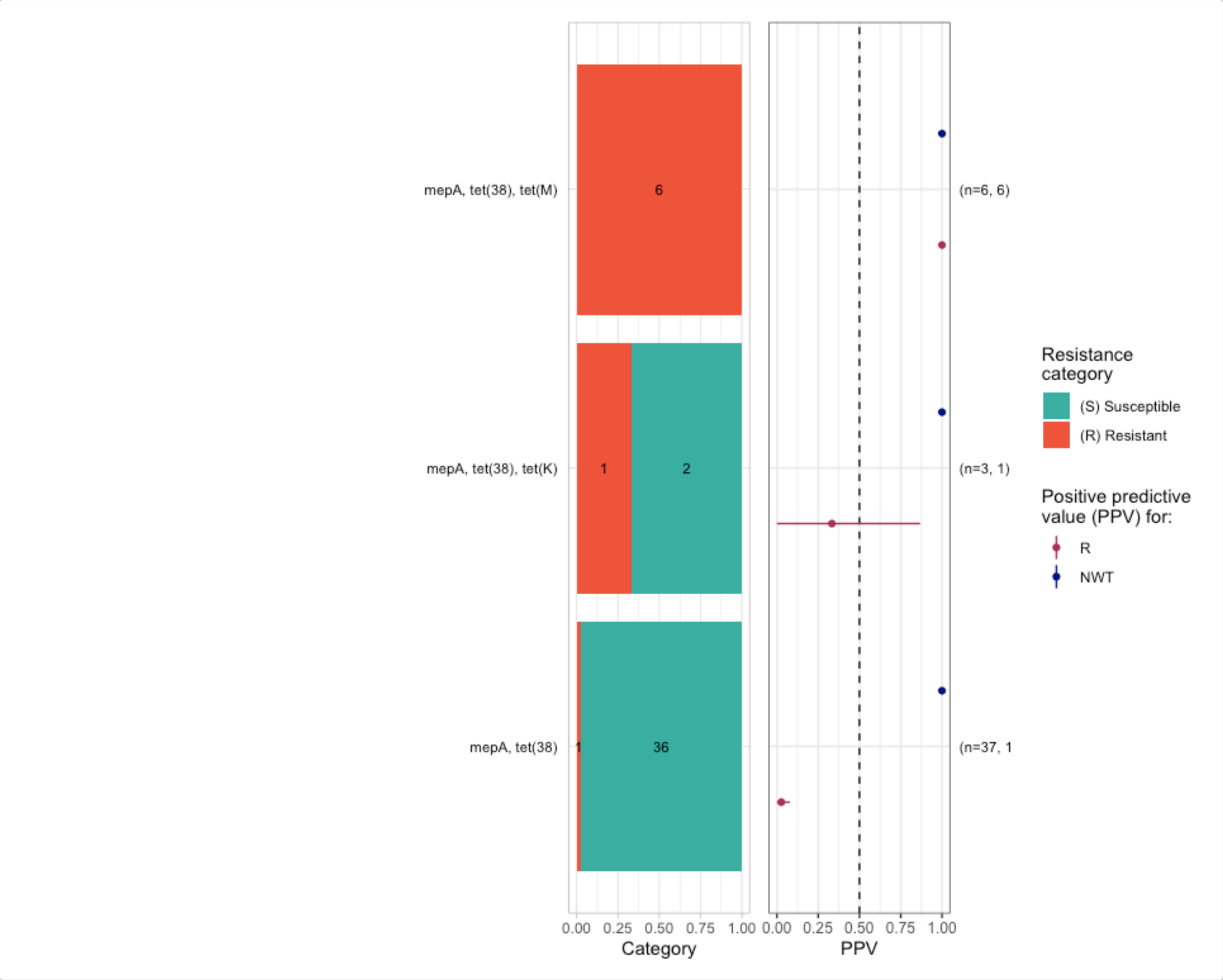


```
tet_ppv$plot
```

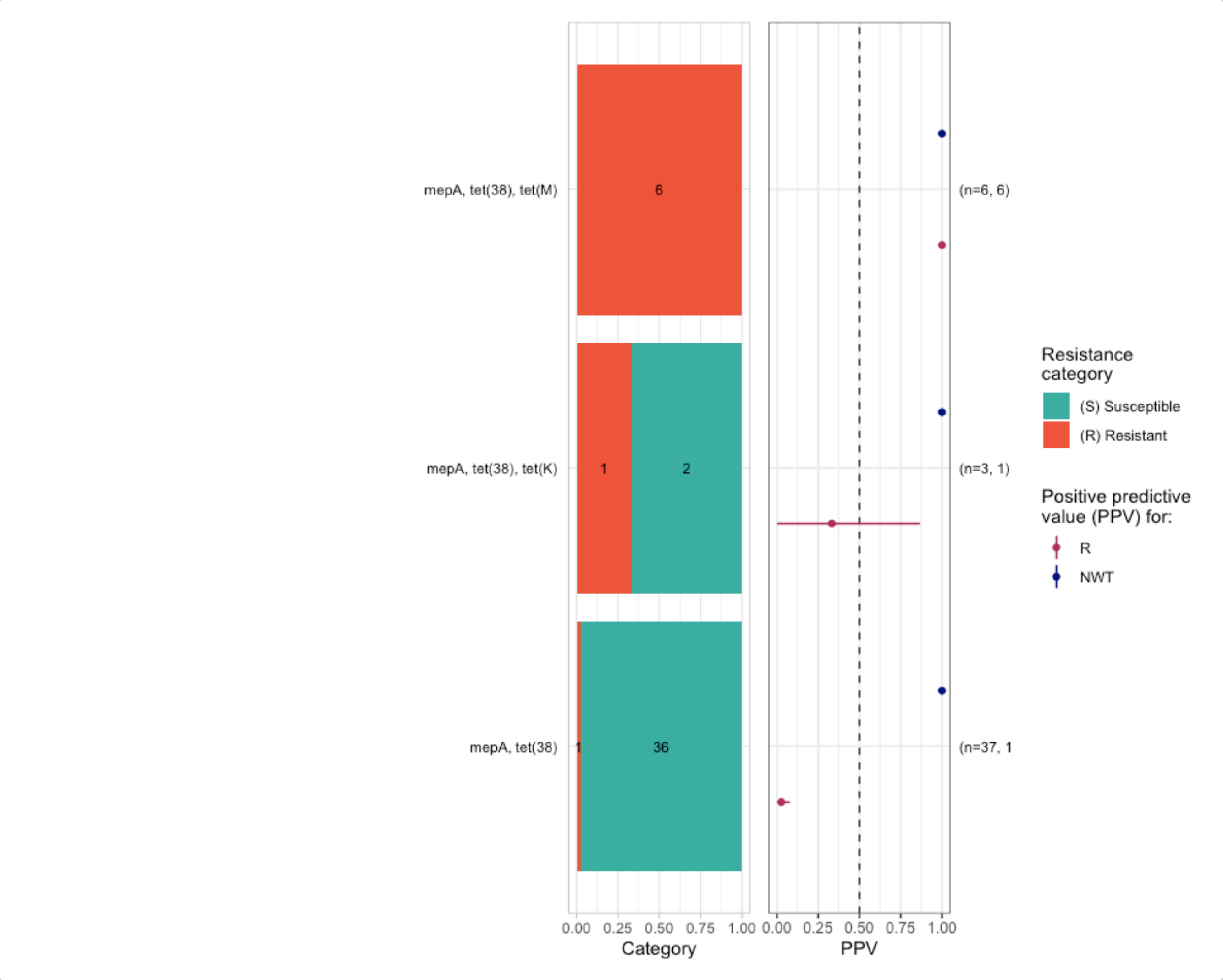


This produces a binary matrix with 1 row per BioSample, for all BioSamples that had both phenotype data for doxycycline in `staph_pheno_ebi_filtered` AND any genotype data (associated with any marker, not just Tetracyclines) in `staph_geno_ebi_filtered`. This ensures that we include samples for which genotyping was performed but returned no hits associated with Tetracyclines, not just those samples that had tetracycline associated markers detected (note that such samples would be missing if we had only downloaded genotype data associated with Tetracyclines).

Columns indicate the input phenotype columns for doxycycline (renamed to standard fields `pheno`, `ecoff`, `mic`), and binary indicators (1=present, 0=absent) for doxycycline phenotypes (`R`, `NWT`) and each genotype marker associated with tetracyclines (in this case `tet(38)`, `tet(K)`, `tet(M)`,`tet(L)`).

For more examples on how to do join geno-pheno analyses and visualisations, so the other AMRgen Vignettes.

### Assessing geno-pheno concordance

### Introduction

`AMRgen` is a comprehensive R package designed to integrate antimicrobial resistance genotype and phenotype data. It provides tools to import AMR genotype data, AST phenotype data, and conduct genotype-phenotype analyses to explore the impact of genotypic markers on phenotype, including phenotype-genotype concordance.

The `concordance()` function in AMRgen compares genotypes (presence of resistance markers) to observed phenotypes (resistant vs susceptible) using a binary matrix obtained with `get_binary_matrix()`. A genotypic prediction variable is defined on the basis of presence of genetic resistance markers, either all markers in the input table or those defined by an input inclusion list or exclusion list (specific marker(s), a minimum number of markers). The user may also filter the markers to be included in the genotypic prediction based on thresholds for solo Positive Predictive Value (PPV, see `solo_ppv()` function) or logistic regression p-values (see `amr_logistic()` function.

This genotypic prediction (the "test") is then compared to the observed phenotypes (the "truth" or "gold standard") using standard classification metrics calculated with the yardstick package (https://yardstick.tidymodels.org/reference/index.html). These are Sensitivity, Specificity, PPV, Negative Predictive Value (NPV), Accuracy, Kappa, and F-measure. Error rates (major error - ME, and very major error - VME) are calculated as per ISO 20776-2 (see FDA definitions). The `concordance()` function supports evaluating both R and NWT outcomes in a single call, with flexible prediction rules and marker inclusion options.

This vignette walks through a workflow to analyse phenotype-genotype concordance using example datasets taken from "A one-year genomic investigation of *Escherichia coli* epidemiology and nosocomial spread at a large US healthcare network" by Mills et al (2022). The antimicrobial susceptibility test results (MIC values and SIR interpretation) were obtained from the EBI AMR portal (https://www.ebi.ac.uk/amr/), and the AMRFinderPlus results from the All the Bacteria project (https://allthebacteria.org/).

Citation: Mills, E.G., Martin, M.J., Luo, T.L. *et al*. A one-year genomic investigation of *Escherichia coli* epidemiology and nosocomial spread at a large US healthcare network. Genome Med 14, 147 (2022). https://doi.org/10.1186/s13073-022-01150-7

#### Start by loading the AMRgen package:

```
# Load AMRgen
library(AMRgen)

# Also load the dplyr package to use the filter function in step 7
# https://dplyr.tidyverse.org/reference/filter.html
library(dplyr)
```

#### 1. Phenotype table

The antimicrobial susceptibility test (AST) data for the 2075 isolates in this study were retrieved from the EBI AMR Portal FTP site, following these steps:

1. Downloaded all E. coli phenotype data with the `download_ebi()` function from AMRgen, passing options `species="Escherichia coli"`, `release = "2025-12"`, and `reformat = TRUE`. Options to reinterpret data based on CLSI/EUCAST breakpoints or ECOFFs were not applied.
2. Downloaded the table with accessions associated with this publication (bioproject PRJNA809394) from the SRA (https://www.ncbi.nlm.nih.gov/Traces/study/?acc=SRP401320&o=acc_s%3Aa). Note: some manual curation was needed as this file contains accessions for 2076 samples.
3. Programmatically selected the AST data for the 2075 E. coli by the "id" column using the "BioSample" column in the accessions table.

The resulting phenotype table was imported to the AMRgen package and called `pheno_eco_2075`.

```
# Check the format of the phenotype table pre-loaded in the AMRgen package
data(pheno_eco_2075)

head(pheno_eco_2075)
#> # A tibble: 6 × 36
#>   id    drug    mic pheno_provided guideline method platform source spp_pheno
#>   <chr> <ab>  <mic> <sir>          <chr>     <chr>  <chr>    <lgl>  <mo>
#> 1 SAMN… AMK:…   <=8 S              CLSI      broth… BD Phoe… NA     B_ESCHR_COL…
#> 2 SAMN… GEN:…   <=2 S              CLSI      broth… BD Phoe… NA     B_ESCHR_COL…
#> 3 SAMN… TOB:…   <=2 S              CLSI      broth… BD Phoe… NA     B_ESCHR_COL…
#> 4 SAMN… AMP:…   <=4 S              CLSI      broth… BD Phoe… NA     B_ESCHR_COL…
#> 5 SAMN… AMC:…     8 S              CLSI      broth… BD Phoe… NA     B_ESCHR_COL…
#> 6 SAMN… TZP:…   <=2 S              CLSI      broth… BD Phoe… NA     B_ESCHR_COL…
#> # ℹ 27 more variables: SRA_accession <chr>, assembly_ID <chr>,
#> #   collection_year <dbl>, ISO_country_code <chr>, host <chr>, host_age <lgl>,
#> #   host_sex <lgl>, isolate <dbl>, isolation_source <chr>,
#> #   isolation_source_category <chr>, isolation_latitude <dbl>,
#> #   isolation_longitude <dbl>, genus <chr>, organism <chr>,
#> #   Updated_phenotype_CLSI <chr>, Updated_phenotype_EUCAST <chr>,
#> #   used_ECOFF <chr>, database <chr>, measurement <chr>, …
```

The phenotype table has one row for each assay measurement, i.e. one per strain/drug combination. The essential columns for a phenotype table to work with `AMRgen` functions are:

- `id`: character string giving the sample name, used to link to sample names in the genotype file
- `spp_pheno`: species in the form of an AMR package `mo` class, in this vignette "B_ESCHR_COLI"
- `drug`: antibiotic name in the form of an AMR package `ab` class
- a phenotype column: S/I/R phenotype calls in the form of an AMR package `sir` class. In this example, SIR phenotype calls are as provided in the EBI AMR portal (`pheno_provided`) as we did not re-interpret data based on CLSI/EUCAST breakpoints during download.

This vignette also uses raw MIC data for the analyses. The corresponding column is:

- `mic`: MIC measurements in the form of an AMR package `mic` class.

#### 2. Genotype table

The AMRFinderPlus results for the 2075 isolates in this study were retrieved from the AllTheBacteria project, following these steps:

1. Downloaded a compressed TSV file containing the aggregated results of running AMRFinderPlus on all samples in the AllTheBacteria dataset from https://osf.io/ck7st (large file)

2. Programmatically selected the results for the 2075 *E. coli* samples by the "Name" column using the "BioSample" column in the accessions table downloaded from the SRA (as for the phenotype table).

```
# Load AMRFinderPlus data to create an object with the key columns needed to work with the
        AMRgen package
data(geno_eco_2075)
geno_eco_2075 <- import_amrfp(geno_eco_2075, "Name")

# Check the format of the processed genotype table
head(geno_eco_2075)
#> # A tibble: 6 × 33
#>   id           marker     gene  mutation drug  drug_class `variation type` node
#>   <chr>        <chr>      <chr> <chr>    <ab>  <chr>      <chr>            <chr>
#> 1 SAMN26304318 pmrB_Y358N pmrB  Tyr358A… COL:… Polymyxins Protein variant… pmrB
#> 2 SAMN26304318 blaEC      blaEC <NA>     NA    Beta-lact… Gene presence d… blaEC
#> 3 SAMN26304318 mdtM       mdtM  <NA>     NA    Efflux     Gene presence d… mdtM
#> 4 SAMN26304318 glpT_E448K glpT  Glu448L… FOS:… Phosphoni… Protein variant… glpT
#> 5 SAMN26304318 acrF       acrF  <NA>     NA    Efflux     Gene presence d… acrF
#> 6 SAMN26304319 blaEC      blaEC <NA>     NA    Beta-lact… Gene presence d… blaEC
#> # ℹ 25 more variables: marker.label <chr>, `Protein identifier` <lgl>,
#> #   `Contig id` <chr>, Start <dbl>, Stop <dbl>, Strand <chr>,
#> #   `Gene symbol` <chr>, `Sequence name` <chr>, Scope <chr>,
#> #   `Element type` <chr>, `Element subtype` <chr>, Class <chr>, Subclass <chr>,
#> #   Method <chr>, `Target length` <dbl>, `Reference sequence length` <dbl>,
#> #   `% Coverage of reference sequence` <dbl>,
#> #   `% Identity to reference sequence` <dbl>, `Alignment length` <dbl>, …
```

The genotype table has one row for each genetic marker detected in an input genome, i.e. one per strain/marker combination. The essential columns for a genotype table to work with `AMRgen` functions are:

- `Name`: character string giving the sample name, used to link to sample names in the phenotype file.
- `marker`: character string giving the name of the genetic marker detected.
- `drug_class`: character string giving the antibiotic class associated with this marker.

NOTE: In this example, at least one AMR marker is present in all 2075 genomes. In contrast, no markers were reported for five genomes in the publication (see Supplementary Table 3).

#### 3. Combine genotype and phenotype data for Ciprofloxacin

The phenotype table includes data for 18 antibiotics from 11 different classes, but we need to analyse concordance one drug at a time.

The function `get_binary_matrix()` is used to extract phenotype data for a specified drug (in this example Ciprofloxacin), and genotype data for markers associated with a specified drug class by AMRFinderPlus (in this example Quinolones). It returns a single dataframe with one row per strain, for the subset of strains that appear in both the genotype and phenotype input tables.

```
# Get matrix combining phenotype data for Ciprofloxacin, binary calls for R/NWT pheno,
# and genotype presence/absence data for all markers associated with Quinolone
eco_cip_matrix <- get_binary_matrix(
  geno_eco_2075,
```

```r
  pheno_eco_2075,
  pheno_drug = "Ciprofloxacin",
  geno_class = "Quinolones",
  sir_col = "pheno_provided",
  keep_assay_values = TRUE,
  keep_assay_values_from = "mic"
)
#>  Defining NWT in binary matrix as I/R vs S, as no ECOFF column defined

# Check the format of the binary matrix
head(eco_cip_matrix)
#> # A tibble: 6 × 36
#>   id           pheno   mic      R   NWT gyrA_D87N gyrA_S83L parC_S80I parE_S458A
#>   <chr>        <sir> <mic> <dbl> <dbl>     <dbl>     <dbl>     <dbl>      <dbl>
#> 1 SAMN26304318 S     <=0.5     0     0         0         0         0          0
#> 2 SAMN26304319 R      >2.0     1     1         1         1         1          1
#> 3 SAMN26304320 S     <=0.5     0     0         0         0         0          0
#> 4 SAMN26304321 S     <=0.5     0     0         0         0         0          0
#> 5 SAMN26304322 S     <=0.5     0     0         0         0         0          0
#> 6 SAMN26304323 S     <=0.5     0     0         0         0         0          0
#> # ℹ 27 more variables: marR_S3N <dbl>, qnrS1 <dbl>, parE_I529L <dbl>,
#> #   parC_E84V <dbl>, qnrB19 <dbl>, parE_L416F <dbl>, parC_S80R <dbl>,
#> #   `aac(6')-Ib-cr5` <dbl>, parE_S458T <dbl>, parE_D475E <dbl>,
#> #   parC_S57T <dbl>, parE_I355T <dbl>, gyrA_S83A <dbl>, parC_E84G <dbl>,
#> #   qnrB2 <dbl>, marR_R77C <dbl>, qnrB6 <dbl>, gyrA_D87Y <dbl>,
#> #   parE_L445H <dbl>, gyrA_D87G <dbl>, qnrB4 <dbl>, soxS_A12S <dbl>,
#> #   parE_I464F <dbl>, parE_E460D <dbl>, qnrB <dbl>, soxR_G121D <dbl>, …
```

Each row in the binary matrix indicates, for one strain, both the phenotypes (with SIR column, mic values, and boolean 1/0 coding of R and NWT status) and the genotypes (one column per marker, with boolean 1/0 coding of marker presence/absence).

The ‘NWT’ variable can be taken either from a precomputed ECOFF-based call of WT=wildtype/NWT=nonwildtype (if passing the option `ecoff_col`), or computed from the S/I/R phenotype as NWT=R/I and WT=S. In this example, NWT was defined as R/I vs S. No ECOFF column was defined because of the nature of the AST data: the minimum MIC values of <=0.5 in the BD Phoenix AST data are above the ECOFF of 0.064 and cannot be interpreted against ECOFF. We can inspect the distribution of the Ciprofloxacin phenotype data to better understand this.

#### 4. Plot Ciprofloxacin phenotype data distribution

The function `assay_by_var()` can be used to plot the distribution of MIC values coloured by a variable. In this case, the S/I/R values were coloured by the column “pheno_provided”, which were interpreted by the authors with the breakpoints from CLSI 2018. We can also compare them to the updated breakpoints from CLSI 2025.

```r
# CLSI 2018 guidelines (as in the publication from Mills et al).
# The breakpoints are provided manually.
assay_by_var(
  pheno_table = pheno_eco_2075,
  pheno_drug = "Ciprofloxacin",
  measure = "mic",
  colour_by = "pheno_provided",
  species = "Escherichia coli",
```

```
  bp_S = 1,
  bp_R = 4
)
```

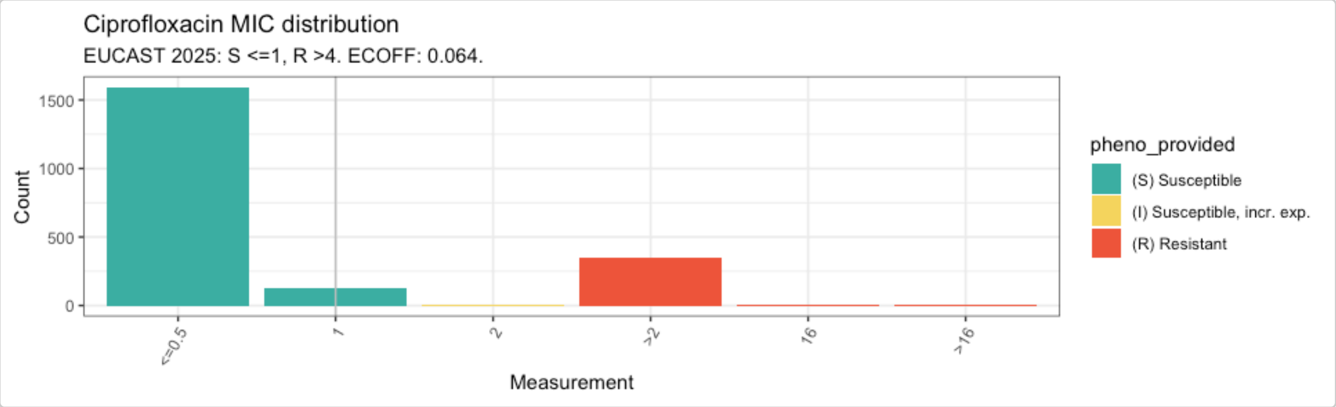


```
# CLSI 2025 guidelines
# The breakpoints are provided by passing the option "guideline"
assay_by_var(
  pheno_table = pheno_eco_2075,
  pheno_drug = "Ciprofloxacin",
  measure = "mic",
  colour_by = "pheno_provided",
  species = "Escherichia coli",
  guideline = "CLSI 2025"
)
#>   MIC breakpoints determined using AMR package: S <= 0.25 and R > 1
```

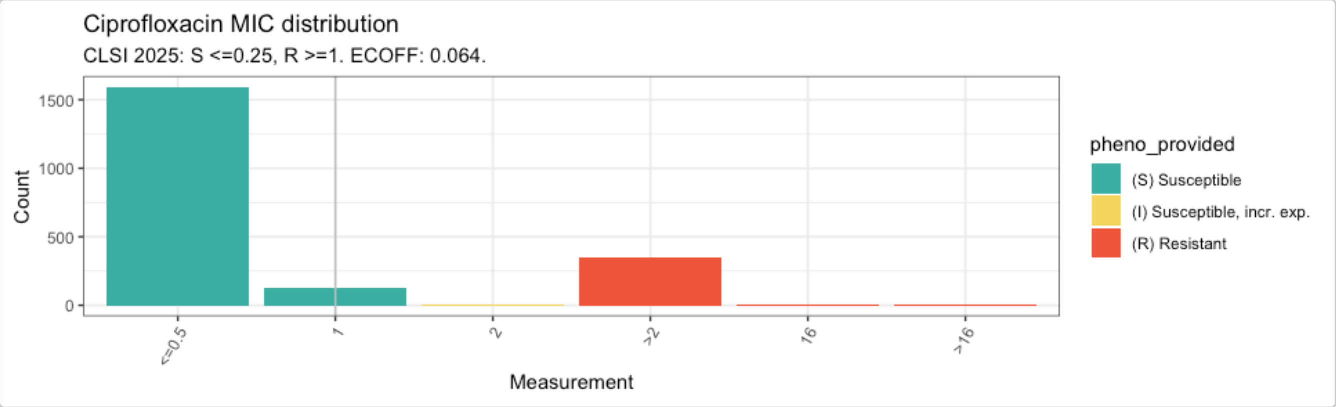


The plots of the distribution of MIC values with breakpoints for Ciprofloxacin from the 2018 CLSI guidelines (S<=1 and R>=4) vs 2025 CLSI guidelines (S<=0.25 and R>=1) confirm that the SIR interpretation was as per the 2018 breakpoints, and that the interpretation would have been different if the AST data had been re-interpreted during download by passing the option `interpret_clsi=TRUE` to the `download_ebi()` function.

The plots also show that the ECOFF value of 0.064 is below the minimum MIC values in the distribution.

#### 5. Calculate concordance between phenotype and genotype

Start with a first concordance calculation including all the AMR markers in the binary matrix.

```r
concordance_cip <- concordance(eco_cip_matrix)

concordance_cip
#> AMR Genotype-Phenotype Concordance
#> Prediction rule: any
#>
#> --- Outcome: R ---
#> Samples: 2075 | Markers: 31
#> Markers used: gyrA_D87N, gyrA_S83L, parC_S80I, parE_S458A, marR_S3N, qnrS1, parE_I529L,
        parC_E84V, qnrB19, parE_L416F, parC_S80R, aac(6')-Ib-cr5, parE_S458T, parE_D475E,
        parC_S57T, parE_I355T, gyrA_S83A, parC_E84G, qnrB2, marR_R77C, qnrB6, gyrA_D87Y,
        parE_L445H, gyrA_D87G, qnrB4, soxS_A12S, parE_I464F, parE_E460D, qnrB,
        soxR_G121D, parC_A56T
#>
#> Confusion Matrix:
#>           Truth
#> Prediction   1   0
#>          1 338 891
#>          0   9 837
#>
#> Metrics:
#>   Sensitivity : 0.9741
#>   Specificity : 0.4844
#>   PPV         : 0.2750
#>   NPV         : 0.9894
#>   Accuracy    : 0.5663
#>   Kappa       : 0.2274
#>   F-measure   : 0.4289
#>   VME         : 0.0259
#>   ME          : 0.5156
#>
#> --- Outcome: NWT ---
#> Samples: 2075 | Markers: 31
#> Markers used: gyrA_D87N, gyrA_S83L, parC_S80I, parE_S458A, marR_S3N, qnrS1, parE_I529L,
        parC_E84V, qnrB19, parE_L416F, parC_S80R, aac(6')-Ib-cr5, parE_S458T, parE_D475E,
        parC_S57T, parE_I355T, gyrA_S83A, parC_E84G, qnrB2, marR_R77C, qnrB6, gyrA_D87Y,
        parE_L445H, gyrA_D87G, qnrB4, soxS_A12S, parE_I464F, parE_E460D, qnrB,
        soxR_G121D, parC_A56T
#>
#> Confusion Matrix:
#>           Truth
#> Prediction   1   0
#>          1 347 882
#>          0   9 837
#>
#> Metrics:
#>   Sensitivity : 0.9747
#>   Specificity : 0.4869
#>   PPV         : 0.2823
#>   NPV         : 0.9894
#>   Accuracy    : 0.5706
#>   Kappa       : 0.2341
#>   F-measure   : 0.4379
#>   VME         : 0.0253
#>   ME          : 0.5131
```

The output shows that 31 markers were identified linked to Ciprofloxacin resistance in this dataset.

The 2x2 confusion matrix shows the following values:

- 338 - no. of true positives (TP), resistant isolates predicted to be resistant by the presence of AMR markers
- 891 - no. of false positives (FP), susceptible isolates predicted to be resistant by the presence of AMR markers
- 837 - no. of true negatives (TN), susceptible isolates predicted to be susceptible by the absence of resistance markers
- 9 - no. of false negatives (FN), resistant isolates predicted to be susceptible by the absence of resistance markers

The sensitivity value (or true positive rate, i.e. the proportion of resistant isolates predicted to be resistant by the presence of AMR markers) is high (>0.95), but the specificity (or true negative rate, i.e. the proportion of susceptible isolates predicted to be susceptible by the absence of resistance markers) is very low (<0.5). The high sensitivity is especially important as predicting a resistant strain as susceptible is more consequential to treatment than finding resistance genes in phenotypically susceptible isolates. This is also reflected in the low Very Major Error (VME) vs the high Major Error (ME).

The PPV, i.e. the proportion of all positive results (true plus false) that are TP, is low. This is due to the higher number of FP (891) than of TP (331), i.e. many Cip-susceptible isolates carry resistance markers. Conversely, the NPV, i.e. the proportion of all negative results (true plus false) that are TN, is high. This is because the number of TN (837) is much higher than the number of false negatives (9).

The concordance stats for outcome R (R vs S/I) are very similar than for outcome NWT (R/I vs S). This can be explained by the small number of isolates interpreted as "I" in this dataset (n=9).

The low specificity value (caused by the 891 Cip-susceptible isolates that carry resistance markers) is not surprising as Ciprofloxacin resistance usually results from the accumulation of mutations, with isolates carrying only one mutation often remaining susceptible. This can be easily visualised with the `amr_upset()` function in `AMRgen`.

#### 6. Compare the presence of markers with susceptibility testing data with an upset plot

The function `amr_upset()` takes the binary matrix table `eco_cip_matrix`, and explores the distribution of MIC assay values for all observed combinations of markers (solo or multiple markers). The resulting upset plot shows the distribution of assay values and S/I/R calls for each observed marker combination, and returns a summary of these distributions (including sample size, median and interquartile range, number and proportion classified as R).

```
# Generate an upset plot comparing ciprofloxacin MIC data with quinolone marker
        combinations,
eco_cip_upset <- amr_upset(
  eco_cip_matrix,
  assay = "mic",
  order = "value"
)
#> Ordering markers by frequency
#> Scale for y is already present.
#> Adding another scale for y, which will replace the existing scale.
#> Scale for y is already present.
#> Adding another scale for y, which will replace the existing scale.
```

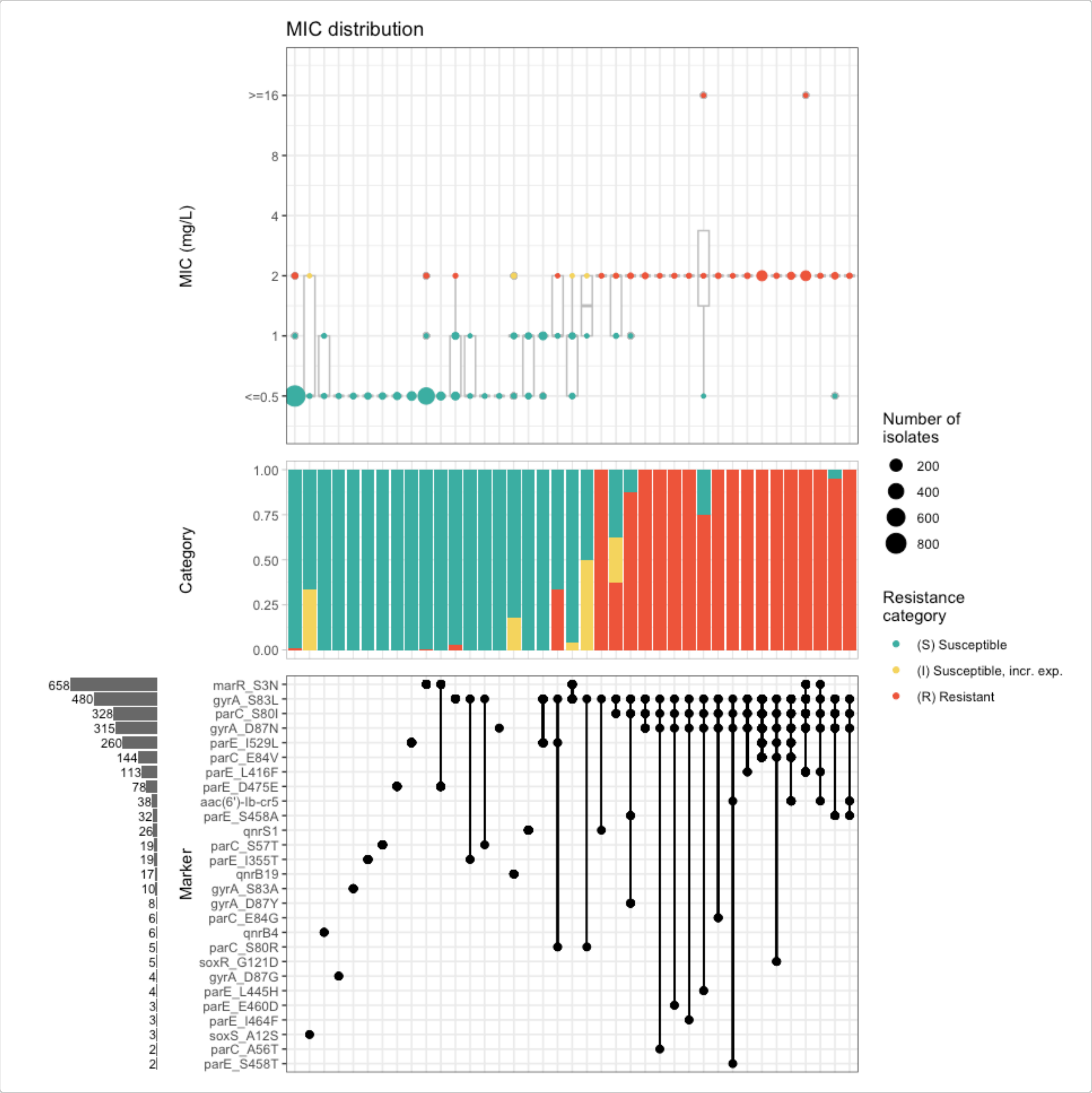


The upset plot shows that the combination of mutations in the Quinolone Resistance Determining Region (QRDR) of the *E. coli* DNA topoisomerase GyrA (gyrA_S83L) and DNA gyrase ParC (parC_S80I), alone or with other mutations, raises the MIC values above the susceptible range, i.e. the combination of this mutations is common in non-susceptpible isolates (I/R).

To identify the combinations of AMR markers that are associated with resistance, we can use the `amr_ppv()` function of the `AMRgen` package

#### 7. Identify markers or combination of markers associated with resistance

The `amr_ppv()` function calculates the possible combinations of markers, and returns the positive predictive value (PPV) for each combination (with 95% CI) and the basic plot elements (including PPV).

```
# Generate a summary plot of PPV for each solo and combination of markers observed in the
        mic assay data and order by decreasing ppv value
eco_cip_ppv <- amr_ppv(eco_cip_matrix,
  assay = "mic",
  order = "ppv"
```

```
)
#> Ordering markers by frequency
```

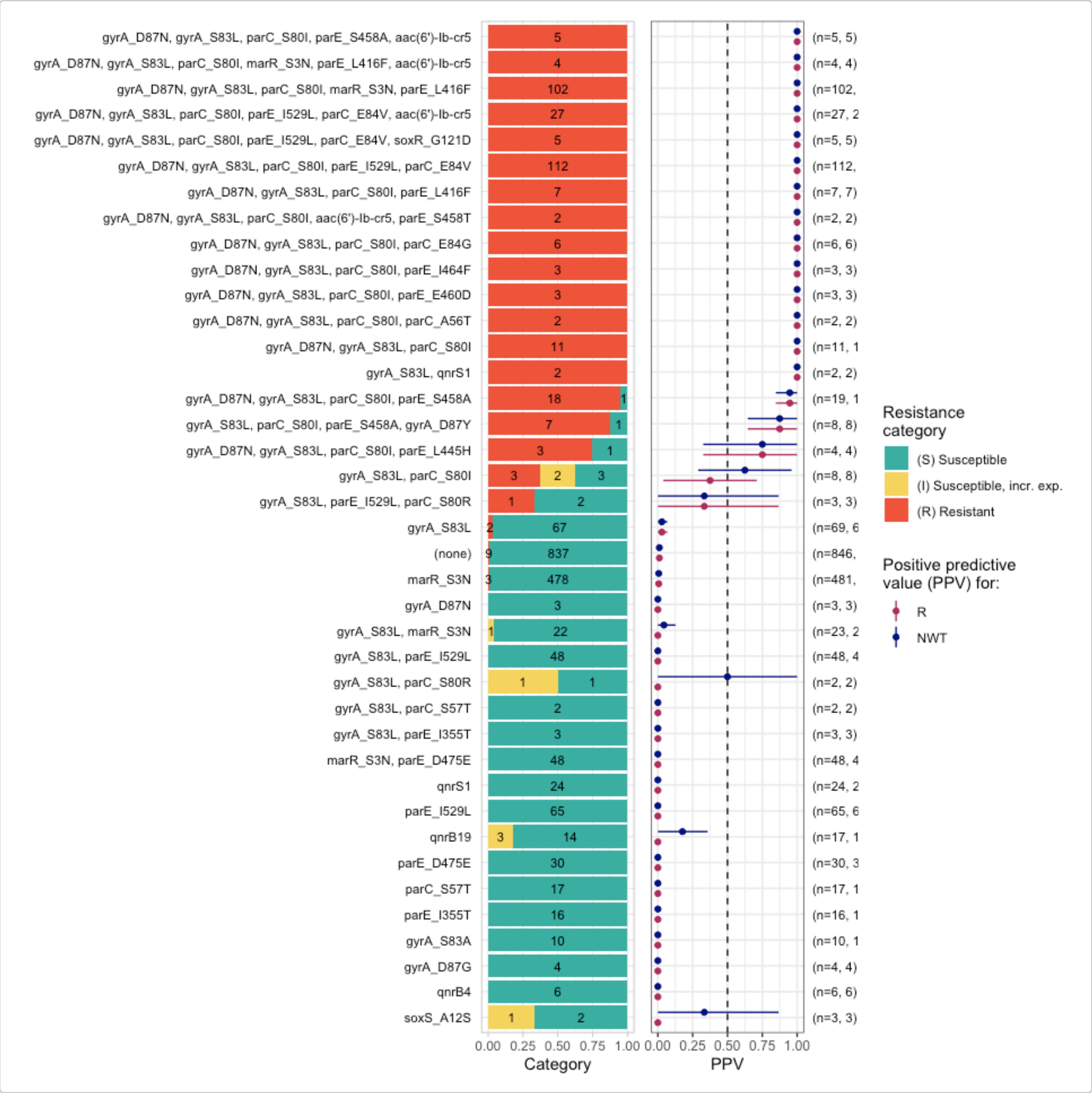


```
# View the column headers of the ppv stats
head(eco_cip_ppv$summary)
#> # A tibble: 6 × 21
#>   marker_list marker_count     n combination_id            R.n  R.ppv R.ci_lower
#>   <chr>              <dbl> <int> <fct>                   <dbl>  <dbl>      <dbl>
#> 1 ""                     0   846 0_0_0_0_0_0_0_0_0_0_0_…     9 0.0106    0.00373
#> 2 "qnrB"                 1     1 0_0_0_0_0_0_0_0_0_0_0_…     0 0          0
#> 3 "soxS_A12S"            1     3 0_0_0_0_0_0_0_0_0_0_0_…     0 0          0
#> 4 "qnrB4"                1     6 0_0_0_0_0_0_0_0_0_0_0_…     0 0          0
#> 5 "gyrA_D87G"            1     4 0_0_0_0_0_0_0_0_0_0_0_…     0 0          0
#> 6 "gyrA_D87Y"            1     1 0_0_0_0_0_0_0_0_0_0_0_…     0 0          0
#> # ℹ 14 more variables: R.ci_upper <dbl>, R.denom <int>, NWT.n <dbl>,
#> #   NWT.ppv <dbl>, NWT.ci_lower <dbl>, NWT.ci_upper <dbl>, NWT.denom <int>,
#> #   median_excludeRangeValues <dbl>, q25_excludeRangeValues <dbl>,
```

```
#> #   q75_excludeRangeValues <dbl>, n_excludeRangeValues <int>,
#> #   median_ignoreRanges <dbl>, q25_ignoreRanges <dbl>, q75_ignoreRanges <dbl>

# Select only combinations with a R ppv value of at least 0.5
ppv_05 <- eco_cip_ppv$summary %>%
  filter(R.ppv >= 0.5)

# View the combinations of markers with a ppv value above 0.5
ppv_05
#> # A tibble: 27 × 21
#>    marker_list          marker_count     n combination_id   R.n R.ppv R.ci_lower
#>    <chr>                       <dbl> <int> <fct>          <dbl> <dbl>      <dbl>
#>  1 aac(6')-Ib-cr5, par…            3     1 0_0_0_0_0_0_0…     1 1          1
#>  2 gyrA_S83L, qnrS1                2     2 0_1_0_0_0_1_0…     2 1          1
#>  3 gyrA_S83L, parC_S80…            3     1 0_1_1_0_0_0_0…     1 1          1
#>  4 gyrA_S83L, parC_S80…            3     1 0_1_1_0_0_0_0…     1 1          1
#>  5 gyrA_S83L, parC_S80…            4     8 0_1_1_1_0_0_0…     7 0.875      0.646
#>  6 gyrA_D87N, gyrA_S83…            3    11 1_1_1_0_0_0_0…    11 1          1
#>  7 gyrA_D87N, gyrA_S83…            4     2 1_1_1_0_0_0_0…     2 1          1
#>  8 gyrA_D87N, gyrA_S83…            4     3 1_1_1_0_0_0_0…     3 1          1
#>  9 gyrA_D87N, gyrA_S83…            4     3 1_1_1_0_0_0_0…     3 1          1
#> 10 gyrA_D87N, gyrA_S83…            4     4 1_1_1_0_0_0_0…     3 0.75       0.326
#> # ℹ 17 more rows
#> # ℹ 14 more variables: R.ci_upper <dbl>, R.denom <int>, NWT.n <dbl>,
#> #   NWT.ppv <dbl>, NWT.ci_lower <dbl>, NWT.ci_upper <dbl>, NWT.denom <int>,
#> #   median_excludeRangeValues <dbl>, q25_excludeRangeValues <dbl>,
#> #   q75_excludeRangeValues <dbl>, n_excludeRangeValues <int>,
#> #   median_ignoreRanges <dbl>, q25_ignoreRanges <dbl>, q75_ignoreRanges <dbl>
```

The analysis confirms that solo AMR markers have low PPV (<0.5) for Ciprofloxacin. In contrast, 27 combinations of between 2 and 6 markers have PPV >= 0.5. The distribution of the number of markers in these combinations is as follows: No. of markers No.of combinations 2 1 3 4 4 9 5 6 6 7

Importantly, out of the 27 combinations, 25 include mutations gyrA_S83L and parC_S80I.

We can use all this information to refine our concordance analysis.

Note: the `amr_ppv()` function applies by default a `min_set_size` threshold of 2, meaning that only solo markers or marker combinations with at least 2 occurrences in the dataset are included in the plots. Nevertheless solo markers or marker combinations that occur only once in the dataset are included in the stats table. In this example, there are 10 marker combinations represented by only 1 isolate in the dataset.

#### 8. Analyse concordance refining the definition of the genotypic prediction variable

We can refine the concordance analysis by setting requirements for the presence of gyrA_S83L and parC_S80I, or for a minimum number of markers. We will try this only for outcome R, as we have seen before that outcome NWT produces very similar concordance stats.

```
# Filter the genotypic prediction by the presence of specific mutations
concordance_cip_markers <- concordance(eco_cip_matrix,
  truth = "R",
  markers = c("gyrA_S83L", "parC_S80I")
)
```

```r
concordance_cip_markers
#> AMR Genotype-Phenotype Concordance
#> Prediction rule: any
#>
#> --- Outcome: R ---
#> Samples: 2075 | Markers: 2
#> Markers used: gyrA_S83L, parC_S80I
#>
#> Confusion Matrix:
#>           Truth
#> Prediction    1    0
#>          1  334  160
#>          0   13 1568
#>
#> Metrics:
#>   Sensitivity : 0.9625
#>   Specificity : 0.9074
#>   PPV         : 0.6761
#>   NPV         : 0.9918
#>   Accuracy    : 0.9166
#>   Kappa       : 0.7440
#>   F-measure   : 0.7943
#>   VME         : 0.0375
#>   ME          : 0.0926

# Filter the  genotypic prediction by the presence of a minimum number of markers
concordance_cip_min <- concordance(eco_cip_matrix,
  truth = "R",
  prediction_rule = 2
)
concordance_cip_min
#> AMR Genotype-Phenotype Concordance
#> Prediction rule: 2
#>
#> --- Outcome: R ---
#> Samples: 2075 | Markers: 31
#> Markers used: gyrA_D87N, gyrA_S83L, parC_S80I, parE_S458A, marR_S3N, qnrS1, parE_I529L,
        parC_E84V, qnrB19, parE_L416F, parC_S80R, aac(6')-Ib-cr5, parE_S458T, parE_D475E,
        parC_S57T, parE_I355T, gyrA_S83A, parC_E84G, qnrB2, marR_R77C, qnrB6, gyrA_D87Y,
        parE_L445H, gyrA_D87G, qnrB4, soxS_A12S, parE_I464F, parE_E460D, qnrB,
        soxR_G121D, parC_A56T
#>
#> Confusion Matrix:
#>           Truth
#> Prediction    1    0
#>          1  333  149
#>          0   14 1579
#>
#> Metrics:
#>   Sensitivity : 0.9597
#>   Specificity : 0.9138
#>   PPV         : 0.6909
#>   NPV         : 0.9912
#>   Accuracy    : 0.9214
#>   Kappa       : 0.7559
#>   F-measure   : 0.8034
```

```
#>   VME         : 0.0403
#>   ME          : 0.0862
```

The specificity, PPV, and ME values have substantially improved for both refinement strategies. The results are fairly similar because 25/27 combinations included the two mutations specified.

#### 9. Refine the concordance analysis by applying a PPV threshold

The PPV analysis in section 7 revealed that solo markers show low PPV for Ciprofloxacin (<= 0.029). Therefore, it is expected that refining the genotypic prediction by a solo PPV threshold would not substantially improve the concordance stats for this antibiotic. Instead, we can try predicting R for all samples with a marker or combination that had PPV >=0.5.

```
# Pass the PPV analysis output and a desired threshold to the `concordance()` function.
concordance_cip_ppv <- concordance(eco_cip_matrix,
  truth = "R",
  prediction_rule = "combo_ppv",
  ppv_results = eco_cip_ppv,
  ppv_threshold = 0.5
)
concordance_cip_ppv
#> AMR Genotype-Phenotype Concordance
#> Prediction rule: combo_ppv
#>
#> --- Outcome: R ---
#> Samples: 2075 | Markers: 24
#> Markers used: aac(6')-Ib-cr5, parE_I355T, qnrB6, gyrA_S83L, qnrS1, parC_S80I, qnrB4,
        gyrA_D87G, parE_S458A, gyrA_D87Y, gyrA_D87N, parC_A56T, parE_E460D, parE_I464F,
        parE_L445H, parC_E84G, parE_S458T, parE_L416F, parC_S57T, qnrB19, parE_I529L,
        parC_E84V, soxR_G121D, marR_S3N
#>
#> Confusion Matrix:
#>           Truth
#> Prediction    1    0
#>          1  329    3
#>          0   18 1725
#>
#> Metrics:
#>   Sensitivity : 0.9481
#>   Specificity : 0.9983
#>   PPV         : 0.9910
#>   NPV         : 0.9897
#>   Accuracy    : 0.9899
#>   Kappa       : 0.9630
#>   F-measure   : 0.9691
#>   VME         : 0.0519
#>   ME          : 0.0017
```

Applying this logic improves the PPV (from 0.691 to 0.991), specificity (from 0.914 to 0.998) and ME (from 0.0862 to 0.00174) without major detriment to sensitivity, NPV or VME.

#### 10. Refine the concordance analysis with the logistic regression

#### model

The `amr_logistic()` function of the `AMRgen` package performs logistic regression to analyse the relationship between genetic markers and phenotype (R, and NWT) for a specified antibiotic. It uses the binary matrix `eco_cip_markers` to fit logistic regression models for R vs S/I and/or NWT vs WT (by comparison to ECOFF), for Ciprofloxacin and a set of associated markers.

By default, only those markers present in at least 10 genomes are included in the logistic regression models, as regression often fails when very rare markers are included. We will use the default threshold but note that it can be adjusted by the user.

The results of the logistic regression analysis can also be used to refine the definition of the genotypic prediction variable by passing the options `prediction_rule="logistic"` and `logreg_results`. In this example, we will apply this to the R output only.

```
# Model a binary Ciprofloxacin phenotype using genetic marker presence/absence data
logreg <- amr_logistic(binary_matrix = eco_cip_matrix)
#> ...Fitting logistic regression model to R using logistf
#>    Filtered data contains 2075 samples (347 => 1, 1728 => 0) and 16 variables.
#> ...Fitting logistic regression model to NWT using logistf
#>    Filtered data contains 2075 samples (356 => 1, 1719 => 0) and 16 variables.
#> Generating plots
#> Plotting 2 models
```

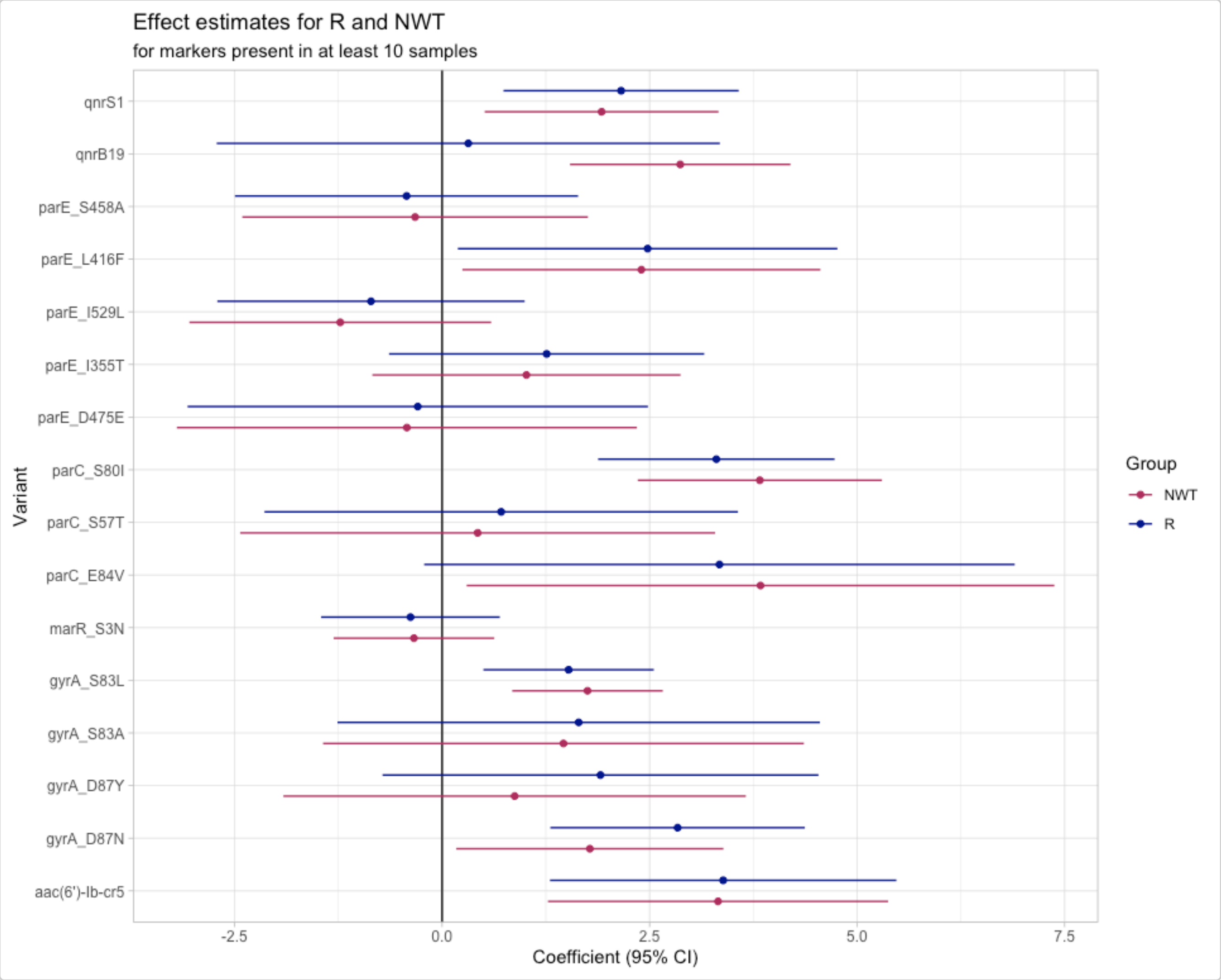

```r
# Apply the logistic regression results to the concordance analysis
concordance_cip_log <- concordance(eco_cip_matrix,
  truth = "R",
  prediction_rule = "logistic",
  logreg_results = logreg
)
concordance_cip_log
#> AMR Genotype-Phenotype Concordance
#> Prediction rule: logistic
#>
#> --- Outcome: R ---
#> Samples: 2075 | Markers: 31
#> Markers used: gyrA_D87N, gyrA_S83L, parC_S80I, parE_S458A, marR_S3N, qnrS1, parE_I529L,
        parC_E84V, qnrB19, parE_L416F, parC_S80R, aac(6')-Ib-cr5, parE_S458T, parE_D475E,
        parC_S57T, parE_I355T, gyrA_S83A, parC_E84G, qnrB2, marR_R77C, qnrB6, gyrA_D87Y,
        parE_L445H, gyrA_D87G, qnrB4, soxS_A12S, parE_I464F, parE_E460D, qnrB,
        soxR_G121D, parC_A56T
#>
#> Confusion Matrix:
#>           Truth
#> Prediction    1    0
#>          1  329    8
#>          0   18 1720
#>
#> Metrics:
#>   Sensitivity : 0.9481
#>   Specificity : 0.9954
#>   PPV         : 0.9763
#>   NPV         : 0.9896
#>   Accuracy    : 0.9875
#>   Kappa       : 0.9545
#>   F-measure   : 0.9620
#>   VME         : 0.0519
#>   ME          : 0.0046
```

The plot showing the logistic regression coefficient and 95% confidence interval shows that QRDR mutations gyrA_D87N, gyrA_S83L, parC_S80I, and parE_L416F, and genes aac(6')-Ib-cr5 and qnrS1 have an effect on the R phenotype (blue). The coefficients are larger than zero and the lower confidence value doesn't cross the zero line on the x-axis.

The `concordance()` function returns our data object with the predictions added in a new column, "R_pred". We can use the `assay_by_var()` function to colour our input MIC distribution by the genotypic prediction.

```r
assay_by_var(concordance_cip_log$data, colour_by = "R_pred")
```

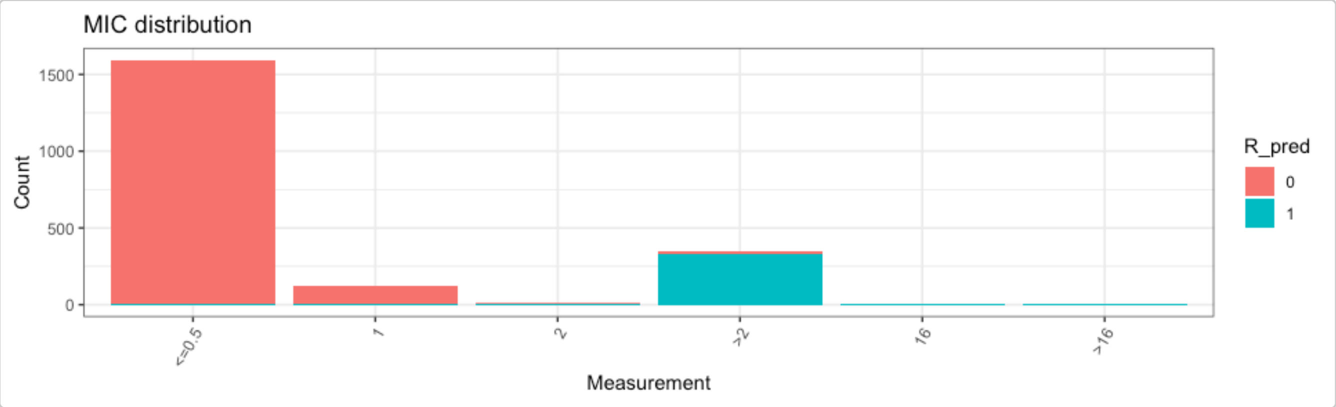


Predictions based on the logistic regression model have similar concordance statistics to those based on combinations with PPV>=0.5.

```
concordance_cip_log$metrics %>%
  left_join(concordance_cip_ppv$metrics, by = c("outcome", "metric"), suffix =
        c(".logistic", ".ppv"))
#> # A tibble: 9 x 4
#>   outcome metric   estimate.logistic estimate.ppv
#>   <chr>   <chr>                <dbl>        <dbl>
#> 1 R       sens               0.948        0.948
#> 2 R       spec               0.995        0.998
#> 3 R       ppv                0.976        0.991
#> 4 R       npv                0.990        0.990
#> 5 R       accuracy           0.987        0.990
#> 6 R       kap                0.954        0.963
#> 7 R       f_meas             0.962        0.969
#> 8 R       VME                0.0519       0.0519
#> 9 R       ME                 0.00463      0.00174
```

We can check the predictions from both methods, vs the observed phenotype, to see how many samples yield different predictions with the two methods.

```
concordance_cip_log$data %>%
  select(id, R_pred) %>%
  left_join(concordance_cip_ppv$data, by = "id", suffix = c(".logistic", ".ppv")) %>%
  count(R_pred.logistic, R_pred.ppv, R)
#> # A tibble: 7 x 4
#>   R_pred.logistic R_pred.ppv     R     n
#>             <int>      <int> <dbl> <int>
#> 1               0          0     0  1720
#> 2               0          0     1    15
#> 3               0          1     1     3
#> 4               1          0     0     5
#> 5               1          0     1     3
#> 6               1          1     0     3
#> 7               1          1     1   326
```

We can also extract and inspect the samples that the logistic model predicted wrongly, to explore their genotypes.

```
# samples with predictions different from the observed phenotype
concordance_cip_log$data %>%
  filter(R_pred != R)
```

```
#> # A tibble: 26 × 37
#>    id     R_pred     R   NWT pheno   mic gyrA_D87N gyrA_S83L parC_S80I parE_S458A
#>    <chr>   <int> <dbl> <dbl> <sir> <mic>     <dbl>     <dbl>     <dbl>      <dbl>
#>  1 SAMN…       1     0     0 S     <=0.5         1         1         1          1
#>  2 SAMN…       0     1     1 R      >2.0         0         0         0          0
#>  3 SAMN…       0     1     1 R      >2.0         0         0         0          0
#>  4 SAMN…       0     1     1 R      >2.0         0         0         0          0
#>  5 SAMN…       0     1     1 R      >2.0         0         0         0          0
#>  6 SAMN…       1     0     0 S       1.0         0         1         1          0
#>  7 SAMN…       1     0     0 S       1.0         0         1         1          0
#>  8 SAMN…       1     0     0 S       1.0         0         1         1          0
#>  9 SAMN…       0     1     1 R      >2.0         0         0         0          0
#> 10 SAMN…       0     1     1 R      >2.0         0         0         0          0
#> # ℹ 16 more rows
#> # ℹ 27 more variables: marR_S3N <dbl>, qnrS1 <dbl>, parE_I529L <dbl>,
#> #   parC_E84V <dbl>, qnrB19 <dbl>, parE_L416F <dbl>, parC_S80R <dbl>,
#> #   `aac(6')-Ib-cr5` <dbl>, parE_S458T <dbl>, parE_D475E <dbl>,
#> #   parC_S57T <dbl>, parE_I355T <dbl>, gyrA_S83A <dbl>, parC_E84G <dbl>,
#> #   qnrB2 <dbl>, marR_R77C <dbl>, qnrB6 <dbl>, gyrA_D87Y <dbl>,
#> #   parE_L445H <dbl>, gyrA_D87G <dbl>, qnrB4 <dbl>, soxS_A12S <dbl>, …

# plot the markers, S/I/R values, and MICs for samples
# that were predicted wrongly from the regression model
concordance_cip_log$data %>%
  filter(R_pred != R) %>%
  select(-R_pred) %>%
  amr_ppv()
#> Ordering markers by frequency
```

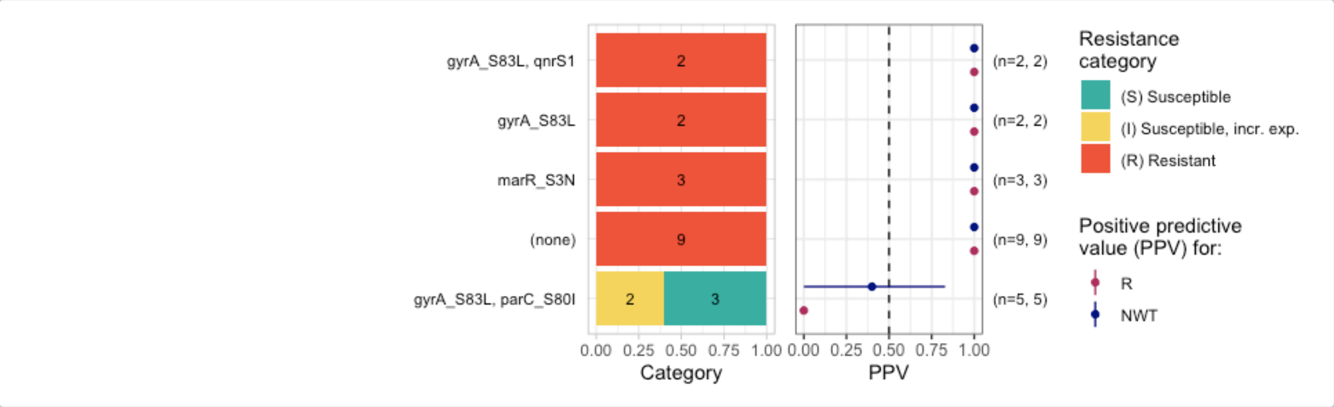


```
#> $plot
```

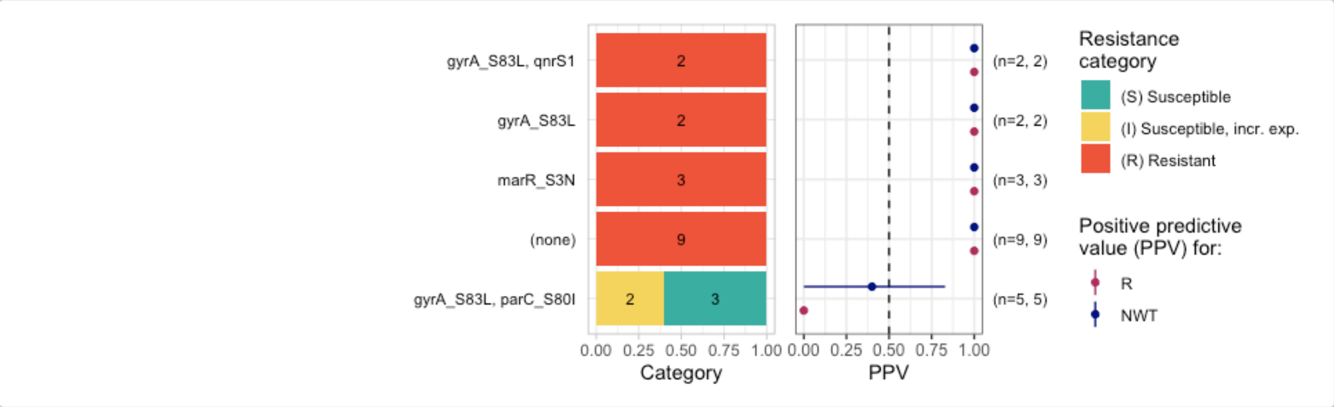

```
#>
#> $binary_matrix
#> # A tibble: 26 × 36
#>    id                R   NWT pheno   mic gyrA_D87N gyrA_S83L parC_S80I parE_S458A
#>    <chr>         <dbl> <dbl> <sir> <mic>     <dbl>     <dbl>     <dbl>      <dbl>
#>  1 SAMN26304359      0     0 S     <=0.5         1         1         1          1
#>  2 SAMN26304504      1     1 R      >2.0         0         0         0          0
#>  3 SAMN26304509      1     1 R      >2.0         0         0         0          0
#>  4 SAMN26304557      1     1 R      >2.0         0         0         0          0
#>  5 SAMN26304572      1     1 R      >2.0         0         0         0          0
#>  6 SAMN26304667      0     0 S       1.0         0         1         1          0
#>  7 SAMN26304713      0     0 S       1.0         0         1         1          0
#>  8 SAMN26304714      0     0 S       1.0         0         1         1          0
#>  9 SAMN26304849      1     1 R      >2.0         0         0         0          0
#> 10 SAMN26305235      1     1 R      >2.0         0         0         0          0
#> # ℹ 16 more rows
#> # ℹ 27 more variables: marR_S3N <dbl>, qnrS1 <dbl>, parE_I529L <dbl>,
#> #   parC_E84V <dbl>, qnrB19 <dbl>, parE_L416F <dbl>, parC_S80R <dbl>,
#> #   `aac(6')-Ib-cr5` <dbl>, parE_S458T <dbl>, parE_D475E <dbl>,
#> #   parC_S57T <dbl>, parE_I355T <dbl>, gyrA_S83A <dbl>, parC_E84G <dbl>,
#> #   qnrB2 <dbl>, marR_R77C <dbl>, qnrB6 <dbl>, gyrA_D87Y <dbl>,
#> #   parE_L445H <dbl>, gyrA_D87G <dbl>, qnrB4 <dbl>, soxS_A12S <dbl>, …
#>
#> $summary
#> # A tibble: 10 × 14
#>    marker_list          marker_count     n combination_id   R.n R.ppv R.ci_lower
#>    <chr>                       <dbl> <int> <fct>          <dbl> <dbl>      <dbl>
#>  1 ""                              0     9 0_0_0_0_0_0_0…     9     1          1
#>  2 "aac(6')-Ib-cr5, pa…            3     1 0_0_0_0_0_0_0…     1     1          1
#>  3 "marR_S3N"                      1     3 0_0_0_0_1_0_0…     3     1          1
#>  4 "gyrA_S83L"                     1     2 0_1_0_0_0_0_0…     2     1          1
#>  5 "gyrA_S83L, parE_I5…            3     1 0_1_0_0_0_0_1…     1     1          1
#>  6 "gyrA_S83L, qnrS1"              2     2 0_1_0_0_0_1_0…     2     1          1
#>  7 "gyrA_S83L, parC_S8…            2     5 0_1_1_0_0_0_0…     0     0          0
#>  8 "gyrA_S83L, parC_S8…            4     1 0_1_1_1_0_0_0…     0     0          0
#>  9 "gyrA_D87N, gyrA_S8…            4     1 1_1_1_0_0_0_0…     0     0          0
#> 10 "gyrA_D87N, gyrA_S8…            4     1 1_1_1_1_0_0_0…     0     0          0
#> # ℹ 7 more variables: R.ci_upper <dbl>, R.denom <int>, NWT.n <dbl>,
#> #   NWT.ppv <dbl>, NWT.ci_lower <dbl>, NWT.ci_upper <dbl>, NWT.denom <int>
```

The logistic regression model wrongly classified as susceptible 16 R samples; these include 9 in which no markers were identified, 3 with marR_S3N only, 2 with gyrA_S83L only, and 2 with gyrA_S83L plus qnrS1. Five S/I samples with gyrA_S83L plus parC_S80I were predicted as R.

### Example using large-scale regional/national surveillance data

**Leonor Sánchez-Busó**

**2026-03-09**

### Analysing *Neisseria gonorrhoeae* geno-pheno data

#### Introduction

This vignette demonstrates three usage examples of `AMRgen` functions to investigate associations between genotype and phenotype data in *Neisseria gonorrhoeae*. Specifically, we illustrate how to:

- Importing AMR genotype data from AMRFinderPlus.
- Import and format antimicrobial susceptibility testing (AST) phenotypic data in the form of minimum inhibitory concentrations (MICs) from a table.
- Explore the distribution of phenotypes across observed combinations of genetic AMR markers.
- Investigate the statistical association of individual (solo) markers and marker combinations with phenotype data.
- Work with MIC data from antibiotics with available EUCAST clinical breakpoints and/or epidemiological cut-offs (ECOFF).
- Evaluate the concordance of phenotypes with observed AMR markers and predict resistance based on this concordance.

Load the necessary libraries before running this vignette:

```
library(AMRgen)
library(dplyr)
library(tidyr)
library(ggplot2)
```

#### Use case 1: Investigation of genotype-phenotype AMR data from Euro-GASP genomic surveys

##### Data preparation

For this example, we have collated whole-genome sequencing data from three European Gonococcal Antimicrobial Surveillance Programme (Euro-GASP) genomic surveys. Raw FASTQ data was downloaded from the European Nucleotide Archive (ENA).

- Euro-GASP 2013, by Harris *et al.* (2018).
  - ENA PRJEB9227, n=1,054 genomes.

- Euro-GASP 2018, by Sánchez-Busó *et al.* (2022).
  - ENA PRJEB34068, n=2,375 genomes.
- Euro-GASP 2020, by Golparian *et al.* (2024).
  - ENA PRJEB58139, n=1,932 genomes.

Assemblies were generated with `SPAdes v3.15.5` (`--careful` mode) and assessed with `Quast v5.1`. Further details on the quality control and assembly pipeline used for this data are described in Sánchez-Serrano *et al.* (2026).

Phenotypic MIC data were obtained from the respective publications and collated into a single table (one row per isolate, one column per antibiotic). This pre-loaded object `eurogasp_pheno_raw` serves as the **phenotype input** for `AMRgen`:

```
head(eurogasp_pheno_raw)
#> # A tibble: 6 × 5
#>   id         Azithromycin Ciprofloxacin Cefixime Ceftriaxone
#>   <chr>             <dbl>         <dbl> <chr>          <dbl>
#> 1 ERR1549755        0.19          8     0.064          0.032
#> 2 ERR1549756        0.25          0.008 0.047          0.023
#> 3 ERR1549757        0.125        32     0.016          0.006
#> 4 ERR1549758        0.38          0.003 0.016          0.003
#> 5 ERR1549759        0.5           0.002 0.016          0.004
#> 6 ERR1471130       NA            NA     0.016          0.012
```

The dataset (total n = 5,361) includes MIC data for:

- Azithromycin (n=5,055 isolates)
- Ciprofloxacin (n=5,360 isolates)
- Cefixime (n=5,361 isolates)
- Ceftriaxone (n=5,361 isolates)

#### Identification of genetic AMR determinants using `AMRFinderPlus`

`AMRFinderPlus v4.0.23` (database version 2025-03-25.1) was run on all assemblies using the following command:

```
amrfinder --threads 4 --print_node -O Neisseria_gonorrhoeae --name ${IN%.fasta} -n $IN -o
        ${IN%.fasta}_amrfp.tsv \
--database /path/to/AMRFinderPlus/databases/4.0/2025-03-25.1
```

> **Note:** The `--print_node` flag must be enabled to include gene hierarchy node names in the output.

All per-sample output files were concatenated into a single table:

```
awk 'FNR>1 || NR==1' *_amrfp.tsv > eurogasps_amrfp.tsv
```

This concatenated table is pre-loaded as `eurogasp_geno_raw`, the **genotype input** for `AMRgen`.

#### Importing genotype and phenotype data into `AMRgen`

Use `import_amrfp()` to parse the AMRFinderPlus output:

```r
eurogasp_geno <- import_amrfp(
  input_table = eurogasp_geno_raw,
  sample_col = "Name"
)
```

For the phenotype file, reshape from wide to long format using `pivot_longer()`. MIC values across antibiotics may be stored as different types (numeric vs. character) due to inequality prefixes such as `<0.016` or `>32`. We coerce all antibiotic columns to character before pivoting to avoid type conflicts while preserving this information for downstream processing by `format_pheno()`:

```r
eurogasp_pheno <- eurogasp_pheno_raw %>%
  mutate(across(c(Azithromycin, Ciprofloxacin, Cefixime, Ceftriaxone), as.character)) %>%
  pivot_longer(
    cols = c(Azithromycin, Ciprofloxacin, Cefixime, Ceftriaxone),
    names_to = "drug",
    values_to = "mic"
  )
```

Then format with `format_pheno()`. Setting `interpret_eucast = TRUE` and `interpret_ecoff = TRUE` adds categorical SIR interpretations (clinical breakpoints) and WT/NWT classifications (ECOFF) fron EUCAST:

```r
eurogasp_ast <- format_pheno(
  input = eurogasp_pheno,
  sample_col = "id",
  species = "Neisseria gonorrhoeae",
  ab_col = "drug",
  mic_col = "mic",
  interpret_eucast = TRUE,
  interpret_ecoff = TRUE
)
#> Adding new micro-organism column 'spp_pheno' (class 'mo') with constant value Neisseria
        gonorrhoeae
#> Parsing column spp_pheno as micro-organism (class 'mo')
#> Parsing column drug as antibiotic (class 'ab')
#> Parsing column mic as class 'mic'
#> Could not find disk_col disk in input table
#> Could not find pheno_col ecoff in input table
#> Could not find pheno_col pheno_eucast in input table
#> Could not find pheno_col pheno_clsi in input table
#> Could not find pheno_col pheno_provided in input table
#> Could not find method_col method in input table
#> Could not find platform_col platform in input table
#> Could not find guideline_col guideline in input table
#> Could not find source_col source in input table
#> Interpreting all data as species: Neisseria gonorrhoeae
```

Importantly, out of the 5,361 strains under study and with phenotype data, genetic determinants of AMR were found for 5,186 by AMRFinderPlus. Negative samples should be added to the genotype table so they are properly accounted for in downstream analyses:

```r
negative_eurogasp <- eurogasp_pheno_raw %>%
  anti_join(eurogasp_geno, by = "id") %>%
  pull(id)
```

```
eurogasp_geno <- eurogasp_geno %>% bind_rows(tibble(id = negative_eurogasp))
```

#### Exploring phenotype distributions and comparing with EUCAST reference data

Round MIC values to the nearest doubling dilution using `as.mic()` with `round_to_next_log2 = TRUE`:

```
eurogasp_double <- eurogasp_ast %>%
  mutate(across(all_of("mic"), ~ AMR::as.mic(.x, round_to_next_log2 = TRUE)))
```

For each antibiotic, we visualise the MIC distribution with `assay_by_var()` and compare it to the EUCAST reference distribution using `compare_mic_with_eucast()`.

##### Azithromycin

```
# Plot the distribution of MIC data in the study
assay_by_var(
  pheno_table = eurogasp_double,
  pheno_drug = "Azithromycin",
  measure = "mic",
  colour_by = "ecoff"
)
```

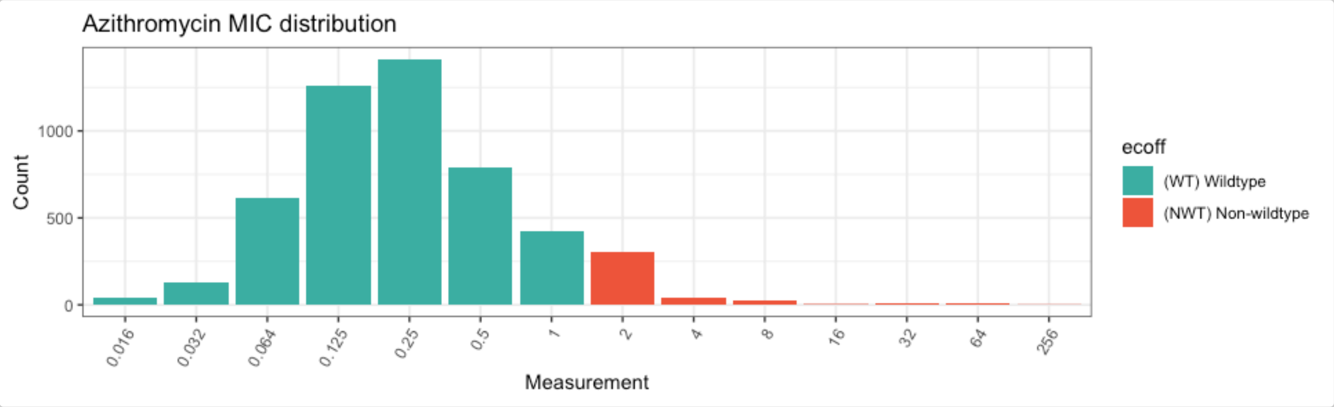


```
# Extract MIC data from the pheno table
azm_data <- eurogasp_double %>%
  filter(drug == "AZM") %>%
  pull(mic)
```

```
# Compare with a reference distribution from EUCAST
azm_comparison <- compare_mic_with_eucast(
  mics = azm_data,
  ab = "Azithromycin",
  mo = "Neisseria gonorrhoeae"
)
```

```r
# automated plot comparing to reference distribution
autoplot(azm_comparison)
```

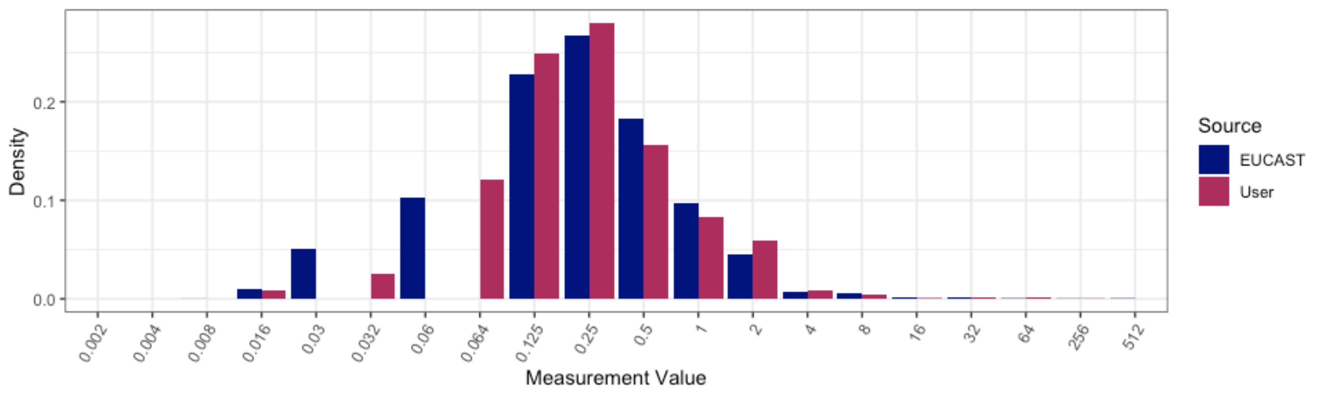


#### Ciprofloxacin

```r
# Plot the distribution of MIC data in the study
# The breakpoint for R is 0.06 but cannot be represented directly as it is not a doubling
        dilution (values in x axis).
assay_by_var(
  pheno_table = eurogasp_double,
  pheno_drug = "Ciprofloxacin",
  measure = "mic",
  colour_by = "pheno_eucast",
  species = "Neisseria gonorrhoeae"
)
#>   MIC breakpoints determined using AMR package: S <= 0.032 and R > 0.06
```

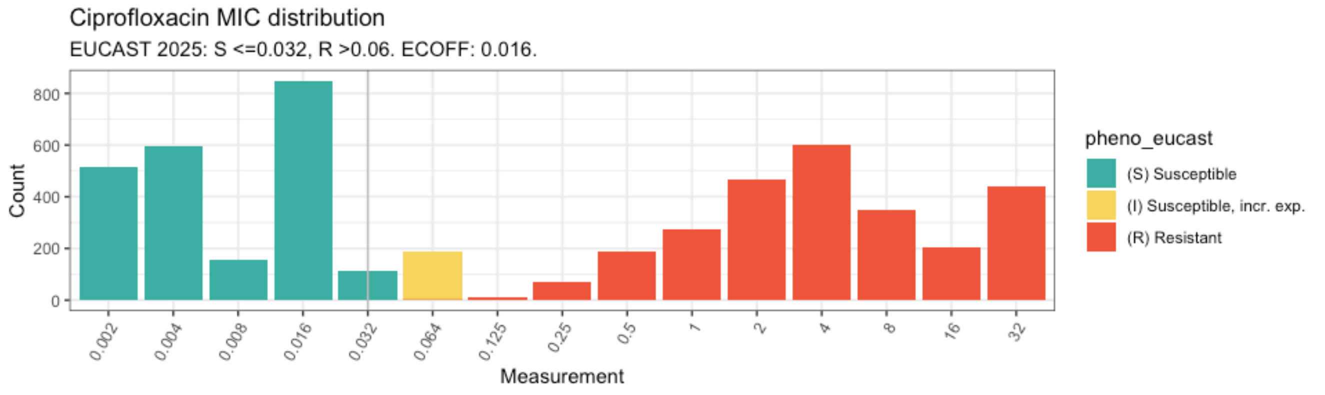


```r
# If the AST data without doublig dilutions is represented, then both breakpoints can be
        plotted.
assay_by_var(
  pheno_table = eurogasp_ast,
  pheno_drug = "Ciprofloxacin",
  measure = "mic",
  colour_by = "pheno_eucast",
  species = "Neisseria gonorrhoeae"
)
#>   MIC breakpoints determined using AMR package: S <= 0.032 and R > 0.06
```

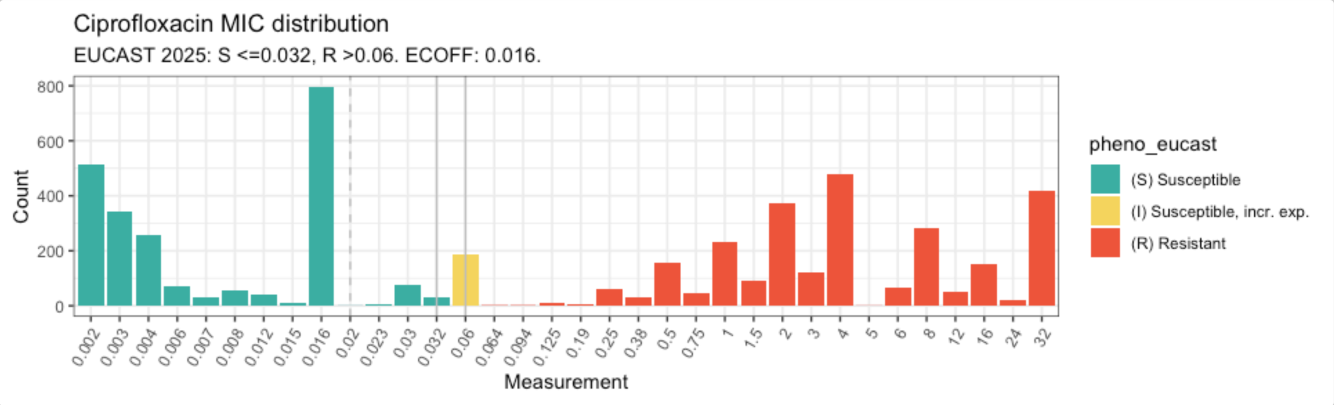


```r
# Extract MIC data from the pheno table
cip_data <- eurogasp_double %>%
  filter(drug == "CIP") %>%
  pull(mic)


# Compare with a reference distribution from EUCAST
cip_comparison <- compare_mic_with_eucast(
  mics = cip_data,
  ab = "Ciprofloxacin",
  mo = "N. gonorrhoeae"
)


# plot the data with the reference distribution
autoplot(cip_comparison)
```

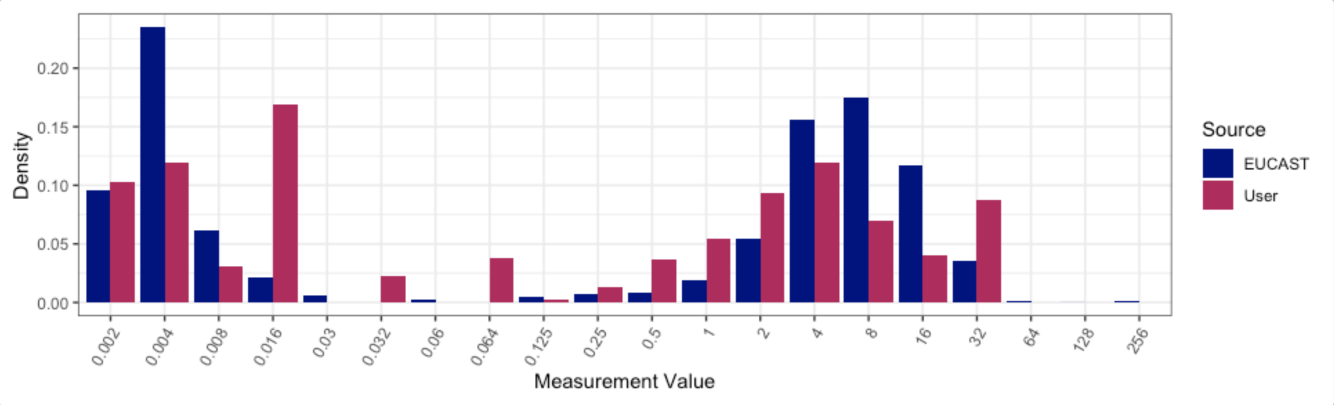


#### Ceftriaxone

```r
# Plot the distribution of MIC data in the study
assay_by_var(
  pheno_table = eurogasp_double,
  pheno_drug = "Ceftriaxone",
  measure = "mic",
  colour_by = "pheno_eucast",
  species = "Neisseria gonorrhoeae"
)
#>   MIC breakpoints determined using AMR package: S <= 0.125 and R > 0.125
```

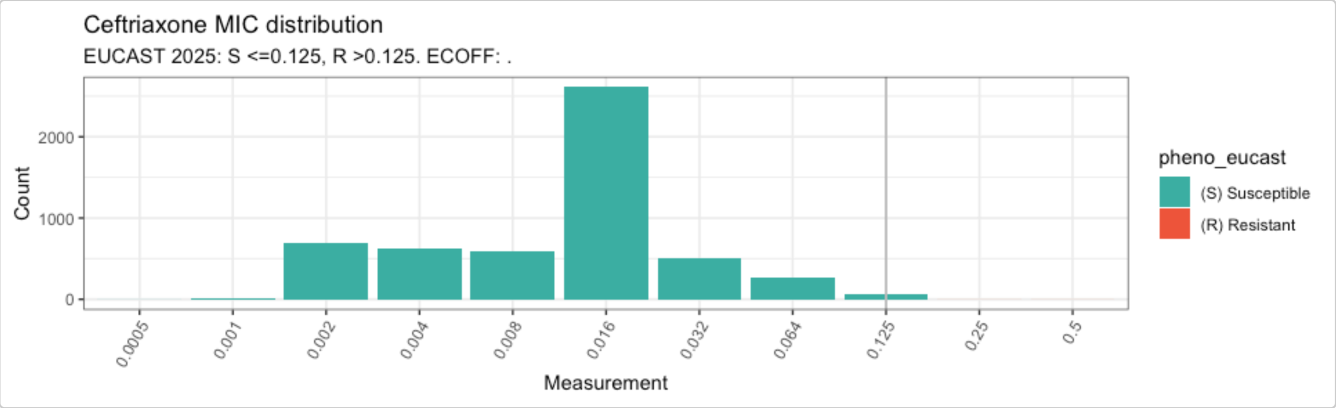


```r
# Extract MIC data from the pheno table
cro_data <- eurogasp_double %>%
  filter(drug == "CRO") %>%
  pull(mic)


# Compare with a reference distribution from EUCAST
cro_comparison <- compare_mic_with_eucast(
  mics = cro_data,
  ab = "Ceftriaxone",
  mo = "N. gonorrhoeae"
)


autoplot(cro_comparison)
```

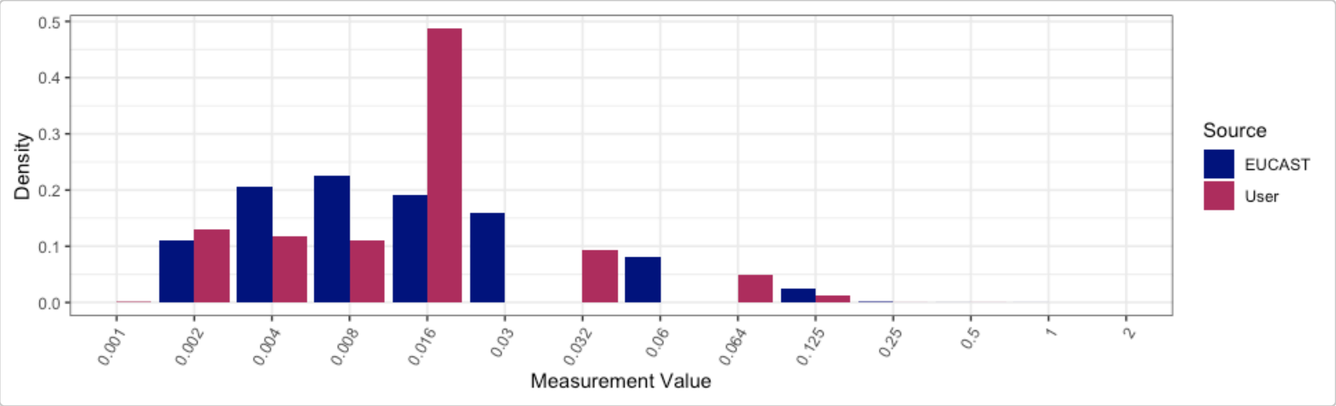


#### Cefixime

```r
# Plot the distribution of MIC data in the study
assay_by_var(
  pheno_table = eurogasp_double,
  pheno_drug = "Cefixime",
  measure = "mic",
  colour_by = "pheno_eucast",
  species = "Neisseria gonorrhoeae"
)
#>   MIC breakpoints determined using AMR package: S <= 0.125 and R > 0.125
```

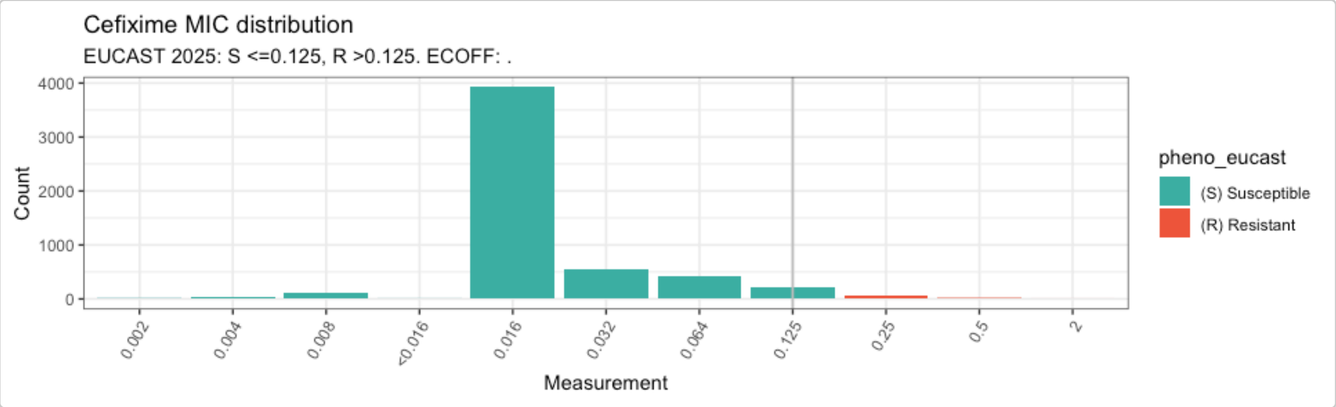


```
# Extract MIC data from the pheno table
cfm_data <- eurogasp_double %>%
  filter(drug == "CFM") %>%
  pull(mic)


# Compare with a reference distribution from EUCAST
cfm_comparison <- compare_mic_with_eucast(
  mics = cfm_data,
  ab = "Cefixime",
  mo = "N. gonorrhoeae"
)


autoplot(cfm_comparison)
```

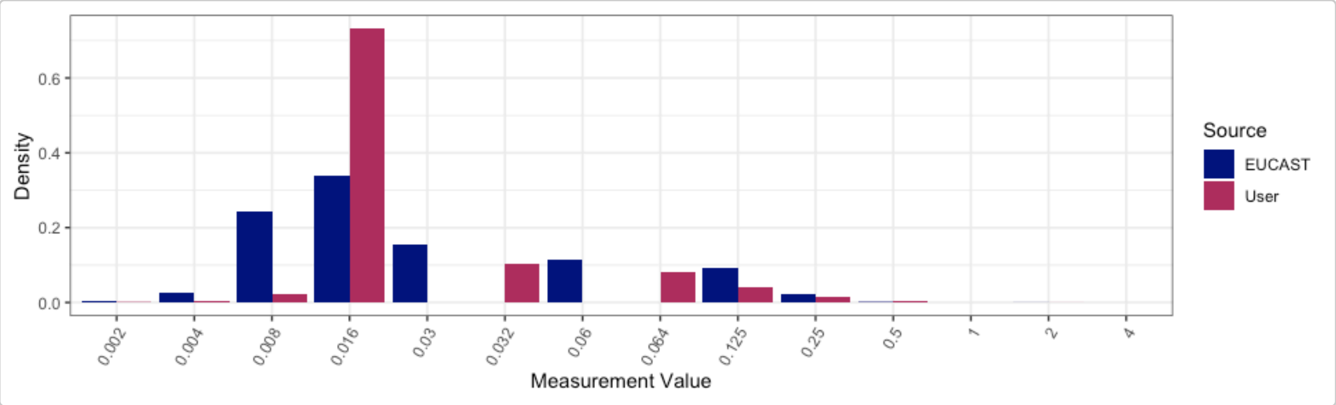


#### Analysing azithromycin genotype-phenotype data

Azithromycin was historically used in dual therapy for gonorrhoea but has been replaced in many countries by ceftriaxone monotherapy due to the expansion of resistant lineages. No EUCAST clinical breakpoint exists for azithromycin in *N. gonorrhoeae*; we therefore use the epidemiological cut-off (ECOFF > 1 mg/L).

Build the binary matrix combining genotype and phenotype:

```
azm_bin <- get_binary_matrix(
  geno_table = eurogasp_geno,
  pheno_table = eurogasp_ast,
  pheno_drug = "Azithromycin",
  geno_class = c("Macrolides", "Lincosamides"),
```

```r
  ecoff_col = "ecoff",
  sir_col = "pheno_eucast",
  keep_assay_values = TRUE,
  keep_assay_values_from = "mic"
)
#>  Defining NWT in binary matrix using ecoff column provided: ecoff
```

Explore the distribution of MIC values across observed marker combinations with `amr_upset()`:

```r
# Calculate upset plots of MIC distributions vs genotype marker combinations
# (specify species and drug in order to look up and plot the ecoff)
azi_upset <- amr_upset(
  binary_matrix = azm_bin,
  assay = "mic",
  min_set_size = 2,
  order = "value",
  pheno_drug = "azithromycin",
  species = "Neisseria gonorrhoeae"
)
#> Removing 306 rows with no phenotype call
#> Ordering markers by frequency
#> Error in executing command: Could not determine MIC breakpoints using AMR package,
        please provide your own breakpoints
#> Error in executing command: Could not determine MIC breakpoints using AMR package,
        please provide your own breakpoints
#> Scale for y is already present.
#> Adding another scale for y, which will replace the existing scale.
#> Scale for y is already present.
#> Adding another scale for y, which will replace the existing scale.
```

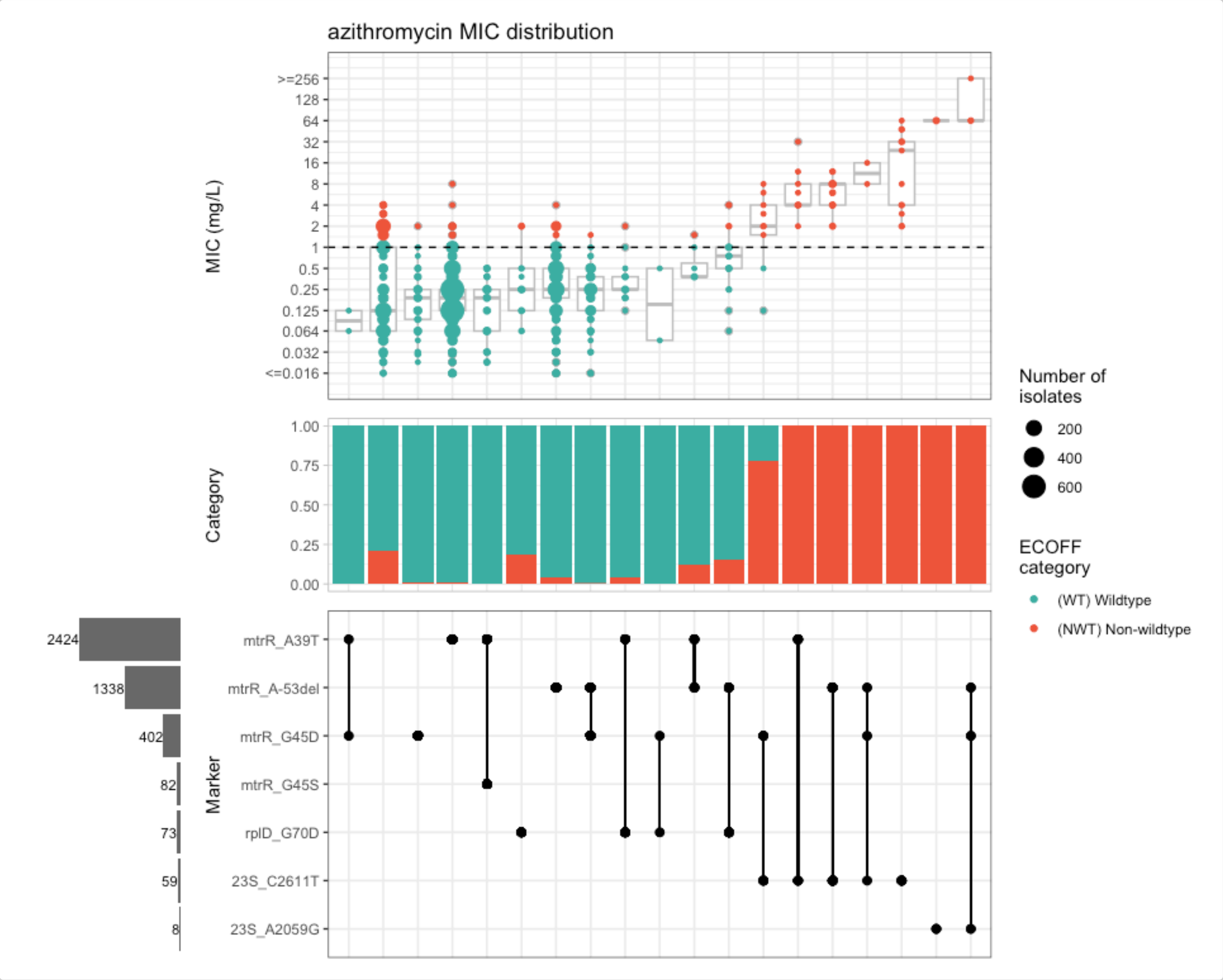


Evaluate the positive predictive value (PPV) of solo markers (i.e. markers occurring in the absence of any other known AMR determinant):

```
azm_solo_ppv <- solo_ppv(
  binary_matrix = azm_bin,
  pheno_drug = "Azithromycin",
  reverse_order = FALSE
)
```

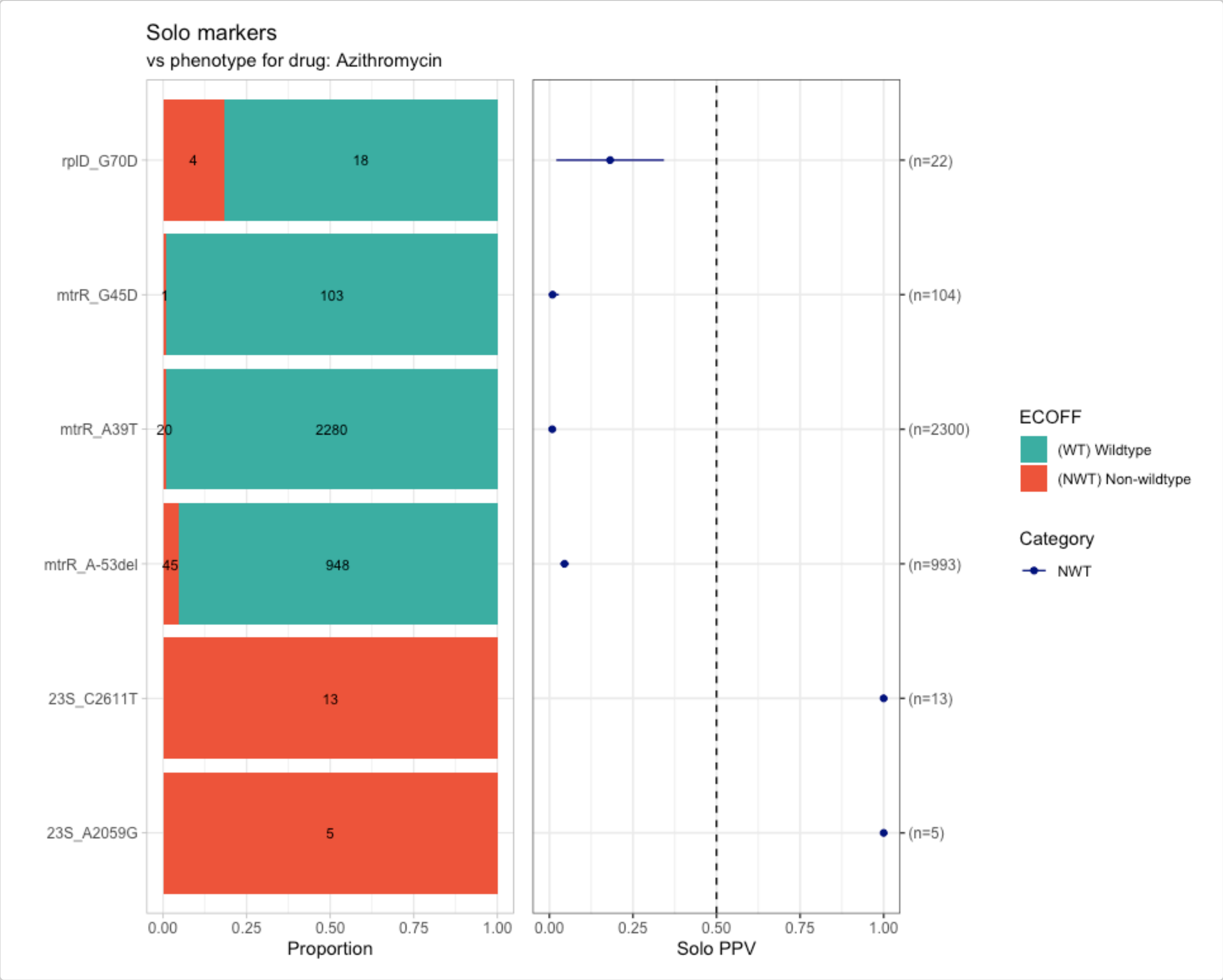


Only known mutations in the *23S* rDNA gene are associated with MIC increases sufficient to reach the NWT category (ECOFF > 1 mg/L). Mutations in the *mtrR* gene or its promoter, or the *rplD* G70D mutation alone, do not. PPV > 0.50 indicates that more than 50% of isolates carrying a given marker fall in the NWT category..

Now evaluate PPVs for marker combinations with `amr_ppv()`:

```
azm_ppv <- amr_ppv(
  binary_matrix = azm_bin,
  order = "value",
  min_set_size = 2,
  pheno_drug = "Azithromycin",
  upset_grid = TRUE,
  plot_assay = TRUE,
  assay = "mic"
)
#> Removing 306 rows with no phenotype call
#> Ordering markers by frequency
#> Scale for y is already present.
#> Adding another scale for y, which will replace the existing scale.
```

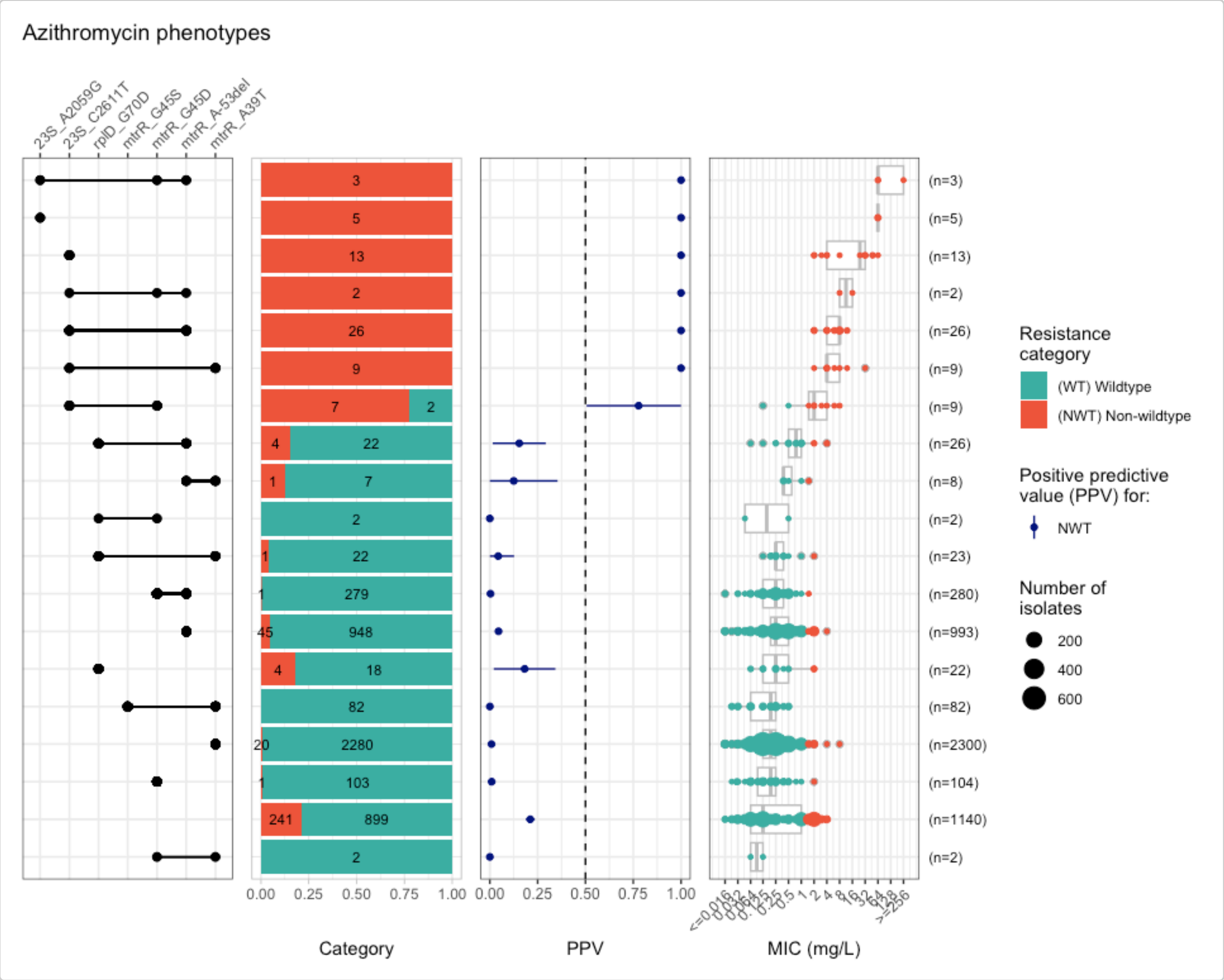


Combinations including *23S* rDNA mutations are the only ones with PPV > 0.5. No combination of *mtrR* with or without *rplD* mutations alone raises MIC above the NWT threshold.

Run a Firth's bias-reduced penalised-likelihood logistic regression with `amr_logistic()`, including markers with a minimum allele frequency (MAF) of ≥ 10 isolates:

```
azm_logist <- amr_logistic(
  binary_matrix = azm_bin,
  pheno_drug = "Azithromycin",
  ecoff_col = "ecoff",
  maf = 10,
  single_plot = TRUE
)
#> ...Fitting logistic regression model to R using logistf
#> ...Fitting logistic regression model to NWT using logistf
#>    Filtered data contains 5055 samples (386 => 1, 4669 => 0) and 7 variables.
#> Generating plots
#> Plotting NWT model only
```

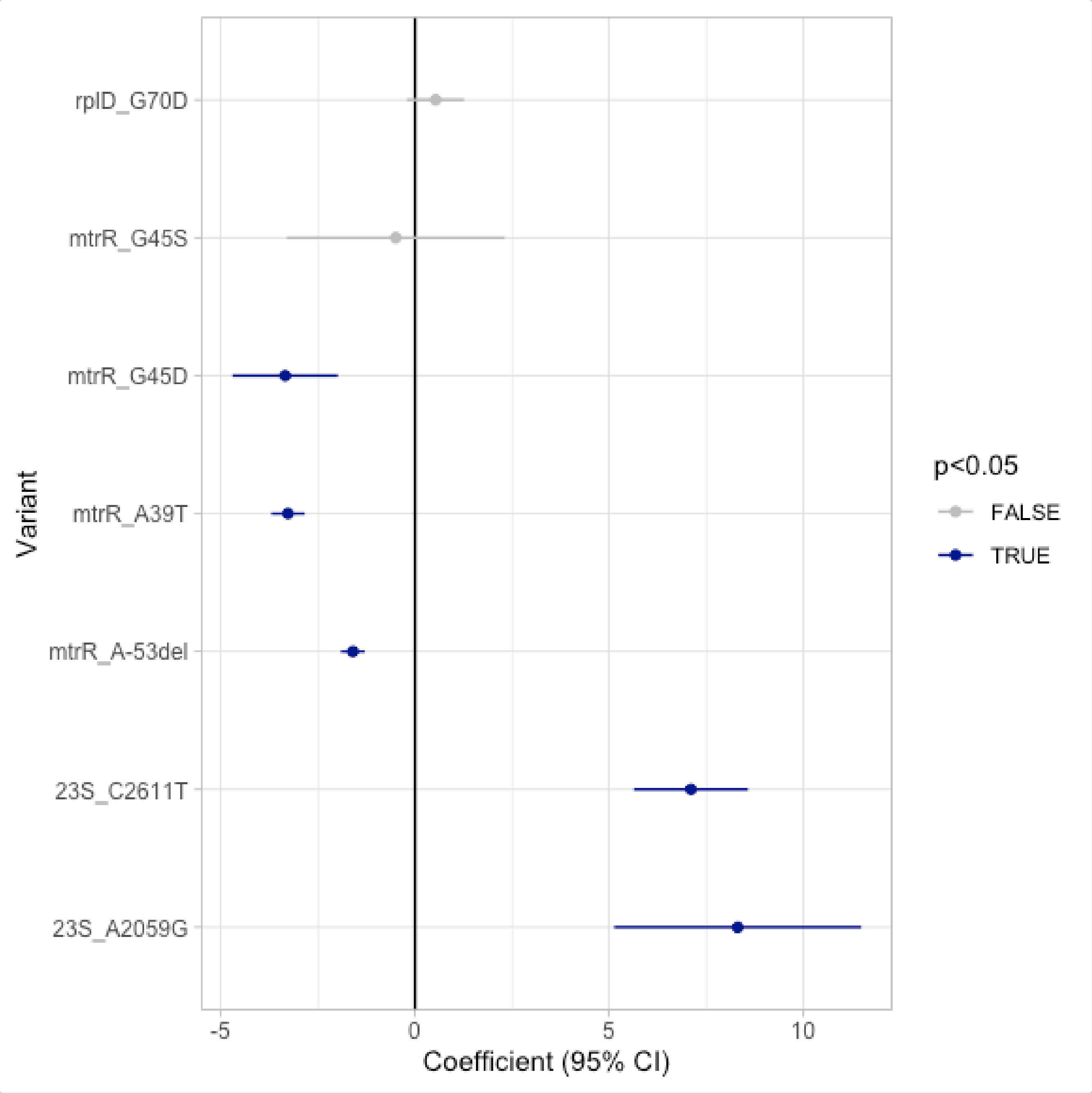


The regression confirms that *mtrR* G45D, A39T, and A-53del do not contribute to increased azithromycin MICs, while *23S* rDNA mutations do.

Calculate concordance and resistance prediction metrics using results from the logistic regression. The results of the `solo_ppv_analisis()` or `amr_ppv()` functions can be provided to the `ppv_results` parameter.

```
azm_concordance <- concordance(
  binary_matrix = azm_bin,
  ppv_results = azm_solo_ppv,
  prediction_rule = "logistic",
  logreg_results = azm_logist
)
azm_concordance
#> AMR Genotype-Phenotype Concordance
#> Prediction rule: logistic
#>
#> --- Outcome: NWT ---
#> Samples: 5055 | Markers: 10
```

```
#> Markers used: mtrR_A39T, mtrR_A-53del, mtrR_G45D, 23S_A2059G, mtrR_G45S, rplD_G70D,
       erm(B), 23S_C2611T, mtrR_G-131A, mtrC_C-120T
#>
#> Confusion Matrix:
#>           Truth
#> Prediction    1    0
#>          1   68    2
#>          0  318 4667
#>
#> Metrics:
#>   Sensitivity : 0.1762
#>   Specificity : 0.9996
#>   PPV         : 0.9714
#>   NPV         : 0.9362
#>   Accuracy    : 0.9367
#>   Kappa       : 0.2814
#>   F-measure   : 0.2982
#>   VME         : 0.8238
#>   ME          : 4e-04
```

The current marker set yields only 17.62% sensitivity (i.e. only 17.62% of resistant isolates are correctly identified), with a very major error (VME) of 82.34%. This indicates that the majority of NWT isolates are not explained by the markers currently available in AMRFinderPlus. Notably, mosaic variants in the *mtrCDE* efflux pump genes — not yet fully characterised and represented in AMR databases — are likely contributing to this discrepancy. Two MtrD mutations associated with decreased azithromycin susceptibility (Ma *et al.*, 2020) have not yet been incorporated into AMRFinderPlus.

Visualise the prediction on the MIC distribution:

```
eurogasp_azm_pred <- eurogasp_ast %>%
  left_join(azm_concordance$data[, c("id", "NWT_pred")], by = "id")

head(eurogasp_azm_pred)
#> # A tibble: 6 × 7
#>   id         drug                 mic ecoff pheno_eucast spp_pheno       NWT_pred
#>   <chr>      <ab>                <mic> <sir> <sir>        <mo>               <int>
#> 1 ERR1549755 AZM: Azithromycin  0.190 WT    NA           B_NESSR_GNRR:…        0
#> 2 ERR1549755 CIP: Ciprofloxacin 8.000 NWT   R            B_NESSR_GNRR:…        0
#> 3 ERR1549755 CFM: Cefixime      0.064 NA    S            B_NESSR_GNRR:…        0
#> 4 ERR1549755 CRO: Ceftriaxone   0.032 NA    S            B_NESSR_GNRR:…        0
#> 5 ERR1549756 AZM: Azithromycin  0.250 WT    NA           B_NESSR_GNRR:…        0
#> 6 ERR1549756 CIP: Ciprofloxacin 0.008 WT    S            B_NESSR_GNRR:…        0

assay_by_var(
  pheno_table = eurogasp_azm_pred,
  pheno_drug = "Azithromycin",
  measure = "mic",
  colour_by = "NWT_pred",
  species = "Neisseria gonorrhoeae",
  colours = c("#3CAEA3", "#ED553B"),
  colour_legend_label = "NWT prediction"
)
#> Error in executing command: Could not determine MIC breakpoints using AMR package,
       please provide your own breakpoints
```

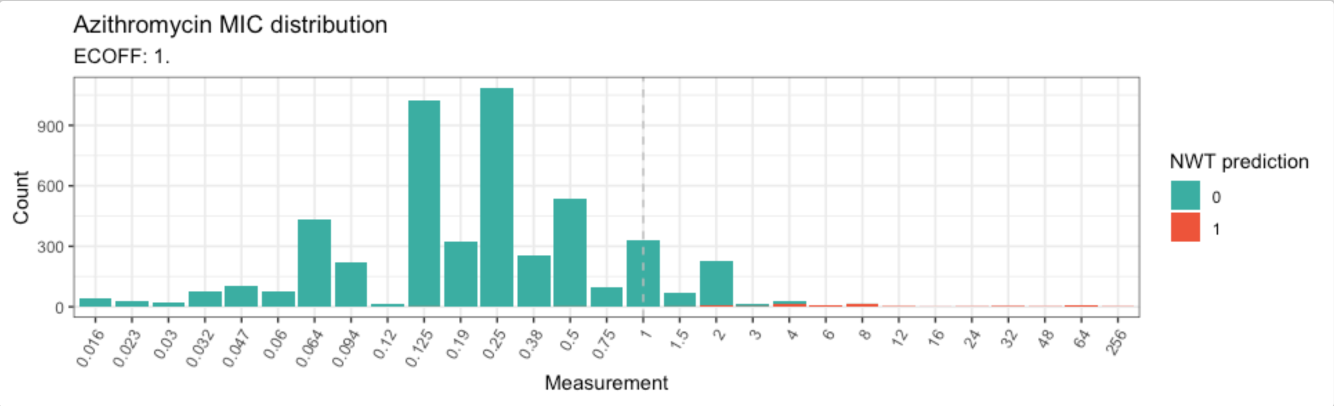


Many isolates with MICs above the ECOFF (1–4 mg/L) are predicted as WT, consistent with uncharacterised efflux pump variants.

#### Analysing ciprofloxacin genotype-phenotype data

Ciprofloxacin resistance is widespread in *N. gonorrhoeae* and is strongly associated with known genetic determinants — primarily mutations in the *gyrA* and *parC* genes. Build the binary matrix and generate upset plots:

```
# Get binary matrix
cip_bin <- get_binary_matrix(
  geno_table = eurogasp_geno,
  pheno_table = eurogasp_ast,
  pheno_drug = "Ciprofloxacin",
  geno_class = "Quinolones",
  sir_col = "pheno_eucast",
  keep_assay_values = TRUE,
  keep_assay_values_from = "mic"
)
#>  Defining NWT in binary matrix using ecoff column provided: ecoff

# Calculate upset plots of MIC distributions vs genotype marker combinations
cip_upset <- amr_upset(
  binary_matrix = cip_bin,
  min_set_size = 1,
  order = "value",
  assay = "mic",
  pheno_drug = "Ciprofloxacin",
  species = "Neisseria gonorrhoeae",
  plot_subtitle = "vs quinolone associated markers"
)
#> Removing 336 rows with no phenotype call
#> Ordering markers by frequency
#>   MIC breakpoints determined using AMR package: S <= 0.032 and R > 0.06
#>   MIC breakpoints determined using AMR package: S <= 0.032 and R > 0.06
#> Scale for y is already present.
#> Adding another scale for y, which will replace the existing scale.
#> Scale for y is already present.
#> Adding another scale for y, which will replace the existing scale.
```

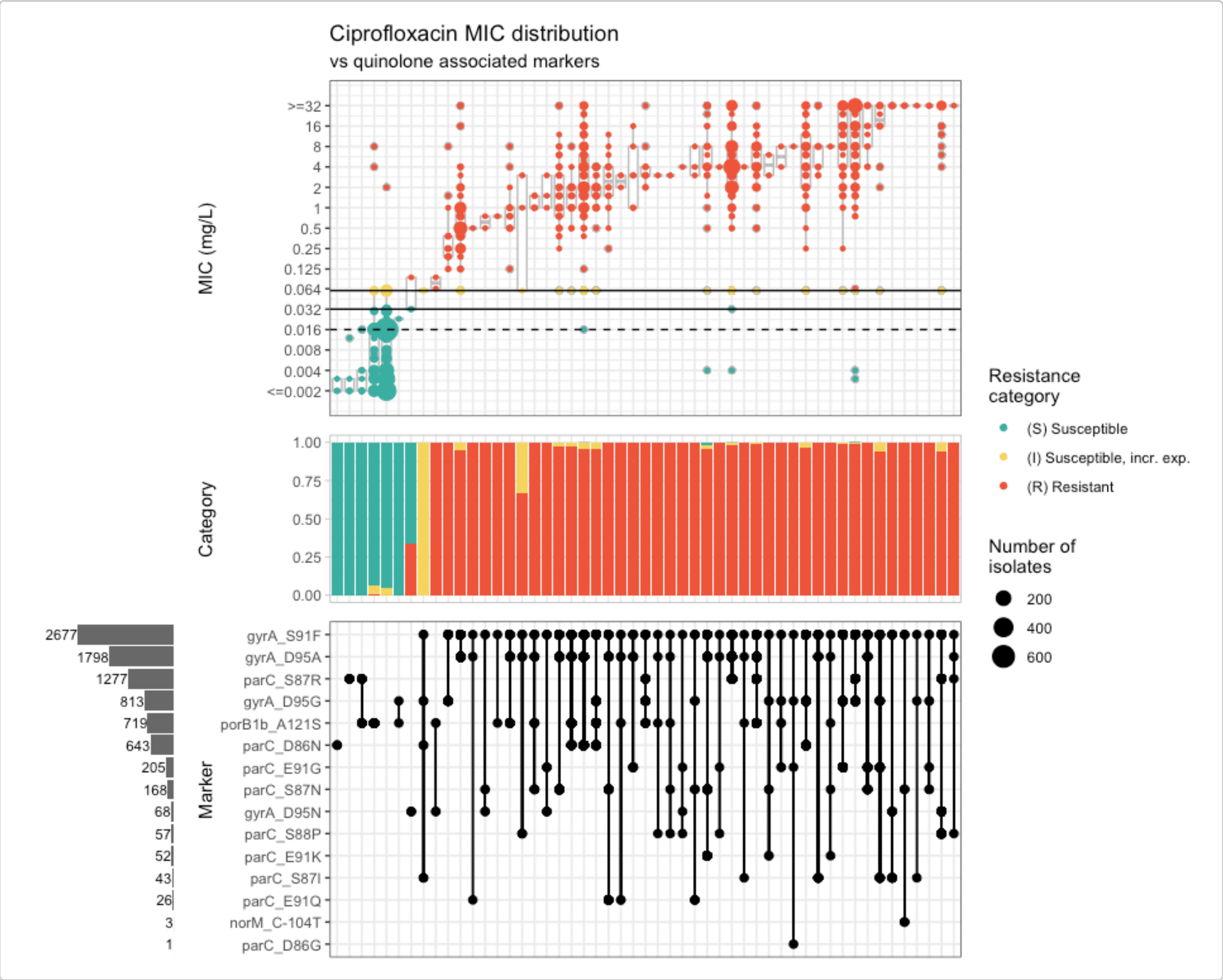


The upset plot shows that most isolates with an MIC above the EUCAST clinical breakpoint (R) carry the *gyrA* S91F mutation, the principal driver of ciprofloxacin resistance. Horizontal black lines in the top panel indicate clinical breakpoints; the dashed line represents the ECOFF.

Only four markers appear in isolation; evaluate their solo PPVs:

```
cip_solo_ppv <- solo_ppv(
  binary_matrix = cip_bin,
  pheno_drug = "Ciprofloxacin",
  reverse_order = FALSE
)
```

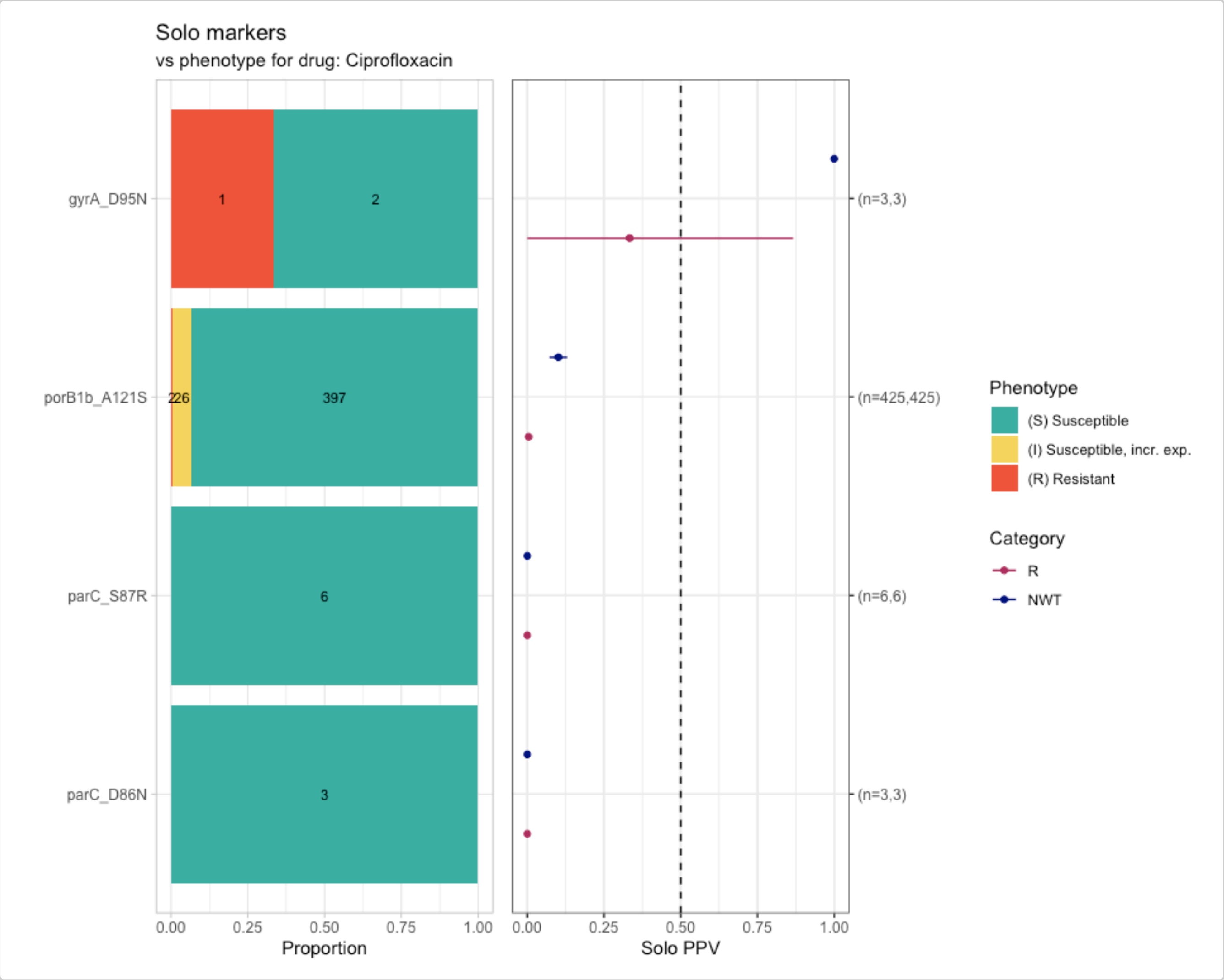


Evaluate PPVs for marker combinations:

```
cip_ppv <- amr_ppv(
  binary_matrix = cip_bin,
  order = "value",
  min_set_size = 2,
  pheno_drug = "Ciprofloxacin",
  upset_grid = TRUE,
  plot_assay = TRUE,
  assay = "mic"
)
#> Removing 336 rows with no phenotype call
#> Ordering markers by frequency
#> Scale for y is already present.
#> Adding another scale for y, which will replace the existing scale.
```

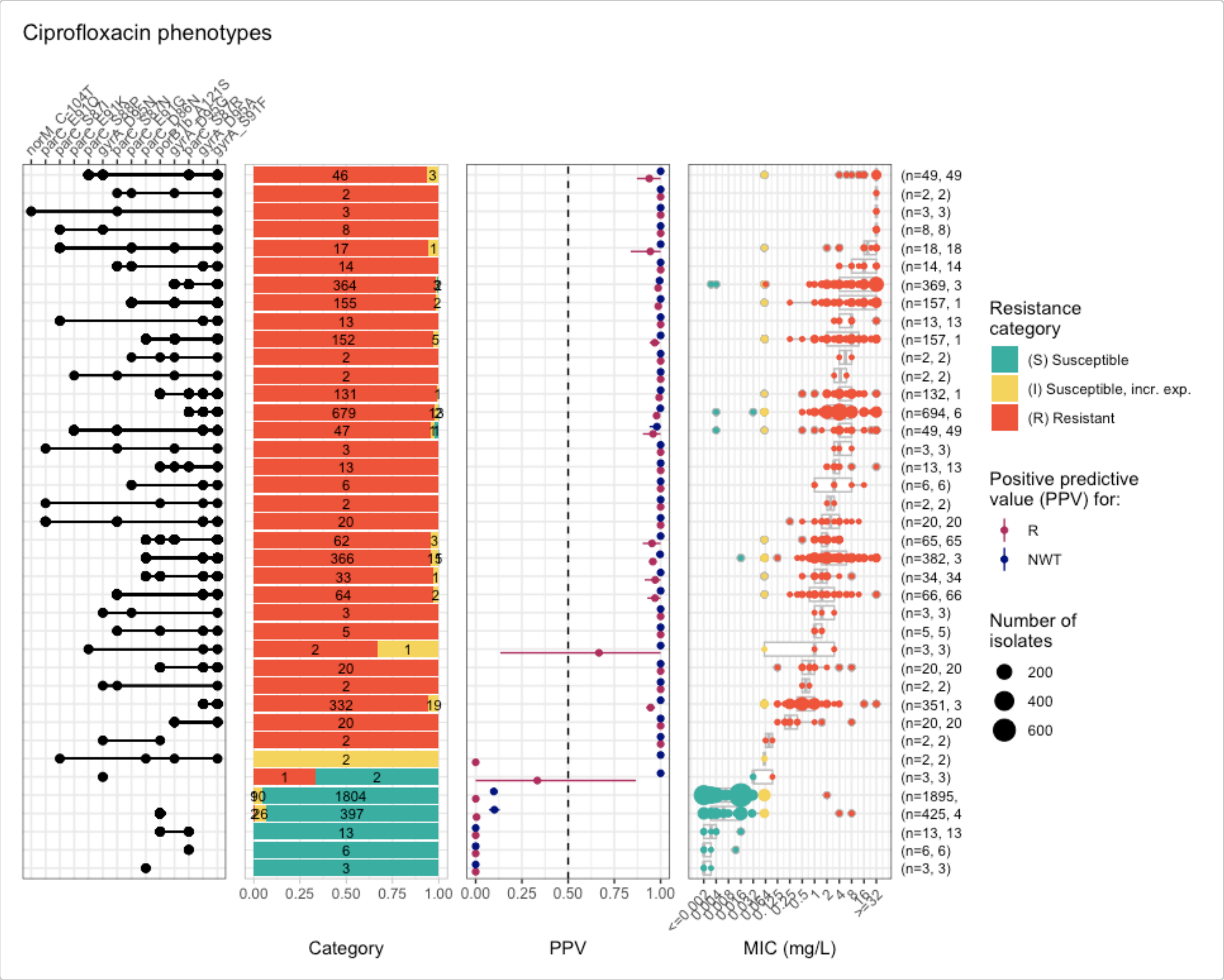


Run logistic regression to assess the individual contribution of each marker:

```
cip_logist <- amr_logistic(
  binary_matrix = cip_bin,
  pheno_drug = "Ciprofloxacin",
  sir_col = "pheno_eucast",
  ecoff_col = "ecoff",
  maf = 10,
  single_plot = TRUE
)
#> ...Fitting logistic regression model to R using logistf
#>    Filtered data contains 5025 samples (2605 => 1, 2420 => 0) and 13 variables.
#> ...Fitting logistic regression model to NWT using logistf
#>    Filtered data contains 5025 samples (2907 => 1, 2118 => 0) and 13 variables.
#> Generating plots
#> Plotting 2 models
```

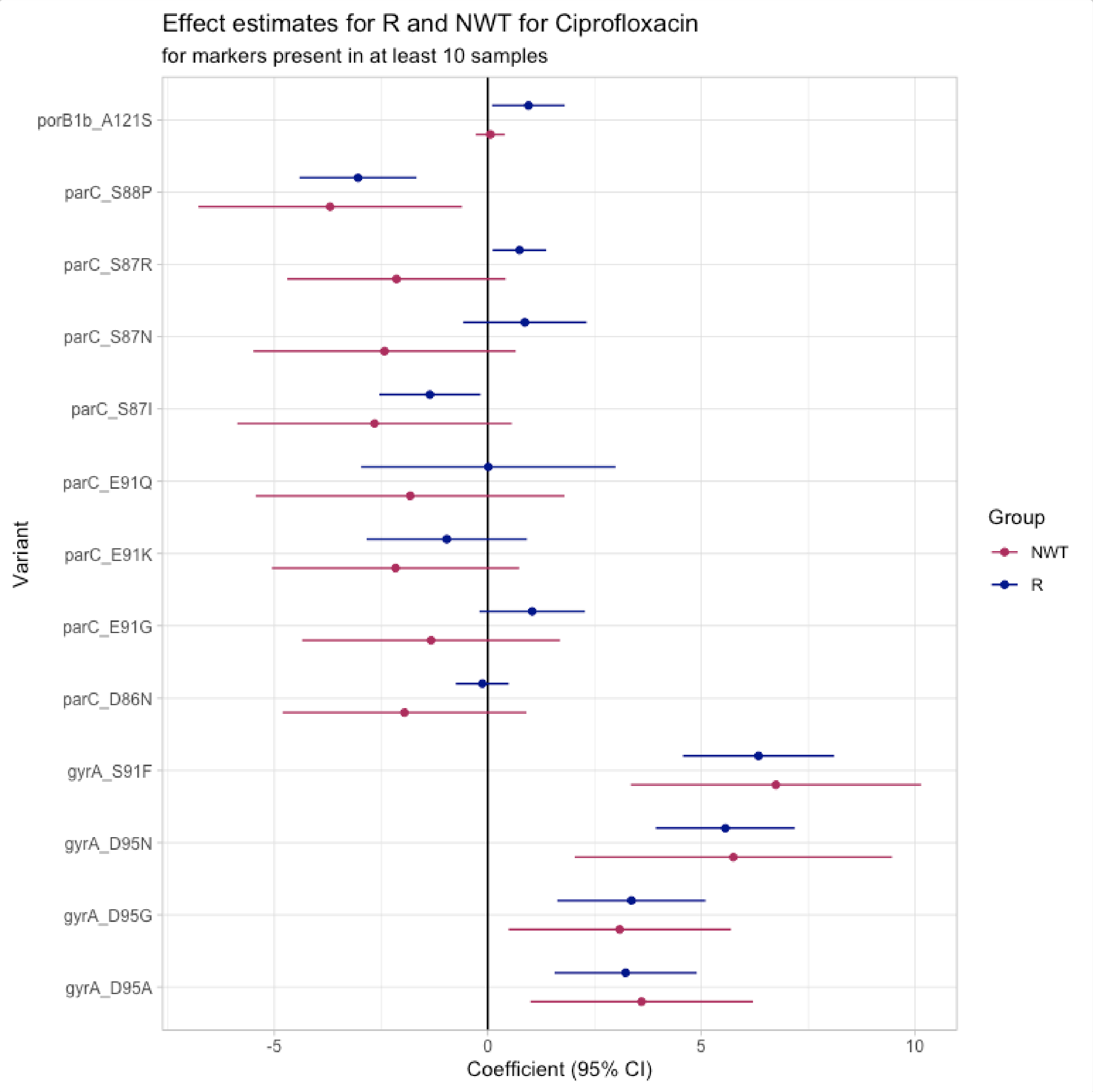


Calculate concordance metrics using the results from the logistic regression:

```
cip_concordance <- concordance(
  binary_matrix = cip_bin,
  ppv_results = cip_solo_ppv,
  prediction_rule = "logistic",
  logreg_results = cip_logist
)
cip_concordance
#> AMR Genotype-Phenotype Concordance
#> Prediction rule: logistic
#>
#> --- Outcome: R ---
#> Samples: 5025 | Markers: 15
#> Markers used: porB1b_A121S, gyrA_D95A, gyrA_S91F, parC_S87R, parC_D86N, gyrA_D95G,
         parC_E91G, parC_E91Q, parC_S87N, norM_C-104T, parC_E91K, gyrA_D95N, parC_S87I,
         parC_S88P, parC_D86G
#>
#> Confusion Matrix:
```

```
#>           Truth
#> Prediction    1    0
#>          1 2601   78
#>          0    4 2342
#>
#> Metrics:
#>   Sensitivity : 0.9985
#>   Specificity : 0.9678
#>   PPV         : 0.9709
#>   NPV         : 0.9983
#>   Accuracy    : 0.9837
#>   Kappa       : 0.9673
#>   F-measure   : 0.9845
#>   VME         : 0.0015
#>   ME          : 0.0322
#>
#> --- Outcome: NWT ---
#> Samples: 5025 | Markers: 15
#> Markers used: porB1b_A121S, gyrA_D95A, gyrA_S91F, parC_S87R, parC_D86N, gyrA_D95G,
         parC_E91G, parC_E91Q, parC_S87N, norM_C-104T, parC_E91K, gyrA_D95N, parC_S87I,
         parC_S88P, parC_D86G
#>
#> Confusion Matrix:
#>           Truth
#> Prediction    1    0
#>          1 2678    5
#>          0  229 2113
#>
#> Metrics:
#>   Sensitivity : 0.9212
#>   Specificity : 0.9976
#>   PPV         : 0.9981
#>   NPV         : 0.9022
#>   Accuracy    : 0.9534
#>   Kappa       : 0.9059
#>   F-measure   : 0.9581
#>   VME         : 0.0788
#>   ME          : 0.0024
```

- Using `R` as outcome, we get a 99.85% sensitivity, 96.55% specificity and a PPV of 97.09% using all markers, with a major error (ME, a susceptible isolate is reported as resistant) of 3.45% and a very major error (VME, a resistant isolate is reported as susceptible) of ~0%.
- Using `NWT` as outcome, we get a 92.34% sensitivity, 99.75% specificity and a PPV of 99.81%, with a ME of ~0% but a VME of 7.67%.

The results from the concordance analysis contain a prediction of R/NWT based on the genotype-phenotype comparison under `$data` in the generated object `cip_concordance`. Extract these prediction columns, named `R_pred` and/or `NWT_pred`, and incorporate them into the `eurogasp_ast` object, which contains the MIC distribution for this antibiotic:

```
eurogasp_cip_pred <- eurogasp_ast %>%
  left_join(cip_concordance$data[, c("id", "R_pred", "NWT_pred")],
    by = "id"
  )
```

```r
head(eurogasp_cip_pred)
#> # A tibble: 6 × 8
#>   id         drug              mic ecoff pheno_eucast spp_pheno    R_pred NWT_pred
#>   <chr>      <ab>            <mic> <sir> <sir>        <mo>          <int>    <int>
#> 1 ERR1549755 AZM: Azithro… 0.190 WT    NA           B_NESSR_GNR…      1        1
#> 2 ERR1549755 CIP: Ciprofl… 8.000 NWT   R            B_NESSR_GNR…      1        1
#> 3 ERR1549755 CFM: Cefixime 0.064 NA    S            B_NESSR_GNR…      1        1
#> 4 ERR1549755 CRO: Ceftria… 0.032 NA    S            B_NESSR_GNR…      1        1
#> 5 ERR1549756 AZM: Azithro… 0.250 WT    NA           B_NESSR_GNR…      0        0
#> 6 ERR1549756 CIP: Ciprofl… 0.008 WT    S            B_NESSR_GNR…      0        0
```

Visualise the distribution of ciprofloxacin MICs by the R/NWT predictions:

```r
assay_by_var(
  pheno_table = eurogasp_cip_pred,
  pheno_drug = "Ciprofloxacin",
  measure = "mic",
  colour_by = "R_pred",
  species = "Neisseria gonorrhoeae",
  colours = c("#3CAEA3", "#ED553B"),
  colour_legend_label = "R prediction"
)
#>   MIC breakpoints determined using AMR package: S <= 0.032 and R > 0.06

assay_by_var(
  pheno_table = eurogasp_cip_pred,
  pheno_drug = "Ciprofloxacin",
  measure = "mic",
  colour_by = "NWT_pred",
  species = "Neisseria gonorrhoeae",
  colours = c("#3CAEA3", "#ED553B"),
  colour_legend_label = "NWT prediction"
)
#>   MIC breakpoints determined using AMR package: S <= 0.032 and R > 0.06
```

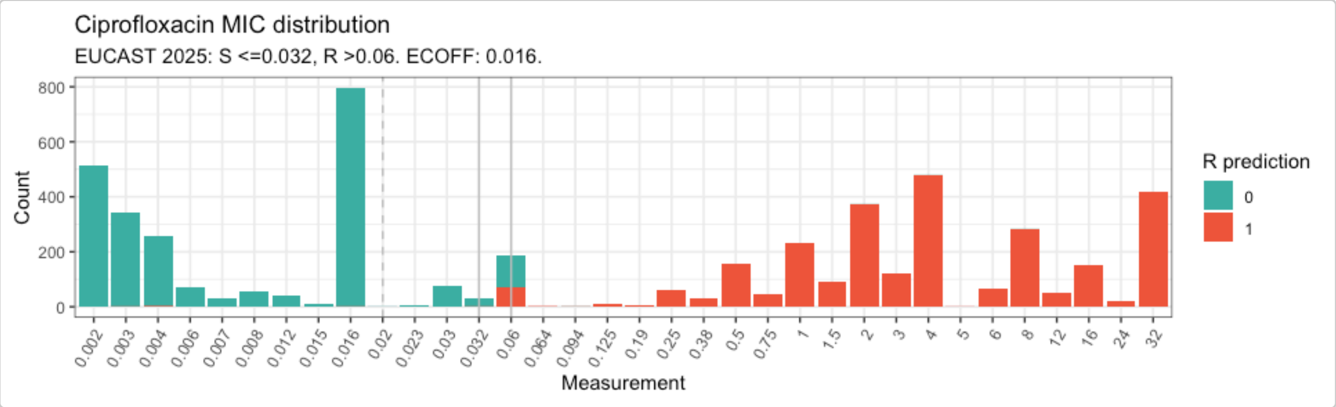


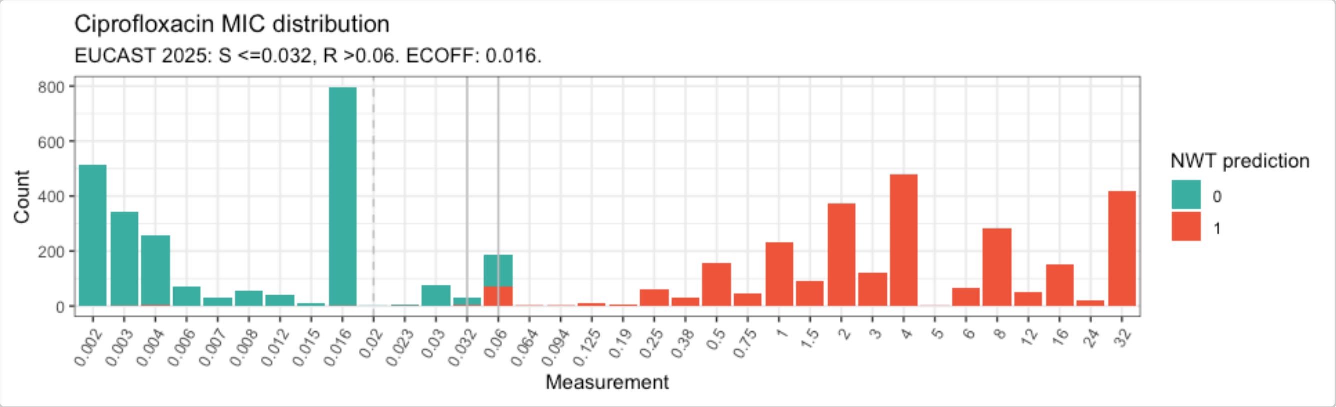


Results demonstrate a very strong genotype-based prediction of ciprofloxacin resistance/susceptibility using the currently available AMR markers.

#### Analysing extended-spectrum cephalosporin genotype-phenotype data

Euro-GASP provides MIC data for both ceftriaxone (first-line monotherapy for gonorrhoea) and cefixime (historically used but largely discarded due to expansion of *penA* mosaics generated through homologous recombination with other *Neisseria* species).

Build binary matrices for each antibiotic:

```
cfm_bin <- get_binary_matrix(
  geno_table = eurogasp_geno,
  pheno_table = eurogasp_ast,
  pheno_drug = "Cefixime",
  geno_class = "Cephalosporins (3rd gen.)",
  sir_col = "pheno_eucast",
  ecoff_col = "ecoff",
  keep_assay_values = TRUE,
  keep_assay_values_from = "mic"
)
#>  Defining NWT in binary matrix using ecoff column provided: ecoff

cro_bin <- get_binary_matrix(
  geno_table = eurogasp_geno,
  pheno_table = eurogasp_ast,
  pheno_drug = "Ceftriaxone",
  geno_class = "Cephalosporins (3rd gen.)",
  sir_col = "pheno_eucast",
  ecoff_col = "ecoff",
  keep_assay_values = TRUE,
  keep_assay_values_from = "mic"
```

```
)
#>  Defining NWT in binary matrix using ecoff column provided: ecoff
```

Generate upset plots:

```
cfm_upset <- amr_upset(
  binary_matrix = cfm_bin,
  min_set_size = 1,
  order = "value",
  assay = "mic",
  pheno_drug = "Cefixime",
  species = "Neisseria gonorrhoeae"
)
#> Ordering markers by frequency
#>   MIC breakpoints determined using AMR package: S <= 0.125 and R > 0.125
#>   MIC breakpoints determined using AMR package: S <= 0.125 and R > 0.125
#> Scale for y is already present.
#> Adding another scale for y, which will replace the existing scale.
#> Scale for y is already present.
#> Adding another scale for y, which will replace the existing scale.
```

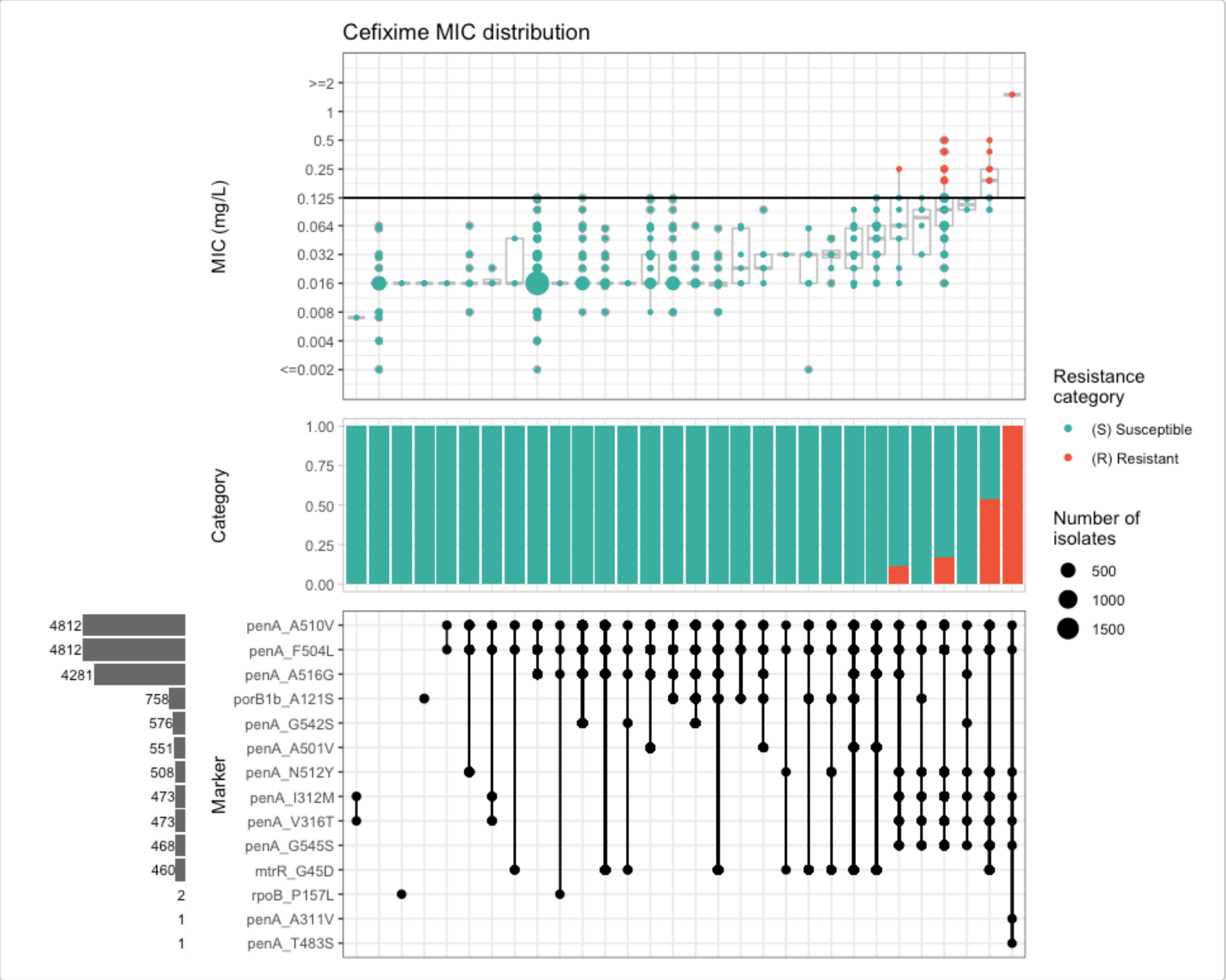


```
cro_upset <- amr_upset(
  binary_matrix = cro_bin,
```

```
  min_set_size = 1,
  order = "value",
  assay = "mic",
  pheno_drug = "Ceftriaxone",
  species = "Neisseria gonorrhoeae"
)
#> Ordering markers by frequency
#>   MIC breakpoints determined using AMR package: S <= 0.125 and R > 0.125
#>   MIC breakpoints determined using AMR package: S <= 0.125 and R > 0.125
#> Scale for y is already present.
#> Adding another scale for y, which will replace the existing scale.Scale for y is
      already present.
#> Adding another scale for y, which will replace the existing scale.
```

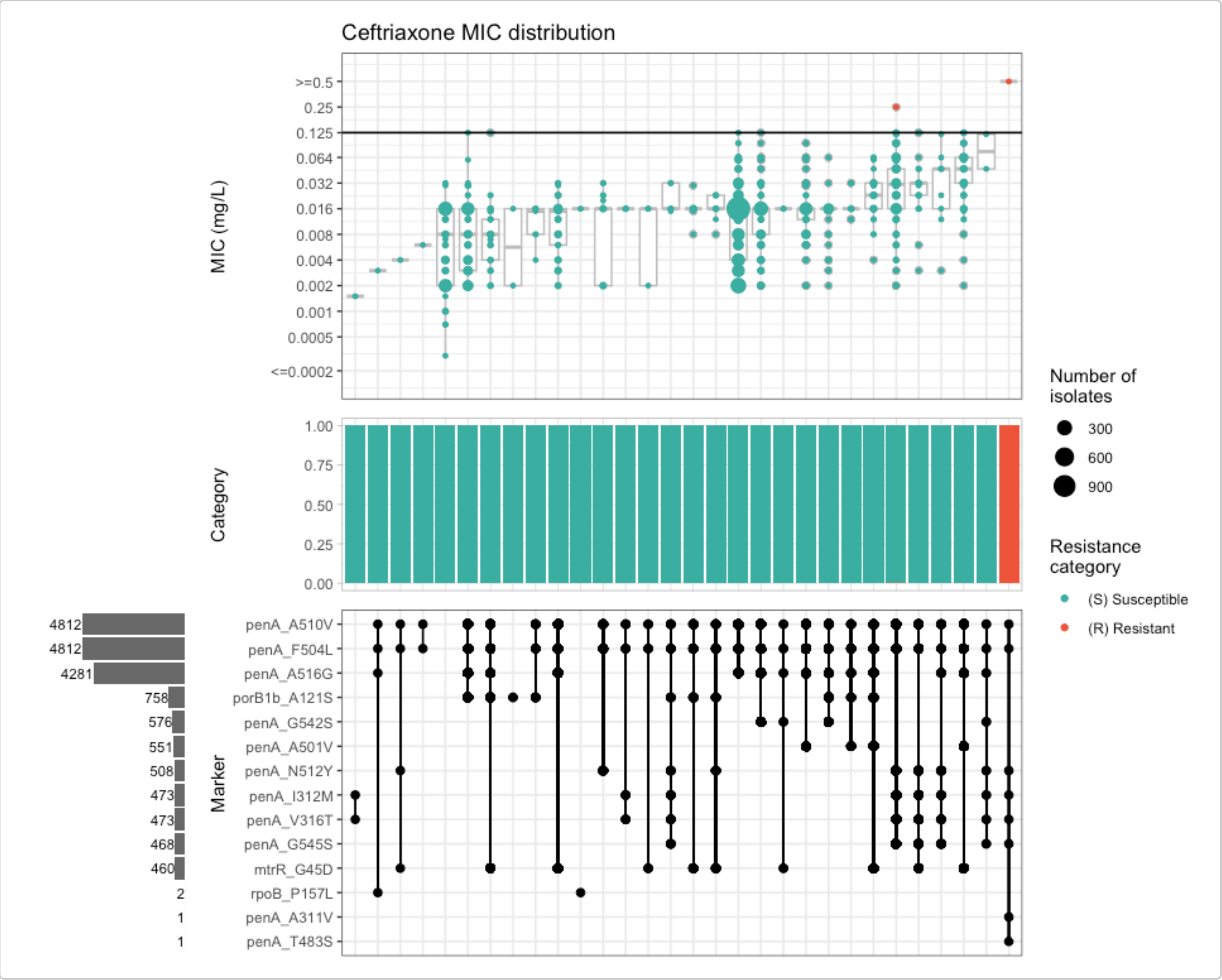


Several mutations in *penA* are associated with increased MICs, however only particular combinations of mutations lead to MICs above the resistant breakpoint. This is very characteristic of *penA* mosaics generated through homologous recombination with other *Neisseria* species. Importantly, not all mosaics increase the MIC above the clinical breakpoint, resulting in a wide MIC range within the susceptible category.

Because most known cephalosporin resistance mutations appear in combination, solo marker analysis provides limited information in this case:

```
cfm_solo_ppv <- solo_ppv(
  binary_matrix = cfm_bin,
  pheno_drug = "Cefixime",
```

```
  geno_class = "Cephalosporins (3rd gen.)",
  sir_col = "pheno_eucast"
)
```

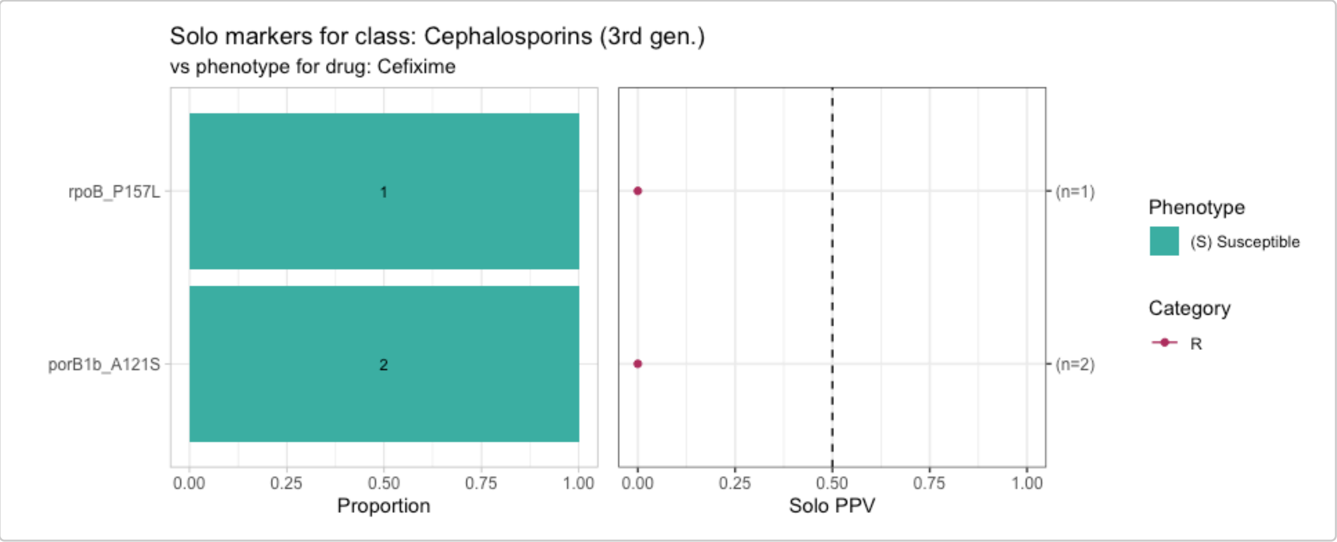


```
cro_solo_ppv <- solo_ppv(
  binary_matrix = cro_bin,
  pheno_drug = "Ceftriaxone",
  geno_class = "Cephalosporins (3rd gen.)",
  sir_col = "pheno_eucast"
)
```

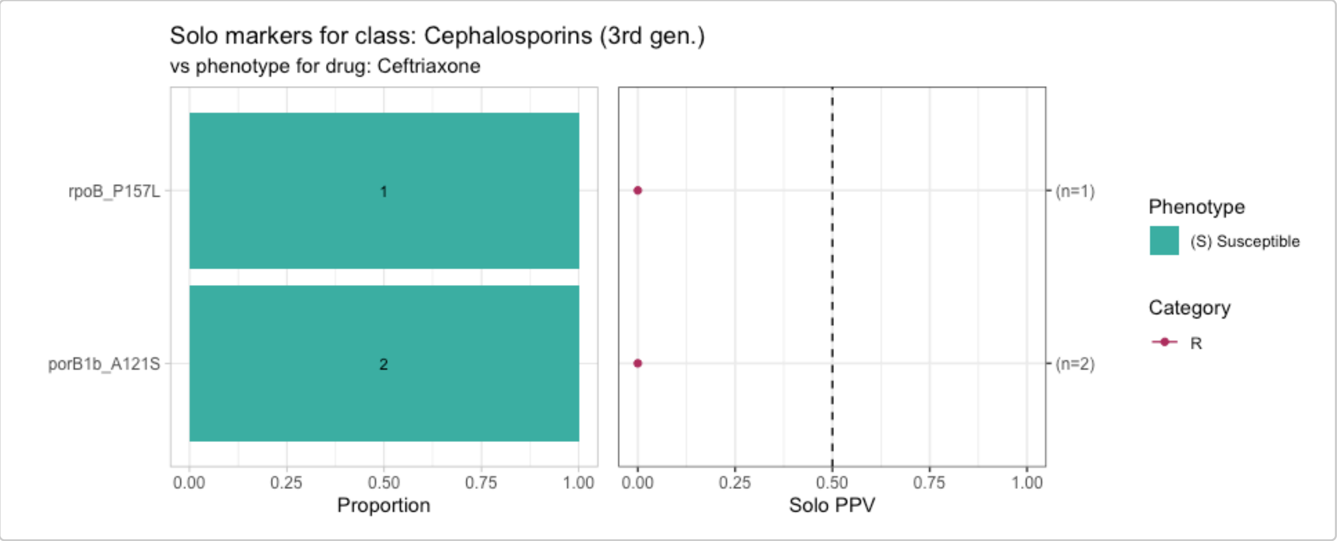


Instead, combination PPVs are a more informative metric:

```
cfm_ppv <- amr_ppv(
  binary_matrix = cfm_bin,
  min_set_size = 2,
  order = "ppv",
  upset_grid = TRUE,
  plot_assay = TRUE,
  assay = "mic"
)
#> Ordering markers by frequency
```

```
#> Scale for y is already present.
#> Adding another scale for y, which will replace the existing scale.
```

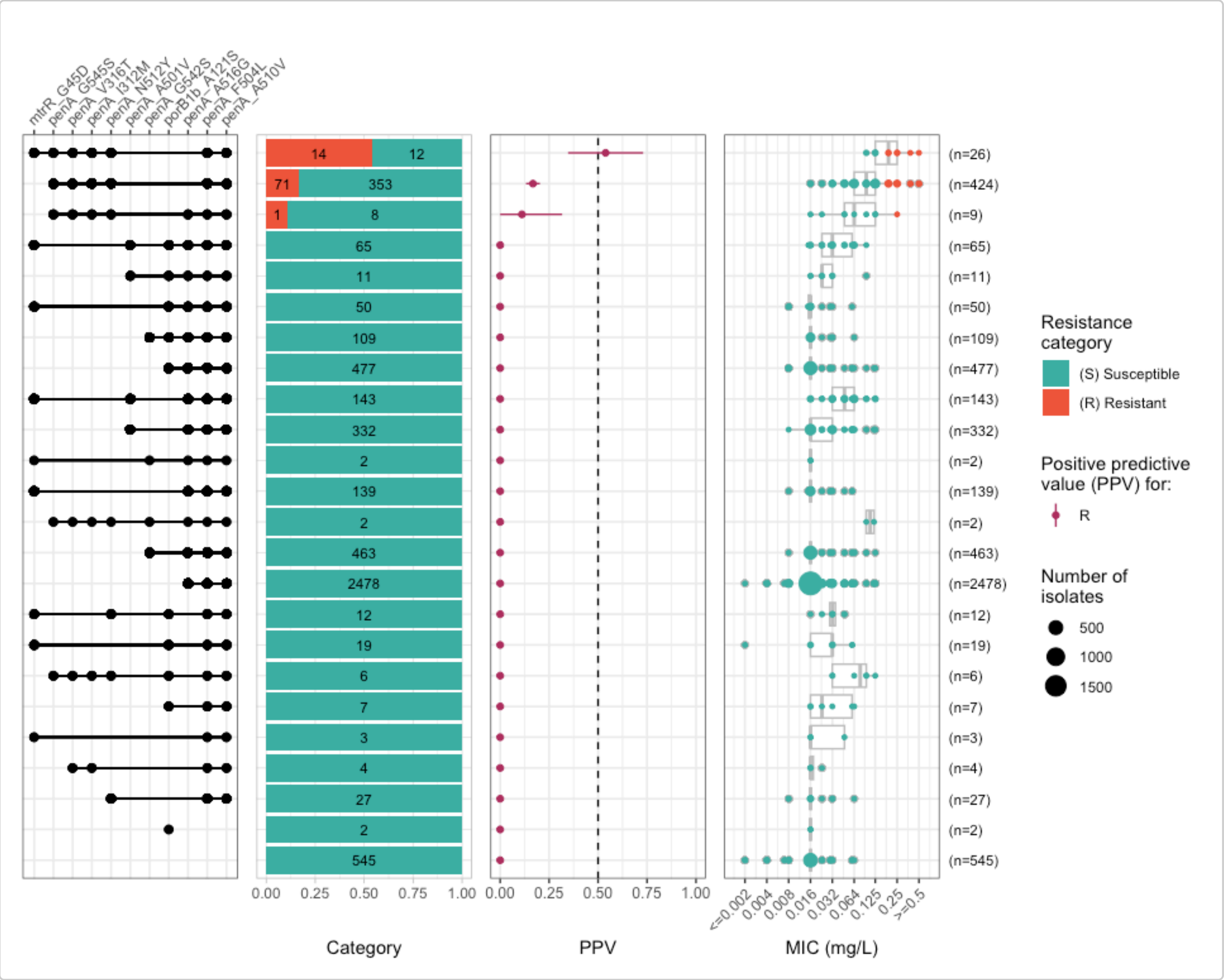


```
cro_ppv <- amr_ppv(
  binary_matrix = cro_bin,
  min_set_size = 1,
  order = "ppv",
  upset_grid = TRUE,
  plot_assay = TRUE,
  assay = "mic"
)
#> Ordering markers by frequency
#> Scale for y is already present.
#> Adding another scale for y, which will replace the existing scale.
```

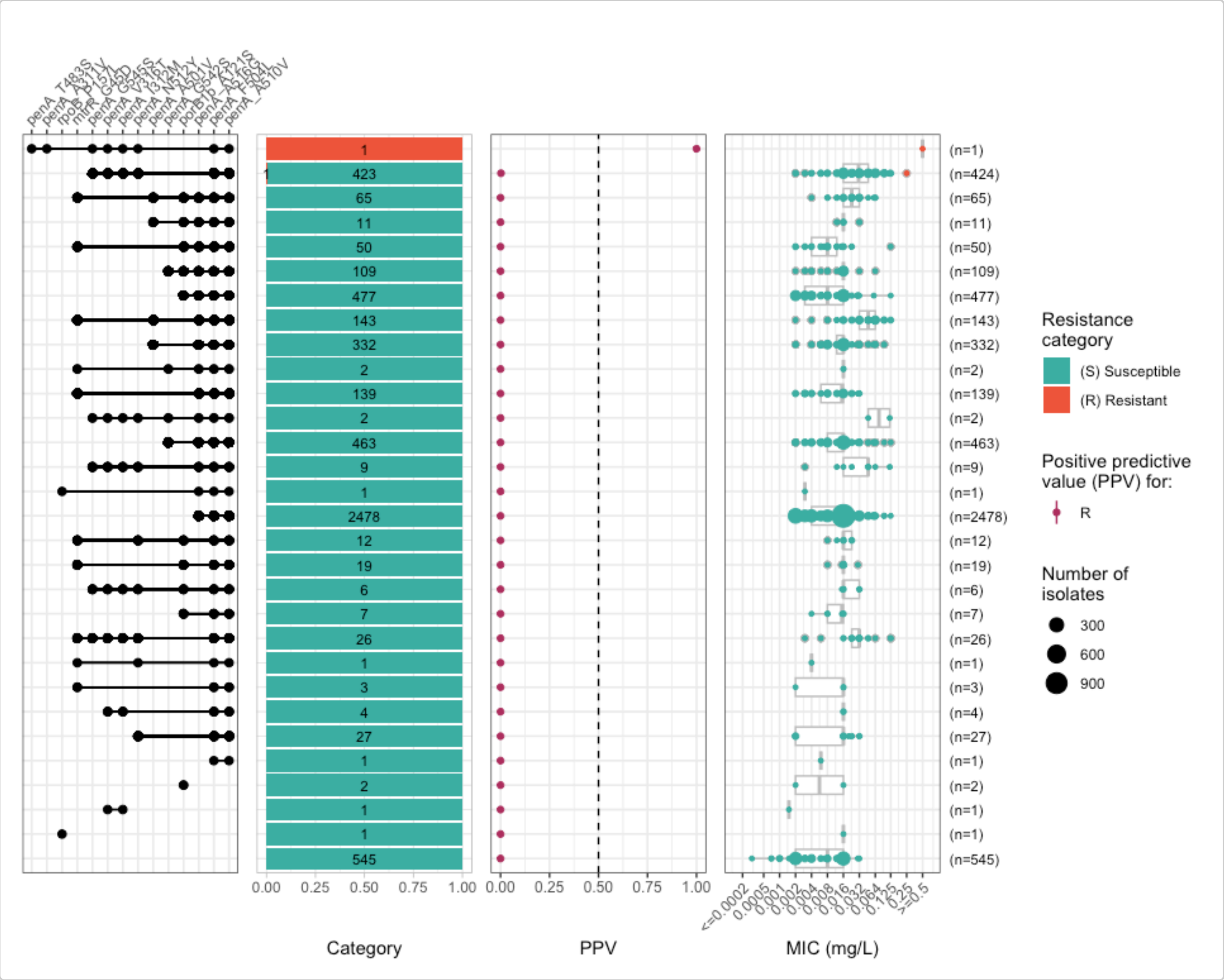


Due to the very limited number of isolates classified as resistant to cefixime or ceftriaxone in this dataset, logistic regression analyses are not feasible here.

#### Use case 2: Study of mosaic PBP2 mutations associated with decreased susceptibility and resistance to ceftriaxone

In this use case, we explore combinations of PBP2 mutations and their associated MIC ranges for mosaic *penA* variants, using a broader dataset that enriches for ceftriaxone-resistant and decreased-susceptibility isolates. The dataset combines AMRFinderPlus results from the following sources (total n = 2,101 genomes):

- Euro-GASP 2020 genomic survey by Golparian *et al.* (2024).
  - ENA PRJEB58139, n=1,932 genomes.
- Ceftriaxone-resistant gonococci from the United Kingdom, by Day *et al.* (2022).
  - ENA PRJEB57389, n=8 genomes.
- Ceftriaxone-resistant gonococci from England, by Fifer *et al.* (2024).
  - ENA PRJEB76977, n=19 genomes.
- Ceftriaxone-resistant gonococci from Asia, by van der Veen *et al.* 2026.
  - ENA PRJEB45627, PRJNA577446, PRJNA776899, PRJNA778600, PRJNA560592, PRJNA874857, PRJNA909328, PRJNA957547, PRJNA1161034, PRJNA1189294, n=113 genomes.
- World Health Organization (WHO) reference genomes, by Unemo *et al.* (2024).
  - ENA PRJNA1067895, n=29 genomes.

Import and format the data:

```r
# Genotype file
ngono_cro_geno <- import_amrfp(ngono_cro_geno_raw, "Name")
#> Input file lacks the expected column: 'Type' (v4.0+) or 'Element type' (pre-v4),
        assuming all rows report AMR markers.

# Phenotype file
ngono_cro_pheno <- ngono_cro_pheno_raw %>%
  pivot_longer(
    cols = c(Ceftriaxone),
    names_to = "drug",
    values_to = "mic"
  )

ngono_cro_ast <- format_pheno(
  input = ngono_cro_pheno,
  sample_col = "id",
  species = "Neisseria gonorrhoeae",
  ab_col = "drug",
  mic_col = "mic",
  interpret_eucast = TRUE
)
#> Adding new micro-organism column 'spp_pheno' (class 'mo') with constant value Neisseria
        gonorrhoeae
#> Parsing column spp_pheno as micro-organism (class 'mo')
#> Parsing column drug as antibiotic (class 'ab')
#> Parsing column mic as class 'mic'
#> Could not find disk_col disk in input table
#> Could not find pheno_col ecoff in input table
#> Could not find pheno_col pheno_eucast in input table
#> Could not find pheno_col pheno_clsi in input table
#> Could not find pheno_col pheno_provided in input table
#> Could not find method_col method in input table
#> Could not find platform_col platform in input table
#> Could not find guideline_col guideline in input table
#> Could not find source_col source in input table
#> Interpreting all data as species: Neisseria gonorrhoeae

# Include empty rows for samples with phenotype but no genotype data
negative_cro <- ngono_cro_pheno_raw %>%
  anti_join(ngono_cro_geno) %>%
  pull(id)
#> Joining with `by = join_by(id)`

ngono_cro_geno <- ngono_cro_geno %>% bind_rows(tibble(id = negative_cro))
```

Build the binary matrix and generate upset plots:

```r
# Get binary matrix
cro_bin_2 <- get_binary_matrix(
  geno_table = ngono_cro_geno,
  pheno_table = ngono_cro_ast,
  pheno_drug = "Ceftriaxone",
  geno_class = "Cephalosporins (3rd gen.)",
  sir_col = "pheno_eucast",
  keep_assay_values = TRUE,
```

```
  keep_assay_values_from = "mic"
)
#>  Defining NWT in binary matrix as I/R vs S, as no ECOFF column defined

# Calculate upset plots of MIC vs genotype marker combinations
cro_upset_2 <- amr_upset(
  binary_matrix = cro_bin_2,
  min_set_size = 1,
  order = "value",
  assay = "mic",
  pheno_drug = "Ceftriaxone",
  species = "Neisseria gonorrhoeae"
)
#> Ordering markers by frequency
#>   MIC breakpoints determined using AMR package: S <= 0.125 and R > 0.125
#>   MIC breakpoints determined using AMR package: S <= 0.125 and R > 0.125
#> Scale for y is already present.
#> Adding another scale for y, which will replace the existing scale.
#> Scale for y is already present.
#> Adding another scale for y, which will replace the existing scale.
```

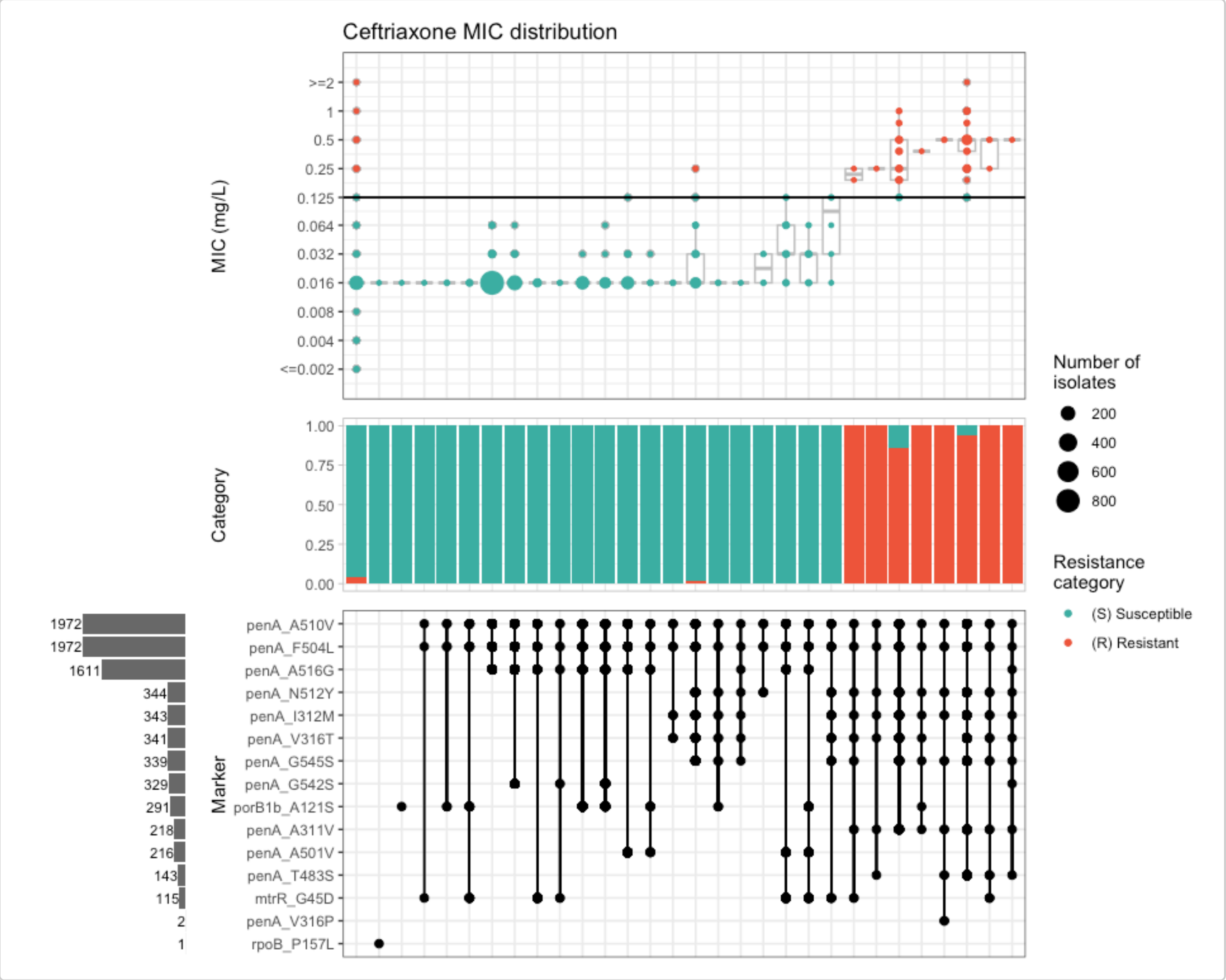


Calculate combination PPVs:

```
cro_ppv_2 <- amr_ppv(
  binary_matrix = cro_bin_2,
```

```
  min_set_size = 1,
  order = "ppv",
  upset_grid = TRUE,
  plot_assay = TRUE,
  assay = "mic"
)
#> Ordering markers by frequency
#> Scale for y is already present.
#> Adding another scale for y, which will replace the existing scale.
```

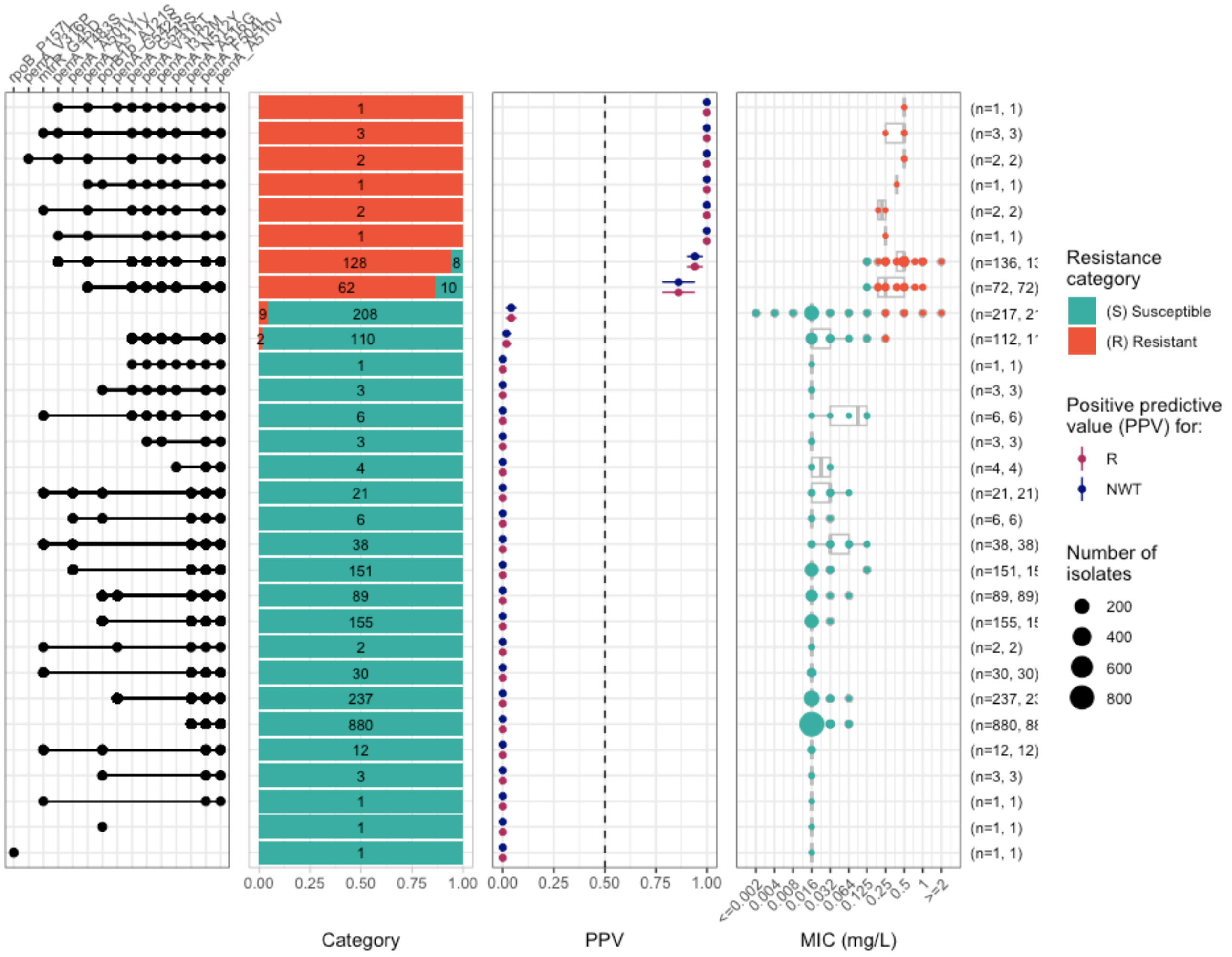


Run logistic regression to evaluate individual marker contributions:

```
cro_logist <- amr_logistic(
  binary_matrix = cro_bin_2,
  pheno_drug = "Ceftriaxone",
  sir_col = "pheno_eucast",
  ecoff_col = "ecoff",
  fit_glm = TRUE,
  maf = 10,
  single_plot = TRUE
)
#> ...Fitting logistic regression model to R using glm
#>    Filtered data contains 2191 samples (211 => 1, 1980 => 0) and 13 variables.
#> Waiting for profiling to be done...
#> ...Fitting logistic regression model to NWT using glm
```

```
#>    Filtered data contains 2191 samples (211 => 1, 1980 => 0) and 13 variables.
#> Waiting for profiling to be done...
#> Generating plots
#> Plotting 2 models
```

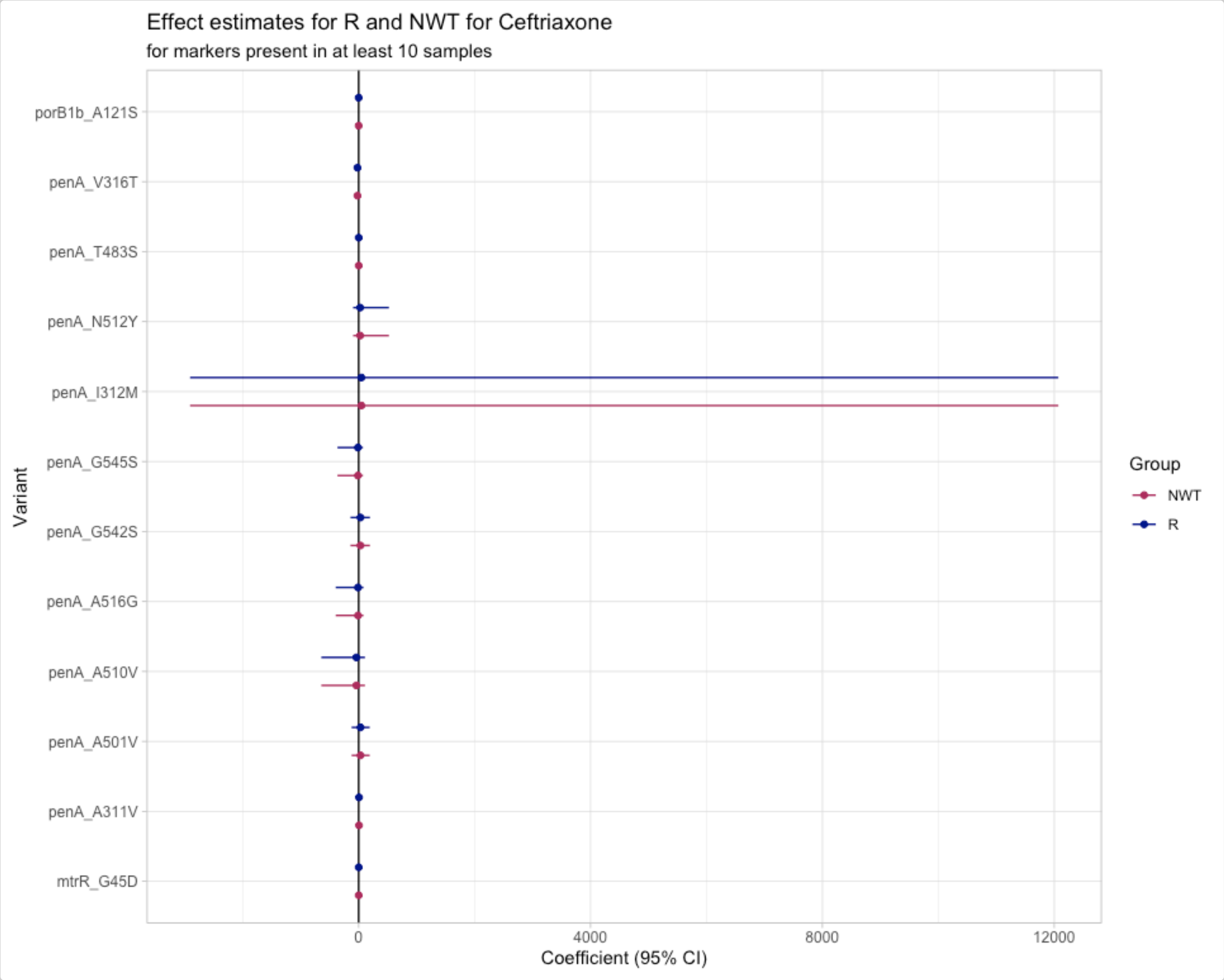


Results confirm that individual *penA* mutations do not independently increase cephalosporin MICs; rather, specific combinations of mutations are required. *penA* I312M has a very wide coefficient interval, likely reflecting their contribution to *penA* mosaics associated with resistance.

#### Use case 3: Investigation of tetracycline resistance and implications for STI prevention strategies

Doxy-PEP (doxycycline post-exposure prophylaxis) involves taking a single dose of doxycycline within 24–72 hours after a sexual risk event to prevent STIs. While evidence supports its short-term efficacy, a modelling study by Reichert *et al.* (2026) suggests that broad doxy-PEP implementation may select resistant lineages, potentially limiting its long-term effectiveness against gonorrhoea.

Here, we analyse 409 *N. gonorrhoeae* genomes from isolates collected in eastern Spain between 2021–2024 (Sánchez-Serrano *et al.*, 2026, ENA PRJEB83795), with available tetracycline MIC data, to explore genetic determinants of tetracycline resistance in a local gonococcal population.

Import and format phenotype data:

```
ngono_tet_pheno <- ngono_tet_pheno_raw %>%
  pivot_longer(
```

```
    cols = c(Tetracycline),
    names_to = "drug",
    values_to = "mic"
  )

ngono_tet_ast <- format_pheno(
  input = ngono_tet_pheno,
  sample_col = "id",
  species = "Neisseria gonorrhoeae",
  ab_col = "drug",
  mic_col = "mic",
  interpret_eucast = TRUE,
  interpret_ecoff = TRUE
)
#> Adding new micro-organism column 'spp_pheno' (class 'mo') with constant value Neisseria
        gonorrhoeae
#> Parsing column spp_pheno as micro-organism (class 'mo')
#> Parsing column drug as antibiotic (class 'ab')
#> Parsing column mic as class 'mic'
#> Could not find disk_col disk in input table
#> Could not find pheno_col ecoff in input table
#> Could not find pheno_col pheno_eucast in input table
#> Could not find pheno_col pheno_clsi in input table
#> Could not find pheno_col pheno_provided in input table
#> Could not find method_col method in input table
#> Could not find platform_col platform in input table
#> Could not find guideline_col guideline in input table
#> Could not find source_col source in input table
#> Interpreting all data as species: Neisseria gonorrhoeae
```

Import genotype data and add empty rows for samples with phenotype information but for which no genetic AMR determinants are identified by AMRFinderPlus. For this example, there are 409 samples with MIC in the phenotype file but AMR determinants were only found for 402.

```
ngono_tet_geno <- import_amrfp(
  input_table = ngono_tet_geno_raw,
  sample_col = "Name"
)

negative_samples <- ngono_tet_ast %>%
  anti_join(ngono_tet_geno) %>%
  pull(id)
#> Joining with `by = join_by(id, drug)`

ngono_tet_geno <- ngono_tet_geno %>% bind_rows(tibble(id = negative_samples))
```

Build the binary matrix and generate upset plots:

```
tet_bin <- get_binary_matrix(
  geno_table = ngono_tet_geno,
  pheno_table = ngono_tet_ast,
  pheno_drug = "Tetracycline",
  geno_class = "Tetracyclines",
  ecoff_col = "ecoff",
```

```
  sir_col = "pheno_eucast",
  keep_assay_values = TRUE,
  keep_assay_values_from = "mic"
)
#>  Defining NWT in binary matrix using ecoff column provided: ecoff

tet_upset <- amr_upset(
  binary_matrix = tet_bin,
  assay = "mic",
  min_set_size = 2,
  order = "value",
  bp_R = 0.5,
  plot_set_size = TRUE,
  print_set_size = TRUE,
  print_category_counts = TRUE
)
#> Ordering markers by frequency
#> Scale for y is already present.
#> Adding another scale for y, which will replace the existing scale.
#> Scale for y is already present.
#> Adding another scale for y, which will replace the existing scale.
```

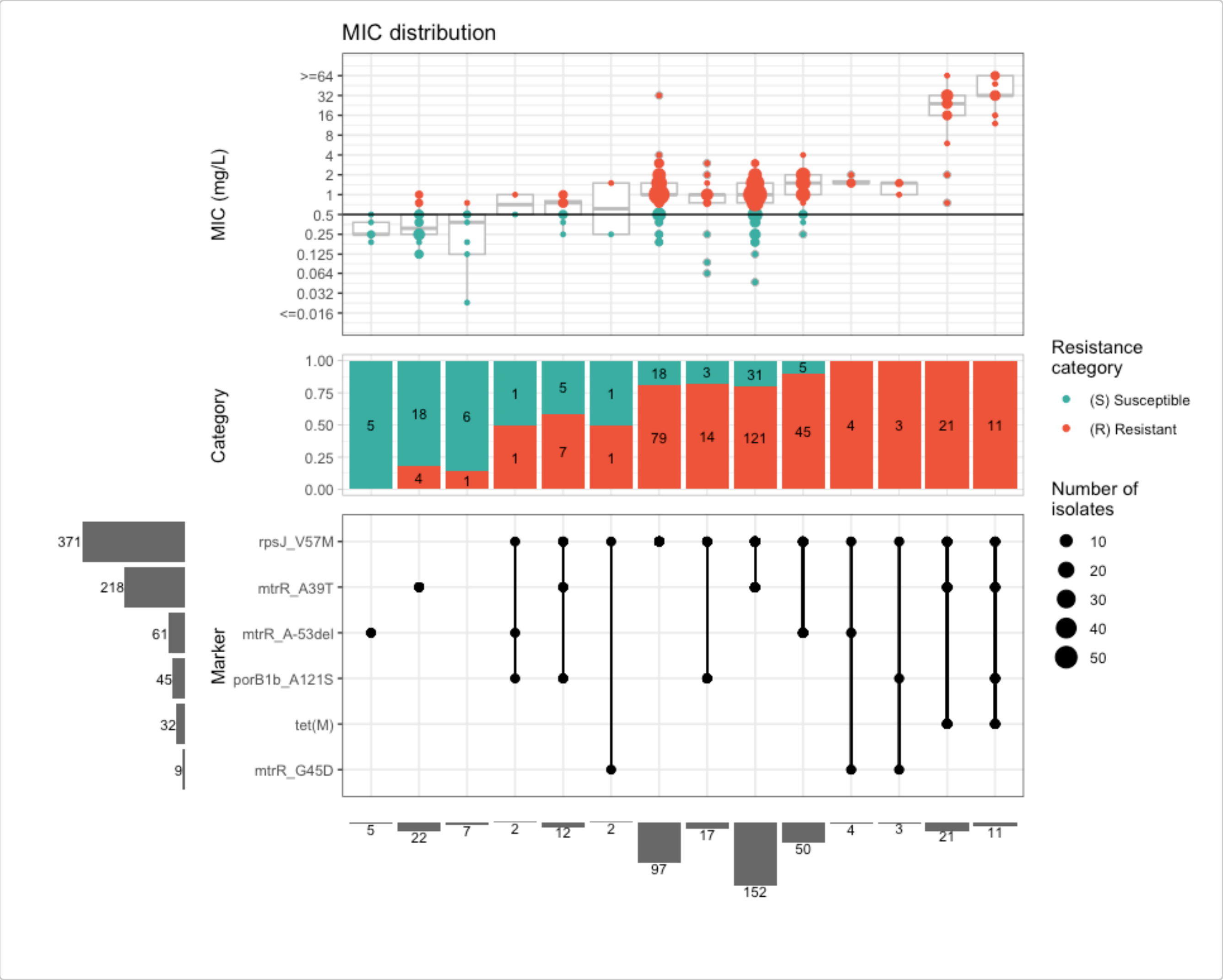


The upset plot shows that MIC increases are primarily explained by the chromosomal *rpsJ* V57M mutation and the *tet(M)* gene, which is carried on a conjugative plasmid.

Calculate prevalence of the two key resistance determinants in the studied population:

```r
pop_size <- 409

ngono_tet_geno %>%
  count(marker) %>%
  filter(marker %in% c("rpsJ_V57M", "tet(M)")) %>%
  mutate(percent = round(100 * n / pop_size, 1))
#> # A tibble: 2 x 3
#>   marker        n percent
#>   <chr>     <int>   <dbl>
#> 1 rpsJ_V57M   373    91.2
#> 2 tet(M)       34     8.3
```

Of the 409 isolates with tetracycline phenotypic data:

- **373 (91.2%)** carry the *rpsJ* V57M chromosomal mutation.
- **34 (8.3%)** carry the *tet(M)* gene on the conjugative plasmid.

Explore marker combinations in the upset summary:

```r
tet_upset$summary %>%
  arrange(desc(marker_count)) %>%
  filter(grepl("rpsJ_V57M", marker_list))
#> # A tibble: 13 x 16
#>    marker_list          marker_count     n combination_id   R.n R.ppv R.ci_lower
#>    <chr>                       <dbl> <int> <fct>          <dbl> <dbl>      <dbl>
#>  1 mtrR_A39T, rpsJ_V57…            4    11 1_1_1_1_0_0       11 1          1
#>  2 rpsJ_V57M, mtrR_A-5…            3     4 0_1_0_0_1_1        4 1          1
#>  3 rpsJ_V57M, porB1b_A…            3     3 0_1_0_1_0_1        3 1          1
#>  4 rpsJ_V57M, porB1b_A…            3     2 0_1_0_1_1_0        1 0.5        0
#>  5 rpsJ_V57M, tet(M), …            3     1 0_1_1_0_1_0        1 1          1
#>  6 mtrR_A39T, rpsJ_V57…            3    12 1_1_0_1_0_0        7 0.583      0.304
#>  7 mtrR_A39T, rpsJ_V57…            3    21 1_1_1_0_0_0       21 1          1
#>  8 rpsJ_V57M, mtrR_G45D            2     2 0_1_0_0_0_1        1 0.5        0
#>  9 rpsJ_V57M, mtrR_A-5…            2    50 0_1_0_0_1_0       45 0.9        0.817
#> 10 rpsJ_V57M, porB1b_A…            2    17 0_1_0_1_0_0       14 0.824      0.642
#> 11 rpsJ_V57M, tet(M)               2     1 0_1_1_0_0_0        1 1          1
#> 12 mtrR_A39T, rpsJ_V57M            2   152 1_1_0_0_0_0      121 0.796      0.732
#> 13 rpsJ_V57M                       1    97 0_1_0_0_0_0       79 0.814      0.737
#> # ℹ 9 more variables: R.ci_upper <dbl>, R.denom <int>,
#> #   median_excludeRangeValues <dbl>, q25_excludeRangeValues <dbl>,
#> #   q75_excludeRangeValues <dbl>, n_excludeRangeValues <int>,
#> #   median_ignoreRanges <dbl>, q25_ignoreRanges <dbl>, q75_ignoreRanges <dbl>

tet_upset$summary %>%
  arrange(desc(marker_count)) %>%
  filter(grepl("tet\\(M\\)", marker_list))
#> # A tibble: 4 x 16
#>   marker_list           marker_count     n combination_id   R.n R.ppv R.ci_lower
#>   <chr>                        <dbl> <int> <fct>          <dbl> <dbl>      <dbl>
#> 1 mtrR_A39T, rpsJ_V57M…            4    11 1_1_1_1_0_0       11     1          1
#> 2 rpsJ_V57M, tet(M), m…            3     1 0_1_1_0_1_0        1     1          1
#> 3 mtrR_A39T, rpsJ_V57M…            3    21 1_1_1_0_0_0       21     1          1
#> 4 rpsJ_V57M, tet(M)                2     1 0_1_1_0_0_0        1     1          1
#> # ℹ 9 more variables: R.ci_upper <dbl>, R.denom <int>,
```

```
#> #   median_excludeRangeValues <dbl>, q25_excludeRangeValues <dbl>,
#> #   q75_excludeRangeValues <dbl>, n_excludeRangeValues <int>,
#> #   median_ignoreRanges <dbl>, q25_ignoreRanges <dbl>, q75_ignoreRanges <dbl>
```

Calculate PPVs for marker combinations:

```
# add species and pheno_drug to retrieve and plot breakpoint
tet_ppv <- amr_ppv(
  binary_matrix = tet_bin,
  min_set_size = 1,
  order = "ppv",
  upset_grid = TRUE,
  plot_assay = TRUE,
  assay = "mic",
  species = "Neisseria gonorrhoeae",
  pheno_drug = "Tetracycline"
)
#> Ordering markers by frequency
#>   MIC breakpoints determined using AMR package: S <= 0.5 and R > 0.5
#>   MIC breakpoints determined using AMR package: S <= 0.5 and R > 0.5
#> Scale for y is already present.
#> Adding another scale for y, which will replace the existing scale.
```

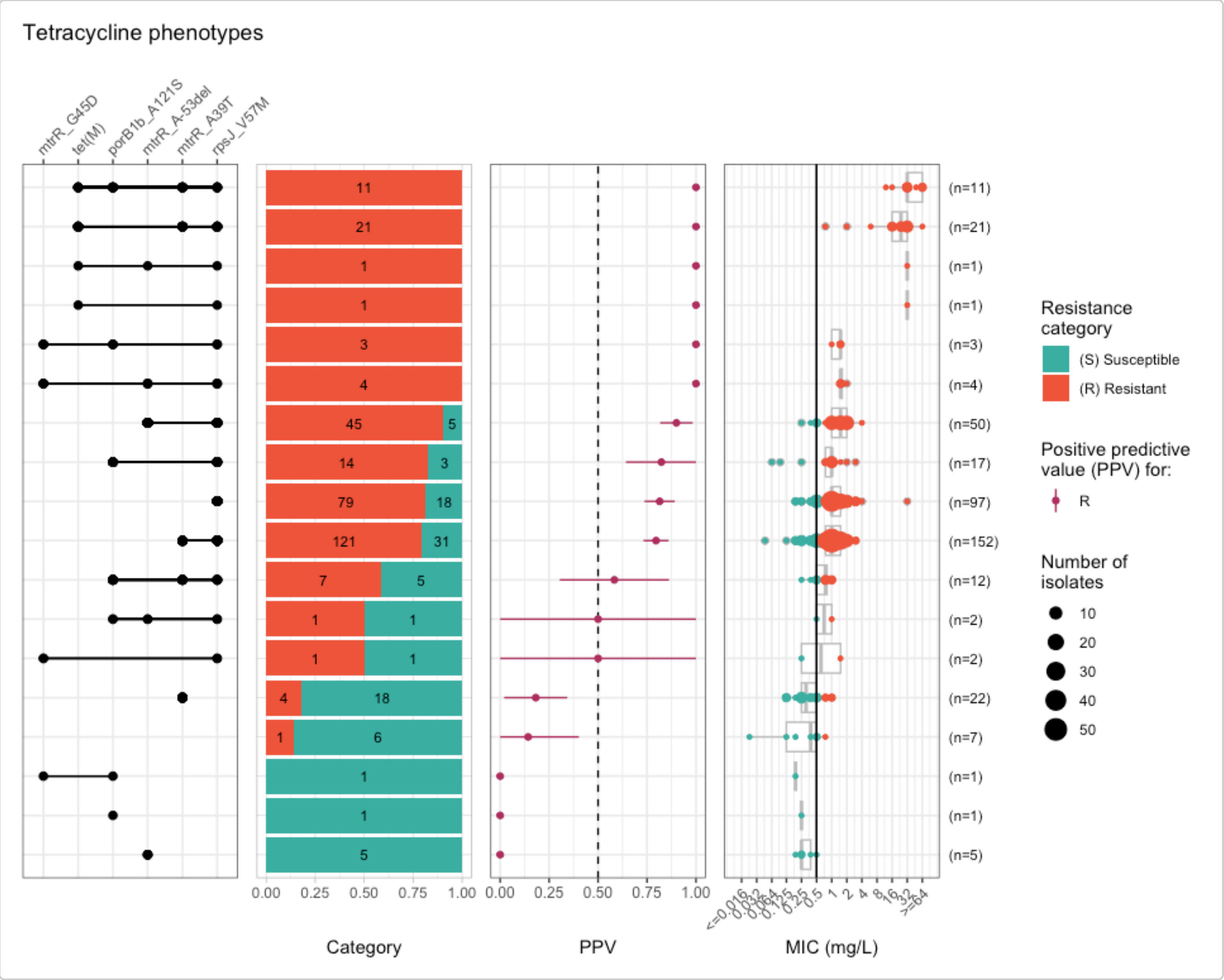


Run logistic regression to confirm the independent contributions of *rpsJ* V57M and *tet(M)*:

```
tet_logist <- amr_logistic(
  binary_matrix = tet_bin,
  pheno_drug = "Tetracycline",
  geno_class = "Tetracyclines",
  sir_col = "pheno_eucast",
  ecoff_col = "ecoff",
  maf = 10,
  single_plot = TRUE
)
#> ...Fitting logistic regression model to R using logistf
#>    Filtered data contains 409 samples (314 => 1, 95 => 0) and 6 variables.
#> ...Fitting logistic regression model to NWT using logistf
#> Generating plots
#> Plotting R model only
```

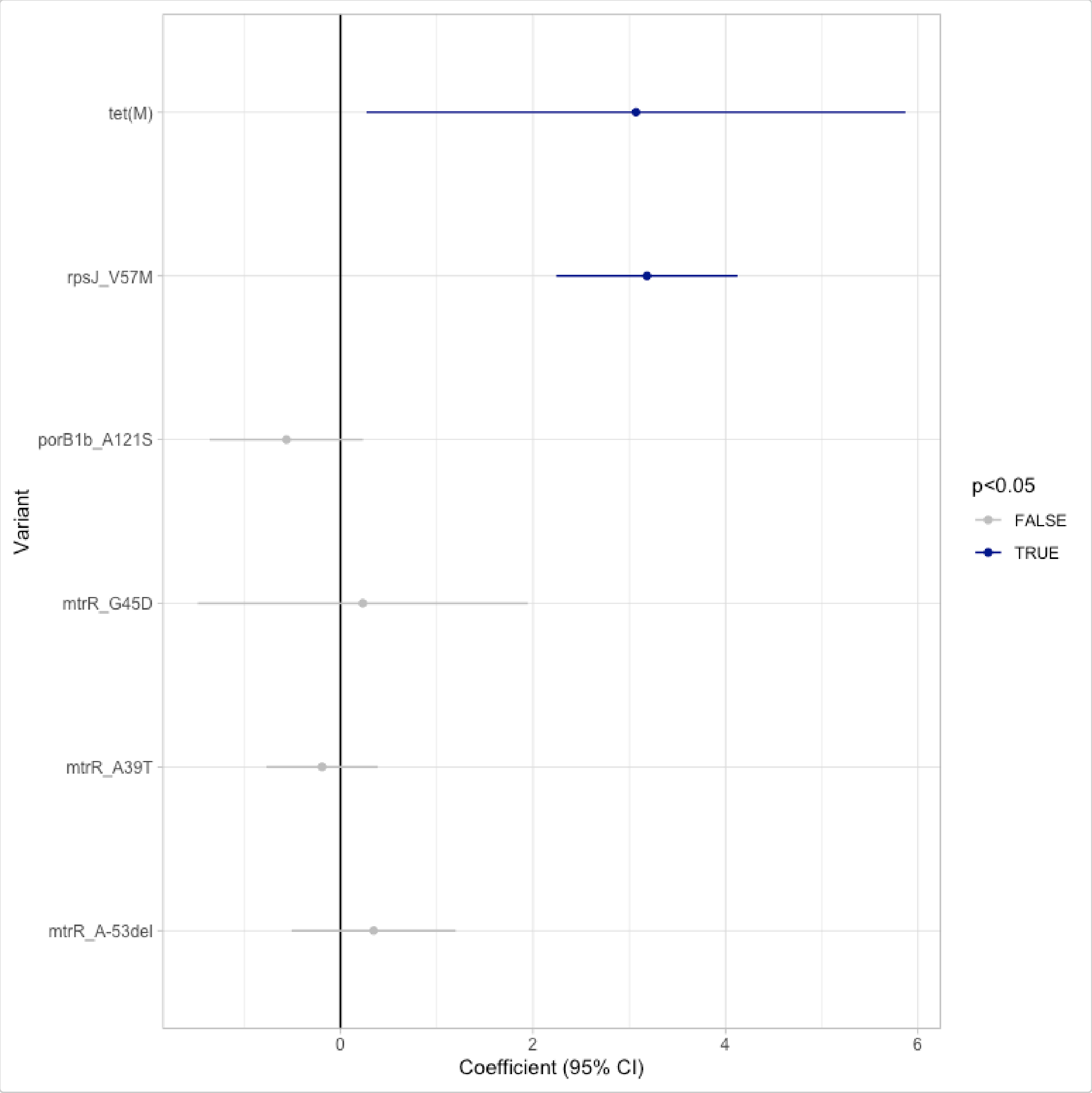


As there is no ECOFF defined for tetracycline in *N. gonorrhoeae*, we use only clinical breakpoints (`truth = "R"`) in the concordance analysis:

```r
tet_concordance <- concordance(
  binary_matrix = tet_bin,
  ppv_results = tet_ppv,
  prediction_rule = "logistic",
  logreg_results = tet_logist,
  truth = "R"
)
tet_concordance
#> AMR Genotype-Phenotype Concordance
#> Prediction rule: logistic
#>
#> --- Outcome: R ---
#> Samples: 409 | Markers: 6
#> Markers used: mtrR_A39T, rpsJ_V57M, tet(M), porB1b_A121S, mtrR_A-53del, mtrR_G45D
#>
#> Confusion Matrix:
#>            Truth
#> Prediction   1   0
#>          1 309  64
#>          0   5  31
#>
#> Metrics:
#>   Sensitivity : 0.9841
#>   Specificity : 0.3263
#>   PPV         : 0.8284
#>   NPV         : 0.8611
#>   Accuracy    : 0.8313
#>   Kappa       : 0.3962
#>   F-measure   : 0.8996
#>   VME         : 0.0159
#>   ME          : 0.6737
```

The current marker set achieves 98.72% sensitivity and 82.84% PPV, but only 28.09 % specificity. This is reflected in a high major error rate (ME = 71.91%), meaning many susceptible isolates are predicted as resistant — a consequence of the very high prevalence of *rpsJ* V57M in isolates that have elevated MICs but remain below the clinical breakpoint.

Visualise the predictions on the MIC distribution:

```r
ngono_tet_pred <- ngono_tet_ast %>%
  left_join(tet_concordance$data[, c("id", "R_pred")],
    by = "id"
  )

assay_by_var(
  pheno_table = ngono_tet_pred,
  pheno_drug = "Tetracycline",
  measure = "mic",
  colour_by = "R_pred",
  species = "Neisseria gonorrhoeae",
  colours = c("#3CAEA3", "#ED553B"),
  colour_legend_label = "R prediction"
)
#>   MIC breakpoints determined using AMR package: S <= 0.5 and R > 0.5
```

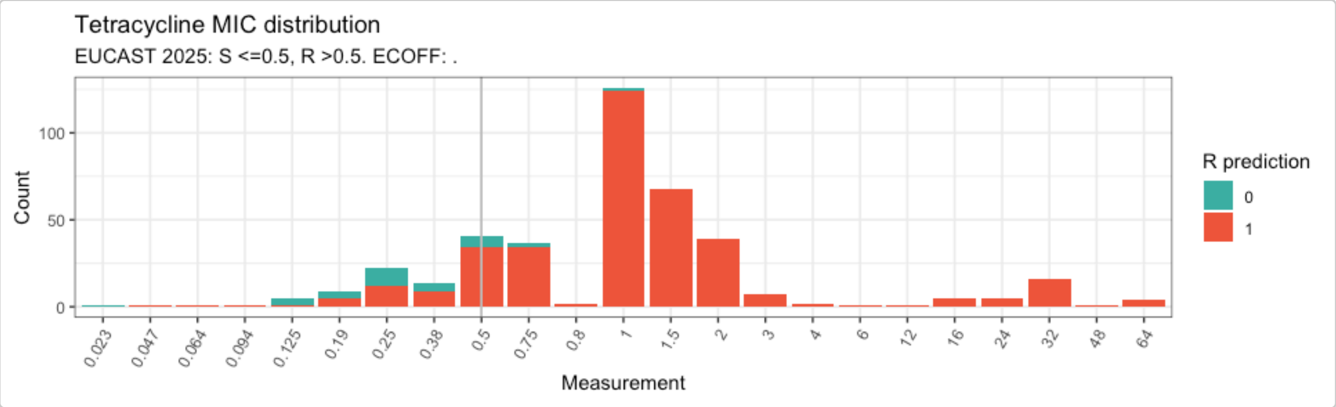


The high prevalence of *rpsJ* V57M (>90% of the population) combined with a significant prevalence of the plasmid-borne *tet(M)* gene — which can be readily acquired by other circulating lineages through conjugation — means that doxy-PEP is likely to select for these pre-existing resistant lineages. Together, these findings support the conclusion that doxy-PEP is unlikely to provide a durable long-term solution for reducing the burden of gonorrhoea infections.

### Example with custom stratification by isolate source

### Analysing a small user-generated geno-pheno dataset of *Salmonella enterica*

#### Introduction

This vignette shows how `AMRgen` can be used to explore a small user-generated dataset with genotypic and phenotypic data for non-typhoidal, non-invasive *Salmonella enterica* isolates. Phenotypic data consists of MIC E-test results for ciprofloxacin, levofloxacin, and moxifloxacin. Genotypic data includes AMRFinderPlus results for quinolone resistance markers.

Start by loading the package:

```
library(AMRgen)
library(dplyr)
library(ggplot2)
library(tidyr)
```

#### Import data

In this example, the data table was exported from the user's laboratory information management system as a single `.csv` file that includes both genotypic and phenotypic results for 115 *S. enterica* isolates, with one line per isolate. There are columns for two categorical variables of interest: isolate source (human, animal, or other) and isolate serovar. Genotype data is stored in a single column, and where multiple quinolone resistance markers were identified, these are listed in a string with semicolon delimiters. Phenotype data is stored in separate columns for each antimicrobial agent that was tested (ciprofloxacin, levofloxacin, moxifloxacin).

Data in this format can be read into R using `read_csv()` and then formatted for use with `AMRgen`. Here, the object `salm_raw` is already pre-loaded with the package so there is no need to read in the `.csv` file.

```
# Importing raw data from .csv file (not needed to run vignette)
salm_raw <- read_csv("Salmonella_pheno_geno_data.csv")
```

```
head(salm_raw)
#> # A tibble: 6 × 7
#>   Sample Source Serovar     CpL_Genotype Ciprofloxacin Levofloxacin Moxifloxacin
#>   <chr>  <chr>  <chr>       <chr>        <chr>         <chr>        <chr>
#> 1 SAL001 Animal other       gyrA_S83F;p… 0.19          0.5          0.5
#> 2 SAL002 Human  S. Enterit… gyrA_S83Y    0.38          1.5          1.5
#> 3 SAL003 Human  other       gyrA_S83F;p… 3             12           24
#> 4 SAL004 other  S. Infantis gyrA_D87G    0.125         0.5          0.5
```

```
#> 5 SAL005 Human  other       gyrA_D87Y    0.094         0.25         0.38
#> 6 SAL006 Human  S. Kentucky gyrA_S83F;g… 8             8            32
```

The AMRgen package requires separate phenotypic and genotypic tables with specific column headers, so the next step is to separate and wrangle these data into the required formats.

### Genotype table

#### Format raw genotype data for AMRgen

To use downstream AMRgen functions, a genotype table must have at a minimum the following columns: Name (unique sample name for each isolate), marker (name of the resistance marker detected), and drug_class (antibiotic class associated with this marker). It also needs to be in long form, with a row for each isolate/marker combination. In our dataset, the genotype information is found in a single column alongside the metadata and phenotypic information, and we can have multiple markers per row (up to four). Here, we carry out the following steps to generate a genotype table that is compatible with AMRgen functions:

- Select only the genotype column (called CpL_Genotype in this dataset) and the column specifying the individual isolate names (called Sample in this dataset)
- Rename the column Sample to Name, which is the default column heading used by AMRgen for identifying individual isolates
- Separate the genotype column CpL_Genotype into individual columns for each marker (up to four per isolate in these data), specifying the delimiter ;
- Convert this wide-form data (one row per isolate) to a long-form table (one row per isolate/marker combination) using pivot_longer
- Add a column specifying that all these markers are associated with resistance to quinolones

```
# Extract geno data, separate by delimiter, pivot longer, and add drug_class column
salm_geno <- salm_raw %>%
  select(Sample, CpL_Genotype) %>%
  rename(Name = Sample) %>%
  separate_wider_delim(CpL_Genotype,
    delim = ";", names = c("Marker_1", "Marker_2", "Marker_3", "Marker_4"),
    too_few = "align_start", cols_remove = TRUE
  ) %>%
  pivot_longer(
    cols = c(Marker_1, Marker_2, Marker_3, Marker_4),
    names_to = "marker_no",
    values_to = "marker",
    values_drop_na = TRUE
  ) %>%
  select(-marker_no) %>%
  mutate(
    drug_class = "Quinolones",
    drug = NA_character_
  ) ## the variable drug appears essential for get_binary_matrix function to work...

# Check the format of the processed genotype table
head(salm_geno)
#> # A tibble: 6 × 4
#>   Name   marker     drug_class drug
#>   <chr>  <chr>      <chr>      <chr>
```

```
#> 1 SAL001 gyrA_S83F Quinolones <NA>
#> 2 SAL001 parC_T57S Quinolones <NA>
#> 3 SAL002 gyrA_S83Y Quinolones <NA>
#> 4 SAL003 gyrA_S83F Quinolones <NA>
#> 5 SAL003 parC_T57S Quinolones <NA>
#> 6 SAL003 qnrB19    Quinolones <NA>
```

#### Summarise genotype data

The `summarise_geno()` function can provide various summaries of the genotype table, including the total number of unique markers and a table showing each marker's prevalence in the dataset.

```
# Create geno_summary object
salm_geno_summary <- summarise_geno(salm_geno)

# Total number of markers, drugs, and drug classes
salm_geno_summary$uniques
#> # A tibble: 3 × 2
#>   column     n_unique
#>   <chr>         <int>
#> 1 marker           26
#> 2 drug              1
#> 3 drug_class        1

# Prevalence of each detected marker, in decreasing abundance
salm_geno_summary$markers %>% arrange(-n)
#> # A tibble: 26 × 4
#>    marker    drug  drug_class     n
#>    <chr>     <chr> <chr>      <int>
#>  1 parC_T57S <NA>  Quinolones    49
#>  2 gyrA_S83F <NA>  Quinolones    18
#>  3 gyrA_S83Y <NA>  Quinolones    18
#>  4 qnrS1     <NA>  Quinolones    18
#>  5 qnrB19    <NA>  Quinolones    16
#>  6 gyrA_D87Y <NA>  Quinolones    14
#>  7 gyrA_D87N <NA>  Quinolones    12
#>  8 gyrA_D87G <NA>  Quinolones    11
#>  9 parC_S80I <NA>  Quinolones     9
#> 10 qnrA1     <NA>  Quinolones     5
#> # ℹ 16 more rows
```

These summaries show that there are 26 distinct genotypic markers for quinolone resistance in this dataset. The most common one is parC_T57S, found in 49 of 115 isolates, followed by gyrA_S83F, gyrA_S83Y, and qnrS1, each found in 18 of 115 isolates. Eleven markers are found only once across all isolates.

### Phenotype table

#### Format raw phenotype data for AMRgen

To use downstream `AMRgen` functions, a phenotype table must be in long form with a single column for all antimicrobial agents that were tested, rather than separate columns for each agent. It also needs to have at a minimum the following columns: `id` (unique sample name for each isolate), `spp_pheno` (species name formatted as per the AMR package `mo` class), `drug` (antimicrobial agent name formatted as per the AMR package `ab` class), a S/I/R phenotype column (e.g. one or more of `pheno_eucast`, `pheno_clsi`, `pheno_provided`, `ecoff`). If raw assay data is to be included, it needs to be in a column called `mic` and/or `disk`.

To convert out dataset to this format, we begin by extracting the columns with the unique isolate names (`Sample`) and those with the MIC results for the three antimicrobial agents that we tested (`Ciprofloxacin`, `Levofloxacin`, `Moxifloxacin`). We also keep the columns with additional metadata (`Source`, `Serovar`), as these are not in the genotype table. The antimicrobial columns need to be forced to character vectors to avoid issues caused by occasional presence of non-numeric prefixes (`>` or `<`). We then pivot the table to long form.

```
# Pheno table: select columns with sample ID, metadata, and antimicrobials tested,
# then pivot to long form
salm_pheno <- salm_raw %>%
  select(Sample, Source, Serovar, Ciprofloxacin, Levofloxacin, Moxifloxacin) %>%
  mutate(across(c(Ciprofloxacin, Levofloxacin, Moxifloxacin), as.character)) %>%
  pivot_longer(
    cols = c(Ciprofloxacin, Levofloxacin, Moxifloxacin),
    names_to = "drug",
    values_to = "MIC.values",
    values_drop_na = TRUE
  )
```

Our phenotype data is not yet in a standard AMR/AMRgen format so we use the helpful `format_pheno` function to add the species name, to format the antimicrobial and MIC columns correctly, and to generate the S/I/R phenotype column `pheno_eucast` by interpreting our Salmonella MIC data against EUCAST breakpoints. We apply these breakpoints across our human/animal/other isolates, as our interest is in resistance phenotypes that are potentially problematic in human infections, including zoonotic ones.

```
salm_ast <- format_pheno(
  input = salm_pheno,
  sample_col = "Sample",
  species = "Salmonella enterica",
  ab_col = "drug",
  mic_col = "MIC.values",
  interpret_eucast = TRUE
)

# Check the format of the processed phenotype table
head(salm_ast)
#> # A tibble: 6 × 7
#>   id     drug                 mic pheno_eucast spp_pheno          Source Serovar
#>   <chr>  <ab>               <mic> <sir>        <mo>               <chr>  <chr>
#> 1 SAL001 CIP: Ciprofloxacin  0.19 R            B_SLMNL_ENTR: Sal… Animal other
#> 2 SAL001 LVX: Levofloxacin   0.50 S            B_SLMNL_ENTR: Sal… Animal other
#> 3 SAL001 MFX: Moxifloxacin   0.50 R            B_SLMNL_ENTR: Sal… Animal other
#> 4 SAL002 CIP: Ciprofloxacin  0.38 R            B_SLMNL_ENTR: Sal… Human  S. Ent…
#> 5 SAL002 LVX: Levofloxacin   1.50 R            B_SLMNL_ENTR: Sal… Human  S. Ent…
#> 6 SAL002 MFX: Moxifloxacin   1.50 R            B_SLMNL_ENTR: Sal… Human  S. Ent…
```

#### Summarise phenotype data

The summarise_pheno function can be used to generate various summaries of the phenotype table, including the total number of samples, drugs, and species, and table of S/I/R counts for each drug in the dataset. Our dataset is limited to a single species with data from only one method (E-test MIC), and we are looking only at the EUCAST breakpoints, but the vignette “Analysing Geno-Pheno Data” shows how the summarise_pheno function can be used for datasets with multiple species, methods, drug classes, and interpretation guidelines.

```
salm_pheno_summary <- summarise_pheno(salm_ast, pheno_cols = c("pheno_eucast"))
#> No disk data colummn provided

# Number of samples, drugs, species, and methods included in phenotype table
salm_pheno_summary$uniques
#> # A tibble: 3 × 2
#>   column    n_unique
#>   <chr>        <int>
#> 1 id             115
#> 2 drug             3
#> 3 spp_pheno        1

# SIR summary table for each drug in the phenotype table
salm_pheno_summary$pheno_counts_list
#> $pheno_eucast
#> # A tibble: 3 × 6
#>   drug               drug_name     spp_pheno                S     R     I
#>   <ab>               <chr>         <chr>                <int> <int> <int>
#> 1 CIP: Ciprofloxacin Ciprofloxacin Salmonella enterica    23    92    NA
#> 2 LVX: Levofloxacin  Levofloxacin  Salmonella enterica    63    25    27
#> 3 MFX: Moxifloxacin  Moxifloxacin  Salmonella enterica    20    95    NA
```

These summaries show that there are 115 isolates of one species in this dataset, with results for three drugs. Similar proportions were resistant to ciprofloxacin and moxifloxacin, whereas a lower proportion was resistant to levofloxacin (using the EUCAST breakpoints).

Now that we have formatted our data into a genotype table (salm_geno) and a phenotype table (salm_ast) that are compatible with AMRgen, we can use its downstream functions for plotting distributions, combining geno/pheno data, modeling binary drug phenotype, assessing positive predictive values of genetic markers, and comparing how combinations of markers influence resistance phenotype. Other vignettes outline these workflows in more detail, so here we show only a basic workflow, focusing on how additional metadata can be included.

### Plot phenotype data distributions

#### Overall phenotype distributions

We begin by looking at the distribution of the MIC values in our dataset for the three antimicrobials that we tested. If desired, these plots could then be combined into a multipanel figure using packages like patchwork or ggpubr.

```
# Plot MIC distributions coloured by S/I/R call
assay_by_var(pheno_table = salm_ast, pheno_drug = "Ciprofloxacin", measure = "mic",
        colour_by = "pheno_eucast")
```

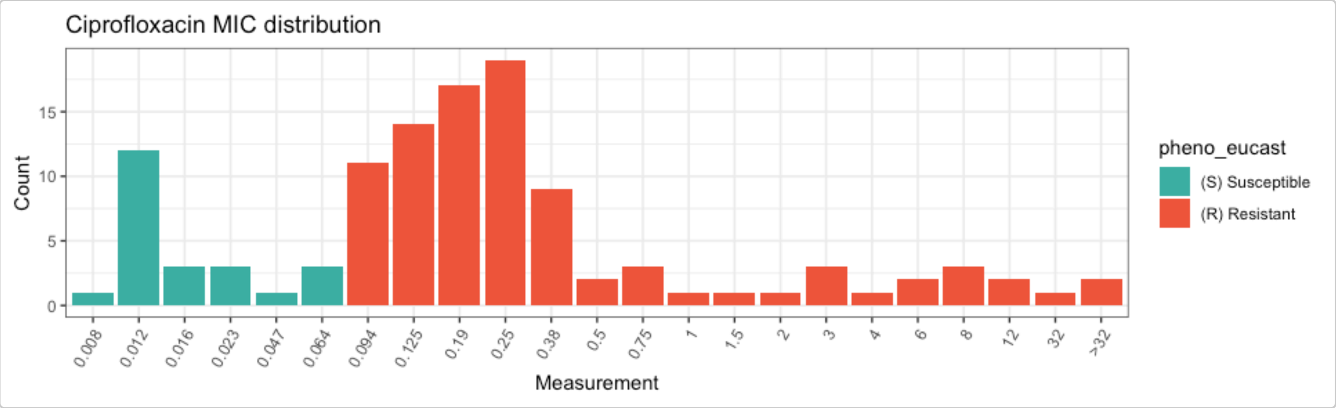


```
assay_by_var(pheno_table = salm_ast, pheno_drug = "Levofloxacin", measure = "mic",
        colour_by = "pheno_eucast")
```

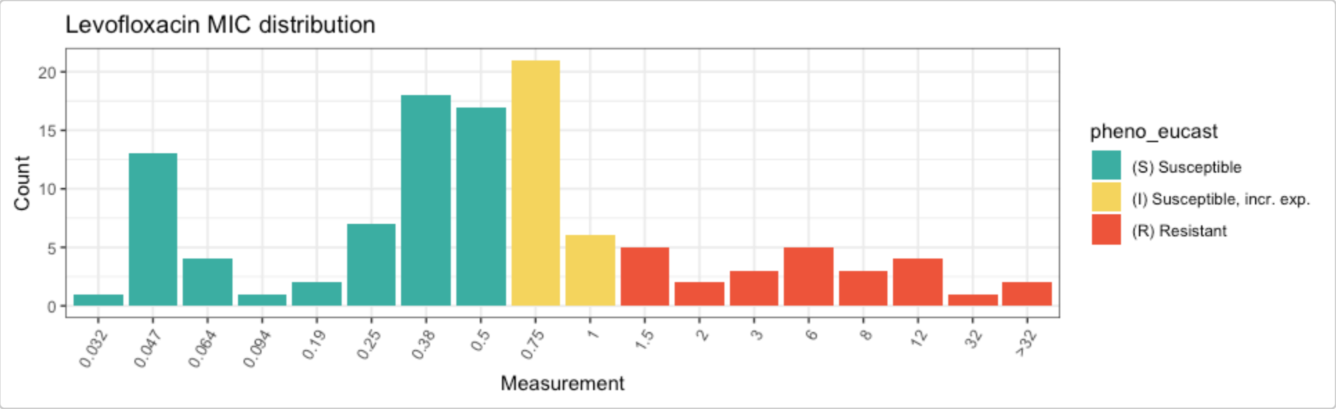


```
assay_by_var(pheno_table = salm_ast, pheno_drug = "Moxifloxacin", measure = "mic",
        colour_by = "pheno_eucast")
```

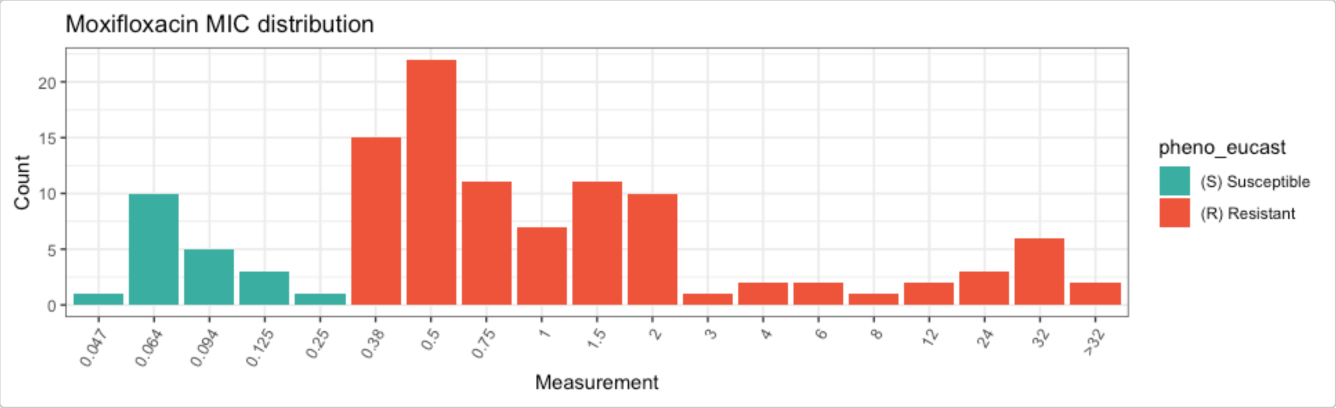


While these plots show the MIC distributions of all isolates in our dataset, we can break this down further by isolation source (human, animal, other) and by *S. enterica* serovar. This can be done in a few different ways, notably by faceting and/or by modifying the colour variable.

#### Phenotype distributions by categorical variable

The `assay_by_var` function can separate plots by an additional categorical variable by specifying `facet_by` within the function. For example, we can split the ciprofloxacin plot by isolation source like this:

```r
assay_by_var(pheno_table = salm_ast, pheno_drug = "Ciprofloxacin", measure = "mic",
        colour_by = "pheno_eucast", facet_by = "Source")
```

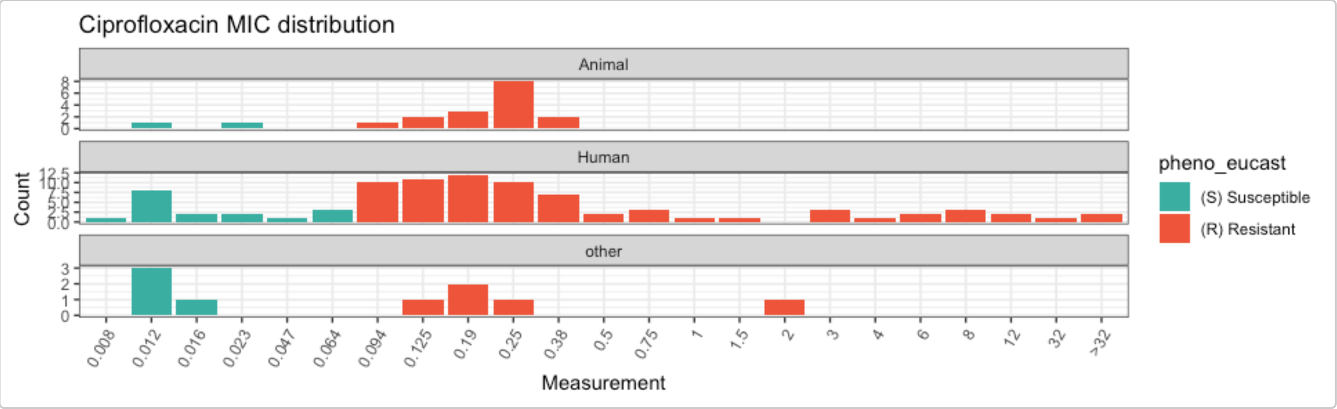


Alternatively, because `assay_by_var` is a `ggplot2` function, it can be extended by adding `ggplot2` layers, including `facet_wrap()` for a single faceting variable (to allow more control over how the facets are displayed) or `facet_grid()` to facet on two variables.

```r
assay_by_var(pheno_table = salm_ast, pheno_drug = "Ciprofloxacin", measure = "mic",
        colour_by = "pheno_eucast") +
  facet_grid(Source ~ Serovar)
```

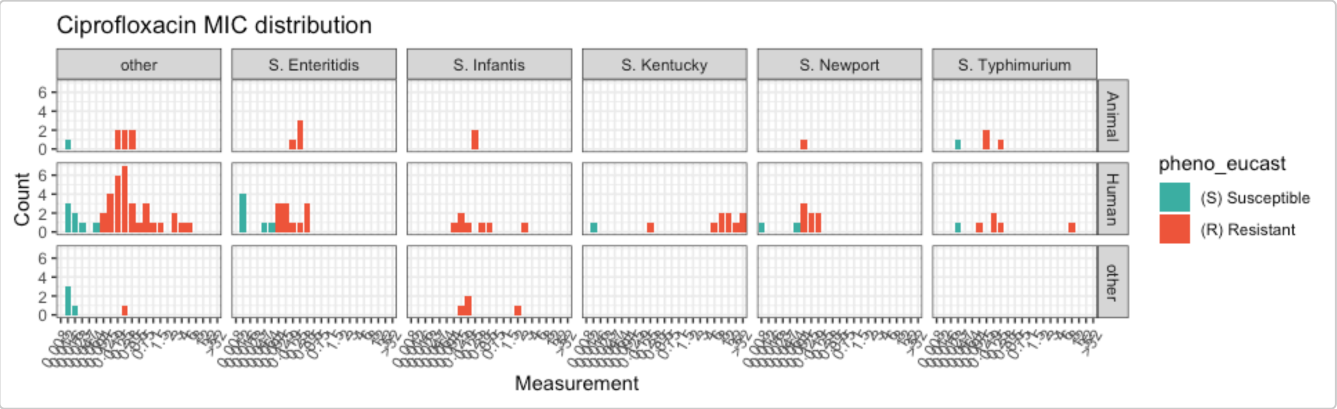


Another option for displaying categorical metadata is to change the colour variable `colour_by` but still show the breakpoints as vertical lines using the `guideline` and `species` calls within `assay_by_var`. If desired, you can also modify other aspects of the plot using standard ggplot2 extensions (e.g. using `viridis` to change the colour palette).

```r
# Specify species and guideline to show breakpoints, but colour bars by isolation source
assay_by_var(pheno_table = salm_ast, pheno_drug = "Ciprofloxacin", measure = "mic",
        colour_by = "Source", species = "Salmonella enterica", guideline = "EUCAST 2025")
        +
  scale_fill_viridis_d(end = 0.8)
#>   MIC breakpoints determined using AMR package: S <= 0.064 and R > 0.064
```

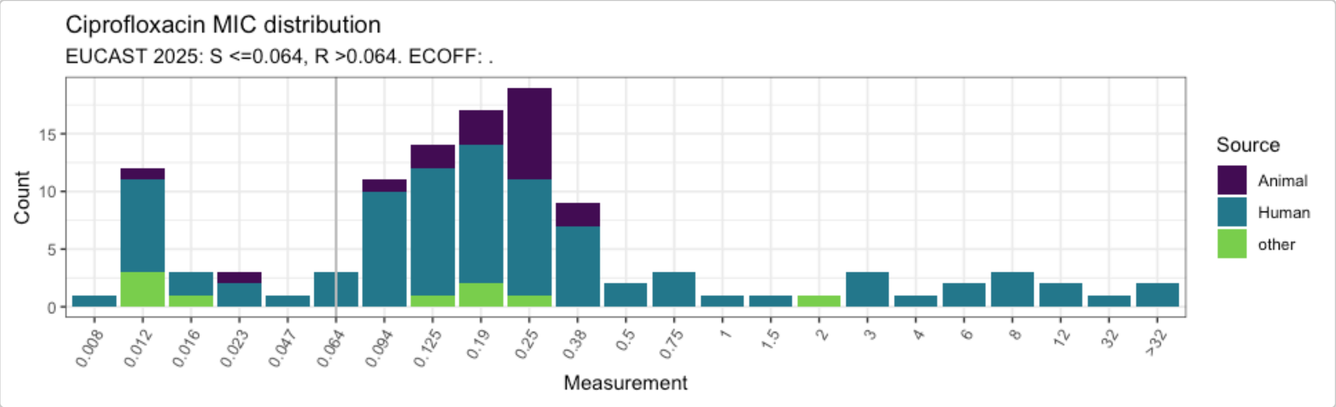


#### Combine ciprofloxacin genotype and phenotype data

Analysis of combined genotype-phenotype data must be carried out separately for each antimicrobial agent. The first step is to generate a combined dataframe for the specified agent from the genotype and phenotype tables, using the `get_binary_matrix()` function. As an example, we will do this for ciprofloxacin with our small *S. enterica* dataset, but this could also be done for levofloxacin and moxifloxacin.

```
### get_binary_matrix throws an error unless the variable drug_class is in salm_geno, even
        if it's empty
cip_bin <- get_binary_matrix(
  salm_geno,
  salm_ast,
  pheno_drug = "Ciprofloxacin",
  geno_class = "Quinolones",
  sir_col = "pheno_eucast",
  keep_assay_values = TRUE,
  keep_assay_values_from = "mic"
)
#>  Defining NWT in binary matrix as I/R vs S, as no ECOFF column defined

# check format
head(cip_bin)
#> # A tibble: 6 × 31
#>    id     pheno   mic     R   NWT gyrA_S83F parC_T57S gyrA_S83Y qnrB19 gyrA_D87G
#>    <chr>  <sir> <mic> <dbl> <dbl>     <dbl>     <dbl>     <dbl>  <dbl>     <dbl>
#> 1 SAL001 R     0.190     1     1         1         1         0      0         0
#> 2 SAL002 R     0.380     1     1         0         0         1      0         0
#> 3 SAL003 R     3.000     1     1         1         1         0      1         0
#> 4 SAL004 R     0.125     1     1         0         0         0      0         1
#> 5 SAL005 R     0.094     1     1         0         0         0      0         0
#> 6 SAL006 R     8.000     1     1         1         0         0      0         0
#> # ℹ 21 more variables: gyrA_D87Y <dbl>, gyrA_D87N <dbl>, qnrS1 <dbl>,
#> #   qnrA1 <dbl>, gyrA_D87X <dbl>, qepA1 <dbl>, parC_S80I <dbl>, gyrA_u <dbl>,
#> #   gyrA_D82N <dbl>, qnrB2 <dbl>, qnrD <dbl>, gyrB_E446D <dbl>, qnrDv1 <dbl>,
#> #   parE_S474G <dbl>, gyrA_Y100H <dbl>, parC_S80X <dbl>, parC_F60L <dbl>,
#> #   parE_D475Y <dbl>, qnrS4 <dbl>, gyrB_L446M <dbl>, parC_A81X <dbl>

# list colnames (alphabetically) to see full list of quinolone markers in data
# (will include additional columns id, pheno, mic, NWT, R, Source, and Serovar)
sort(colnames(cip_bin))
#>  [1] "gyrA_D82N"   "gyrA_D87G"   "gyrA_D87N"   "gyrA_D87X"   "gyrA_D87Y"
```

```
#>  [6] "gyrA_S83F"   "gyrA_S83Y"   "gyrA_u"      "gyrA_Y100H" "gyrB_E446D"
#> [11] "gyrB_L446M" "id"          "mic"         "NWT"         "parC_A81X"
#> [16] "parC_F60L"   "parC_S80I"   "parC_S80X"   "parC_T57S"   "parE_D475Y"
#> [21] "parE_S474G" "pheno"       "qepA1"       "qnrA1"       "qnrB19"
#> [26] "qnrB2"       "qnrD"        "qnrDv1"      "qnrS1"       "qnrS4"
#> [31] "R"
```

This binary matrix can be used for numerous downstream analyses, as described in other vignettes. Here, we show how some can be modified with additional metadata variables.

The `get_binary_matrix()` output (`cip_bin`) did not retain our additional metadata variables `Source` and `Serovar`. To use these variables in downstream analyses, we first extract them from `salm_pheno` and then join to `cip_bin` to create `cip_bin_meta`, which can be used in plotting functions (warning: statistical model functions may not work correctly with these additional metadata so `cip_bin` should be used in those instead of `cip_bin_meta`).

```
salm_metadata <- salm_pheno %>%
  select(Sample, Source, Serovar) %>%
  rename(id = Sample)

cip_bin_meta <- left_join(cip_bin, salm_metadata)
#> Joining with `by = join_by(id)`
```

#### Plot ciprofloxacin phenotype by number of mutations and Serovar/Source

Joining the metadata to `cip_bin` allows us to colour the MIC distribution plot by the number of mutations in *gyrA* and facet by `Serovar`, highlighting that all S. Infantis isolates had one *gyrA* mutation and all S. Kentucky isolates had two *gyrA* mutations, whereas other serovars had variable numbers of mutations.

```
# count the number of gyrA mutations per genome
gyrA_mut <- cip_bin_meta %>%
  dplyr::mutate(gyrA_mut = rowSums(across(contains("gyrA_") & where(is.numeric)), na.rm =
        T)) %>%
  select(mic, gyrA_mut, Source, Serovar)

# plot the MIC distribution, coloured by count of gyrA mutations
mic_by_gyrA_count <- assay_by_var(gyrA_mut, measure = "mic", colour_by = "gyrA_mut",
        colour_legend_label = "Number of\ngyrA mutations", measure_axis_label = "MIC
        (mg/L)", pheno_drug = "Ciprofloxacin", colours = viridisLite::viridis(5)[c(4, 3,
        2)]) + facet_wrap(~Serovar)

# add title with italicised species and drug names
mic_by_gyrA_count + ggtitle(expression(paste(
  "Ciprofloxacin MIC in ",
  italic("Salmonella"),
  " serovars, by number of ",
  italic("gyrA"),
  " mutations"
)))
```

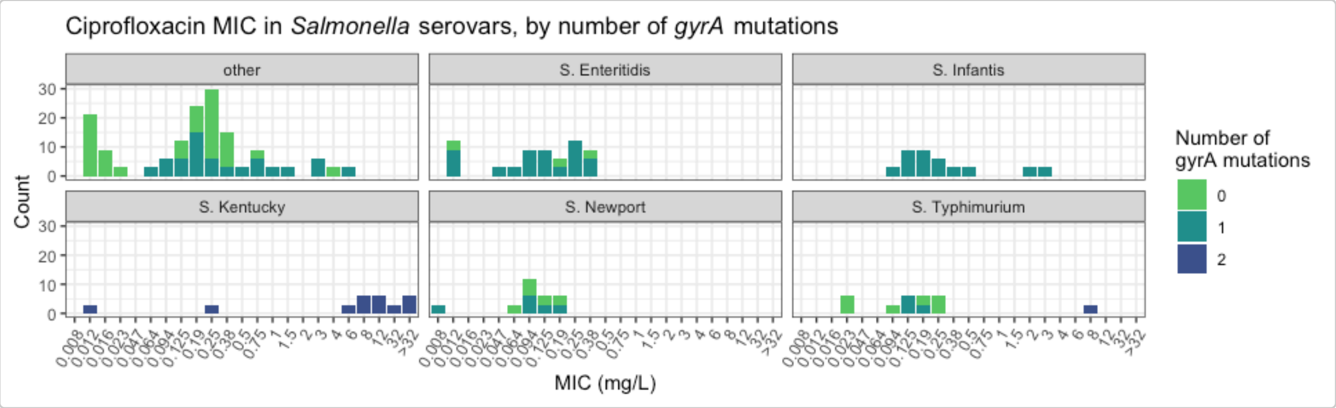


Similarly, we can plot the total number of markers per isolate and facet by `Source`.

```
# count the number of genetic determinants per genome
marker_count <- cip_bin_meta %>%
  mutate(marker_count = rowSums(across(where(is.numeric) & !any_of(c("R", "NWT"))), na.rm
      = T)) %>%
  select(mic, marker_count, Source, Serovar)

# plot the MIC distribution, coloured by count of associated genetic markers
mic_by_marker_count <- assay_by_var(marker_count, measure = "mic", colour_by =
        "marker_count", colour_legend_label = "Total number\nof markers", pheno_drug =
        "Ciprofloxacin", colours = viridisLite::viridis(max(marker_count$marker_count) +
        1)) +
  facet_wrap(~Source, ncol = 1)

mic_by_marker_count
```

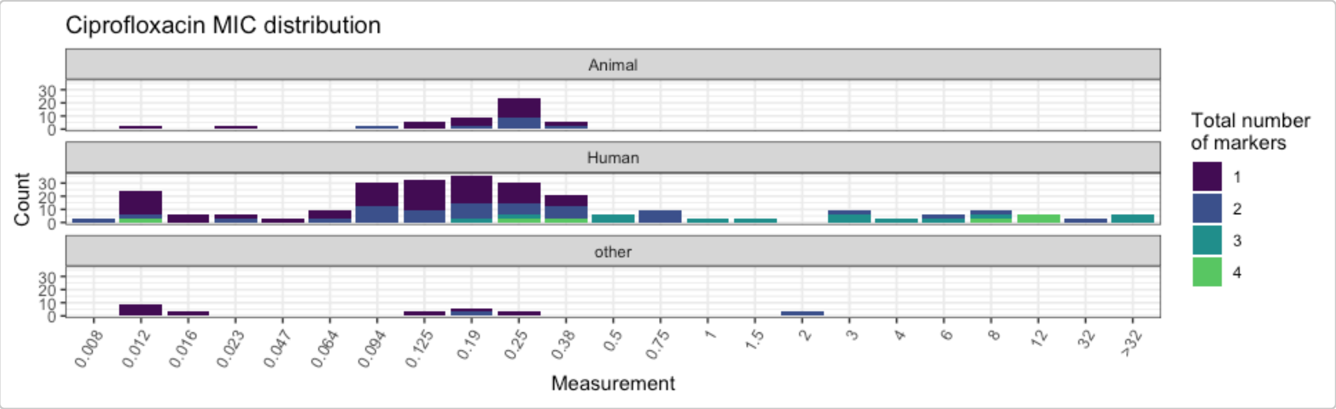


We can also use the `boxplot=TRUE` option in `assay_by_var()` to see the MIC distribution as boxplots, stratified by number of markers. This also summarises the median and interquartile range of MIC values, per marker count so we can quantify as well as visualise the impact of number of mutations on MIC.

```
# plot the MIC distributions as boxplots, stratified by number of markers
mic_boxplot_by_marker_count <- assay_by_var(marker_count, measure = "mic", colour_by =
        "marker_count", colour_legend_label = "Total number\nof markers", pheno_drug =
        "Ciprofloxacin", colours = viridisLite::viridis(max(marker_count$marker_count) +
        1), boxplot = T)

mic_boxplot_by_marker_count$plot
```

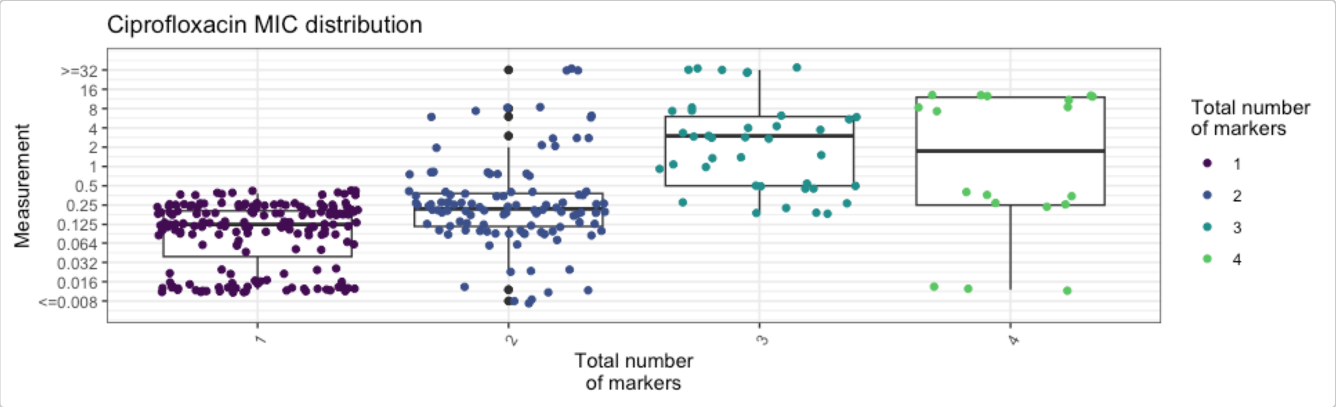


```
mic_boxplot_by_marker_count$stats
#> # A tibble: 4 × 6
#>   marker_count     n median geom_mean   q25   q75
#>          <dbl> <int>  <dbl>     <dbl> <dbl> <dbl>
#> 1            1   180  0.125    0.0884 0.041 0.205
#> 2            2   108  0.22     0.266  0.117 0.38
#> 3            3    39  3        2.22   0.5   6
#> 4            4    18  4.19     1.05   0.25 12
```

We can also use the `boxplot=TRUE` option in `assay_by_var()` to see the MIC distribution as boxplots, stratified by number of markers. This also summarises the median and interquartile range of MIC values, per marker count so we can quantify as well as visualise the impact of number of mutations on MIC.

```
# plot the MIC distributions as boxplots, stratified by number of markers
mic_boxplot_by_marker_count_source <- assay_by_var(marker_count, measure = "mic",
        colour_by = "marker_count", colour_legend_label = "Total number\nof markers",
        pheno_drug = "Ciprofloxacin", colours =
        viridisLite::viridis(max(marker_count$marker_count) + 1), facet_by = "Source",
        boxplot = T)

mic_boxplot_by_marker_count_source$plot
```

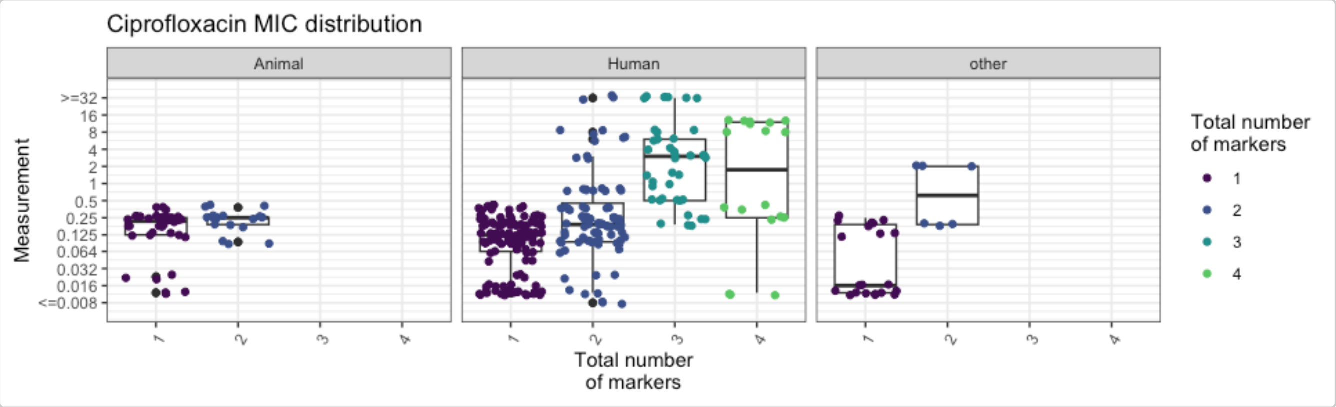


#### Plot ciprofloxacin phenotype by combinations of markers

We can look at how combinations of markers are associated with phenotypic ciprofloxacin MIC by generating an UpSet plot with the `amr_upset` function. This function does not accept extra metadata columns so we use `cip_bin` instead of `cip_bin_meta` here. As our dataset is quite small, we keep all the combinations, including those with a single isolate (`min_set_size = 1`).

```r
# Compare ciprofloxacin MIC data with quinolone marker combinations,
#    using the binary matrix we constructed earlier via get_binary_matrix()
cipro_mic_upset <- amr_upset(
  cip_bin,
  min_set_size = 1,
  assay = "mic",
  order = "value"
)
#> Ordering markers by frequency
#> Scale for y is already present.
#> Adding another scale for y, which will replace the existing scale.
#> Scale for y is already present.
#> Adding another scale for y, which will replace the existing scale.
```

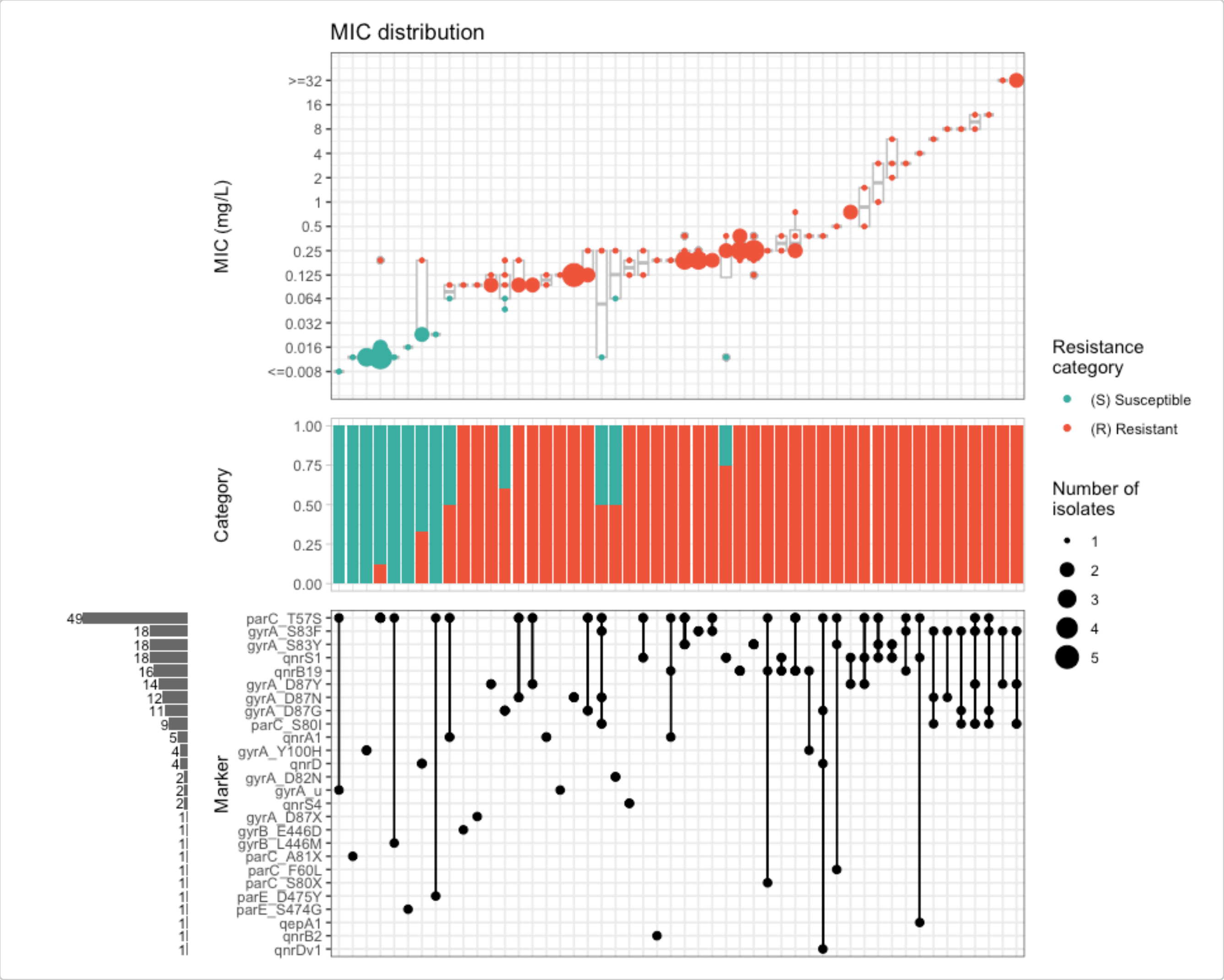


We can generate UpSet plots of a subset of isolates by filtering on one of our additional metadata variables and then running `amr_upset()`. For example, we can focus on the animal isolates only:

```r
cip_bin_animal <- cip_bin_meta %>%
  filter(Source == "Animal") %>%
  select(-Source, -Serovar)

cipro_mic_upset_animal <- amr_upset(
  cip_bin_animal,
  min_set_size = 1,
  assay = "mic",
```

```
  order = "value"
)
#> Ordering markers by frequency
#> Scale for y is already present.
#> Adding another scale for y, which will replace the existing scale.
#> Scale for y is already present.
#> Adding another scale for y, which will replace the existing scale.
```

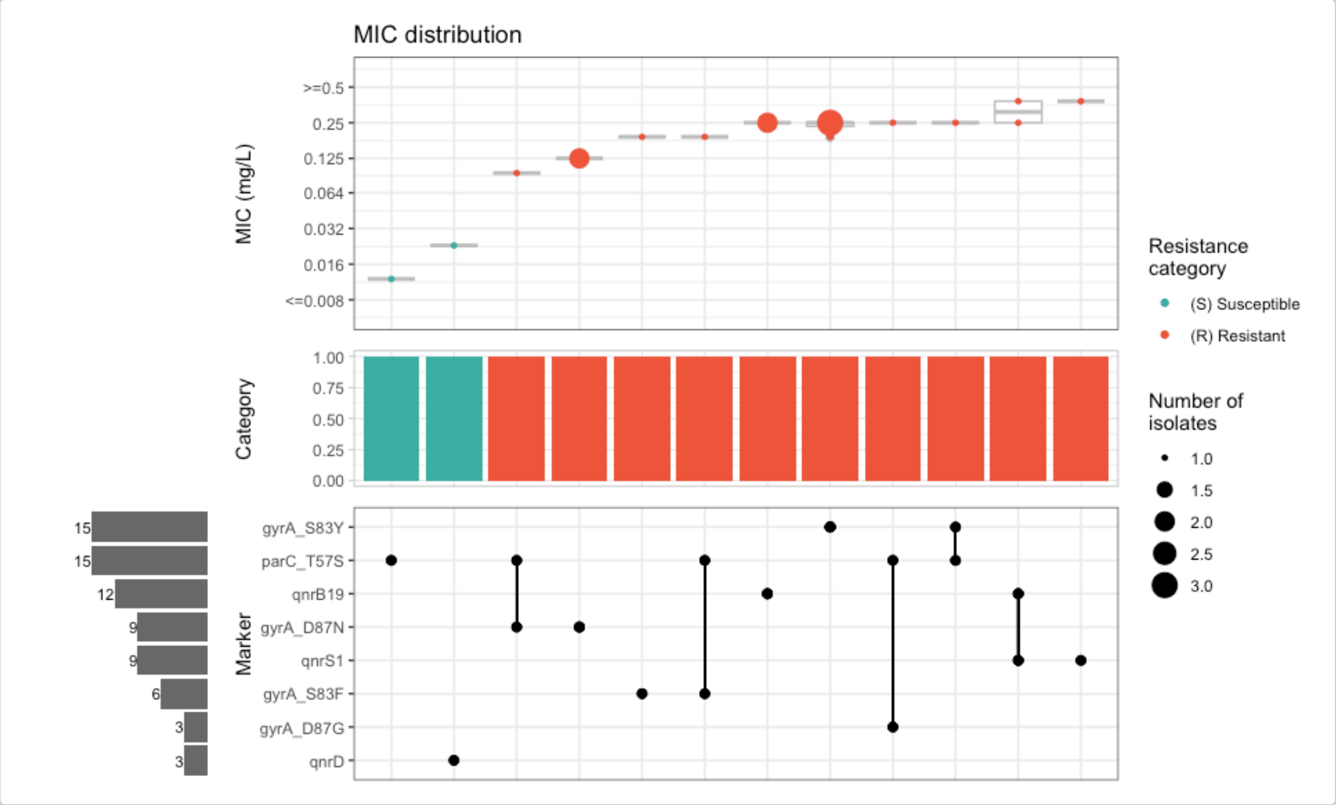

### Example with custom classification of genotype hits

**Natacha Couto**

**2026-03-12**

### Analysing clindamycin resistance in *Staphylococcus aureus*

#### Introduction

This is document summarises the analysis of clindamycin resistance in *S. aureus*. More specifically, we will look at the distribution of clindamycin resistance in *S. aureus* isolates from a dataset of clinical samples.

Start by loading the package:

```
library(AMRgen)
library(ggplot2)
library(dplyr)
```

#### Data preparation

For this example, we have collated genotype-phenotype data from NCBI and EBI through the ESGEM-AMR Staphylococcus subgroup. Phenotypic data were collated into a single table (one row per isolate). The genotype data was generated using AMRFinderPlus. The pre-loaded objects `pheno_CLI_public` and `afp_CLI_public` serve as the **input** for `AMRgen`. First, we will create a new `variant.label` column that includes the marker gene name and its closest accession number, and also indicate whether the match to the accession is exact or not.

```
# Create the new marker.label column
afp_CLI_public <- afp_CLI_public %>%
  mutate(exact_match = if_else(
    `% Coverage of reference sequence` == 100 &
      `% Identity to reference sequence` == 100.00,
    "", "x_"
  )) %>%
  mutate(node_hit = paste0(node, "__", exact_match, `Accession of closest sequence`)) %>%
  mutate(variant.label = case_when(
    `variation type` == "Inactivating mutation detected" ~ paste0(node_hit, ":-"),
    !is.na(mutation) ~ paste0(node_hit, ":", mutation),
    TRUE ~ node_hit
  ))
```

#### Genotype-phenotype analysis

Now let's create some upset plots, to see how genes, and their allelic variants, relate to clindamycin MICs.

```
# Visualise with UpSet plot (markers)
cli_mic_upset <- amr_upset(
  geno_table = afp_CLI_public,
  pheno_table = pheno_CLI_public,
  marker_col = "marker.label",
  pheno_drug = "Clindamycin",
  min_set_size = 2,
  assay = "mic",
  order = "value",
  print_set_size = TRUE,
  plot_set_size = TRUE,
  print_category_counts = TRUE,
  bp_S = 0.25,
  bp_R = 0.5
)
```

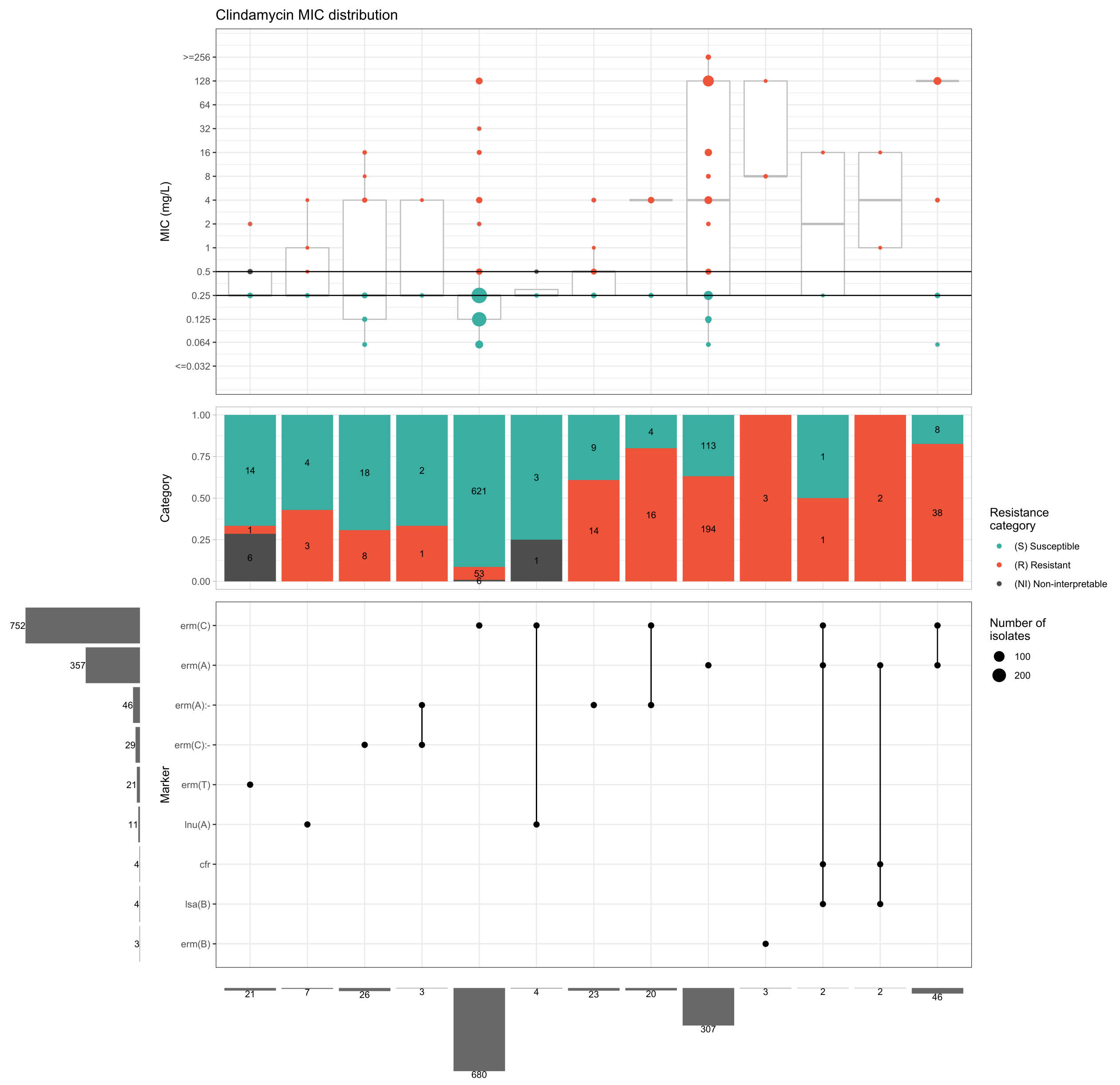


```r
# Visualise with UpSet plot (markers)
# order markers alphabetically so we can more easily see where there are different
        variants of the same gene
cli_mic_upset_variant <- amr_upset(
  geno_table = afp_CLI_public,
  pheno_table = pheno_CLI_public,
  marker_col = "variant.label",
  pheno_drug = "Clindamycin",
  min_set_size = 2,
  assay = "mic",
  order = "value",
  marker_order = "alpha", # order markers alphabetically
  print_set_size = TRUE,
  plot_set_size = TRUE,
  print_category_counts = TRUE,
  bp_S = 0.25,
  bp_R = 0.5
)
```

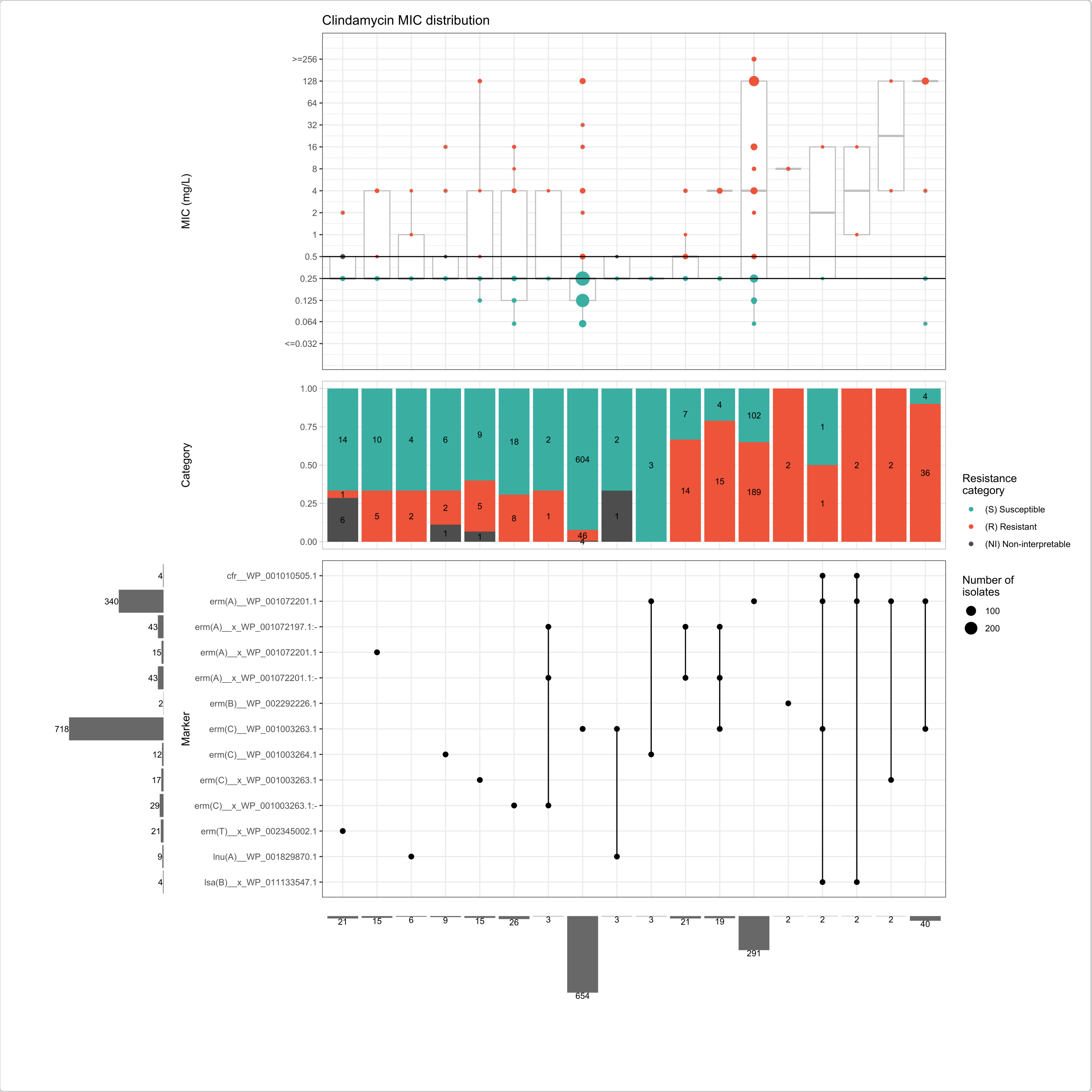


#### Solo PPV analysis

Next, we will calculate the solo positive predictive value (soloPPV) for each marker and variant. This will allow us to see which markers and variants are most predictive of clindamycin resistance.

```
# soloPPV analysis for markers
# order markers alphabetically so we can more easily see where there are different
        variants of the same gene
cli_soloPPV <- solo_ppv(
  geno_table = afp_CLI_public,
  pheno_table = pheno_CLI_public,
  marker_col = "marker.label",
  pheno_drug = "Clindamycin",
  order_ppv = FALSE
)
```

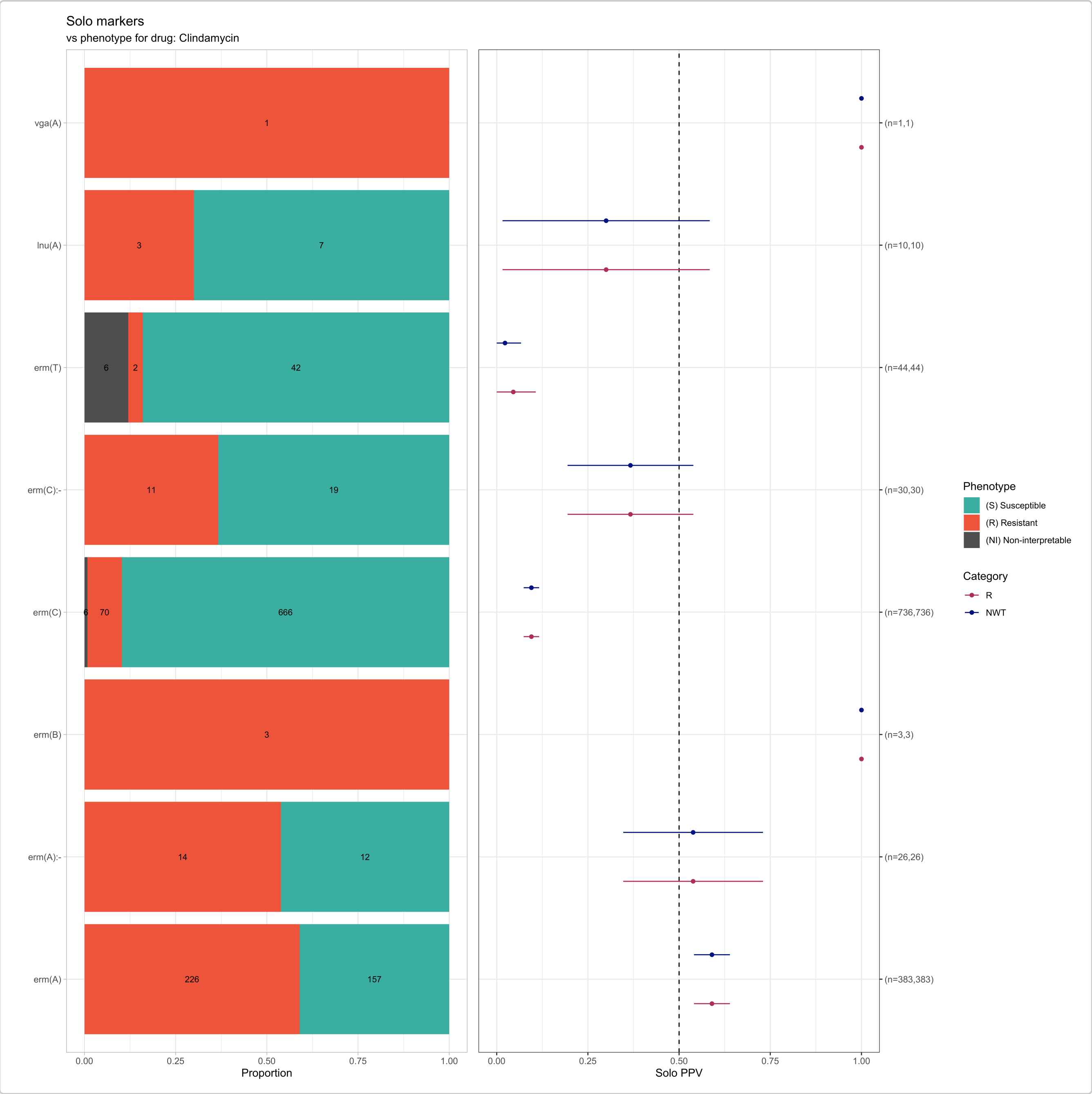


```r
# soloPPV analysis for variants
# order markers alphabetically so we can more easily see where there are different
        variants of the same gene
cli_soloPPV_variant <- solo_ppv(
  geno_table = afp_CLI_public,
  pheno_table = pheno_CLI_public,
  marker_col = "variant.label",
  pheno_drug = "Clindamycin",
  order_ppv = FALSE
)
```

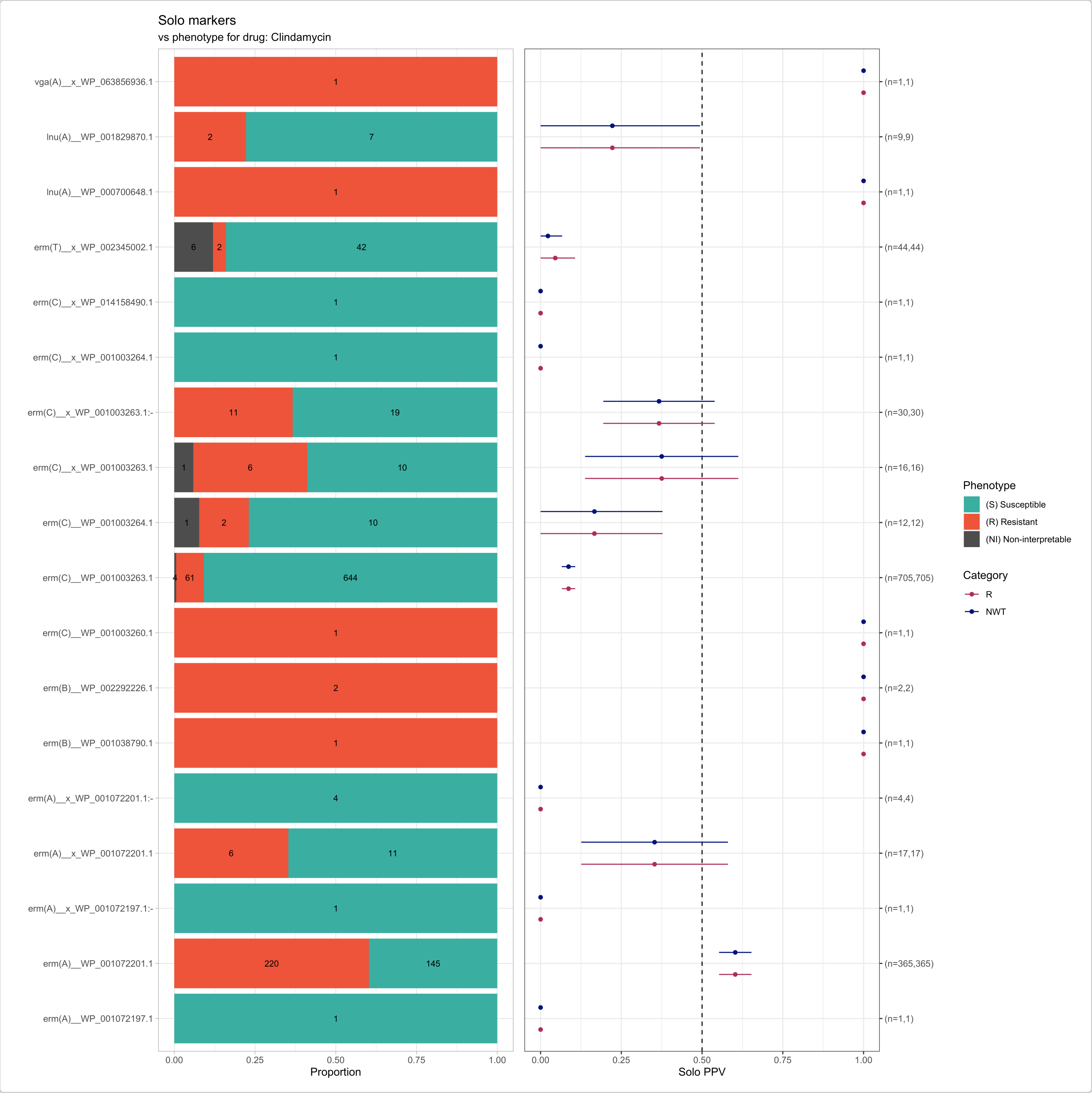


```
cli_soloPPV_variant$solo_stats
## # A tibble: 38 × 8
##    marker                    category     x     n    ppv     se ci.lower ci.upper
##    <chr>                     <chr>    <dbl> <int>  <dbl>  <dbl>    <dbl>    <dbl>
##  1 erm(A)__WP_001072197.1    R            0     1 0      0        0         0
##  2 erm(A)__WP_001072201.1    R          220   365 0.603  0.0256   0.553     0.653
##  3 erm(A)__x_WP_001072197.… R            0     1 0      0        0         0
##  4 erm(A)__x_WP_001072201.1 R            6    17 0.353  0.116    0.126     0.580
##  5 erm(A)__x_WP_001072201.… R            0     4 0      0        0         0
##  6 erm(B)__WP_001038790.1    R            1     1 1      0        1         1
##  7 erm(B)__WP_002292226.1    R            2     2 1      0        1         1
##  8 erm(C)__WP_001003260.1    R            1     1 1      0        1         1
##  9 erm(C)__WP_001003263.1    R           61   705 0.0865 0.0106   0.0658    0.107
## 10 erm(C)__WP_001003264.1    R            2    12 0.167  0.108    0         0.378
## # ℹ 28 more rows
```

We can see from this analysis that the variants of the markers have different associations with clindamycin resistance. For example, the isolates with exact matches to erm(C)_WP001003263.1 or erm(C)_WP001003264.1 are associated with low PPV for resistance (8.5%, n=60/705 and 17%, n=2/12, respectively). This highlights the importance of considering the specific variants of the markers when predicting clindamycin resistance. Note: this discrepancy may be caused by inducible resistance, which is not always captured by a standard AST test. This is because the resistance gene may not be expressed under the conditions of the AST test, but can be induced in the presence of certain antibiotics (e.g., erythromycin). This is a known phenomenon for clindamycin resistance in *S. aureus*, where the presence of an erm gene can lead to inducible resistance that may not be detected in a standard AST test.

#### MIC distribution associated with allelic variants

We can use the `assay_by_var()` function to visualise the MIC distributions associated with different variants of the same gene. First we need to join the key genotype fields into the phenotype table, and then use `assay_by_var()` to plot MIC assay measures grouped by variant and faceted into one panel per gene, coloured by S/I/R phenotype.

```
cli_geno_pheno <- afp_CLI_public %>%
  filter(`variation type` == "Gene presence detected") %>%
  mutate(variant_hit = if_else(exact_match == "",
    `Accession of closest sequence`,
    paste0(`Accession of closest sequence`, "_x")
  )) %>%
  select(id, marker.label, variant_hit) %>%
  left_join(pheno_CLI_public)
## Joining with `by = join_by(id)`
## Warning in left_join(., pheno_CLI_public): Detected an unexpected many-to-many
        relationship between `x` and `y`.
## ℹ Row 7609 of `x` matches multiple rows in `y`.
## ℹ Row 3161 of `y` matches multiple rows in `x`.
## ℹ If a many-to-many relationship is expected, set `relationship =
##   "many-to-many"` to silence this warning.
```

```
cli_mic_byhit <- assay_by_var(cli_geno_pheno,
  group_by = "variant_hit",
  boxplot = T, colour_by = "pheno_eucast"
)$plot +
  facet_wrap(~marker.label, scales = "free_y", ncol = 2) +
  coord_flip() +
  theme(legend.position = "none")

cli_mic_byhit + ggtitle(expression(paste(
  "Clindamycin MIC in ",
  italic("S. aureus"),
  ", by gene symbol and closest reference"
)))
```

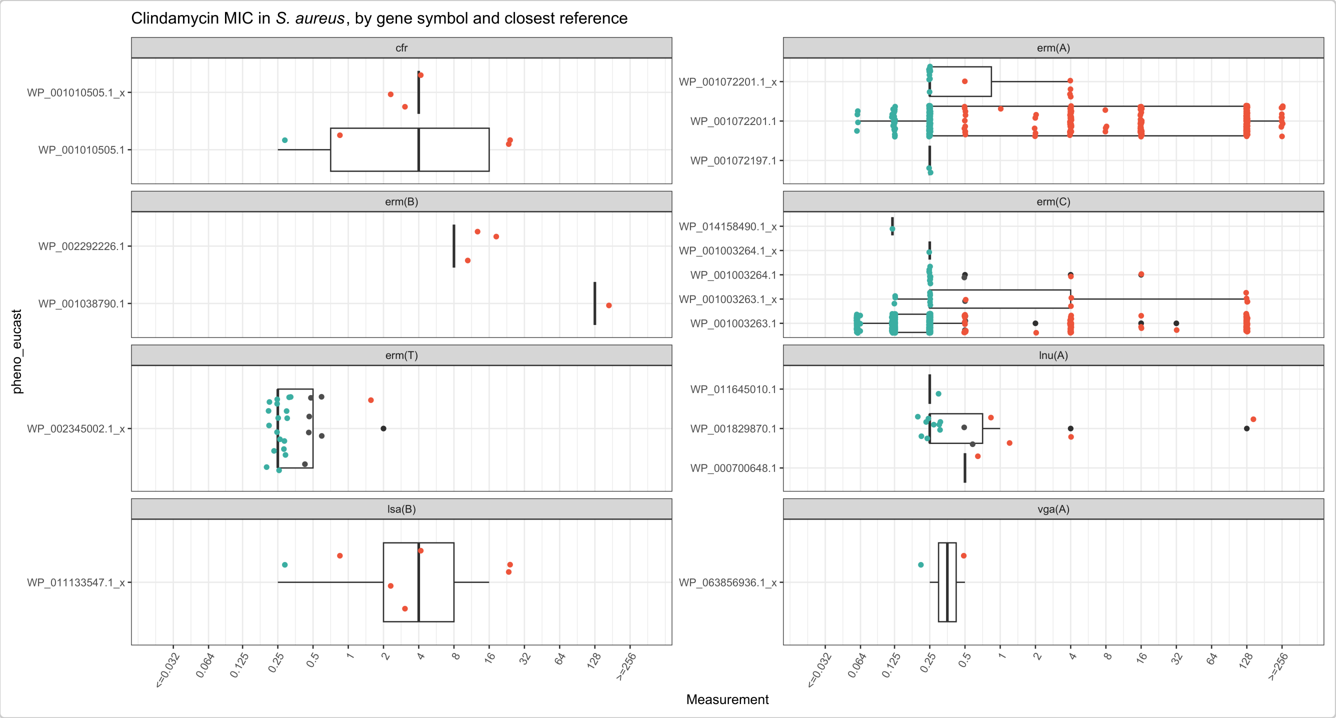


#### Association of specific variants with sequence types

Finally, we can also look at the association of specific marker variants with sequence types, if these data are available. This can be done by creating a binary matrix for the variants and then visualizing the associations with a bubble plot.

```
# Identify "High-Frequency" STs because we have many STs with only a few samples, which can make the plot cluttered and less informative. By filtering to include only STs with a certain number of samples (e.g., n >= 100), we can focus on the most prevalent STs in the dataset, which are likely to provide more meaningful insights into the associations between specific variants and sequence types.
high_freq_sts <- ST_data_CLI %>%
  count(ST) %>%
  filter(n >= 100) %>%
  pull(ST)

# Merge with ST data
cli_accession_st <- afp_CLI_public %>%
  inner_join(ST_data_CLI) %>%
  filter(ST %in% high_freq_sts)

# Calculate prevalence of variant in each ST
balloon_data <- cli_accession_st %>%
  group_by(ST, variant.label) %>%
  summarise(n_samples = n_distinct(id), .groups = "drop") %>%
  left_join(ST_data_CLI %>% count(ST, name = "total_st_n"), by = "ST") %>%
  mutate(percent_prevalence = (n_samples / total_st_n) * 100) %>%
  filter(percent_prevalence >= 10)

# Create the Balloon Plot showing only marker-ST combinations with at least 10% prevalence
ggplot(balloon_data, aes(x = ST, y = variant.label)) +
  geom_point(aes(size = n_samples, color = percent_prevalence)) +
  scale_color_viridis_c(option = "viridis", direction = -1) +
  scale_size_continuous(range = c(1, 10)) +
```

```
theme_minimal() +
theme(
  axis.text.x = element_text(size = 16, angle = 45, hjust = 1),
  axis.text.y = element_text(size = 16),
  legend.title = element_text(size = 16),
  legend.text = element_text(size = 14)
) +
labs(
  title = "Clindamycin Markers by Major Sequence Types",
  subtitle = "Analysis of STs with n >= 100",
  x = "Sequence Type (ST)",
  y = "Marker variant",
  size = "Sample Count",
  color = "Prevalence (%)"
)
```

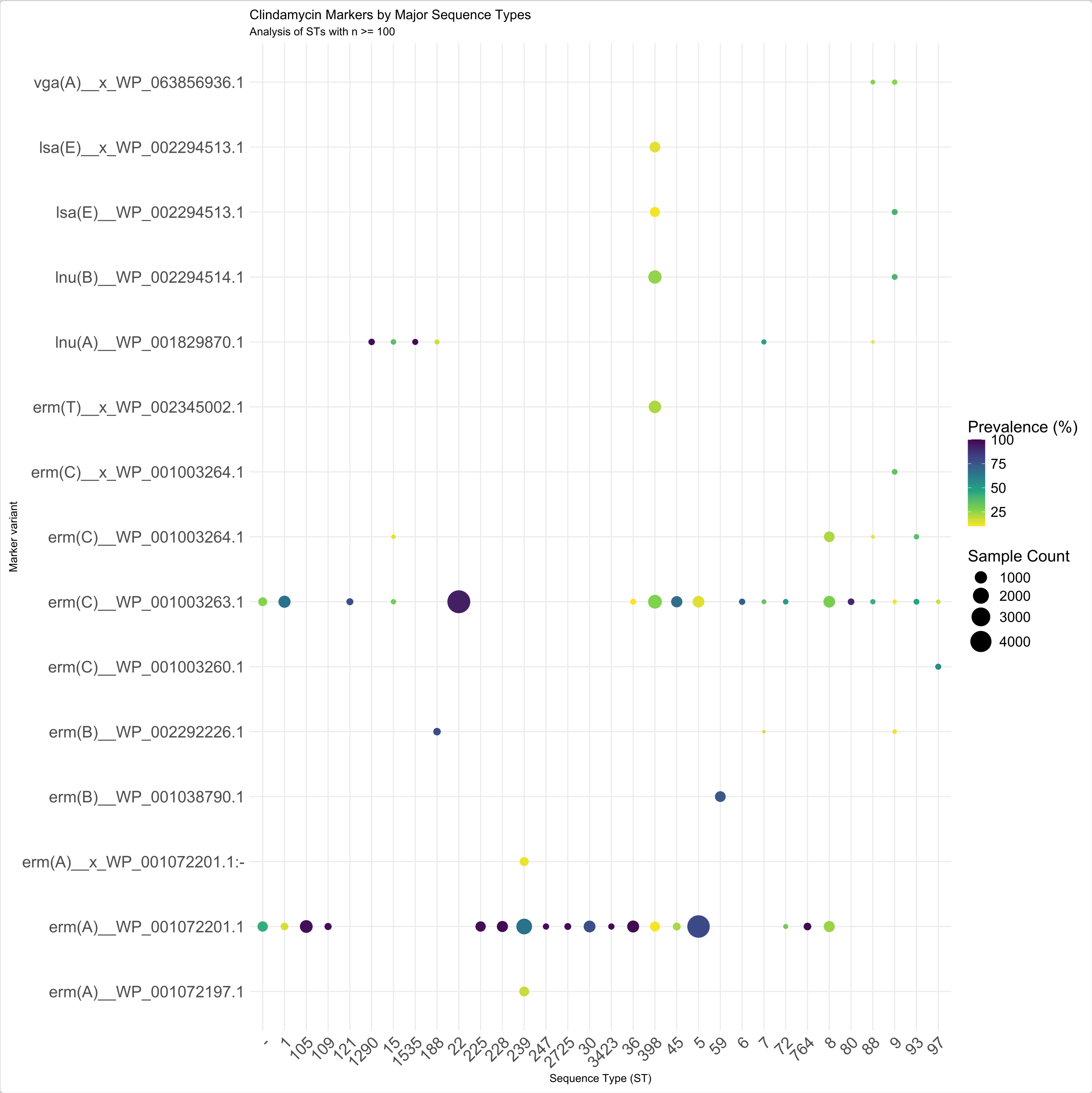


We can see from this last figure, that while erm(C)_WP0012364.1 is quite rare, erm(C)_WP0012363.1 is prevalent in certain STs, such as ST22. This suggests that there may be a clonal association of this variant with this ST, indicating resistance in this clone might be inducible and therefore more difficult to identify using standard AST.

### Analysing the impact of deletion variants on susceptibility

Richard Goodman

### Exploring catB3 deletion variants and impact on chloramphenicol susceptibility in *Escherichia coli*

### 1. Introduction

Infections caused by extended-spectrum beta-lactamase (ESBL)-producing Enterobacterales (ESBL-E) are a critical global health threat, often leaving clinicians with few treatment options beyond last-resort carbapenems.

In Malawi, first-line treatment for sepsis shifted from chloramphenicol (CHL) to ceftriaxone (CRO) in 2004, this was followed by a notable re-emergence of CHL susceptibility as its clinical use declined (Musicha et al., 2017).

A recent study (Graf et al., 2024) revealed that this "resensitisation" is frequently driven by the stable degradation of resistance genes rather than their total loss from the population. Specifically, insertion sequences like IS*26* and IS*5* have been identified as key drivers; IS*26* causes truncations in *catB3* (creating the non-functional variant *catB4*), while IS*5* can integrate into the promoter of *catA1*, effectively silencing its transcription.

Here we analyse a matched phenotype/genotype dataset used in the (Graf et al., 2024) study and publicly available datasets from NCBI to investigate these genotype-phenotype mismatches with the AMRgen package.

#### 1.1 Sourcing data from the DASSIM study

One of the datasets used to highlight genotype-phenotype mismatches in the Graf et al., 2024 paper was the "Developing an antimicrobial strategy for sepsis in Malawi" (DASSIM) dataset (Lewis et al. 2022).

The DASSIM study was an observational study of patients with sepsis admitted to Queen Elizabeth Central Hospital, Blantyre, Malawi. The aim was to understand the drivers of acquisition and long term carriage of ESBL-E in sepsis survivors. The DASSIM dataset contains faecal samples from community patients, inpatients and sepsis patients.

All genomic data was short-read sequenced on Illumina platforms at the Wellcome Sanger Institute.

Antimicrobial sensitivity testing (AST) was carried out on a subset of isolates using the disc-diffusion method using British Society for Antimicrobial Chemotherapy (BSAC) guidelines (https://bsac.org.uk/). AST was carried out for meropenem, amikacin, chloramphenicol, ciprofloxacin, co-trimoxazole and gentamicin.

However this dataset only contains the interpreted phenotype data (S/I/R) which is what we work with in this example.

#### 1.1a DASSIM Genotype data

We downloaded genome data from the following papers:

> Colonization dynamics of extended-spectrum beta-lactamase-producing Enterobacterales in the gut of Malawian adults Nat. Microbiol. 7, 1593–1604 (2022). Joseph M Lewis, Madalitso Mphasa, Rachel Banda, Matthew Beale, Eva Heinz, Jane Mallewa, Christopher Jewell, Nicholas R Thomson, Nicholas A Feasey

> Genomic analysis of extended-spectrum beta-lactamase (ESBL) producing Escherichia coli colonising adults in Blantyre, Malawi reveals previously undescribed diversity Microb. Genom. 9, mgen001035 (2023). Joseph M Lewis, Madalitso Mphasa, Rachel Banda, Matthew Beale, Jane Mallewa, Catherine Anscombe, Allan Zuza, Adam P Roberts, Eva Heinz, Nicholas Thomson, Nicholas A Feasey

> Genomic and antigenic diversity of colonising Klebsiella pneumoniae isolates mirrors that of invasive isolates in Blantyre, Malawi Microb. Genom. 8, 000778 (2022). Joseph M Lewis, Madalitso Mphasa, Rachel Banda, Matthew Beale, Jane Mallewa, Eva Heinz, Nicholas Thomson, Nicholas A Feasey

The genomes are deposited in the European Nucleotide Archive (ENA) under the project IDs PRJEB26677 and PRJEB36486.

The fastq files were downloaded, processed with cutadapt, filtered to >Q20 with FASTQC, and assembled with SPAdes v3.11.1 as described in Graf et al., 2024.

Antimicrobial resistance genes (ARGs) were called with AMRFinderPlus v4.0.23 to produce the data frame (DASSIM_geno), which is included in the AMRgen package as a data object `DASSIM_geno`.

##### 1.1b DASSIM Phenotype data

We downloaded the AST data from the blantyreESBL Github from Dr. Joe Lewis at https://github.com/joelewis101/blantyreESBL/raw/refs/heads/main/data/btESBL_pheno.rda.

This is included in the AMRgen package as a data object: `btESBL_pheno`.

Additional metadata relating to the isolates can be found in Supplementary Data 1 of the following paper:

> Molecular mechanisms of re-emerging chloramphenicol susceptibility in extended-spectrum beta-lactamase-producing Enterobacterales. Nat Commun 15, 9019 (2024). Fabrice E Graf, Richard N Goodman, Sarah Gallichan, Sally Forrest, Esther Picton-Barlow, Alice J Fraser, Minh-Duy Phan, Madalitso Mphasa, Alasdair T M Hubbard, Patrick Musicha, Mark A Schembri, Adam P Roberts, Thomas Edwards, Joseph M Lewis, Nicholas A Feasey.

This data is included in the AMRgen package as data object: `DASSIM_pheno_raw`.

### 2. Analysis of the DASSIM dataset

#### 2.1 Setting up R

First we set up R and load our libraries.

```r
library(AMRgen)
library(dplyr)
library(tidyr)
library(ggplot2)
```

#### 2.2 Format the phenotype data

The phenotype data in `btESBL_pheno` needs to be reformatted to long format, and sequence identifiers imported from `DASSIM_pheno_raw` so we can match the phenotype data to the genotypes.

```r
# Convert the S/I/R phenotype data to long format for easy use with AMRgen functions
DASSIM_pheno <- btESBL_pheno %>%
  pivot_longer(
    names_to = "drug",
    values_to = "pheno",
    cols = "amikacin":"meropenem"
  ) %>%
  mutate( # Standardise the terms to S, I, and R
    pheno = case_when(
      tolower(pheno) %in% c("sensitive", "susceptible", "s") ~ "S",
      tolower(pheno) %in% c("intermediate", "i") ~ "I",
      tolower(pheno) %in% c("resistant", "r") ~ "R",
      TRUE ~ NA_character_ # Any other unrecognised values become NA
    )
  ) %>%
  mutate(pheno = AMR::as.sir(pheno))

# add the sequence identifier from DASSIM_pheno_raw so we can match to genotype data
DASSIM_pheno <- DASSIM_pheno %>%
  left_join(DASSIM_pheno_raw %>% select(Strain_ID, seq, ST), join_by("supplier_name" ==
      "Strain_ID")) %>%
  rename(id = seq) %>%
  relocate(id) %>%
  mutate(mic = NA)

head(DASSIM_pheno)
#> # A tibble: 6 × 7
#>   id         supplier_name organism drug            pheno    ST mic
#>   <chr>      <chr>         <chr>    <chr>           <sir> <dbl> <lgl>
#> 1 ERR3426052 CAB10K        E. coli  amikacin        S       656 NA
#> 2 ERR3426052 CAB10K        E. coli  chloramphenicol S       656 NA
#> 3 ERR3426052 CAB10K        E. coli  ciprofloxacin   S       656 NA
#> 4 ERR3426052 CAB10K        E. coli  cotrimoxazole   R       656 NA
#> 5 ERR3426052 CAB10K        E. coli  gentamicin      S       656 NA
#> 6 ERR3426052 CAB10K        E. coli  meropenem       S       656 NA
```

#### 2.3 Check the genotype data

```r
head(DASSIM_geno)
#> # A tibble: 6 × 32
#>   id           marker      gene  mutation drug  drug_class `variation type` node
#>   <chr>        <chr>       <chr> <chr>    <ab>  <chr>      <chr>            <chr>
#> 1 26141_1_134 aadA5       aadA5 <NA>     STR1… Aminoglyc… Gene presence d… aadA5
#> 2 26141_1_134 dfrA17      dfrA… <NA>     NA    Trimethop… Gene presence d… dfrA…
#> 3 26141_1_134 arr         arr   <NA>     RFM:… Rifamycins Inactivating mu… arr
#> 4 26141_1_134 catB3       catB3 <NA>     CHL:… Phenicols  Gene presence d… catB3
#> 5 26141_1_134 blaOXA-1    blaO… <NA>     NA    Cephalosp… Gene presence d… blaO…
#> 6 26141_1_134 aac(6')-Ib… aac(… <NA>     AMK:… Aminoglyc… Gene presence d… aac(…
#> # ℹ 24 more variables: marker.label <chr>, `Protein id` <lgl>,
#> #   `Contig id` <chr>, Start <dbl>, Stop <dbl>, Strand <chr>,
#> #   `Gene symbol` <chr>, `Element name` <chr>, Scope <chr>, Type <chr>,
#> #   Subtype <chr>, Class <chr>, Subclass <chr>, Method <chr>,
#> #   `Target length` <dbl>, `Reference sequence length` <dbl>,
#> #   `% Coverage of reference` <dbl>, `% Identity to reference` <dbl>,
#> #   `Alignment length` <dbl>, `Closest reference accession` <chr>, …
```

#### 2.4 PPV Analysis of DASSIM dataset

The function `amr_ppv()` predicts positive predictive value of genetic markers (i.e. genes/mutations) for resistance among strains that carry these markers.

We will look at the markers associated with the antibiotics used in the AST assays for the DASSIM study.

- chloramphenicol
- amikacin
- gentamicin
- cotrimoxazole
- meropenem

For each of these first we create a binary matrix using `get_binary_matrix()` which takes our genotype (AMRfinderplus etc.) and phenotype (AST profile) datasets.

We can then plot the PPV graphs using `amr_ppv()`.

```r
# Get binary matrix
DASSIM_CHL_bin_mat <- get_binary_matrix(DASSIM_geno, DASSIM_pheno, pheno_drug =
        "chloramphenicol", sir_col = "pheno")
# Plot ppv
DASSIM_CHL_PPV <- amr_ppv(DASSIM_CHL_bin_mat, pheno_drug = "Chloramphenicol", sir_col =
        "pheno", upset_grid = FALSE)
```

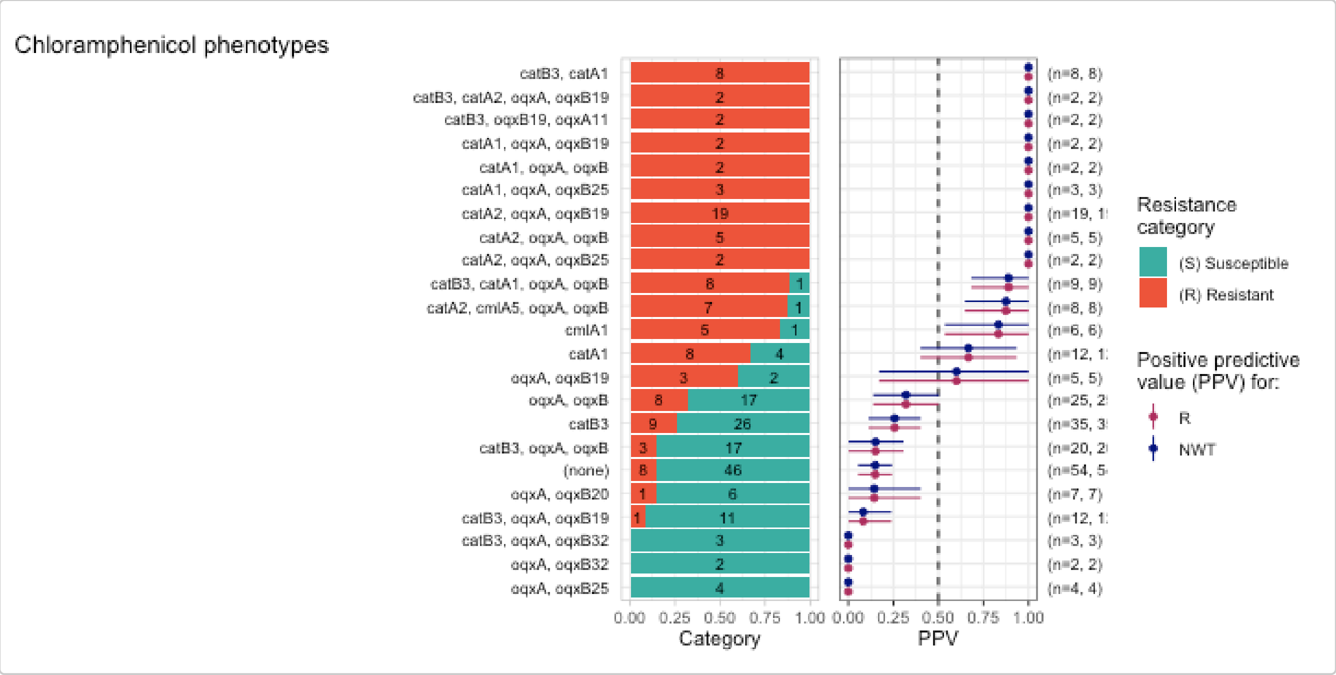


This result clearly shows how the detection of *catB3* is not a good predictor of resistance, whereas the detection of *catA2* is.

Now let's check how well aminoglycoside markers predict resistance to amikacin and gentamicin.

```
DASSIM_AMK_bin_mat <- get_binary_matrix(DASSIM_geno, DASSIM_pheno, pheno_drug =
        "amikacin", sir_col = "pheno")
DASSIM_AMK_PPV <- amr_ppv(DASSIM_AMK_bin_mat, pheno_drug = "amikacin", sir_col = "pheno",
        upset_grid = TRUE)
```

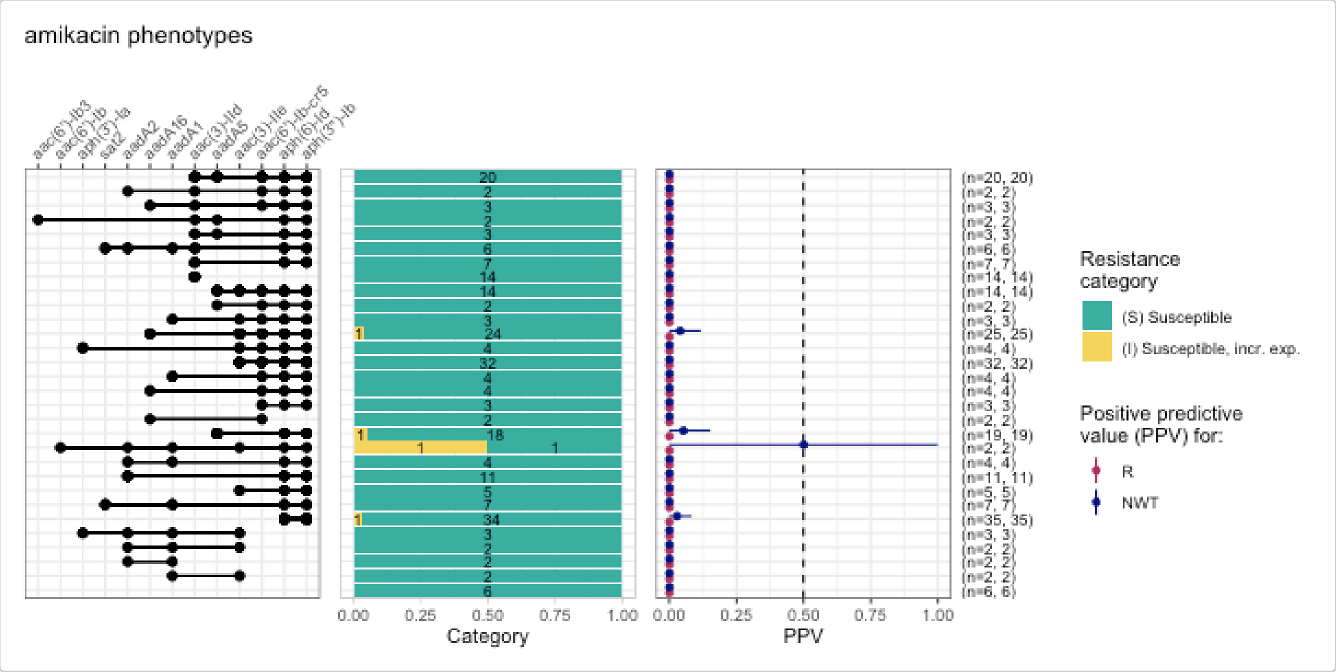


```
DASSIM_GEN_bin_mat <- get_binary_matrix(DASSIM_geno, DASSIM_pheno, pheno_drug =
        "gentamicin", sir_col = "pheno")
DASSIM_GEN_PPV <- amr_ppv(DASSIM_GEN_bin_mat, pheno_drug = "gentamicin", sir_col =
        "pheno", upset_grid = TRUE)
```

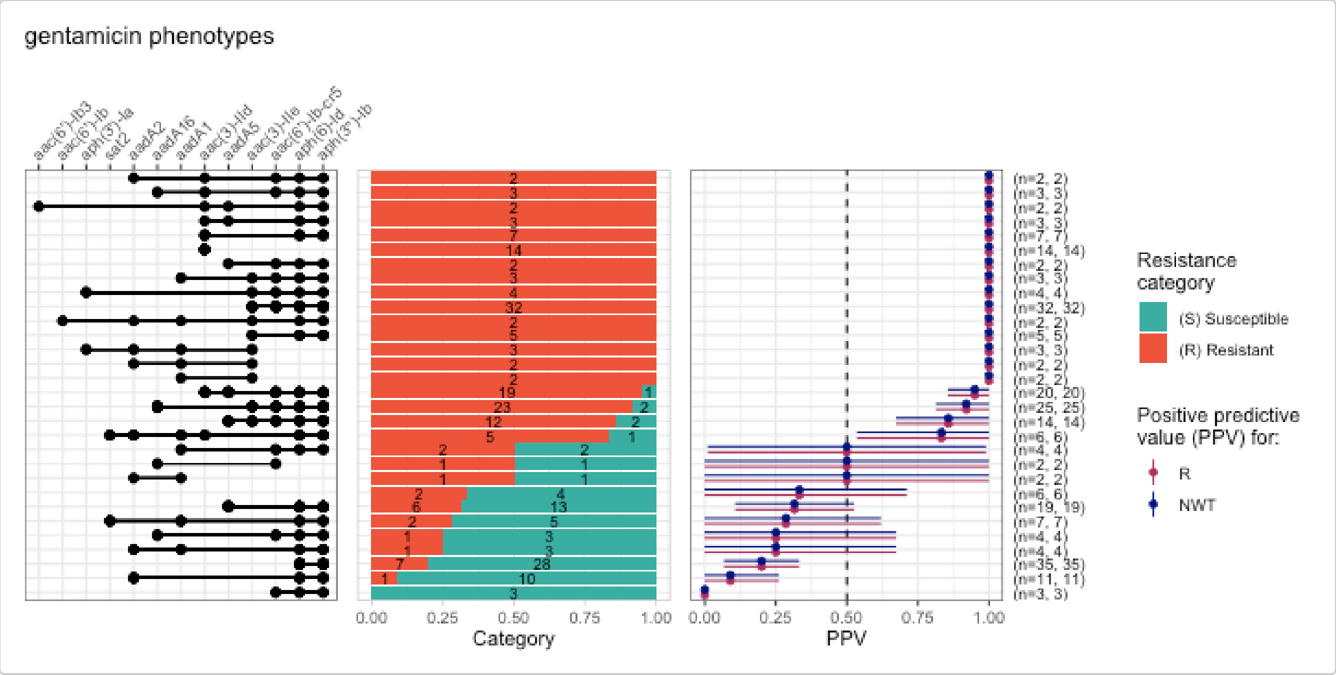


Here we see many of the aminoglycoside-associated resistance genes detected in these genomes are predictive of resistance to gentamicin, but none are associated with amikacin resistance. This is because amikacin is a semi-synthetic drug with an addition of a specific side chain, called the L-hydroxyaminobutyryl amide (HABA) group. This HABA side chain blocks the Aminoglycoside-Modifying Enzymes (AMEs) from reaching the sites on the molecule where they would normally attach their deactivating tags.

Now let's check how well markers associated with trimethoprims or sulfonamides predict resistance to co-trimoxazole.

```
DASSIM_SXT_bin_mat <- get_binary_matrix(DASSIM_geno, DASSIM_pheno, pheno_drug =
        "cotrimoxazole", sir_col = "pheno")
DASSIM_SXT_PPV <- amr_ppv(DASSIM_SXT_bin_mat, pheno_drug = "cotrimoxazole", sir_col =
        "pheno", upset_grid = TRUE)
```

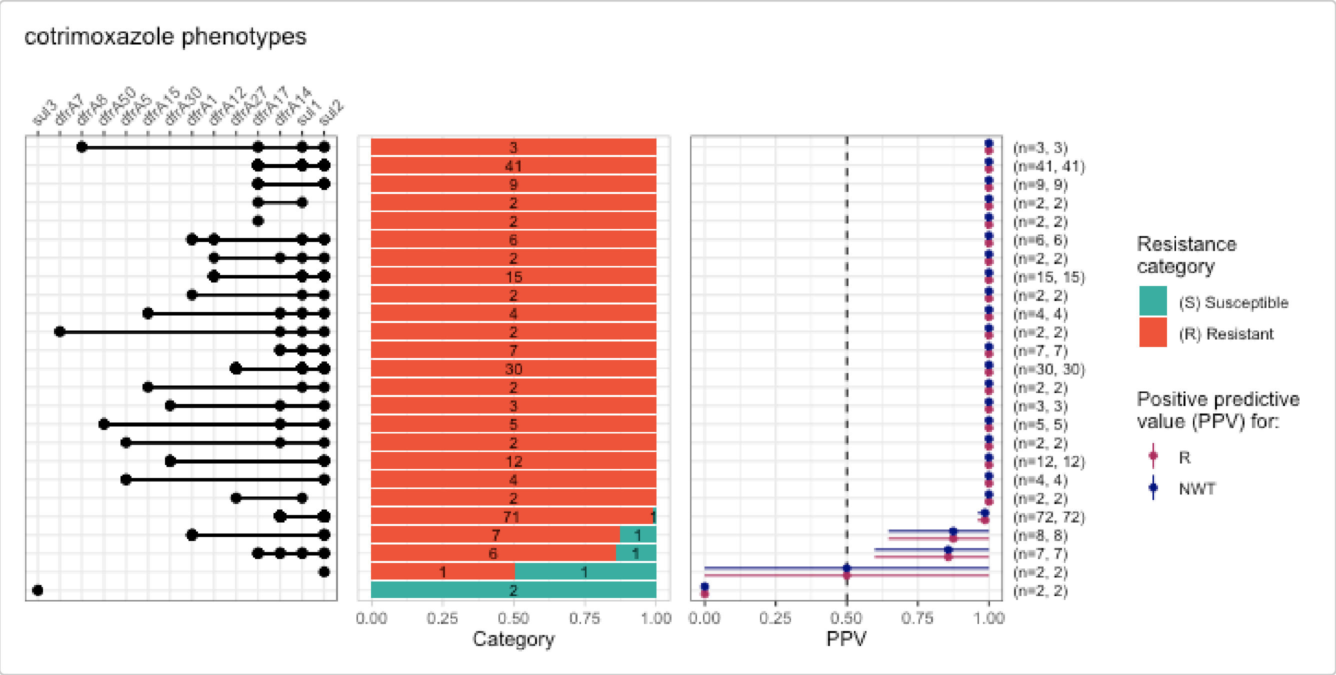


Co-trimoxazole is a combination drug made of sulfamethoxazole and trimethoprim. It is prescribed prophylactically in Malawi for HIV-positive individuals (Everett et al. 2011). Since it is a combination drug it requires both Sulfamethoxazole resistance genes (e.g., *sul*) and Trimethoprim resistance genes (e.g., *dfrA*).

Meropenem is a last resort carbapenem antibiotic. `amr_ppv()` by default returns all markers associated with beta-lactam resistance, for comparison with meropenem phenotypes. However while many beta-lactamases were detected, none are known carbapenemases (e.g., *bla*$_{NDM}$, *bla*$_{KPC}$, *bla*$_{VIM}$, *bla*$_{IMP}$, *bla*$_{OXA\text{-}48\text{-}like}$). Consistent with this only a single isolate, carrying multiple beta-lactamases, was phenotyped as resistant to meropenem.

```
DASSIM_MEM_bin_mat <- get_binary_matrix(DASSIM_geno, DASSIM_pheno, pheno_drug =
        "meropenem", sir_col = "pheno")
DASSIM_MEM_PPV <- amr_ppv(DASSIM_MEM_bin_mat, pheno_drug = "meropenem", sir_col = "pheno",
        upset_grid = TRUE)
```

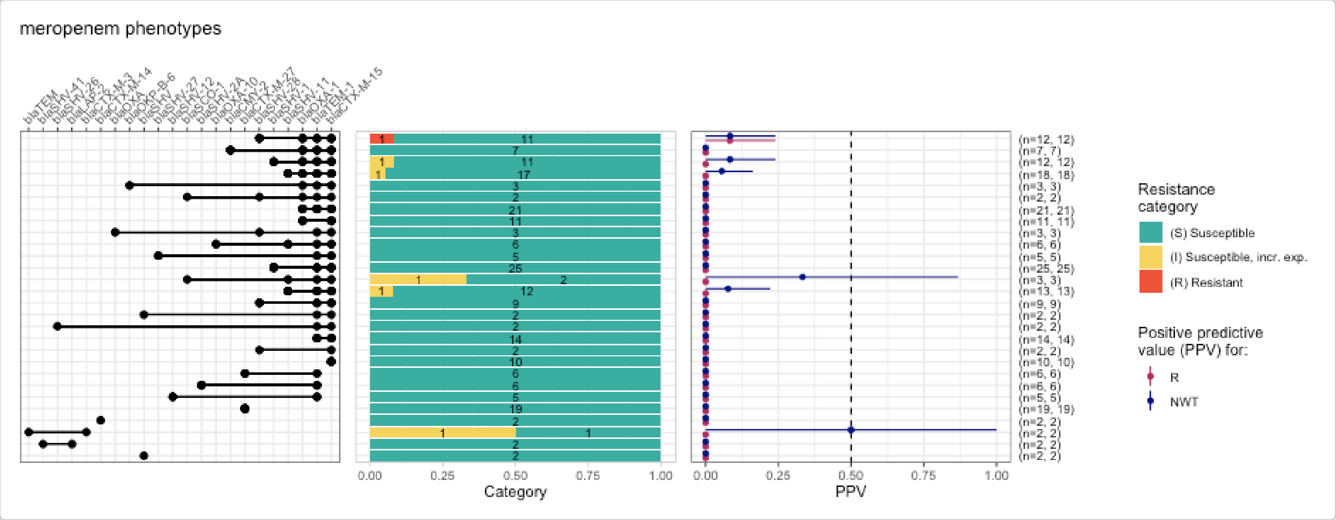


This analysis highlights how AMRgen can be used to explore genotype/phenotype associations for specific genetic markers related to a variety of antibiotics.

Next we'll explore the chloramphenicol susceptibility associated with the *catB3* gene with more functions of the AMRgen package. For this we need raw phenotype data (e.g., MIC or disc diffusion), so we will go to NCBI to download public datasets.

### 3. Analysis of publicly available phenotype and genotype data for chloramphenicol

#### 3.1 Importing public data from NCBI

The National Center for Biotechnology Information (NCBI) provides tools for analysing antimicrobial resistance (AMR).

**Phenotypes: NCBI AST**

The Antibiotic Susceptibility Test (AST) Browser serves as a centralised resource for viewing and filtering phenotypic susceptibility data, allowing researchers to correlate specific bacterial isolates with their phenotypic resistance profiles.

AST data can be retrieved directly from NCBI using the AMRgen functions `download_ncbi_pheno()` (slow but does not require authorisation) or `query_ncbi_bq_pheno()` (very fast, requires a Google Cloud account), or via the NCBI AST Browser. For more details see the Analysing Geno-Pheno Data vignette.

```r
# Download E. coli phenotype data from NCBI, filtering for chloramphenicol, and re-
        interpret with CLSI breakpoints
ecoli_pheno_ncbi_via_biosample <- download_ncbi_pheno(
  species = "E. coli",
  pheno_drug = "chloramphenicol",
  reformat = TRUE,
  interpret_clsi = TRUE
)


# Download E. coli AST data from NCBI via Google Cloud, filtering for chloramphenicol, and
        re-interpret with CLSI breakpoints

install.packages("bigrquery")
library(bigrquery)
bigrquery::bq_auth()

# replace xxx with your project id
ecoli_pheno_ncbi_via_cloud <- query_ncbi_bq_pheno(
  taxgroup = "E.coli and Shigella",
  pheno_drug = "chloramphenicol",
  project_id = "xxx"
)

ecoli_pheno_ncbi_via_cloud_interpreted <- import_ncbi_pheno(ecoli_pheno_ncbi_via_cloud,
  interpret_clsi = TRUE
)
```

Alternatively, we can navigate to the *NCBI Antibiotic Susceptibility Test (AST) Browser* in a web browser and search for

```
chloramphenicol AND Escherichia
```

https://www.ncbi.nlm.nih.gov/pathogens/ast#chloramphenicol%20AND%20Escherichia

Save this as a `tsv` file: `CHL_Ecoli_asts.tsv`, and import it using the AMRgen function `import_pheno`.

```r
# import phenotype data
NCBI_Ecoli_pheno_chl <- import_pheno("data-raw/CHL_Ecoli_asts.tsv", format = "ncbi")
```

A copy of this imported data (downloaded March 2026) is included in AMRgen as data frame `NCBI_Ecoli_pheno_chl`.

```r
head(NCBI_Ecoli_pheno_chl)
#> # A tibble: 6 × 25
#>   id     drug      mic  disk guideline method platform pheno_provided spp_pheno
#>   <chr> <ab>    <mic> <dsk> <chr>     <chr>  <chr>    <chr>          <mo>
#> 1 SAMN… CHL:… <=0.03    NA EUCAST    broth… <NA>     not defined    B_ESCHR_COL…
#> 2 SAMN… CHL:…    4.00   NA EUCAST    broth… <NA>     not defined    B_ESCHR_COL…
#> 3 SAMN… CHL:…    2.00   NA EUCAST    broth… <NA>     not defined    B_ESCHR_COL…
#> 4 SAMN… CHL:… <=0.03    NA EUCAST    broth… <NA>     not defined    B_ESCHR_COL…
#> 5 SAMN… CHL:…    2.00   NA EUCAST    broth… <NA>     not defined    B_ESCHR_COL…
#> 6 SAMN… CHL:…    1.00   NA EUCAST    broth… <NA>     not defined    B_ESCHR_COL…
```

```
#> # ℹ 16 more variables: `Organism group` <chr>, `Scientific name` <chr>,
#> #   `Isolation type` <chr>, Location <chr>, `Isolation source` <chr>,
#> #   Isolate <chr>, Antibiotic <chr>, `Resistance phenotype` <chr>,
#> #   `Measurement sign` <chr>, `MIC (mg/L)` <dbl>, `Disk diffusion (mm)` <dbl>,
#> #   `Laboratory typing platform` <chr>, Vendor <chr>,
#> #   `Laboratory typing method version or reagent` <chr>,
#> #   `Testing standard` <chr>, `Create date` <dttm>
```

###### Genotypes: MicroBIGG-E

The Microbial Browser for Identification of Genetic and Genomic Elements (MicroBIGG-E) is a specialised portal within the NCBI Pathogen Detection system that enables users to query a database of over 46,000 isolates to identify specific AMR genes and point mutations.

Navigate to the *NCBI Pathogen Detection Microbial Browser for Identification of Genetic and Genomic Elements (MicroBIGG-E)* in a web browser and search for:

```
chloramphenicol AND Escherichia
```

https://www.ncbi.nlm.nih.gov/pathogens/microbigge/#chloramphenicol%20AND%20Escherichia

Save this as a `tsv` file: `CHL-R_Ecoli_microbigge.tsv`, and import it using the AMRgen function `import_gheno`.

```
# import phenotype data
MICROBIGGE_Ecoli_CHLR <- import_geno("data-raw/CHL-R_Ecoli_microbigge.tsv", format =
        "amrfp", sample_col = "BioSample")
```

A copy of this imported data (downloaded March 2026) is included in AMRgen as data frame `MICROBIGGE_Ecoli_CHLR`.

```
head(MICROBIGGE_Ecoli_CHLR)
#> # A tibble: 6 × 27
#>   id           marker gene  mutation drug_agent drug_class `variation type` node
#>   <chr>        <chr>  <chr> <chr>    <ab>       <chr>      <chr>            <chr>
#> 1 SAMN008293… catA1  catA1 <NA>     CHL: Chlo… Phenicols  Gene presence d… catA1
#> 2 SAMN018857… catA1  catA1 <NA>     CHL: Chlo… Phenicols  Gene presence d… catA1
#> 3 SAMN026875… catA1  catA1 <NA>     CHL: Chlo… Phenicols  Gene presence d… catA1
#> 4 SAMN028020… catA1  catA1 <NA>     CHL: Chlo… Phenicols  Gene presence d… catA1
#> 5 SAMN028018… catB3  catB3 <NA>     CHL: Chlo… Phenicols  Inactivating mu… catB3
#> 6 SAMN031982… catA1  catA1 <NA>     CHL: Chlo… Phenicols  Gene presence d… catA1
#> # ℹ 19 more variables: marker.label <chr>, `Scientific name` <chr>,
#> #   Protein <chr>, Isolate <chr>, Contig <chr>, Start <dbl>, Stop <dbl>,
#> #   Strand <chr>, `Element symbol` <chr>, `Element name` <chr>, Type <chr>,
#> #   Scope <chr>, Subtype <chr>, Class <chr>, Subclass <chr>, Method <chr>,
#> #   `% Coverage of reference` <dbl>, `% Identity to reference` <dbl>,
#> #   subclass_to_parse <chr>
```

#### 3.2 Filter data to samples with chloramphenicol phenotype data, and chloramphenicol genotypic markers detected

```
# filter AST data, re-interpret using CLSI breakpoints
AST_pheno <- NCBI_Ecoli_pheno_chl %>%
  filter(id %in% MICROBIGGE_Ecoli_CHLR$id) %>%
  interpret_pheno(interpret_clsi = TRUE)
#> Warning: There was 1 warning in `mutate()`.
#> ℹ In argument: `across(...)`.
#> Caused by warning:
#> ! Some MICs were converted to the nearest higher log2 level, following the CLSI
#> interpretation guideline.
MB_CHLR_geno <- MICROBIGGE_Ecoli_CHLR %>% filter(id %in% NCBI_Ecoli_pheno_chl$id)

# check how many samples we have
length(unique(AST_pheno$id))
#> [1] 410

# check the genes
MB_CHLR_geno %>% count(gene)
#> # A tibble: 6 × 2
#>   gene       n
#>   <chr> <int>
#> 1 catA1    213
#> 2 catA2     16
#> 3 catB3    278
#> 4 cmlA1    350
#> 5 cmlA5     12
#> 6 cmlA6      6

# filter to find samples with catB3
MB_CATB3_geno <- MB_CHLR_geno %>% filter(gene == "catB3")
MB_nonCATB3_geno <- MB_CHLR_geno %>% filter(gene != "catB3")

# filter AST data to samples with catB3 and no other chloramphenicol markers
AST_CATB3_pheno <- AST_pheno %>%
  filter(id %in% MB_CATB3_geno$id) %>%
  filter(!(id %in% MB_nonCATB3_geno$id))

# check how many samples we have with catB3
length(unique(AST_CATB3_pheno$id))
#> [1] 116
```

#### 3.3 Exploring chloramphenicol phenotype distributions with assay_by_var()

Now we can explore phenotype distributions based on MIC data using the `assay_by_var()` function. We visualise the MIC distribution with `assay_by_var()` on all chloramphenicol AST data.

```
AST_pheno <- AST_pheno %>%
  mutate(across(all_of("mic"), ~ AMR::as.mic(.x, round_to_next_log2 = TRUE)))

assay_by_var(
  pheno_table = AST_pheno,
  pheno_drug = "chloramphenicol",
```

```
  measure = "mic",
  colour_by = "pheno_clsi",
  species = "Escherichia coli"
)
```

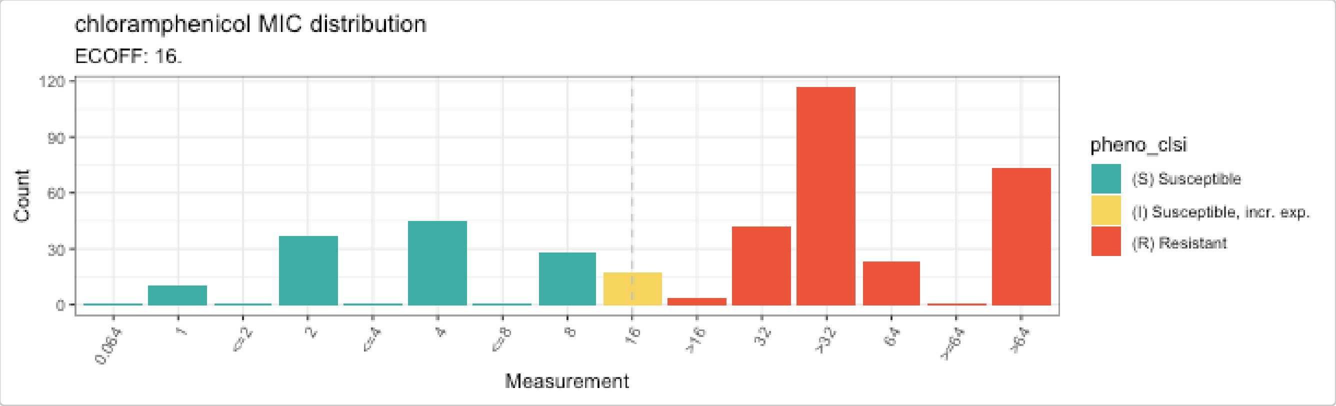


Next we can visualise the MIC distribution with `assay_by_var()` on the AST data of isolates containing the *catB3* gene only

```
# CATB3 specific

assay_by_var(
  pheno_table = AST_CATB3_pheno,
  pheno_drug = "CHL",
  measure = "mic",
  colour_by = "pheno_clsi",
  species = "Escherichia coli"
) +
  labs(title = "Chloramphenicol MIC distribution for isolates with catB3 only")
```

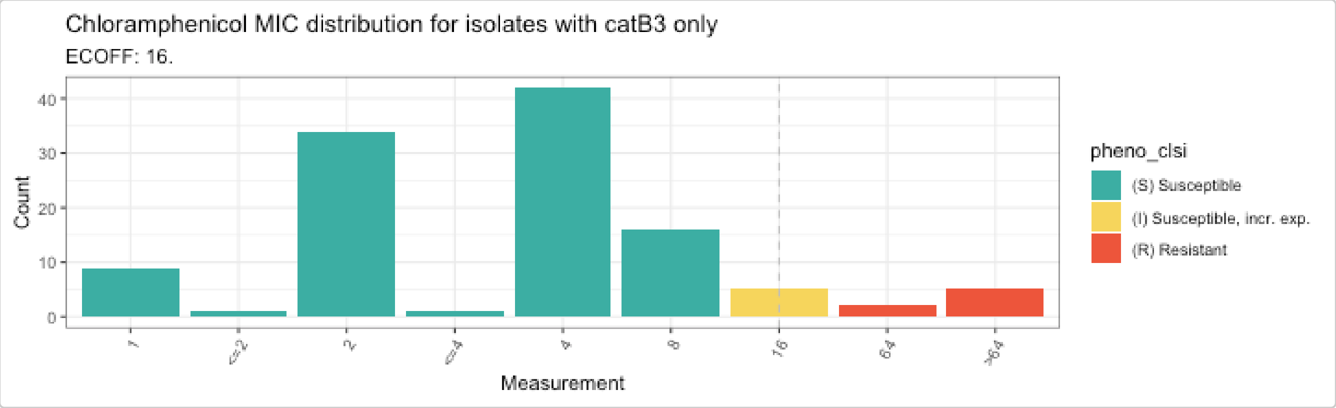


The distribution shifts to the right and towards sensitive for isolates with the *catB3* gene only, when compared to those with any chloramphenicol-associated gene.

We can then split the plot based on whether the the *catB3* gene is truncated or not. This is calculated as a percentage of coverage, with 100% aligning across the entire length of the gene and <100% showing a truncation or deletion (see [Graf et al., 2024] for more about the IS*26* mediated truncation of *catB3*).

```
# add genotype data to the phenotype table for isolates with catB3 alone
AST_CATB3_pheno_2 <- MB_CATB3_geno %>%
  select(id, gene, `% Coverage of reference`) %>%
  distinct(id, .keep_all = TRUE) %>%
  right_join(AST_CATB3_pheno, by = "id")
```

```r
# check coverage, all values are either 100% or 66.7-70%
AST_CATB3_pheno_2 %>% count(`% Coverage of reference`)
#> # A tibble: 4 x 2
#>   `% Coverage of reference`     n
#>                       <dbl> <int>
#> 1                      66.7     1
#> 2                      69.5    25
#> 3                      70      84
#> 4                     100       6

# define a grouping variable 'truncation' indicating samples with full coverage vs <=70%
AST_CATB3_pheno_3 <- AST_CATB3_pheno_2 %>%
  mutate(truncation = ifelse(`% Coverage of reference` > 70, "100% catB3 coverage", "<= 70
        % catB3 coverage"))

# plot the MIC distribution for these 2 groups
MIC_dist_by_cov <- assay_by_var(
  pheno_table = AST_CATB3_pheno_3,
  pheno_drug = "Chloramphenicol",
  measure = "mic",
  colour_by = "pheno_clsi",
  facet_by = "truncation",
  measure_axis_label = "MIC (mg/L)",
  colour_legend_label = "Phenotype (CLSI)"
)

MIC_dist_by_cov
```

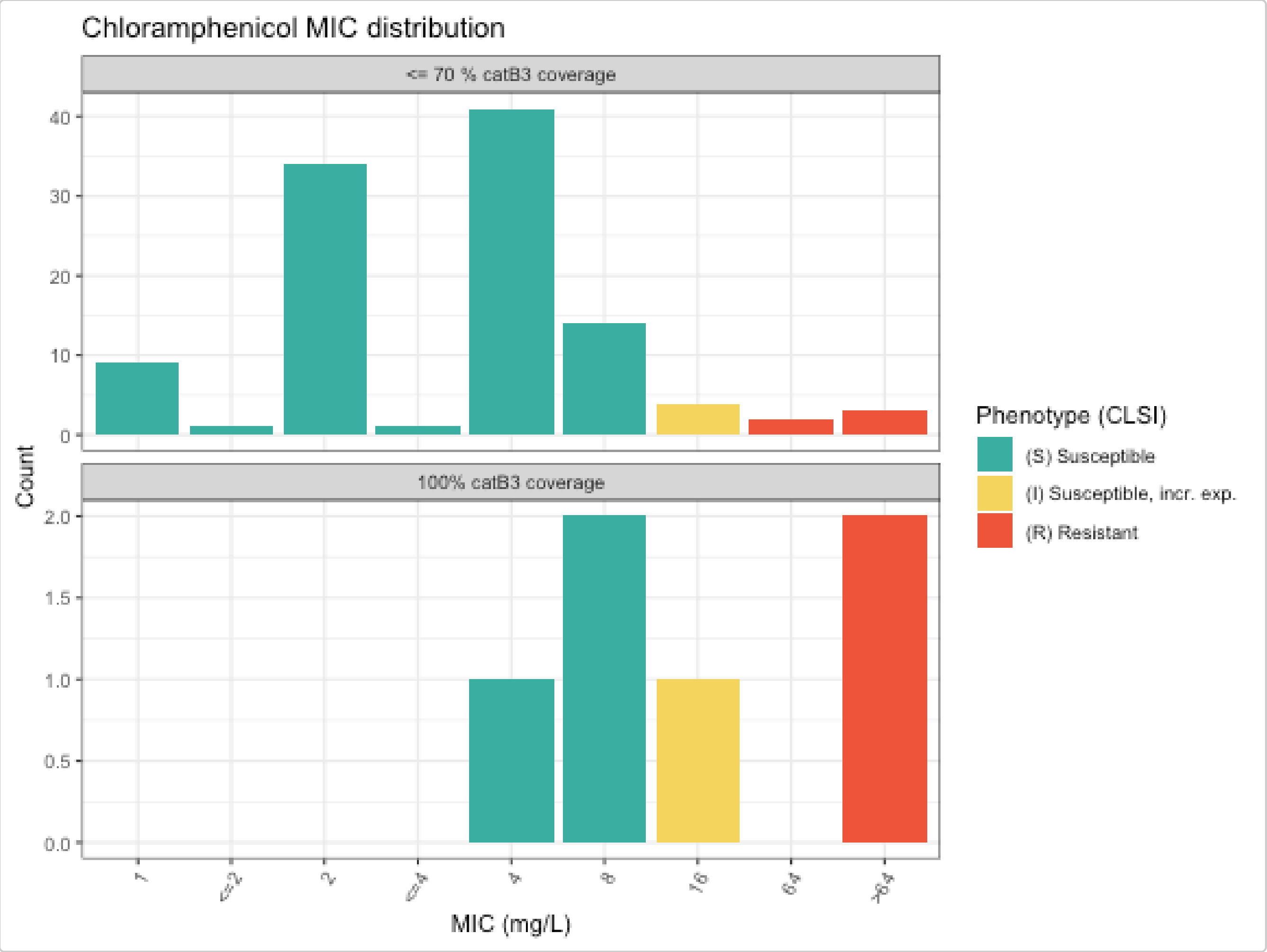


```
# check counts and median MIC per group
AST_CATB3_pheno_3 %>%
  group_by(truncation) %>%
  summarise(median = median(mic, na.rm = T), n = n(), R = sum(pheno_clsi == "S"))
#> # A tibble: 2 × 4
#>   truncation              median     n     R
#>   <chr>                    <dbl> <int> <int>
#> 1 100% catB3 coverage         12     6     3
#> 2 <= 70 % catB3 coverage       4   110   100
```

As we can see, isolates with truncated *catB3* genes (n=110) have median MIC of 4 mg/L, and most (n=100) were classed as susceptible (CLSI breakpoint ≤8 mg/L; note there are no EUCAST breakpoints). In contrast, those with full-length *catB3* genes (n=6) had higher values, and only 3 were classed as susceptible.

#### 3.4 Analysing genotype and phenotype data with `amr_ppv()`

Now let's look at the geno-pheno associations across all the isolates with matched data. We first build a binary matrix using the genotype and phenotype tables as input with `get_binary_matrix()`

Then we can view it as an upset grid, with a upset plot, SIR stacked barplot and positive predictive value (ppv) using the `amr_ppv()` function.

```
MB_CHLR_geno <- MB_CHLR_geno %>%
  mutate(marker = ifelse(marker == "catB3" & `% Coverage of reference` < 100, "catB3
        (truncated)", marker))

CHL_bin_mat <- get_binary_matrix(MB_CHLR_geno,
  AST_pheno,
  pheno_drug = "CHL",
  geno_class = c("Phenicols"),
  sir_col = "pheno_clsi",
  keep_assay_values = TRUE
)

CHL_PPV <- amr_ppv(CHL_bin_mat,
  pheno_drug = "Chloramphenicol",
  geno_class = c("Phenicols"),
  sir_col = "pheno_clsi",
  upset_grid = TRUE,
  assay = "mic",
  plot_assay = TRUE,
  order = "value"
)
```

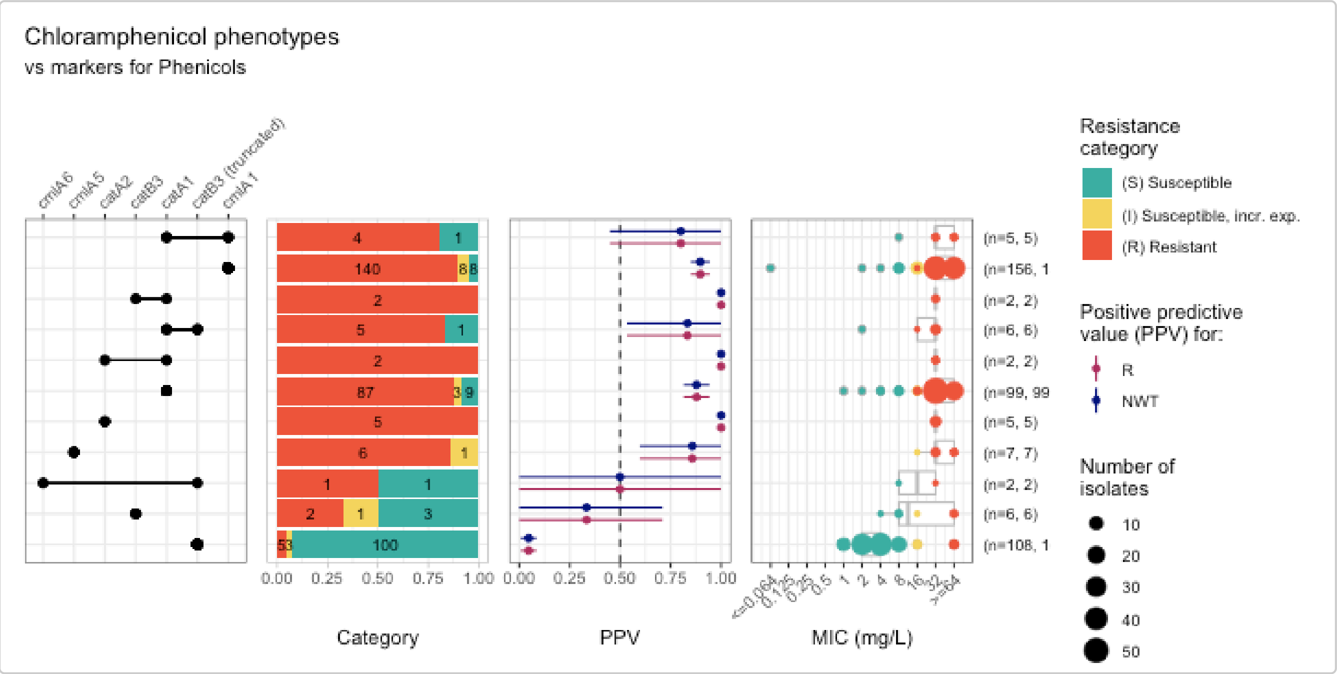


#### 3.5 Logistic regression

We can plot a coefficient plot using the `amr_logistic()` function to show the statistical relationship between chloramphenicol resistance genes and phenotypic resistance to chloramphenicol.

We just need the binary matrix as input (from `get_binary_matrix()`) and we can define the antibiotic, the column containing the ecoff and a threshold for the number of samples a marker is present in.

```
CHL_logist <- amr_logistic(
  binary_matrix = CHL_bin_mat,
  pheno_drug = "chloramphenicol",
  ecoff_col = "ecoff",
  maf = 10,
```

```
  single_plot = TRUE
)
```

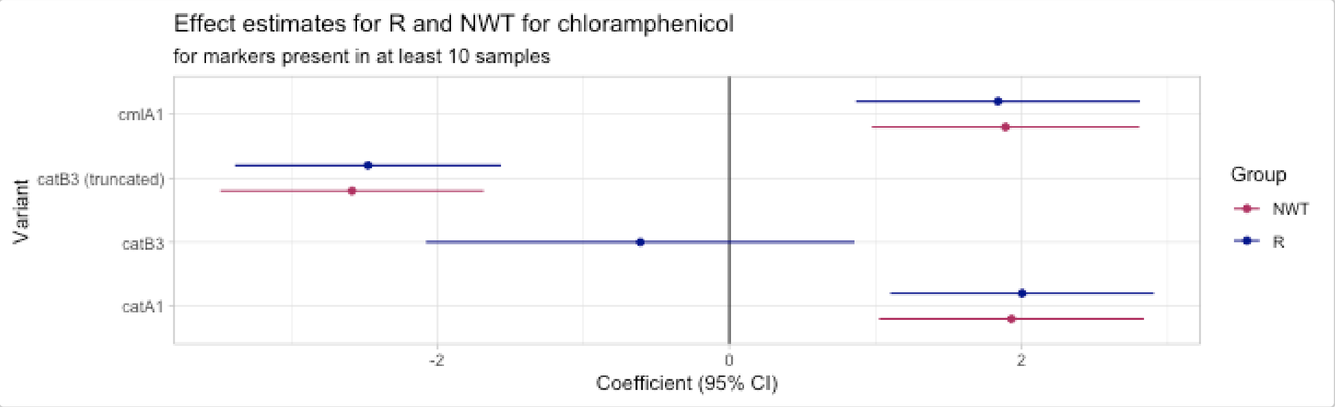


```
# model coefficients
CHL_logist$modelR
#> # A tibble: 5 × 5
#>   marker              est ci.lower ci.upper         pval
#>   <chr>             <dbl>    <dbl>    <dbl>        <dbl>
#> 1 (Intercept)       0.295   -0.555    1.14  0.496
#> 2 cmlA1             1.84     0.866    2.81  0.000211
#> 3 catA1             2.00     1.10     2.91  0.0000134
#> 4 catB3            -0.611   -2.08     0.857 0.415
#> 5 catB3 (truncated) -2.47   -3.38    -1.56  0.0000000980
```

This plot shows that both *catA1* and *cmlA1* have a strong positive association with phenotypic resistance whereas truncated *catB3* has a strong negative association with resistance (i.e. a stronger association with susceptibility to chloramphenicol).

Experimental evolution analysis by [Graf et al., 2024]() suggested that these *catB3* silencing events are highly stable under antibiotic pressure, reinforcing the potential for chloramphenicol to be reintroduced as a targeted reserve agent for ESBL-E infections in low-resource settings.

### Example exploring gene vs mutation combinations

**Kara Tsang**

### Analysing meropenem resistance *Klebsiella pneumoniae*

This vignette demonstrates an example of how to investigate associations between *Klebsiella pneumoniae* meropenem antibiotic susceptibility testing (AST) data and AMR genotype data (Kleborate, AMRFinderPlus and RGI), including examining the orthogonal effects of acquired carbapenemase genes and porin mutations.

We will be using the AST data and AMR genotyping outputs for n=1490 isolates from the European Survey of Carbapenemase-Producing Enterobacteriaceae (EuSCAPE) published as ‘Epidemic of carbapenem-resistant *Klebsiella pneumoniae* in Europe is driven by nosocomial spread’ by David, S., et al. Nature Microbiology, 2016.

Whole genome sequence reads were downloaded from NCBI Bioproject PRJEB10018 / European Nucleotide Archive ERP011196, trimmed using Trim Galore v0.5.0, and assembled using Unicycler v0.5.0. The assembled genomes were then run through each AMR genotyper:

- Kleborate v3.1.3
- Kleborate version development branch - commit #4ec1dcb on March 17, 2026
- Resistance Gene Idenfier (RGI v6.0.6) - using the Comprehensive Antibiotic Resistance Database (CARD, v4.0.1)

AMRFinderPlus results were generated by EMBL-EBI Antimicrobial Resistance Portal using - AMRFinderPlus v4.0 with NCBI Reference Gene Catalog database version 2025-07-16.1.

#### Load the required packages

```
library(AMRgen)
library(dplyr)
library(ggplot2)
library(tidyr)
library(stringr)
```

### Phenotype Data

The `download_ebi()` function lets you load phenotype data and interpret susceptible, intermediate, and resistant phenotypes using EUCAST breakpoints and ECOFF. A copy of the data objected produced below is available in the AMRgen package as `kp_mero_euscape`.

```
# Download Klebsiella pneumoniae AST data from EBI, filtering for meropenem and re-
        interpret with EUCAST breakpoints and ECOFF

kp_mero <- download_ebi(
```

```r
  pheno_drug = "meropenem",
  species = "Klebsiella pneumoniae",
  reformat = TRUE,
  interpret_eucast = TRUE,
  interpret_ecoff = TRUE
)

# Filter for isolates in EuSCAPE paper (PMID: 31358985)
kp_mero_euscape <- kp_mero %>% filter(grepl("31358985", source))

# There are assemblies from NCBI that are flagged for contamination and supposed to be
        excluded. For example, see SAMEA3729690
        (https://www.ncbi.nlm.nih.gov/datasets/genome/?biosample=SAMEA3729690)

contaminated_assemblies <- c("SAMEA3729690", "SAMEA3721062", "SAMEA3721052",
        "SAMEA3720966", "SAMEA3673128", "SAMEA3538742", "SAMEA3721188", "SAMEA3649589",
        "SAMEA3538652", "SAMEA3649503", "SAMEA3538911", "SAMEA3727711", "SAMEA3649452",
        "SAMEA3649453", "SAMEA3649454", "SAMEA3649467", "SAMEA3721063", "SAMEA3538862",
        "SAMEA3538667", "SAMEA3673004", "SAMEA3729818", "SAMEA3729660", "SAMEA3673078",
        "SAMEA3673097")

# Remove contaminated assemblies from phenotype list
kp_mero_euscape <- kp_mero_euscape %>%
  filter(!id %in% contaminated_assemblies)
```

Check the data frame

```r
head(kp_mero_euscape)
#> # A tibble: 6 × 43
#>   id         drug    mic  disk pheno_provided pheno_eucast ecoff guideline method
#>   <chr>      <ab>   <mic> <dsk> <sir>          <sir>        <sir> <chr>     <chr>
#> 1 SAMEA372… MEM:…  2.00    NA NA             S            NWT   <NA>      broth…
#> 2 SAMEA372… MEM:…  4.00    NA NA             I            NWT   <NA>      broth…
#> 3 SAMEA372… MEM:…  0.12    NA NA             S            WT    <NA>      broth…
#> 4 SAMEA372… MEM:…  1.00    NA NA             S            NWT   <NA>      broth…
#> 5 SAMEA372… MEM:…  4.00    NA NA             I            NWT   <NA>      broth…
#> 6 SAMEA372… MEM:…  8.00    NA NA             I            NWT   <NA>      broth…
#> # ℹ 34 more variables: platform <chr>, source <chr>, spp_pheno <mo>,
#> #   SRA_accession <chr>, assembly_ID <chr>, collection_year <int>,
#> #   ISO_country_code <chr>, host <chr>, host_age <chr>, host_sex <chr>,
#> #   isolate <chr>, isolation_source <chr>, isolation_source_category <chr>,
#> #   isolation_latitude <chr>, isolation_longitude <chr>, genus <chr>,
#> #   organism <chr>, Updated_phenotype_CLSI <chr>,
#> #   Updated_phenotype_EUCAST <chr>, used_ECOFF <chr>, database <chr>, …
```

### Phenotype Data Summary

Summarize the downloaded phenotype data and plot the minimum inhibitory concentration (MIC) distributions with EUCAST breakpoints and ECOFF.

```r
# Summary of meropenem phenotype data including S/I/R count using EUCAST breakpoint and
        ECOFF
summarise_pheno(kp_mero_euscape, pheno_cols = c("pheno_eucast", "ecoff"))
#> $uniques
#> # A tibble: 7 × 2
```

```
#>   column    n_unique
#>   <chr>        <int>
#> 1 id            1490
#> 2 drug             1
#> 3 spp_pheno        1
#> 4 method           1
#> 5 platform         1
#> 6 guideline        1
#> 7 source           2
#>
#> $drugs
#> # A tibble: 1 × 4
#>   drug           drug_name spp_pheno                mic
#>   <ab>           <chr>     <chr>                  <int>
#> 1 MEM: Meropenem Meropenem Klebsiella pneumoniae  1490
#>
#> $details
#> # A tibble: 2 × 8
#>   drug           drug_name spp_pheno      method platform guideline source   mic
#>   <ab>           <chr>     <chr>          <chr>  <chr>    <chr>     <chr>  <int>
#> 1 MEM: Meropenem Meropenem Klebsiella pn… broth… <NA>     <NA>      31358…   628
#> 2 MEM: Meropenem Meropenem Klebsiella pn… broth… <NA>     <NA>      31358…   862
#>
#> $pheno_counts_list
#> $pheno_counts_list$pheno_eucast
#> # A tibble: 1 × 6
#>   drug           drug_name spp_pheno                  S     I     R
#>   <ab>           <chr>     <chr>                  <int> <int> <int>
#> 1 MEM: Meropenem Meropenem Klebsiella pneumoniae    973   126   391
#>
#> $pheno_counts_list$ecoff
#> # A tibble: 1 × 5
#>   drug           drug_name spp_pheno                 WT   NWT
#>   <ab>           <chr>     <chr>                  <int> <int>
#> 1 MEM: Meropenem Meropenem Klebsiella pneumoniae    710   780

# MIC distribution coloured by phenotype interpretation using EUCAST breakpoint
assay_by_var(
  pheno_table = kp_mero_euscape,
  pheno_drug = "Meropenem",
  measure = "mic",
  colour_by = "pheno_eucast",
  species = "Klebsiella pneumoniae"
)
#>   MIC breakpoints determined using AMR package: S <= 2 and R > 8
#>   NOTE: Multiple breakpoint entries, for different sites: Non-meningitis; Meningitis.
       Using the one with the highest S breakpoint (Non-meningitis).
```

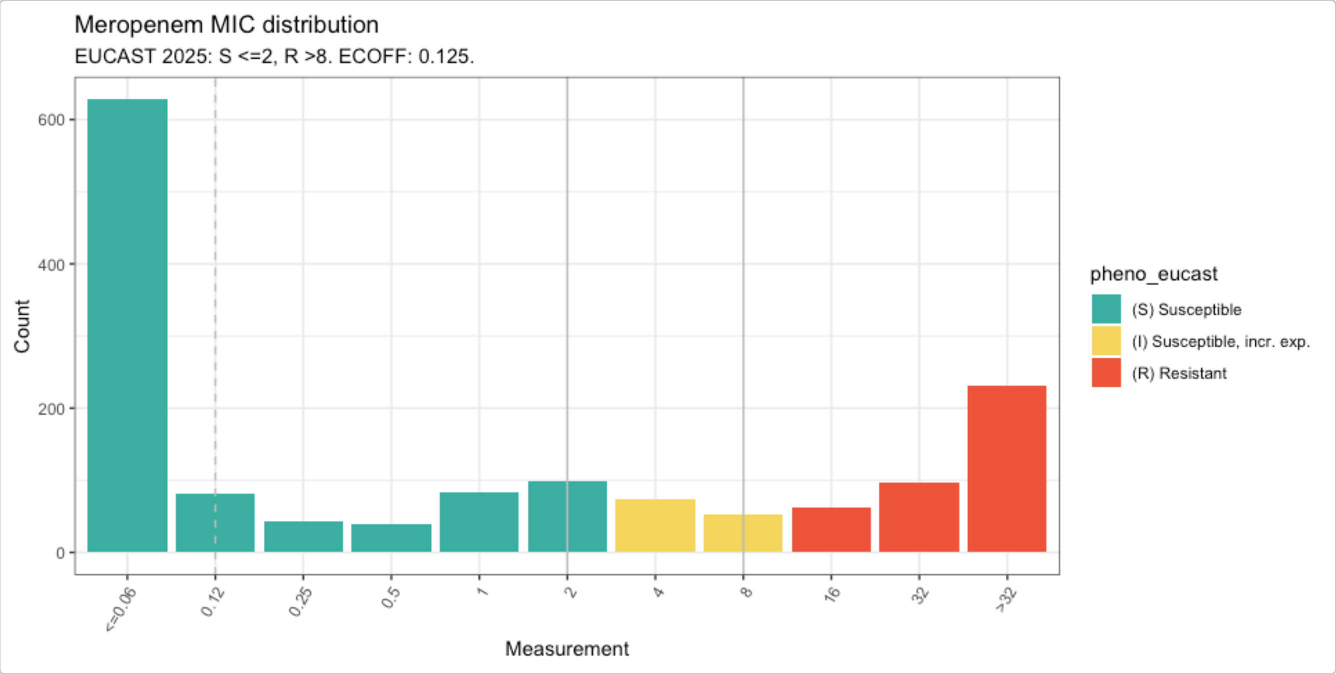


```
# Summary of meropenem phenotypes using ECOFF
kp_mero_euscape %>% count(ecoff)
#> # A tibble: 2 × 2
#>   ecoff     n
#>   <sir> <int>
#> 1 WT      710
#> 2 NWT     780

# MIC distribution coloured by ECOFF
assay_by_var(
  pheno_table = kp_mero_euscape,
  pheno_drug = "Meropenem",
  measure = "mic",
  colour_by = "ecoff",
  species = "Klebsiella pneumoniae"
)
#>   MIC breakpoints determined using AMR package: S <= 2 and R > 8
#>   NOTE: Multiple breakpoint entries, for different sites: Non-meningitis; Meningitis.
        Using the one with the highest S breakpoint (Non-meningitis).
```

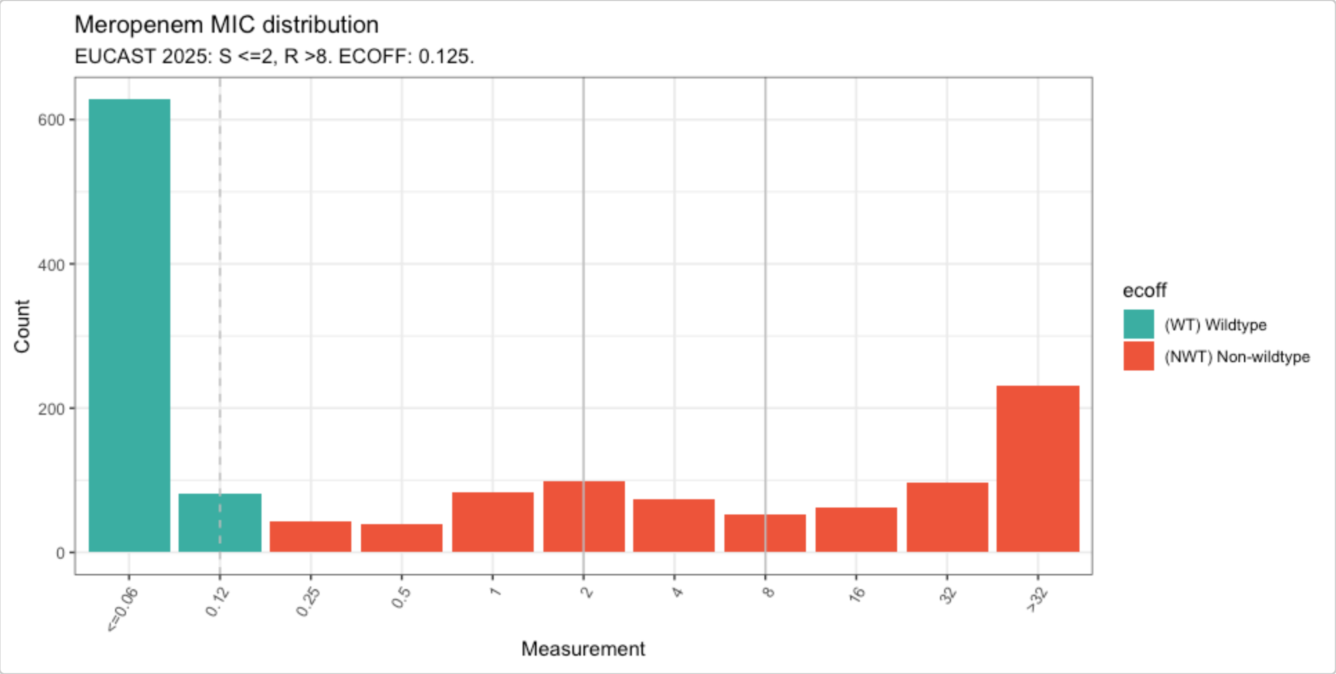


### Genotypes from Kleborate

Kleborate screens *Klebsiella pneumoniae* species complex (KpSC) genome assemblies to identify sequence types (MLST), species, antimicrobial resistance (AMR) genes, virulence loci (e.g., yersiniabactin, aerobactin), and capsule/LPS serotypes (K and O antigens), published here. It can be run on the command line or via Pathogenwatch.

#### Import Kleborate Genotype Data

The `import_kleborate()` function imports the output table from Kleborate, extracts the AMR genotyping data, and formats it to be used with AMRgen functions.

Mutation notation in Kleborate changed after v3.1.3 to adhere to HGVS Nomenclature, so:

- To import Kleborate output <=v3.1.3 (using informal nomenclature (e.g. [gene]-[mutation], [gene]-X%, OmpK36GD)), in the `import_kleborate()` function, set `hgvs = FALSE`.
- To import Kleborate output >v3.1.3 (using HGVS Nomenclature), in the `import_kleborate()` function, set to `hgvs=TRUE` (which is already the default option).

We are importing the latest version of Kleborate which is in the development branch - commit #4ec1dcb from March 17, 2026. This version uses HGVS Nomenclature for describing mutations and includes an updated AMR database compared to the most recent Kleborate release v3.2.4.

A table of Kleborate results generated for the EuSCAPE genomes is available in the AMRgen package as `kleborate_raw`. Let's import this to AMRgen genotype table format and summarise the content:

```
# Updated Kleborate results from the development branch as of March 17, 2026 (commit
        #4ec1dcb)
head(kleborate_raw, n = 10)
#> # A tibble: 10 × 122
#>    strain     species species_match contig_count    N50 largest_contig total_size
#>    <chr>      <chr>   <chr>                <dbl>  <dbl>          <dbl>      <dbl>
#>  1 SAMEA349… Klebsi… strong                 141 230759         470757    5578320
#>  2 SAMEA349… Klebsi… strong                  88 370309         938079    5384685
#>  3 SAMEA349… Klebsi… strong                  90 238750         529125    5446454
#>  4 SAMEA349… Klebsi… strong                 144 207582         663698    5574298
```

```
#>  5 SAMEA349… Klebsi… strong                  142 263498         678692    5486238
#>  6 SAMEA349… Klebsi… strong                   79 285199         991412    5529803
#>  7 SAMEA349… Klebsi… strong                  280 178980         585359    5817055
#>  8 SAMEA349… Klebsi… strong                  108 209418         517450    5379124
#>  9 SAMEA349… Klebsi… strong                  134 371444         984005    5558705
#> 10 SAMEA349… Klebsi… strong                  142 197944         636773    5497421
#> # ℹ 115 more variables: GC_content <dbl>, ambiguous_bases <chr>,
#> #   QC_warnings <chr>, ST <chr>, gapA <dbl>, infB <dbl>, mdh <dbl>, pgi <dbl>,
#> #   phoE <dbl>, rpoB <dbl>, tonB <dbl>, YbST <chr>, Yersiniabactin <chr>,
#> #   ybtS <chr>, ybtX <chr>, ybtQ <chr>, ybtP <chr>, ybtA <chr>, irp2 <chr>,
#> #   irp1 <chr>, ybtU <chr>, ybtT <chr>, ybtE <chr>, fyuA <chr>,
#> #   spurious_ybt_hits <chr>, CbST <chr>, Colibactin <chr>, clbA <chr>,
#> #   clbB <chr>, clbC <chr>, clbD <chr>, clbE <chr>, clbF <chr>, clbG <chr>, …

# Import Kleborate
kleborate_dev <- import_kleborate(kleborate_raw)

# View summary of genotypes
summarise_geno(kleborate_dev)
#> $uniques
#> # A tibble: 6 × 2
#>   column         n_unique
#>   <chr>             <int>
#> 1 id                 1490
#> 2 marker              470
#> 3 drug                  1
#> 4 drug_class           13
#> 5 gene                293
#> 6 variation type        4
#>
#> $per_type
#> # A tibble: 4 × 6
#>   `variation type`                  id marker  drug drug_class  gene
#>   <chr>                          <int>  <int> <int>      <int> <int>
#> 1 Gene presence detected          1490    288     1         13   288
#> 2 Inactivating mutation detected   569    167     1          2     3
#> 3 Nucleotide variant detected       15      1     1          1     1
#> 4 Protein variant detected         874     14     1          2     3
#>
#> $drugs
#> # A tibble: 13 × 5
#>    drug  drug_class                markers samples  hits
#>    <lgl> <chr>                       <int>   <int> <int>
#>  1 NA    Aminoglycosides                68    1060  2709
#>  2 NA    Beta-lactams                   76    1457  2833
#>  3 NA    Carbapenems                   153     770  1477
#>  4 NA    Cephalosporins (3rd gen.)      18     648   668
#>  5 NA    Macrolides                     15     460   924
#>  6 NA    Phenicols                      16     479   572
#>  7 NA    Phosphonics                    17    1485  1490
#>  8 NA    Polymyxins                     33     138   138
#>  9 NA    Quinolones                     26    1021  2665
#> 10 NA    Rifamycins                      3     119   128
#> 11 NA    Sulfonamides                   13     917  1096
#> 12 NA    Tetracyclines                  12     514   554
```

```
#> 13 NA    Trimethoprims                   21     941  1110
#> 
#> $markers
#> # A tibble: 471 × 5
#>    marker   drug  drug_class                `variation type`          n
#>    <chr>    <lgl> <chr>                     <chr>                 <int>
#>  1 ACC-4.v1 NA    Beta-lactams              Gene presence detected     2
#>  2 CMY-13   NA    Beta-lactams              Gene presence detected     1
#>  3 CMY-16   NA    Beta-lactams              Gene presence detected    31
#>  4 CMY-2.v2 NA    Beta-lactams              Gene presence detected     1
#>  5 CMY-30   NA    Cephalosporins (3rd gen.) Gene presence detected     1
#>  6 CMY-4.v1 NA    Beta-lactams              Gene presence detected     5
#>  7 CMY-6    NA    Beta-lactams              Gene presence detected     2
#>  8 CTX-M-1  NA    Cephalosporins (3rd gen.) Gene presence detected     1
#>  9 CTX-M-14 NA    Cephalosporins (3rd gen.) Gene presence detected    16
#> 10 CTX-M-15 NA    Cephalosporins (3rd gen.) Gene presence detected   568
#> # ℹ 461 more rows
```

#### Kleborate Genotype and Phenotype Summary

Summarize how many markers are associated with the beta-lactam and carbapenem drug class since Kleborate only operates at a drug class level.

```
summarise_geno_pheno(kleborate_dev, kp_mero_euscape,
  pheno_cols = c("pheno_eucast", "ecoff")
)
#> $overlapping_samples
#> [1] 1490
#> 
#> $drugs_with_pheno
#> # A tibble: 2 × 6
#>   drug                n drug_class   drug_name spp_pheno               mic
#>   <ab>            <int> <chr>        <chr>     <chr>                 <int>
#> 1 MEM: Meropenem   1490 Carbapenems  Meropenem Klebsiella pneumoniae  1490
#> 2 MEM: Meropenem   1490 Beta-lactams Meropenem Klebsiella pneumoniae  1490
#> 
#> $geno_hits
#> # A tibble: 2 × 6
#>   drug  drug_name drug_class   markers samples  hits
#>   <lgl> <chr>     <chr>          <int>   <int> <int>
#> 1 NA    <NA>      Beta-lactams      76    1457  2833
#> 2 NA    <NA>      Carbapenems      153     770  1477
#> 
#> $geno_markers
#> # A tibble: 229 × 6
#>    marker   drug  drug_name drug_class   `variation type`           n
#>    <chr>    <lgl> <chr>     <chr>        <chr>                  <int>
#>  1 ACC-4.v1 NA    <NA>      Beta-lactams Gene presence detected     2
#>  2 CMY-13   NA    <NA>      Beta-lactams Gene presence detected     1
#>  3 CMY-16   NA    <NA>      Beta-lactams Gene presence detected    31
#>  4 CMY-2.v2 NA    <NA>      Beta-lactams Gene presence detected     1
#>  5 CMY-4.v1 NA    <NA>      Beta-lactams Gene presence detected     5
#>  6 CMY-6    NA    <NA>      Beta-lactams Gene presence detected     2
```

```
#>   7 CTX-M-33 NA     <NA>      Carbapenems  Gene presence detected     1
#>   8 DHA-1    NA     <NA>      Beta-lactams Gene presence detected    78
#>   9 IMP-1    NA     <NA>      Carbapenems  Gene presence detected     3
#>  10 KPC-12   NA     <NA>      Carbapenems  Gene presence detected     1
#> # ℹ 219 more rows
#>
#> $pheno_counts_list
#> $pheno_counts_list$ecoff
#> # A tibble: 1 × 5
#>   drug           drug_name spp_pheno                WT   NWT
#>   <ab>           <chr>     <chr>                 <int> <int>
#> 1 MEM: Meropenem Meropenem Klebsiella pneumoniae   710   780
#>
#> $pheno_counts_list$pheno_eucast
#> # A tibble: 1 × 6
#>   drug           drug_name spp_pheno                 S     I     R
#>   <ab>           <chr>     <chr>                 <int> <int> <int>
#> 1 MEM: Meropenem Meropenem Klebsiella pneumoniae   973   126   391
```

### Phenotypes vs Kleborate Genotypes

#### Generate Binary Matrix for Kleborate AMR Markers

Most AMRgen analysis functions require a binary matrix with one sample per row, and columns indicating the phenotype and genotype data in columns, where a `1` indicates the presence and `0` indicates the absence of the phenotype or genotypic marker in that sample. This is produced using the `get_binary_matrix` function:

```
kleborate_binary_matrix <- get_binary_matrix(
  geno_table = kleborate_dev,
  pheno_table = kp_mero_euscape,
  pheno_drug = "Meropenem",
  geno_class = c("Carbapenems"),
  sir_col = "pheno_eucast",
  keep_assay_values = TRUE,
  keep_assay_values_from = "mic",
  marker_col = "marker.label"
)
#>  Defining NWT in binary matrix using ecoff column provided: ecoff
```

```
head(kleborate_binary_matrix, n = 10)
#> # A tibble: 10 × 24
#>     id    pheno ecoff    mic     R   NWT `OmpK36..-` `OmpK35..-` `NDM-1` `OXA-48`
#>     <chr> <sir> <sir>  <mic> <dbl> <dbl>       <dbl>       <dbl>   <dbl>    <dbl>
#>   1 SAME… I     NWT     4.00     0     1           1           0       0        0
#>   2 SAME… S     WT    <=0.06     0     0           0           0       0        0
#>   3 SAME… S     NWT     1.00     0     1           1           1       0        0
#>   4 SAME… I     NWT     4.00     0     1           1           0       0        0
#>   5 SAME… I     NWT     4.00     0     1           1           1       0        0
#>   6 SAME… S     WT    <=0.06     0     0           0           0       0        0
#>   7 SAME… S     WT    <=0.06     0     0           0           0       0        0
#>   8 SAME… S     NWT     2.00     0     1           1           0       0        0
```

```
#>  9 SAME… S      WT     <=0.06     0     0           0           0       0        0
#> 10 SAME… S      NWT     0.50     0     1           1           0       0        0
#> # ℹ 14 more variables: `OmpK36..c.25C>T` <dbl>, `KPC-3` <dbl>,
#> #   OmpK36..p.134_135insGD <dbl>, `KPC-2` <dbl>, OmpK36..p.135_136insD <dbl>,
#> #   `OXA-204` <dbl>, `VIM-1` <dbl>, OmpK36..p.136_137insTD <dbl>,
#> #   `VIM-4` <dbl>, `KPC-12` <dbl>, `OXA-232` <dbl>, `CTX-M-33` <dbl>,
#> #   `OXA-162` <dbl>, `IMP-1` <dbl>
```

#### Solo PPV Analysis for Kleborate AMR Markers

To understand the individual contribution of an AMR marker found “solo” (i.e., in the absence of another carbapenem resistance determinant), we use the `solo_ppv()` function. The `combined_plot` is a visual representation of each AMR marker found solo, the phenotypic distribution of isolates, and the positive predictive values (PPVs). The `solo_stats` table provides the PPVs, standard error (`se`), lower confidence interval (`ci.lower`), and upper confidence interval (`ci.upper`).

```
soloPPV_kleborate_mero <- solo_ppv(binary_matrix = kleborate_binary_matrix)
```

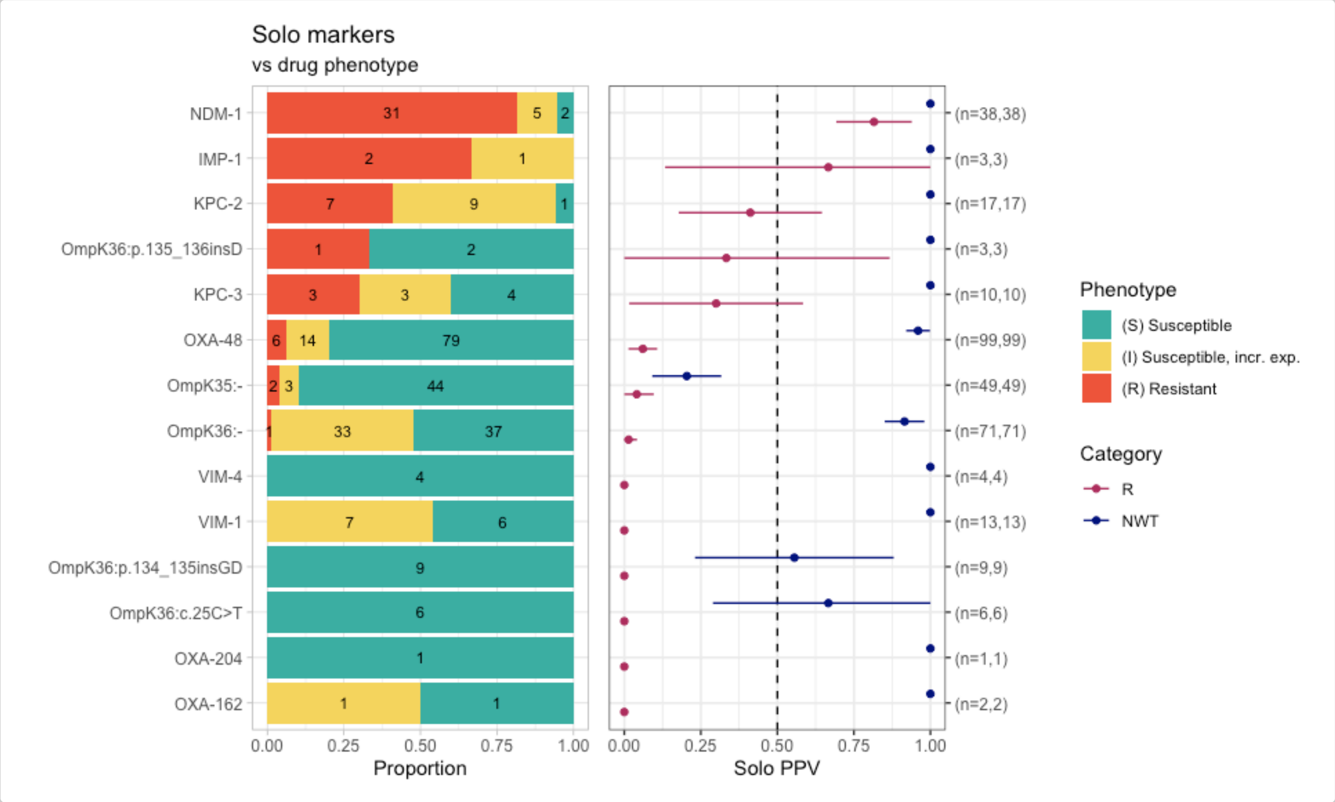


```
soloPPV_kleborate_mero$solo_stats
#> # A tibble: 28 × 8
#>    marker                category     x     n    ppv      se ci.lower ci.upper
#>    <chr>                 <chr>    <dbl> <int>  <dbl>   <dbl>    <dbl>    <dbl>
#>  1 OXA-162               R            0     2 0       0           0        0
#>  2 OXA-204               R            0     1 0       0           0        0
#>  3 OmpK36:c.25C>T        R            0     6 0       0           0        0
#>  4 OmpK36:p.134_135insGD R            0     9 0       0           0        0
#>  5 VIM-1                 R            0    13 0       0           0        0
#>  6 VIM-4                 R            0     4 0       0           0        0
#>  7 OmpK36:-              R            1    71 0.0141 0.0140      0        0.0415
#>  8 OmpK35:-              R            2    49 0.0408 0.0283      0        0.0962
```

```
#>  9 OXA-48                 R             6     99 0.0606 0.0240    0.0136    0.108
#> 10 KPC-3                  R             3     10 0.3    0.145     0.0160    0.584
#> # ℹ 18 more rows
```

Here we can see that the only genotypes whose presence alone, in the absence of any other markers, confers resistance are the carbapenemase genes NDM-1 and IMP-1. Porin mutations alone are not associated with resistance.

#### Combinatorial PPV Analysis for Kleborate AMR Markers

To understand the contribution of AMR markers found in combination with one another, we use the `amr_ppv()` function. The `plot` is a visual summary of each AMR marker combination observed in an UpSet plot format, including phenotypic distribution and PPVs for each combination. The `summary` table includes each AMR marker combination observed, including number of resistant isolates, positive predictive values, and median assay values (and interquartile range) where relevant.

```
comboPPV_kleborate_mero <- amr_ppv(
  binary_matrix = kleborate_binary_matrix,
  order = "value",
  min_set_size = 1,
  pheno_drug = "Meropenem",
  upset_grid = TRUE,
  plot_assay = TRUE,
  assay = "mic"
)
#> Ordering markers by frequency
#> Scale for y is already present.
#> Adding another scale for y, which will replace the existing scale.
```

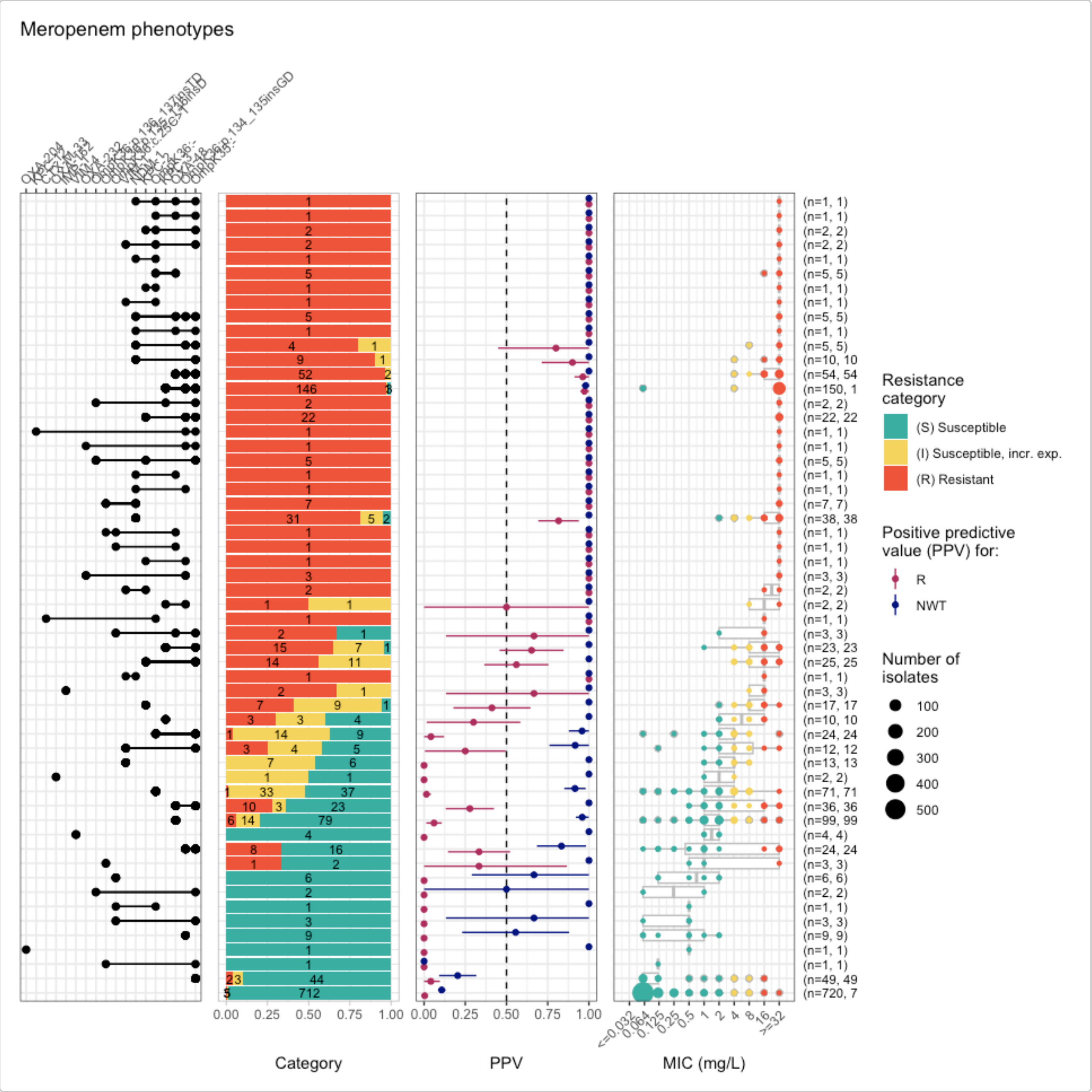


```
comboPPV_kleborate_mero$summary
#> # A tibble: 56 × 21
#>    marker_list        marker_count     n combination_id   R.n    R.ppv R.ci_lower
#>    <chr>                     <dbl> <int> <fct>          <dbl>    <dbl>      <dbl>
#>  1 ""                            0   720 0_0_0_0_0_0_0…     3  0.00417          0
#>  2 "IMP-1"                       1     3 0_0_0_0_0_0_0…     2  0.667        0.133
#>  3 "OXA-162"                     1     2 0_0_0_0_0_0_0…     0  0                0
#>  4 "VIM-4"                       1     4 0_0_0_0_0_0_0…     0  0                0
#>  5 "VIM-1"                       1    13 0_0_0_0_0_0_0…     0  0                0
#>  6 "OXA-204"                     1     1 0_0_0_0_0_0_0…     0  0                0
#>  7 "OmpK36:p.135_136…            1     3 0_0_0_0_0_0_0…     1  0.333            0
#>  8 "KPC-2"                       1    17 0_0_0_0_0_0_0…     7  0.412        0.178
#>  9 "KPC-2, VIM-1"                2     2 0_0_0_0_0_0_0…     2  1                1
#> 10 "OmpK36:p.134_135…            1     9 0_0_0_0_0_0_1…     0  0                0
#> # ℹ 46 more rows
#> # ℹ 14 more variables: R.ci_upper <dbl>, R.denom <int>, NWT.n <dbl>,
#> #   NWT.ppv <dbl>, NWT.ci_lower <dbl>, NWT.ci_upper <dbl>, NWT.denom <int>,
#> #   median_excludeRangeValues <dbl>, q25_excludeRangeValues <dbl>,
```

```
#> #   q75_excludeRangeValues <dbl>, n_excludeRangeValues <int>,
#> #   median_ignoreRanges <dbl>, q25_ignoreRanges <dbl>, q75_ignoreRanges <dbl>
```

#### UpSet Plot for Kleborate AMR Markers

Similar to the previous `amr_ppv()` function, the `amr_upset()` function will generate a `summary` table and `plot` that shows the combinations of AMR markers found in the isolates and their phenotypic distribution. The UpSet `plot` produced by `amr_upset()` is similar to that generated using `amr_ppv()` function, but oriented vertically and without the PPV panel. Restricting the plot to marker combinations observed at least 3 times in the dataset (`min_set_size=3`) makes it a little easier to see what's going on.

```
kp_mic_upset_kleborate <- amr_upset(kleborate_binary_matrix,
  assay = "mic", species = "Klebsiella pneumoniae", min_set_size = 3
)
#> Ordering markers by frequency
#> Scale for y is already present.
#> Adding another scale for y, which will replace the existing scale.
#> Scale for y is already present.
#> Adding another scale for y, which will replace the existing scale.
```

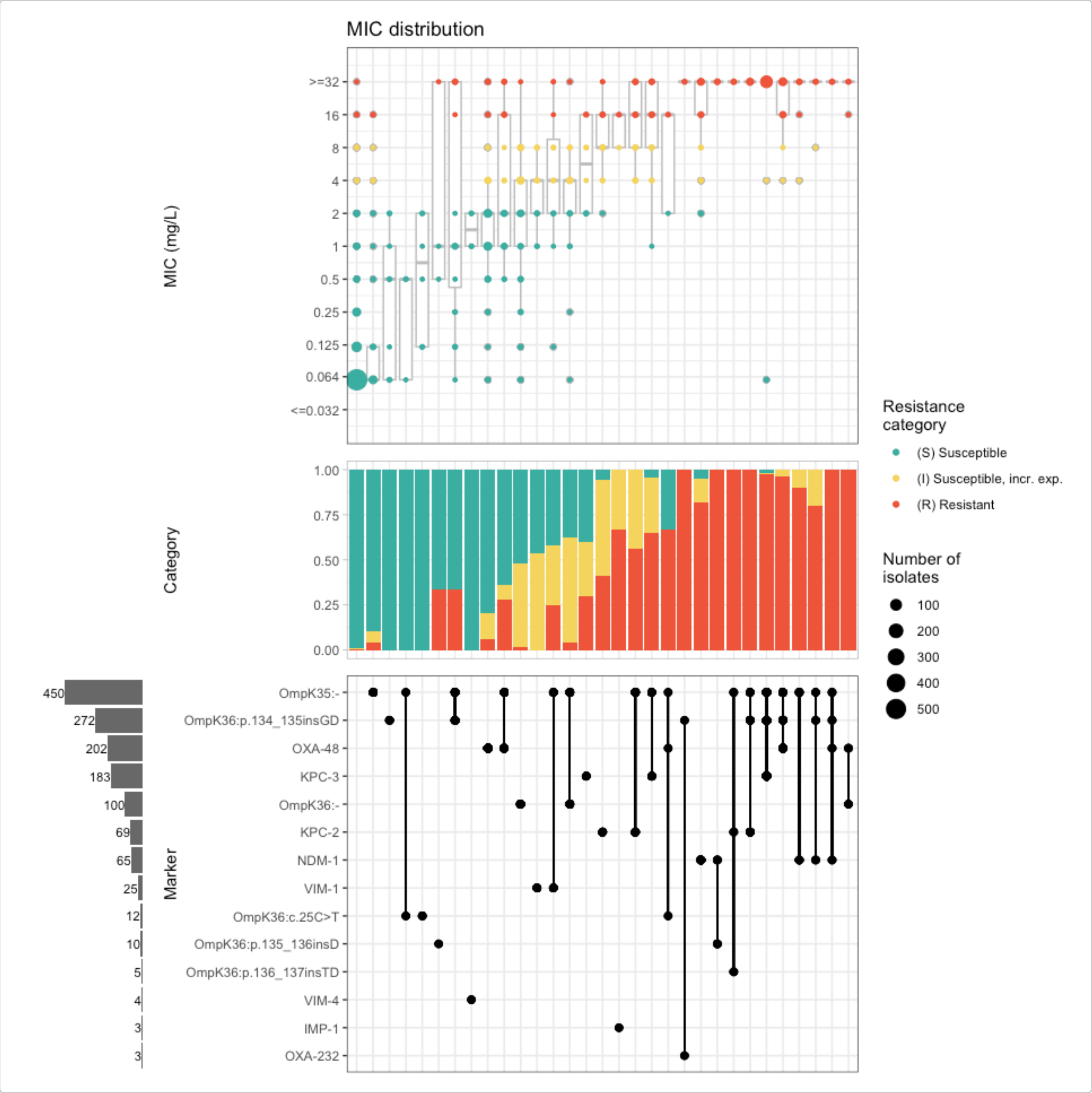


It’s still quite complex to follow the plot and spot patterns, as we have carbapenemase genes mixed in with mutations, making it difficult to compare the effect of a given gene with and without a mutation (and vice versa).

To simplify things, let’s group the various insertion mutations together.

```
# modify marker labels in the genotype table
kleborate_dev_plotting <- kleborate_dev %>%
  mutate(marker.label = case_when(
    marker.label == "OmpK36:-" ~ "OmpK36Δ",
    marker.label == "OmpK35:-" ~ "OmpK35Δ",
    grepl("OmpK36:p", marker.label) ~ "OmpK36ins",
    TRUE ~ marker.label
  ))

# calculate binary matrix from this updated genotype table, within the amr_upset function
kp_mic_upset_kleborate2 <- amr_upset(
  geno_table = kleborate_dev_plotting,
  pheno_table = kp_mero_euscape,
```

```r
  pheno_drug = "Meropenem",
  geno_class = c("Carbapenems"),
  sir_col = "pheno_eucast",
  marker_col = "marker.label",
  assay = "mic",
  species = "Klebsiella pneumoniae",
  min_set_size = 3
)
#> Generating geno-pheno binary matrix
#>  Defining NWT in binary matrix using ecoff column provided: ecoff
#> Ordering markers by frequency
#>   MIC breakpoints determined using AMR package: S <= 2 and R > 8
#>   NOTE: Multiple breakpoint entries, for different sites: Non-meningitis; Meningitis.
       Using the one with the highest S breakpoint (Non-meningitis).
#>   MIC breakpoints determined using AMR package: S <= 2 and R > 8
#>   NOTE: Multiple breakpoint entries, for different sites: Non-meningitis; Meningitis.
       Using the one with the highest S breakpoint (Non-meningitis).
#> Scale for y is already present.
#> Adding another scale for y, which will replace the existing scale.
#> Scale for y is already present.
#> Adding another scale for y, which will replace the existing scale.
```

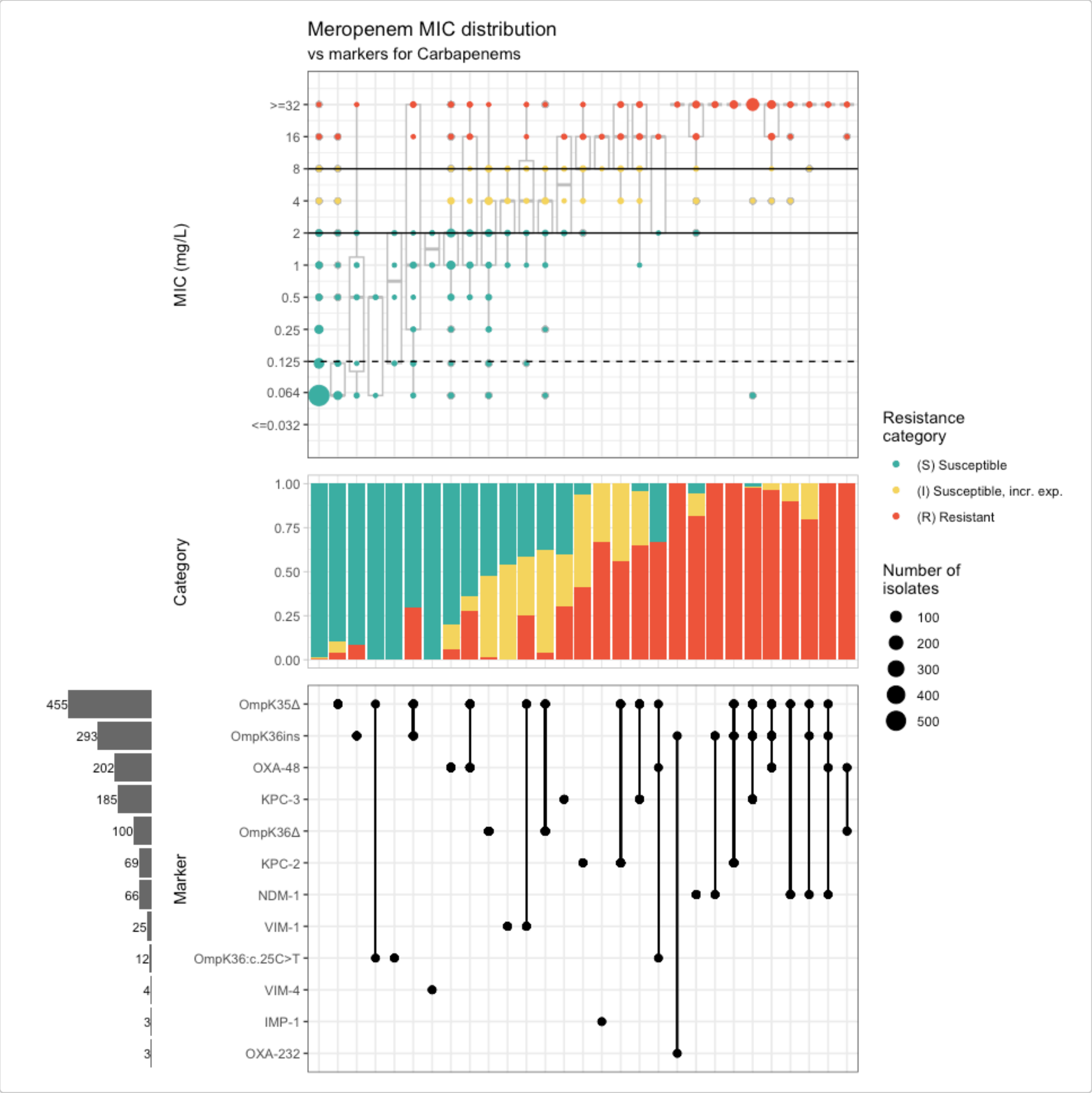


#### MIC boxplot for Kleborate enzymes vs porin mutations

Now let's use the `assay_by_var()` function to explore the MIC distribution stratified by enzyme, and by mutation. Setting `boxplot=T` we can view boxplots of MIC, grouped and coloured by mutation (setting `colour_by="Omp_mutations"`), and faceted to one plotting panel per carbapenemase gene (setting `facet_by = "Bla_Carb_acquired"`). This also returns summary stats (median, interquartile range for MIC) stratified by gene and mutation. By specifying `species="Klebsiella pneumoniae'`, we can also retrieve the clinical breakpoints and ECOFF for meropenem and add these to the plot.

```
# pivot genotype table to wide format (one row per sample) with separate columns for each
        class of marker, and add the MIC values for each sample
kleborate_dev_wide_mic <- kleborate_dev_plotting %>%
  filter(marker.label != "OmpK36:c.25C>T") %>% # filter out synonymous SNP
  select(id, Kleborate_Class, marker.label) %>%
  pivot_wider(
    id_cols = id,
    names_from = Kleborate_Class,
```

```
    values_from = marker.label,
    values_fn = ~ paste(.x, collapse = ","),
    values_fill = "-"
  ) %>%
  left_join(kp_mero_euscape)
#> Joining with `by = join_by(id)`

head(kleborate_dev_wide_mic)
#> # A tibble: 6 × 61
#>   id           AGly_acquired Fcyn_acquired Flq_acquired Phe_acquired Sul_acquired
#>   <chr>        <chr>         <chr>         <chr>        <chr>        <chr>
#> 1 SAMEA34989… aac(3)-IIa.v… fosA10*?      qnrB4,aac(6… catA1        sul1
#> 2 SAMEA34989… -             fosA5*        -            -            -
#> 3 SAMEA34989… -             fosA10*?      -            catA1        sul2
#> 4 SAMEA34989… aac(3)-IIa.v… fosA10*?      qnrB4,aac(6… catA1        sul1
#> 5 SAMEA34989… aac(3)-IIa.v… fosA5         -            -            -
#> 6 SAMEA34989… -             fosA5         -            -            -
#> # ℹ 55 more variables: Tmt_acquired <chr>, Bla_acquired <chr>,
#> #   Bla_ESBL_acquired <chr>, Bla_chr <chr>, Omp_mutations <chr>,
#> #   Flq_mutations <chr>, MLS_acquired <chr>, Tet_acquired <chr>,
#> #   Col_mutations <chr>, Rif_acquired <chr>, Bla_Carb_acquired <chr>,
#> #   Col_acquired <chr>, Bla_inhR_acquired <chr>, drug <ab>, mic <mic>,
#> #   disk <dsk>, pheno_provided <sir>, pheno_eucast <sir>, ecoff <sir>,
#> #   guideline <chr>, method <chr>, platform <chr>, source <chr>, …

# this table is now ready to use with assay_by_var, to flexibly explore MIC distribution
        by genotype
kleborate_mic_by_gene_mutation <- assay_by_var(kleborate_dev_wide_mic,
  pheno_drug = "Meropenem", colour_by = "Omp_mutations",
  facet_by = "Bla_Carb_acquired", species = "Klebsiella pneumoniae",
  colour_legend_label = "Porin status", boxplot = T, measure_axis_label = "MIC (mg/L)"
)
#>   MIC breakpoints determined using AMR package: S <= 2 and R > 8
#>   NOTE: Multiple breakpoint entries, for different sites: Non-meningitis; Meningitis.
        Using the one with the highest S breakpoint (Non-meningitis).

kleborate_mic_by_gene_mutation$plot
```

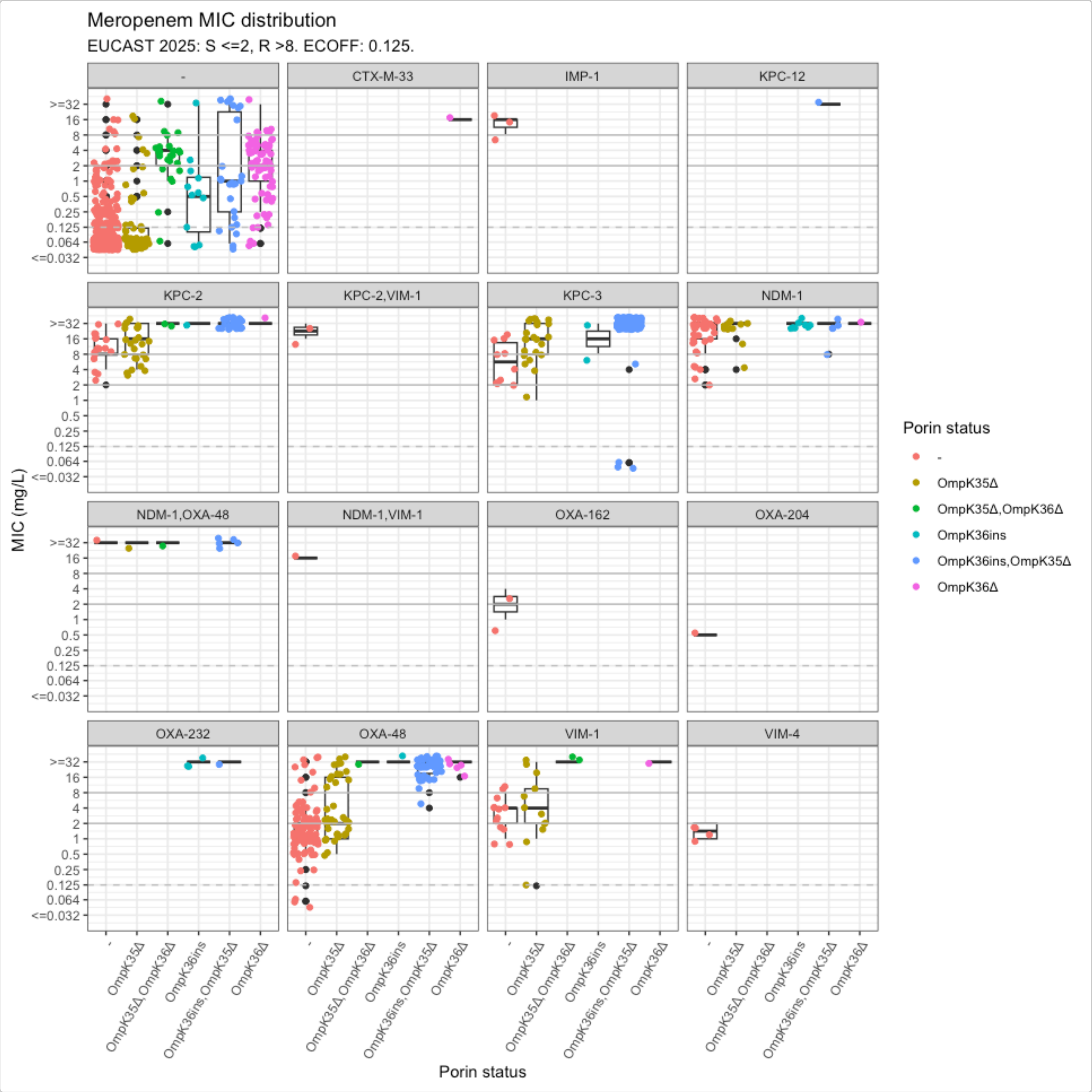


The plot is crowded with some carbapenemase/combinations that are very rare, let's collapse into enzyme families, exclude isolates that have multiple carbapenemases (n=11), and remove the single CTX gene.

```
kleborate_dev_wide_mic_trim <- kleborate_dev_wide_mic %>%
  filter(!grepl(",", Bla_Carb_acquired)) %>% # exclude isolates with multiple
      carbapenemases
  mutate(Bla_Carb_acquired = substr(Bla_Carb_acquired, 1, 3)) %>% # first 3 letters give
      gene family name
  filter(Bla_Carb_acquired != "CTX")

kleborate_mic_by_gene_mutation <- assay_by_var(kleborate_dev_wide_mic_trim,
  pheno_drug = "Meropenem", colour_by = "Omp_mutations",
  facet_by = "Bla_Carb_acquired", species = "Klebsiella pneumoniae",
  colour_legend_label = "Porin status",
  boxplot = T, measure_axis_label = "MIC (mg/L)"
)
#>   MIC breakpoints determined using AMR package: S <= 2 and R > 8
#>   NOTE: Multiple breakpoint entries, for different sites: Non-meningitis; Meningitis.
      Using the one with the highest S breakpoint (Non-meningitis).
```

```
kleborate_mic_by_gene_mutation$plot
```

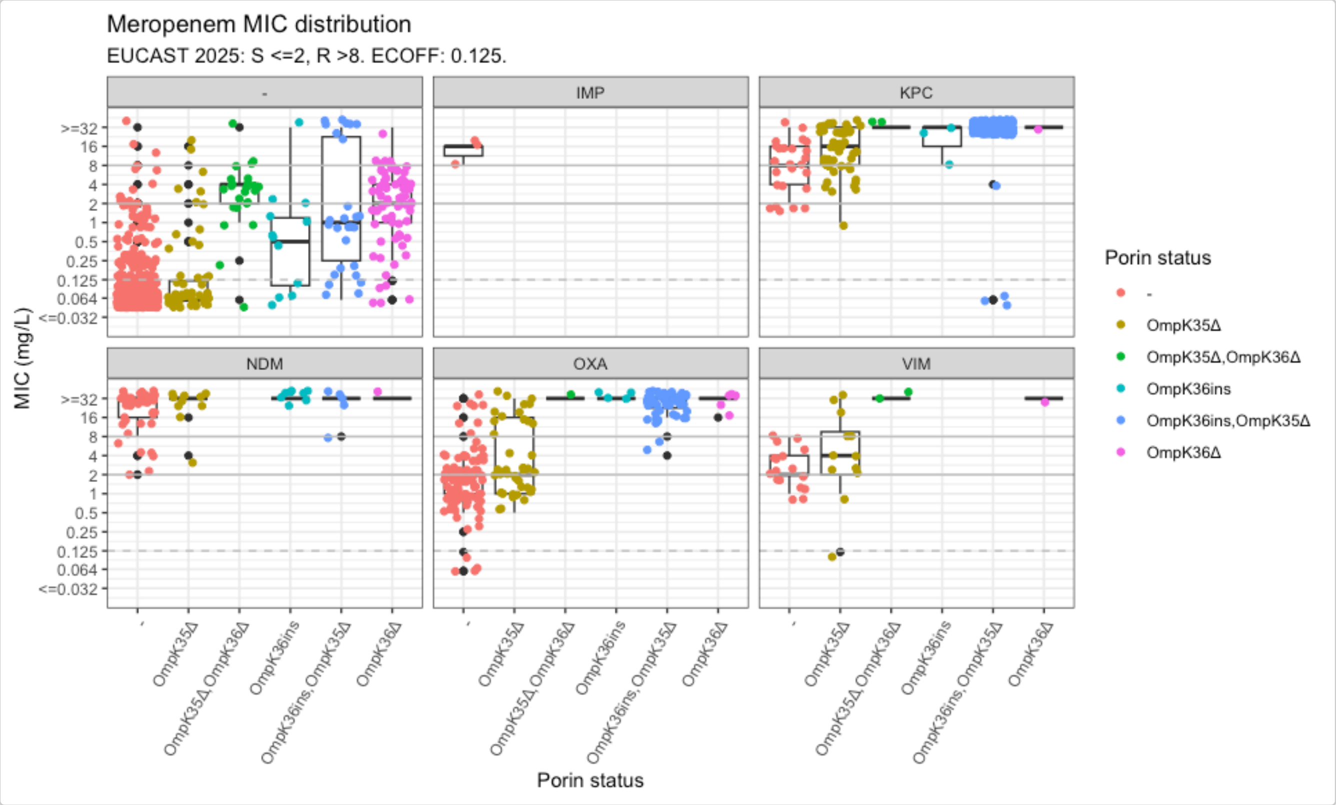


Now we can see the impact of the mutations in the absence of any carbapenemase gene (panel labelled "-"); the MIC distribution associated with each carbapenemase in the absence of any mutation (red points in all other plots); and the impact of mutations in the presence of each type of carbapenemase (other colours).

```
kleborate_mic_by_gene_mutation$stats %>% head(19)
#> # A tibble: 19 × 7
#>    Omp_mutations     Bla_Carb_acquired      n median geom_mean   q25   q75
#>    <chr>             <chr>              <int>  <dbl>     <dbl> <dbl> <dbl>
#>  1 -                 -                    726   0.06    0.0842 0.06   0.06
#>  2 -                 IMP                    3  16      12.7   12     16
#>  3 -                 KPC                   27   8       7.80   4     16
#>  4 -                 NDM                   38  32      18.9   16     32
#>  5 -                 OXA                  103   2       1.56   1      2
#>  6 -                 VIM                   17   2       2.55   2      4
#>  7 OmpK35Δ           -                     52   0.06    0.150  0.06   0.12
#>  8 OmpK35Δ           KPC                   48  16      13.3    8     32
#>  9 OmpK35Δ           NDM                   10  32      24.3   32     32
#> 10 OmpK35Δ           OXA                   39   2       3.47   1     16
#> 11 OmpK35Δ           VIM                   12   4       3.99   2     10
#> 12 OmpK35Δ,OmpK36Δ -                     24   4       2.74   2      4
#> 13 OmpK35Δ,OmpK36Δ KPC                    2  32      32     32     32
#> 14 OmpK35Δ,OmpK36Δ OXA                    1  32      32     32     32
#> 15 OmpK35Δ,OmpK36Δ VIM                    2  32      32     32     32
#> 16 OmpK36ins         -                     12   0.5     0.523  0.105  1.25
#> 17 OmpK36ins         KPC                    3  32      20.2   20     32
#> 18 OmpK36ins         NDM                    8  32      32     32     32
#> 19 OmpK36ins         OXA                    4  32      32     32     32
```

Reformat the stats table into wide format to more clearly see the effects of porin vs. carbapenemase status on MIC. Median and geometric mean (in brackets) values are grouped by porin vs. carbapenemase status, each corresponding to unique boxplotted distribution.

```
kleborate_mic_by_gene_mutation_table <- kleborate_mic_by_gene_mutation$stats %>%
  mutate(geom_mean = round(geom_mean, 1)) %>%
  mutate(median_mean = paste0(median, " (", geom_mean, ")")) %>%
  mutate(OmpK35 = case_when(
    grepl("OmpK35", Omp_mutations) ~ "\u0394",
    TRUE ~ "-"
  )) %>%
  mutate(OmpK36 = case_when(
    grepl("OmpK36ins", Omp_mutations) ~ "Insertion",
    grepl("OmpK36\u0394", Omp_mutations) ~ "\u0394",
    TRUE ~ "-"
  )) %>%
  mutate(Bla_Carb_acquired = str_replace_all(Bla_Carb_acquired, "-", "None")) %>%
  select(OmpK35, OmpK36, Bla_Carb_acquired, median_mean)

median_MIC_table <- kleborate_mic_by_gene_mutation_table %>%
  pivot_wider(
    names_from = Bla_Carb_acquired,
    values_from = median_mean,
    values_fill = "-"
  )

median_MIC_table
#> # A tibble: 6 x 8
#>   OmpK35 OmpK36    None       IMP         KPC         NDM         OXA         VIM
#>   <chr>  <chr>     <chr>      <chr>       <chr>       <chr>       <chr>       <chr>
#> 1 -      -         0.06 (0.1) 16 (12.7)   8 (7.8)     32 (18.9)   2 (1.6)     2 (2.6)
#> 2 Δ      -         0.06 (0.1) -           16 (13.3)   32 (24.3)   2 (3.5)     4 (4)
#> 3 Δ      Δ         4 (2.7)    -           32 (32)     -           32 (32)     32 (32)
#> 4 -      Insertion 0.5 (0.5)  -           32 (20.2)   32 (32)     32 (32)     -
#> 5 Δ      Insertion 1 (1.5)    -           32 (28.5)   32 (24.3)   32 (25.8)   -
#> 6 -      Δ         2 (1.8)    -           32 (32)     32 (32)     32 (27.9)   32 (32)
```

Making the table aesthetically pleasing using the gt package (to reproduce Figure 5c in the AMRgen paper).

```
# If you have the gt package, you can use it to make the table aesthetically pleasing
median_MIC_table_aes <- median_MIC_table %>%
  gt() %>%
  cols_label(
    OmpK35 = html("<b>OmpK35</b>"),
    OmpK36 = html("<b>OmpK36</b>"),
    None = html("<b>None</b>"),
    IMP = html("<b>IMP</b>"),
    KPC = html("<b>KPC</b>"),
    NDM = html("<b>NDM</b>"),
    OXA = html("<b>OXA</b>"),
    VIM = html("<b>VIM</b>"),
  ) %>%
  tab_spanner(
```

```r
  label = html("<b>Porin status</b>"),
  columns = c("OmpK35", "OmpK36")
) %>%
tab_spanner(
  label = html("<b>Carbapenemase status</b>"),
  columns = c("None", "KPC", "NDM", "OXA", "VIM", "IMP")
) %>%
fmt_missing(
  columns = everything(),
  missing_text = "-"
) %>%
# Background color based on EUCAST breakpoints
data_color(
  columns = c("None", "KPC", "NDM", "OXA", "VIM", "IMP"),
  fn = function(x) {
    numeric_vals <- as.numeric(str_extract(x, "^[0-9.]+"))

    case_when(
      is.na(numeric_vals) ~ "transparent",
      numeric_vals <= 2 ~ "#3CAEA3",
      numeric_vals <= 8 ~ "#F6D55C",
      numeric_vals > 8 ~ "#ED553B"
    )
  }
) %>%
# Changing orange cells to have white font so it is easier to see the numbers
data_color(
  columns = c("None", "KPC", "NDM", "OXA", "VIM", "IMP"),
  fn = function(x) {
    numeric_vals <- as.numeric(str_extract(x, "^[0-9.]+"))

    case_when(
      is.na(numeric_vals) ~ "black",
      numeric_vals > 8 ~ "white",
      TRUE ~ "black"
    )
  },
  apply_to = "text"
) %>%
# aligning columns
cols_align(
  align = "center",
  columns = everything()
) %>%
# text options
tab_options(
  table.font.names = "Arial",
  table.font.size = 14,
  heading.title.font.size = 16,
  table.border.top.color = "black",
  table.border.bottom.color = "black"
) %>%
# column widths
cols_width(
  OmpK35 ~ px(125),
```

```
    OmpK36 ~ px(125),
    everything() ~ px(100)
  )

# Adding title and legend for colour
median_MIC_table_aes <- median_MIC_table_aes %>%
  tab_header(
    title = html("<b>Median (Mean) Meropenem Minimum Inhibitory Concentration (mg/L)
        </b>"),
    subtitle = html(
      "<span style='font-size:12px;'>
      <b>EUCAST clinical breakpoint:</b>
      <span style='color:#3CAEA3;'>■</span> S (Susceptible, ≤ 2 mg/L)  
      <span style='color:#F6D55C;'>■</span> I (Susceptible, Increased exposure; 4-8 mg/L)
         
      <span style='color:#ED553B;'>■</span> R (Resistant, > 8 mg/L)
      </span>"
    )
  )

median_MIC_table_aes
```

### Genotypes from Kleborate v3.1.3

#### Import Kleborate (<= v3.1.3) Genotype Data

We will now compare the latest version of Kleborate (which we've been using up until this point) development branch; commit #4ec1dcb to an older version of Kleborate v3.1.3. The older versions (<=v3.1.3) of Kleborate uses informal nomenclature to describe mutations (e.g. [gene]-[mutation], [gene]-X%, OmpK36GD), whereas the updated version follows the HGVS nomenclature standard.

A table of Kleborate v3.1.3 results generated for the EuSCAPE genomes is available in the AMRgen package as `kleborate_raw_v313`. Let's import it and summarise its contents:

```
# View Kleborate output v3.1.3 (using informal nomenclature (e.g. [gene]-[mutation],
        [gene]-X%, OmpK36GD))
head(kleborate_raw_v313, n = 10)
#> # A tibble: 10 × 113
#>    strain    species species_match contig_count    N50 largest_contig total_size
#>    <chr>     <chr>   <chr>                <dbl>  <dbl>          <dbl>      <dbl>
#>  1 SAMEA349… Klebsi… strong                 141 230759         470757    5578320
#>  2 SAMEA349… Klebsi… strong                  88 370309         938079    5384685
#>  3 SAMEA349… Klebsi… strong                  90 238750         529125    5446454
#>  4 SAMEA349… Klebsi… strong                 144 207582         663698    5574298
#>  5 SAMEA349… Klebsi… strong                 142 263498         678692    5486238
#>  6 SAMEA349… Klebsi… strong                  79 285199         991412    5529803
#>  7 SAMEA349… Klebsi… strong                 280 178980         585359    5817055
#>  8 SAMEA349… Klebsi… strong                 108 209418         517450    5379124
#>  9 SAMEA349… Klebsi… strong                 134 371444         984005    5558705
#> 10 SAMEA349… Klebsi… strong                 142 197944         636773    5497421
#> # ℹ 106 more variables: ambiguous_bases <chr>, QC_warnings <chr>, ST <chr>,
#> #   gapA <dbl>, infB <dbl>, mdh <dbl>, pgi <dbl>, phoE <dbl>, rpoB <dbl>,
#> #   tonB <dbl>, YbST <chr>, Yersiniabactin <chr>, ybtS <chr>, ybtX <chr>,
#> #   ybtQ <chr>, ybtP <chr>, ybtA <chr>, irp2 <chr>, irp1 <chr>, ybtU <chr>,
```

```
#> #   ybtT <chr>, ybtE <chr>, fyuA <chr>, spurious_ybt_hits <chr>, CbST <chr>,
#> #   Colibactin <chr>, clbA <chr>, clbB <chr>, clbC <chr>, clbD <chr>,
#> #   clbE <chr>, clbF <chr>, clbG <chr>, clbH <chr>, clbI <chr>, clbL <chr>, …

# Use import_kleborate() function and set `hgvs = FALSE` for Kleborate outputs generated
        from <=v3.1.3 (using non-HGVS nomenclature)
kleborate_v313 <- import_kleborate(
  input_table = kleborate_raw_v313,
  sample_col = "strain",
  hgvs = FALSE
)

# Summarize genotype table
summarise_geno(kleborate_v313, sample_col = "id", marker_col = "marker.label")
#> $uniques
#> # A tibble: 6 × 2
#>   column         n_unique
#>   <chr>             <int>
#> 1 id                 1489
#> 2 marker.label        263
#> 3 drug                  1
#> 4 drug_class           13
#> 5 gene                263
#> 6 variation type        4
#>
#> $per_type
#> # A tibble: 4 × 6
#>   `variation type`                 id marker.label  drug drug_class  gene
#>   <chr>                         <int>        <int> <int>      <int> <int>
#> 1 Gene presence detected         1489          246     1         13   246
#> 2 Inactivating mutation detected  568            3     1          2     3
#> 3 Nucleotide variant detected      15            1     1          1     1
#> 4 Protein variant detected        874           13     1          2    13
#>
#> $drugs
#> # A tibble: 13 × 5
#>    drug  drug_class                markers samples  hits
#>    <lgl> <chr>                       <int>   <int> <int>
#>  1 NA    Aminoglycosides                59    1057  3140
#>  2 NA    Beta-lactams                   76    1457  2833
#>  3 NA    Carbapenems                    17     766  1459
#>  4 NA    Cephalosporins (3rd gen.)      18     648   668
#>  5 NA    Macrolides                     14     460   923
#>  6 NA    Phenicols                      12     469   558
#>  7 NA    Phosphonics                     1       3     3
#>  8 NA    Polymyxins                      3     138   138
#>  9 NA    Quinolones                     25    1020  2422
#> 10 NA    Rifamycins                      2     114   114
#> 11 NA    Sulfonamides                    9     913  1090
#> 12 NA    Tetracyclines                   9     503   541
#> 13 NA    Trimethoprims                  18     886  1017
#>
#> $markers
#> # A tibble: 263 × 5
#>    marker.label drug  drug_class                `variation type`           n
```

```
#>    <chr>        <lgl> <chr>                   <chr>                  <int>
#>  1 ACC-4        NA    Beta-lactams            Gene presence detected     2
#>  2 CMY-13       NA    Beta-lactams            Gene presence detected     1
#>  3 CMY-16       NA    Beta-lactams            Gene presence detected    31
#>  4 CMY-2.v2     NA    Beta-lactams            Gene presence detected     1
#>  5 CMY-30       NA    Cephalosporins (3rd gen.) Gene presence detected     1
#>  6 CMY-4.v1     NA    Beta-lactams            Gene presence detected     5
#>  7 CMY-6        NA    Beta-lactams            Gene presence detected     2
#>  8 CTX-M-1      NA    Cephalosporins (3rd gen.) Gene presence detected     1
#>  9 CTX-M-14     NA    Cephalosporins (3rd gen.) Gene presence detected    16
#> 10 CTX-M-15     NA    Cephalosporins (3rd gen.) Gene presence detected   568
#> # ℹ 253 more rows
```

### Compare Kleborate versions

Comparing only the carbapenem resistance determinants from an updated version Kleborate (development branch) to a previous version (v3.1.3).

```
# Grouping Kleborate development branch carbapenem resistance determinants, so that there
        is one row per sample
kleborate_dev_markers_grouped <- kleborate_dev %>%
  filter(Kleborate_Class == "Omp_mutations" | Kleborate_Class == "Bla_Carb_acquired") %>%
  select(id, marker.label) %>%
  rename(kleborate = marker.label) %>%
  group_by(id) %>%
  summarise(
    Kleborate_dev_markers = kleborate %>%
      sort() %>%
      str_c(collapse = ";")
  )

# Since we know that Kleborate v3.1.3 does not use HGVS notation, we will change the names
        of the OmpK36 mutations to match HGVS notation for comparison purposes
# After, we group them so there is one row per sample
kleborate_v313_markers_grouped <- kleborate_v313 %>%
  filter(Kleborate_Class == "Omp_mutations" | Kleborate_Class == "Bla_Carb_acquired") %>%
  select(id, marker.label) %>%
  mutate(
    marker.label = str_replace_all(marker.label, "OmpK36GD", "OmpK36:p.134_135insGD"),
    marker.label = str_replace_all(marker.label, "OmpK36_c25t", "OmpK36:c.25C>T"),
    marker.label = str_replace_all(marker.label, "OmpK36TD", "OmpK36:p.136_137insTD")
  ) %>%
  rename(kleborate = marker.label) %>%
  group_by(id) %>%
  summarise(
    Kleborate_v313_markers = kleborate %>%
      sort() %>%
      str_c(collapse = ";")
  )

# Joining Kleborate version tables for comparison
compare_kleborate_versions <- full_join(
  kleborate_dev_markers_grouped,
  kleborate_v313_markers_grouped
```

```
)
#> Joining with `by = join_by(id)`

# Comparing Kleborate versions and creating two new columns to show what each version is
        missing
compare_kleborate_versions <- compare_kleborate_versions %>%
  rowwise() %>%
  mutate(
    dev_missing = {
      v313_vec <- str_split(Kleborate_v313_markers, ";")[[1]]
      dev_vec <- str_split(Kleborate_dev_markers, ";")[[1]]
      missing <- setdiff(v313_vec, dev_vec)
      if (length(missing) == 0) NA_character_ else str_c(missing, collapse = ";")
    },
    v313_missing = {
      v313_vec <- str_split(Kleborate_v313_markers, ";")[[1]]
      dev_vec <- str_split(Kleborate_dev_markers, ";")[[1]]
      missing <- setdiff(dev_vec, v313_vec)
      if (length(missing) == 0) NA_character_ else str_c(missing, collapse = ";")
    }
  ) %>%
  ungroup()

# Table listing the AMR markers that are missing from Kleborate v3.1.3
compare_kleborate_versions %>% count(v313_missing, sort = TRUE)
#> # A tibble: 4 x 2
#>   v313_missing            n
#>   <chr>               <int>
#> 1 <NA>                  752
#> 2 OmpK36:p.135_136insD   12
#> 3 OmpK35:-                4
#> 4 OmpK36:-                2

# Table listing the AMR markers that are missing from the updated Kleborate version
compare_kleborate_versions %>% count(dev_missing, sort = TRUE)
#> # A tibble: 1 x 2
#>   dev_missing     n
#>   <chr>       <int>
#> 1 <NA>          770
```

We can see that there are no carbapenem resistance determinants missed by the updated Kleborate development version, in fact, the new version now additionally calls OmpK36:p.135_136insD (n=12), and OmpK35:- (n=4), and OmpK36:- (n=2) that were previously unidentified.

Noting that there are no new carbapenem resistance genes identified in this new version of Kleborate which includes an updated AMR database.

Up until this point, we have only explored using different versions of Kleborate as the AMR genotyper. The next few sections will explore using different AMR genotypers and comparing their results.

First up, we have AMRFinderPlus!

### Genotypes from AMRFinderPlus

NCBI has developed AMRFinderPlus, a tool that identifies AMR genes, resistance-associated point mutations, and select other classes of genes using protein annotations and/or assembled nucleotide sequence, published here. It can only be used via command line and also has organism-specific options.

#### Import AMRFinderPlus Genotype Data

The download_ebi() can download AMRFinderPlus genotype data from the EBI AMR portal, filter your species of interest, and reformat into AMRgen genotype table. The AMRFinderPlus data being used here is from the 2025-12 release using AMRFinderPlus version v4.0. Noting that as of AMRFinderPlus (v4.2.5), there is additional functionality to identify putatively function disrupting mutations in genes (i.e., ompk35 and ompk36 loss and truncations) that lead to resistance, which Kleborate already identifies.

```
# Download Klebsiella pneumoniae genotype AMRFinderPlus data and re-format the data into
        an AMRgen genotype table
amrfp <- download_ebi(
  data = "genotype", species = "Klebsiella pneumoniae",
  reformat = T
)

# Filter for isolates in EuSCAPE paper with meropenem phenotypes and remove contaminated
        samples
kp_mero_amrfp <- kp_mero_amrfp %>%
  filter(id %in% kp_mero_euscape$id) %>%
  filter(!id %in% contaminated_assemblies)
```

A copy of this data frame is avaiable in the AMRgen pacakge as `kp_mero_amrfp`:

```
head(kp_mero_amrfp)
#> # A tibble: 6 × 34
#>   id         marker gene  mutation drug_agent drug_class marker.label assembly_ID
#>   <chr>      <chr>  <chr> <chr>    <ab>       <chr>      <chr>        <chr>
#> 1 SAMEA364… ompK3… ompK… Asp135A… NA         Carbapene… ompK36:Asp1… GCA_900500…
#> 2 SAMEA364… gyrA_… gyrA  Ser83Ile NA         Quinolones gyrA:Ser83I… GCA_900500…
#> 3 SAMEA364… fosA   fosA  -        FOS: Fosf… Phosphoni… fosA         GCA_900500…
#> 4 SAMEA364… parC_… parC  Ser80Ile NA         Quinolones parC:Ser80I… GCA_900500…
#> 5 SAMEA364… oqxB   oqxB  -        NA         Phenicols  oqxB         GCA_900500…
#> 6 SAMEA364… oqxB   oqxB  -        NA         Quinolones oqxB         GCA_900500…
#> # ℹ 26 more variables: genus <chr>, species <chr>, organism <chr>,
#> #   isolate <chr>, taxon_id <int>, region <chr>, region_start <int>,
#> #   region_end <int>, strand <chr>, `_bin` <int>, id2 <chr>, gene_symbol <chr>,
#> #   amr_element_symbol <chr>, element_type <chr>, element_subtype <chr>,
#> #   class <chr>, subclass <chr>, split_subclass <chr>, antibiotic_name <chr>,
#> #   antibiotic_ontology <chr>, antibiotic_ontology_link <chr>,
#> #   evidence_accession <chr>, evidence_type <chr>, evidence_link <chr>, …

# Summary of carbapenem resistance determinants
summarise_geno(kp_mero_amrfp)
#> $uniques
#> # A tibble: 4 × 2
#>   column     n_unique
#>   <chr>         <int>
#> 1 id             1490
#> 2 marker          237
#> 3 drug_class       20
```

```
#> 4 gene           218
#>
#> $per_type
#> NULL
#>
#> $drugs
#> # A tibble: 20 × 4
#>    drug_class                markers samples     n
#>    <chr>                       <int>   <int> <int>
#>  1 Aminoglycosides                35    1020  6046
#>  2 Beta-lactams                   45    1424  2353
#>  3 Carbapenems                    16     669   956
#>  4 Cephalosporins                  1      23    48
#>  5 Cephalosporins (3rd gen.)      36     788  1479
#>  6 Efflux                          1    1483  1483
#>  7 Glycopeptides                   1      49    49
#>  8 Lincosamides                    3       3     5
#>  9 Macrolides                      7     479  6818
#> 10 Other                           1       2     4
#> 11 Penicillins                     1       4    16
#> 12 Phenicols                      25    1458  3802
#> 13 Phosphonics                     5    1488  1491
#> 14 Polymyxins                     12      40    40
#> 15 Quinolones                     40    1468  4959
#> 16 Rifamycins                      3     110   119
#> 17 Streptogramins                  2      83   249
#> 18 Sulfonamides                    3     931  1206
#> 19 Tetracyclines                  12     605   699
#> 20 Trimethoprims                  12     539   563
#>
#> $markers
#> # A tibble: 261 × 3
#>    marker        drug_class          n
#>    <chr>         <chr>           <int>
#>  1 aac(3)-IId    Aminoglycosides   122
#>  2 aac(3)-IIe    Aminoglycosides   379
#>  3 aac(3)-IIg    Aminoglycosides    14
#>  4 aac(3)-IVa    Aminoglycosides    45
#>  5 aac(3)-Ia     Aminoglycosides     5
#>  6 aac(3)-VIa    Aminoglycosides     1
#>  7 aac(6')-IIc   Aminoglycosides   189
#>  8 aac(6')-Ib    Aminoglycosides  3100
#>  9 aac(6')-Ib'   Aminoglycosides    11
#> 10 aac(6')-Ib-cr Aminoglycosides    24
#> # ℹ 251 more rows
```

#### Compare AMRFinderPlus to Kleborate genotype results

We are going to compare the AMRFinderPlus results that we just downloaded and the Kleborate development branch results. We know that there are differences between AMRFinderPlus and Kleborate development branch in detecting porin defects:

- Kleborate development branch does not identify OmpK35_E132K. As per NCBI Reference Gene Catalog, the citation related to this mutation is PMID: 20660684. In the paper, the mutation

  (ompK35_E132K) is detected in a strain (AIS080884) with an ompk36 mutation (Ser255Thr) that is categorized as low carbapenem resistance. There is no other other literature that experimentally tests this mutation alone, other papers only report the presence of the mutation (often in combination with other mutations/carbapenemases).
- AMRFinderPlus (<v4.2.5) only does not detect nucleotide mutations (e.g., OmpK36_c25t), nor does it detect loss of OmpK35/36 (e.g., OmpK35:- or OmpK36:-).

As such, these differences make it difficult to compare, so we will simplify and remove `OmpK35:-` and `OmpK36:-` from the Kleborate development branch results.

```
# To count and see the names of the carbapenem resistance determinants
kp_mero_amrfp %>%
  filter(drug_class == "Carbapenems") %>%
  count(marker, sort = TRUE)
#> # A tibble: 16 × 2
#>    marker            n
#>    <chr>         <int>
#>  1 ompK36_D135DGD   281
#>  2 blaOXA-48        219
#>  3 blaKPC-3         194
#>  4 blaKPC-2          77
#>  5 blaNDM-1          73
#>  6 blaVIM-1          32
#>  7 blaVIM-4          25
#>  8 ompK35_E132K      22
#>  9 ompK36_D135DD     12
#> 10 ompK36_T136TDT     9
#> 11 blaOXA-232         4
#> 12 blaIMP-1           3
#> 13 blaOXA-162         2
#> 14 blaNDM             1
#> 15 blaOXA-204         1
#> 16 blaOXA-427         1

# Massaging AMRfp marker names to match Kleborate names
amrfp_simplified <- kp_mero_amrfp %>%
  filter(drug_class == "Carbapenems") %>%
  mutate(
    marker_amrfp = str_replace_all(marker, "bla", ""),
    marker_amrfp = str_replace_all(marker_amrfp, "ompK36_D135DGD",
        "OmpK36:p.134_135insGD"),
    marker_amrfp = str_replace_all(marker_amrfp, "ompK36_D135DD", "OmpK36:p.135_136insD"),
    marker_amrfp = str_replace_all(marker_amrfp, "ompK36_T136TDT",
        "OmpK36:p.136_137insTD")
  ) %>%
  select(id, marker_amrfp) %>%
  rename(AMRfp = marker_amrfp) %>%
  group_by(id) %>%
  summarise(
    AMRfp_markers = AMRfp %>%
      sort() %>%
      str_c(collapse = ";")
  )

# Filtering Kleborate AMR markers
```

```r
# Excluding `OmpK35:-` and `OmpK36:-` since we know that AMRfinderplus (<v4.2.5) does not
        detect loss/truncations of OmpK35 and OmpK36
kleborate_dev_simplified <- kleborate_dev %>%
  filter(Kleborate_Class == "Omp_mutations" | Kleborate_Class == "Bla_Carb_acquired") %>%
  select(id, marker.label) %>%
  rename(kleborate = marker.label) %>%
  filter(!kleborate %in% c("OmpK35:-", "OmpK36:-")) %>%
  group_by(id) %>%
  summarise(
    Kleborate_markers = kleborate %>%
      sort() %>%
      str_c(collapse = ";")
  )

# Joining AMRFinderPlus and Kleborate tables
compare_amrfp_kleborate <- full_join(
  amrfp_simplified,
  kleborate_dev_simplified
)
#> Joining with `by = join_by(id)`

# Comparing AMRFinderPlus and Kleborate tables and creating two new columns to show what
        AMRFinderPlus is missing and what Kleborate is missing
compare_amrfp_kleborate <- compare_amrfp_kleborate %>%
  rowwise() %>%
  mutate(
    Kleborate_dev_missing = {
      amr_vec <- str_split(AMRfp_markers, ";")[[1]]
      kleb_vec <- str_split(Kleborate_markers, ";")[[1]]
      missing <- setdiff(amr_vec, kleb_vec)
      if (length(missing) == 0) NA_character_ else str_c(missing, collapse = ";")
    },
    AMRfp_missing = {
      amr_vec <- str_split(AMRfp_markers, ";")[[1]]
      kleb_vec <- str_split(Kleborate_markers, ";")[[1]]
      missing <- setdiff(kleb_vec, amr_vec)
      if (length(missing) == 0) NA_character_ else str_c(missing, collapse = ";")
    }
  ) %>%
  ungroup()

# Table listing the AMR markers that are missing from Kleborate (but detected in
        AMRFinderPlus)
compare_amrfp_kleborate %>% count(Kleborate_dev_missing, sort = TRUE)
#> # A tibble: 11 × 2
#>    Kleborate_dev_missing      n
#>    <chr>                  <int>
#>  1 <NA>                     610
#>  2 ompK35_E132K              22
#>  3 VIM-4                     21
#>  4 OXA-48                    10
#>  5 KPC-3                      7
#>  6 OmpK36:p.134_135insGD      3
#>  7 KPC-2                      2
#>  8 NDM-1                      2
#>  9 NDM                        1
```

```
#> 10 OXA-427                  1
#> 11 VIM-1                    1

# Table listing the AMR markers that are missing from AMRFinderPlus (but detected in
        Kleborate)
compare_amrfp_kleborate %>% count(AMRfp_missing, sort = TRUE)
#> # A tibble: 4 × 2
#>   AMRfp_missing      n
#>   <chr>          <int>
#> 1 <NA>             663
#> 2 OmpK36:c.25C>T    15
#> 3 CTX-M-33           1
#> 4 KPC-12             1
```

Since we know there are differences in porin defect detection between Kleborate and AMRFinderPlus, we can focus on the carbapenemase detection. AMRFinderPlus is missing the detection of `CTX-M-33` in one genome and `KPC-12` in another genome.

#### CTX-M-33

However, AMRFinderPlus is not "missing" `CTX-M-33` and `KPC-12` in their database. In the case of `CTX-M-33`, it is detected in n=1 genome and is annotated as conferring resistance to `Cephalosporins (3rd gen.)` instead of `Carbapenems`, which is why it has been excluded from the AMR genotype table since we filtered for `drug_class=="Carbapenems"`.

```
# CTX-M-33 is annotated as conferring resistance to Cephalosporins (3rd gen.) and is
        identified by AMRFinderPlus in Sample SAMEA3721133
kp_mero_amrfp %>%
  filter(gene == "blaCTX-M-33") %>%
  select(id, gene, drug_class)
#> # A tibble: 1 × 3
#>   id           gene        drug_class
#>   <chr>        <chr>       <chr>
#> 1 SAMEA3721133 blaCTX-M-33 Cephalosporins (3rd gen.)

# Confirming that CTX-M-33 is identified in SAMEA3721133 using Kleborate development
        branch
compare_amrfp_kleborate %>%
  filter(AMRfp_missing == "CTX-M-33") %>%
  select(id, AMRfp_markers, Kleborate_markers)
#> # A tibble: 1 × 3
#>   id           AMRfp_markers Kleborate_markers
#>   <chr>        <chr>         <chr>
#> 1 SAMEA3721133 <NA>          CTX-M-33

# Checking phenotype of SAMEA3721133
kp_mero_euscape %>%
  filter(id == "SAMEA3721133") %>%
  select(id, mic, pheno_eucast, ecoff)
#> # A tibble: 1 × 4
#>   id             mic pheno_eucast ecoff
#>   <chr>        <mic> <sir>        <sir>
#> 1 SAMEA3721133    16 R            NWT
```

The primary literature that describes CTX-M-33 by Galani, et al. describes the clinical strain of *E. coli* 2439 harbouring CTX-M-33, *E. coli* RC85 recipient, and *E. coli* 2439 transconjugant (aka E. coli RC85 recipient + CTX-M-33). The authors performed antibiotic susceptibility tests on *E. coli* RC85 recipient and *E. coli* 2439 transconjugant which showed that MIC for third gen cephalosporins increased, but the MIC for imipenem remained the when harbouring CTX-M-33. In the Comprehensive Antibiotic Resistance Database (CARD, v4.0.1) CTX-M-33 is annotated as conferring resistance to cephalosporins (not carbapenems). The EuSCAPE isolate (SAMEA3721133 from above) only harbours CTM-M-33 and is meropenem resistant. Based on this conflicting evidence, it is unclear if `CTX-M-33` in *K. pneumoniae* confers resistance to carbapenems. The experimental work was performed in an *E. coli* strain and not *K. pneumoniae* and only imipenem was the only carbapenem tested. Additional experimental work and epidemiological support from *K. pneumoniae* strains harbouring CTX-M-33 with carbapenem susceptibility test results is needed to understand the substrate activity of CTX-M-33.

#### KPC-12

Similarly, in the case of `KPC-12`, it is detected in n=1 genome using AMRFinderPlus and is annotated as conferring resistance to `Cephalosporins (3rd gen.)` instead of `Carbapenems`, which is why it has been excluded from the genotype table since we filtered for `drug_class=="Carbapenems"`.

```
# KPC-12 is annotated as conferring resistance to Cephalosporins (3rd gen.)  and is
        identified by AMRFinderPlus in Sample SAMEA3649729
kp_mero_amrfp %>%
  filter(gene == "blaKPC-12") %>%
  select(id, gene, drug_class)
#> # A tibble: 1 × 3
#>   id           gene      drug_class
#>   <chr>        <chr>     <chr>
#> 1 SAMEA3649729 blaKPC-12 Cephalosporins (3rd gen.)

# Confirming that KPC-12 is identified in SAMEA3649729 using Kleborate. It also harbours
        OmpK36:p.134_135insGD
compare_amrfp_kleborate %>%
  filter(AMRfp_missing == "KPC-12") %>%
  select(id, AMRfp_markers, Kleborate_markers)
#> # A tibble: 1 × 3
#>   id           AMRfp_markers         Kleborate_markers
#>   <chr>        <chr>                 <chr>
#> 1 SAMEA3649729 OmpK36:p.134_135insGD KPC-12;OmpK36:p.134_135insGD

# Checking phenotype of SAMEA3649729
kp_mero_euscape %>%
  filter(id == "SAMEA3649729") %>%
  select(id, mic, pheno_eucast, ecoff)
#> # A tibble: 1 × 4
#>   id             mic pheno_eucast ecoff
#>   <chr>        <mic> <sir>        <sir>
#> 1 SAMEA3649729    32 R            NWT
```

The only primary literature discussing KPC-12 is by Han, et al. where they show that a strain of *E. coli* DH5alpha + empty plasmid vs. *E. coli* DH5alpha + plasmid with `KPC-12` does not change meropenem MIC (0.06mg/L) with small elevation in imipenem (0.25 vs. 1 mg/L) and ertapenem (<=0.12mg/L vs. 0.25 mg/L) MICs. Whereas, *E. coli* DH5alpha + empty plasmid vs. *E. coli* DH5alpha + plasmid with `KPC-12` elevates the MICs for ceftriaxone (<=0.25 vs. >=64 mg/L, 3rd gen cephalosporin) and cefuroxime (8 vs. 256 mg/L, 2nd gen cephalosporin). This experiment was performed in an *E. coli* strain, not *K. pneumoniae*. In the EuSCAPE dataset (from above), there is only one isolate

(SAMEA3649729) with KPC-12, which harbours both KPC-12 and OmpK36:p.134_135insGD and is meropenem resistant. Based on this conflicting evidence, it is not clear whether `KPC-12` should be changed to conferring resistance to 3rd generation cephalosporins or carbapenems. Similar to Kleborate, CARD v4.0.1 also has KPC-12 annotated as a carbapenemase. Further experimental work performed in a *K. pneumoniae* strain and having additional evidence from *K. pneumoniae* strains with genotype-phenotype data can help strengthen our understanding of its substrate specificity.

We will not continue to investigate what is missing in the Kleborate development branch, compared to AMRFinderPlus, for the purpose and lengthiness of this vignette. These two examples act as ways to investigate differences in AMR genotypers and how ultimately, choosing a specific genotyper will impact the foundation upon which we understand AMR genotype-phenotype relationships.

Next up, we have the Resistance Gene Identifier (RGI)!

### Genotypes from Resistance Gene Identifier (RGI)

RGI identifies resistance determinants from protein or nucleotide data using homology and mutation models, published here. The software uses reference data from the Comprehensive Antibiotic Resistance Database. It can be run via the website or on the command line.

CARD is an ontology-drive database including resistance genes, their products, and the antibiotics they confer resistance towards. CARD operates at both a drug class and drug-specific level, where curators can establish `gene confers_resistance_to_drug antibiotic` relationships. Lastly, CARD includes both intrinsic/core and acquired resistance determinants.

#### Import Resistance Gene Identifier (RGI) results

Import RGI results using the `import_rgi()` function. This function imports and processes genotyping results from RGI extracting antimicrobial resistance determinants and mapping them to standardised drug classes/antibiotics. It also shortens determinant names using `CARD Short Names` as provided by CARD (https://card.mcmaster.ca/download in `aro_index.tsv`).

**Note**: Check the number of genomes that you expect using `summarise_geno()`. RGI text output will be empty if there are no AMR determinants identified in the submitted genome, so you have to either:

1. Add sample IDs with no AMR determinants into the `ORF_ID` column of the RGI text output that you are importing, or
2. List your sample IDs in a vector using the `samples_no_amr=` parameter in the `import_rgi()` function. For example, `import_rgi(rgi_EuSCAPE_raw, samples_no_amr = c("SampleA_noAMR", "SampleB_noAMR", "SampleC_noAMR"))`

The data frame `rgi_EuSCAPE_raw` included in the AMRgen package provides CARD RGI calls for the EuSCAPE genomes:

```
# Sample IDs with no AMR determinants have been added to rgi_EuSCAPE_raw under the
        `ORF_ID` column with the rest of the rows left blank
tail(rgi_EuSCAPE_raw, n = 10)
#> # A tibble: 10 × 26
#>    ORF_ID Contig Start  Stop Orientation Cut_Off Pass_Bitscore Best_Hit_Bitscore
#>    <chr>   <dbl> <dbl> <dbl> <chr>       <chr>           <dbl>             <dbl>
#>  1 SAMEA…     NA    NA    NA <NA>        <NA>               NA                NA
#>  2 SAMEA…     NA    NA    NA <NA>        <NA>               NA                NA
#>  3 SAMEA…     NA    NA    NA <NA>        <NA>               NA                NA
#>  4 SAMEA…     NA    NA    NA <NA>        <NA>               NA                NA
```

```
#>   5 SAMEA…     NA    NA    NA <NA>        <NA>                NA                 NA
#>   6 SAMEA…     NA    NA    NA <NA>        <NA>                NA                 NA
#>   7 SAMEA…     NA    NA    NA <NA>        <NA>                NA                 NA
#>   8 SAMEA…     NA    NA    NA <NA>        <NA>                NA                 NA
#>   9 SAMEA…     NA    NA    NA <NA>        <NA>                NA                 NA
#>  10 SAMEA…     NA    NA    NA <NA>        <NA>                NA                 NA
#> # ℹ 18 more variables: Best_Hit_ARO <chr>, Best_Identities <dbl>, ARO <dbl>,
#> #   Model_type <chr>, SNPs_in_Best_Hit_ARO <chr>, Other_SNPs <chr>,
#> #   `Drug Class` <chr>, `Resistance Mechanism` <chr>, `AMR Gene Family` <chr>,
#> #   `Percentage Length of Reference Sequence` <dbl>, ID <chr>, Model_ID <dbl>,
#> #   Nudged <lgl>, Note <lgl>, Hit_Start <dbl>, Hit_End <dbl>, Antibiotic <chr>,
#> #   AST_Source <chr>

# Import RGI output with n=1490 isolates
rgi <- import_rgi(rgi_EuSCAPE_raw)

# Summarize genotype data
summarise_geno(rgi)
#> $uniques
#> # A tibble: 5 × 2
#>   column         n_unique
#>   <chr>             <int>
#> 1 id                 1490
#> 2 marker              252
#> 3 drug                109
#> 4 drug_class           30
#> 5 variation type        3
#> 
#> $per_type
#> # A tibble: 3 × 5
#>   `variation type`            id marker  drug drug_class
#>   <chr>                    <int>  <int> <int>      <int>
#> 1 Gene presence detected    1430    233    98         29
#> 2 Protein variant detected  1445     18    37         13
#> 3 <NA>                        45      1     1          1
#> 
#> $drugs
#> # A tibble: 114 × 5
#>    drug                 drug_class       markers samples  hits
#>    <chr>                <chr>              <int>   <int> <int>
#>  1 2'-N-ethylnetilmicin Aminoglycosides        8     462   492
#>  2 5-episisomicin       Aminoglycosides        3      37    37
#>  3 6'-N-ethylnetilmicin Aminoglycosides        6     449   456
#>  4 AMK                  Aminoglycosides       18    1445  3846
#>  5 AMP                  Aminopenicillins      22    1445 10675
#>  6 AMX                  Aminopenicillins       6     747  1110
#>  7 APR                  Aminoglycosides        1       5     5
#>  8 ARB                  Aminoglycosides        4      72    73
#>  9 AST                  Aminoglycosides        2      17    17
#> 10 ATM                  Monobactams            1       1     1
#> # ℹ 104 more rows
#> 
#> $markers
#> # A tibble: 815 × 5
#>    marker     drug                drug_class      `variation type`            n
```

```
#>    <chr>      <chr>               <chr>          <chr>                  <int>
#>  1 AAC(3)-IIc 2'-N-ethylnetilmicin Aminoglycosides Gene presence detected    16
#>  2 AAC(3)-IIc 6'-N-ethylnetilmicin Aminoglycosides Gene presence detected    16
#>  3 AAC(3)-IIc DKB                Aminoglycosides Gene presence detected    16
#>  4 AAC(3)-IIc GEN                Aminoglycosides Gene presence detected    16
#>  5 AAC(3)-IIc NET                Aminoglycosides Gene presence detected    16
#>  6 AAC(3)-IIc SIS                Aminoglycosides Gene presence detected    16
#>  7 AAC(3)-IIc TOB                Aminoglycosides Gene presence detected    16
#>  8 AAC(3)-IId 2'-N-ethylnetilmicin Aminoglycosides Gene presence detected    88
#>  9 AAC(3)-IId 6'-N-ethylnetilmicin Aminoglycosides Gene presence detected    88
#> 10 AAC(3)-IId DKB                Aminoglycosides Gene presence detected    88
#> # ℹ 805 more rows
```

#### Generate Binary Matrix for RGI AMR Markers

```
rgi_binary_matrix <- get_binary_matrix(
  geno_table = rgi,
  pheno_table = kp_mero_euscape,
  pheno_drug = "Meropenem",
  geno_class = c("Carbapenems"),
  sir_col = "pheno_eucast",
  marker_col = "marker.label",
  keep_assay_values = TRUE,
  keep_assay_values_from = "mic"
)
#>  Defining NWT in binary matrix using ecoff column provided: ecoff
```

#### Solo PPV Analysis for RGI AMR Markers

```
# No solo markers error when you run solo_ppv()! Since CARD/RGI includes intrinsic and
        acquired resistance determinants, there could be intrinsic / core resistance
        determinants that are found across most (if not all) genomes which obstructs our
        view of carbapenem resistance determinants found alone.

soloPPV_rgi_mero <- solo_ppv(binary_matrix = rgi_binary_matrix)
```

As such, we will exclude the core/intrinsic resistance determinants, using their prevalence and exclude any determinants identified across more than 80% of genomes.

```
rgi_binary_matrix_prev80 <- rgi_binary_matrix %>%
  select(where(~ {
    if (is.numeric(.x)) {
      prop_ones <- mean(.x == 1, na.rm = TRUE) # fraction of 1s
      prop_ones <= 0.80 # keep only if ≤ 80% prevalent across all genomes
    } else {
      TRUE
    }
  }))
```

Try running the `solo_ppv()` function again.

```r
soloPPV_rgi_mero <- solo_ppv(binary_matrix = rgi_binary_matrix_prev80)
```

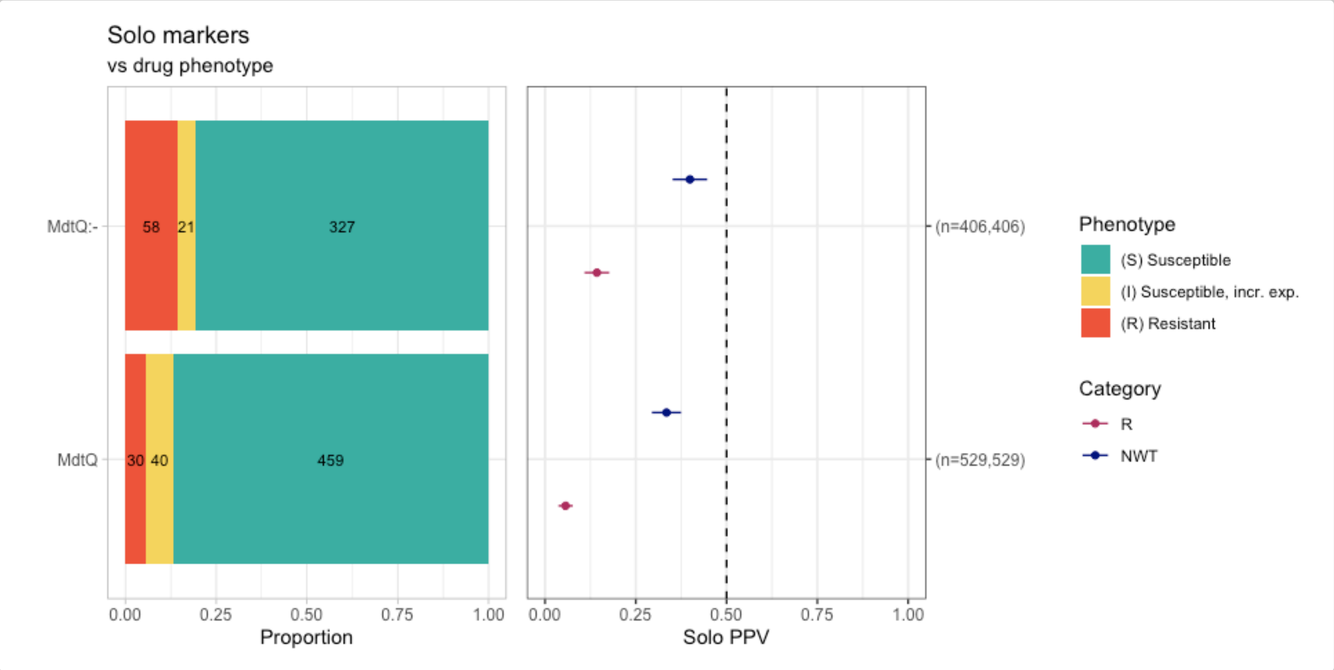


```r
# Count number of genomes that have either `MdtQ` or `MdtQ:-`
sum(rgi_binary_matrix_prev80$MdtQ == "1" | rgi_binary_matrix_prev80$`MdtQ..-` == "1",
        na.rm = TRUE)
#> [1] 1429
```

Only MdtQ and MdtQ variants (MdtQ:-) were identified in the absence of other carbapenem resistance determinants. MdtQ is an outer-membrane porin identified in *K. pneumoniae* - however the primary paper by Fan, et al. only reports a clinical strain with MdtQ resistant to carbapenems, but does not show antibiotic susceptibility tests for proper controls of the same strain with MdtQ vs. without MdtQ. In addition, 96% (n=1429/1490) of the EuSCAPE *K. pneumoniae* genomes have MdtQ or a variant of MdtQ, therefore it could be considered a core gene. In summary, because is no compelling evidence that MdtQ confers resistance to carbapenems and that it is likely a core gene, we will exclude it from further analyses.

```r
# Exclude MdtQ and MdtQ:- from the binary matrix
rgi_binary_matrix_prev80 <- rgi_binary_matrix_prev80 %>% select(-MdtQ, -`MdtQ..-`)
```

Try running the `solo_ppv()` function... again.

```r
soloPPV_rgi_mero <- solo_ppv(binary_matrix = rgi_binary_matrix_prev80)
```

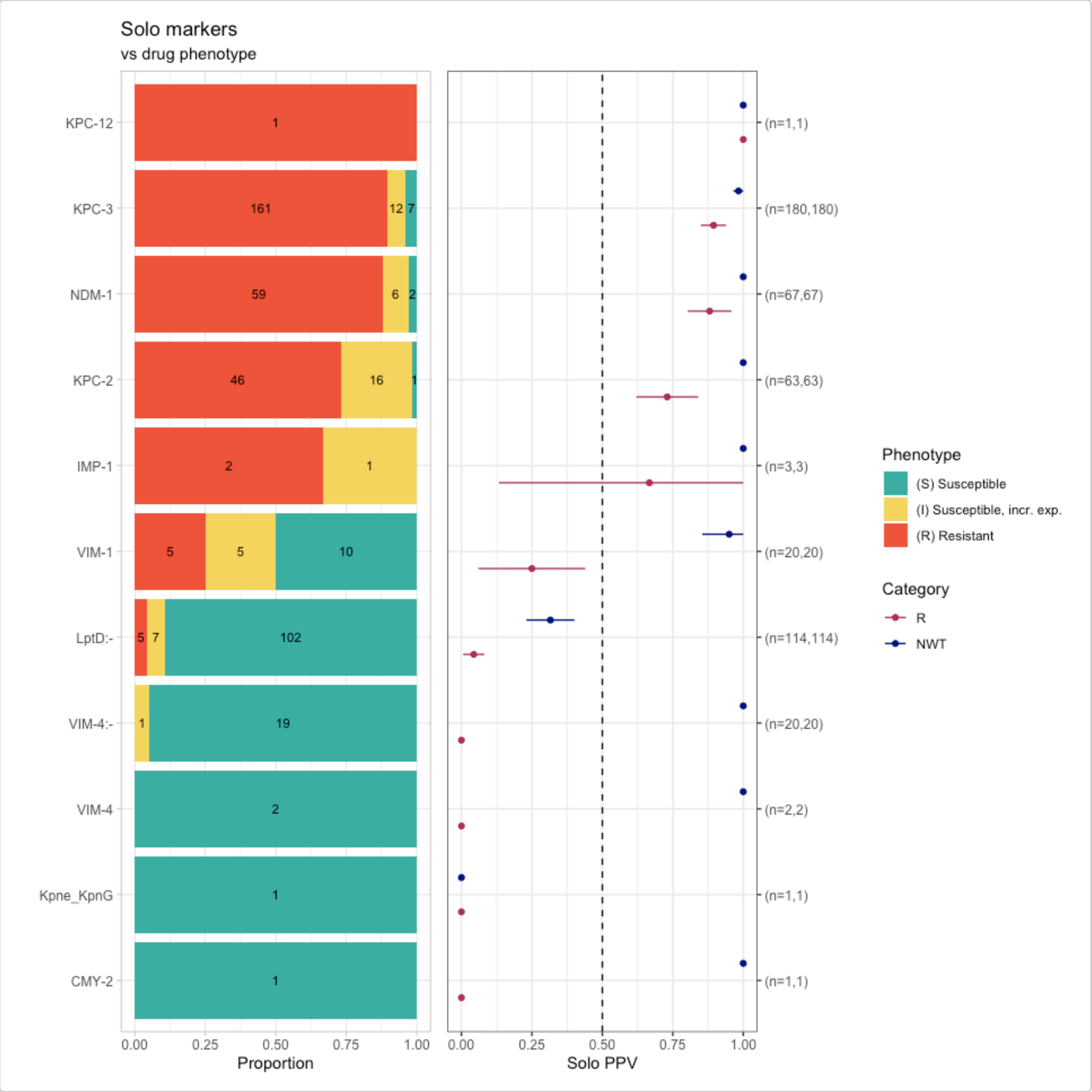


We can finally see carbapenem resistance determinants alone! Since RGI does not detect porin defects, many AMR markers are found alone compared to Kleborate's and AMRFinderPlus' `solo_ppv()` (see below). For example, NDM-1 alone was found in 59 resistant genomes (using RGI), 31 resistant genomes (using Kleborate), and 41 resistant genomes (using AMRFinderPlus).

```
soloPPV_kleborate_mero
#> $solo_stats
#> # A tibble: 28 × 8
#>    marker               category     x     n    ppv      se ci.lower ci.upper
#>    <chr>                <chr>    <dbl> <int>  <dbl>   <dbl>    <dbl>    <dbl>
#>  1 OXA-162              R            0     2 0      0        0        0
#>  2 OXA-204              R            0     1 0      0        0        0
#>  3 OmpK36:c.25C>T       R            0     6 0      0        0        0
#>  4 OmpK36:p.134_135insGD R           0     9 0      0        0        0
#>  5 VIM-1                R            0    13 0      0        0        0
#>  6 VIM-4                R            0     4 0      0        0        0
#>  7 OmpK36:-             R            1    71 0.0141 0.0140   0        0.0415
#>  8 OmpK35:-             R            2    49 0.0408 0.0283   0        0.0962
#>  9 OXA-48               R            6    99 0.0606 0.0240   0.0136   0.108
```

```
#> 10 KPC-3                R              3     10 0.3     0.145     0.0160    0.584
#> # ℹ 18 more rows
#>
#> $combined_plot
```

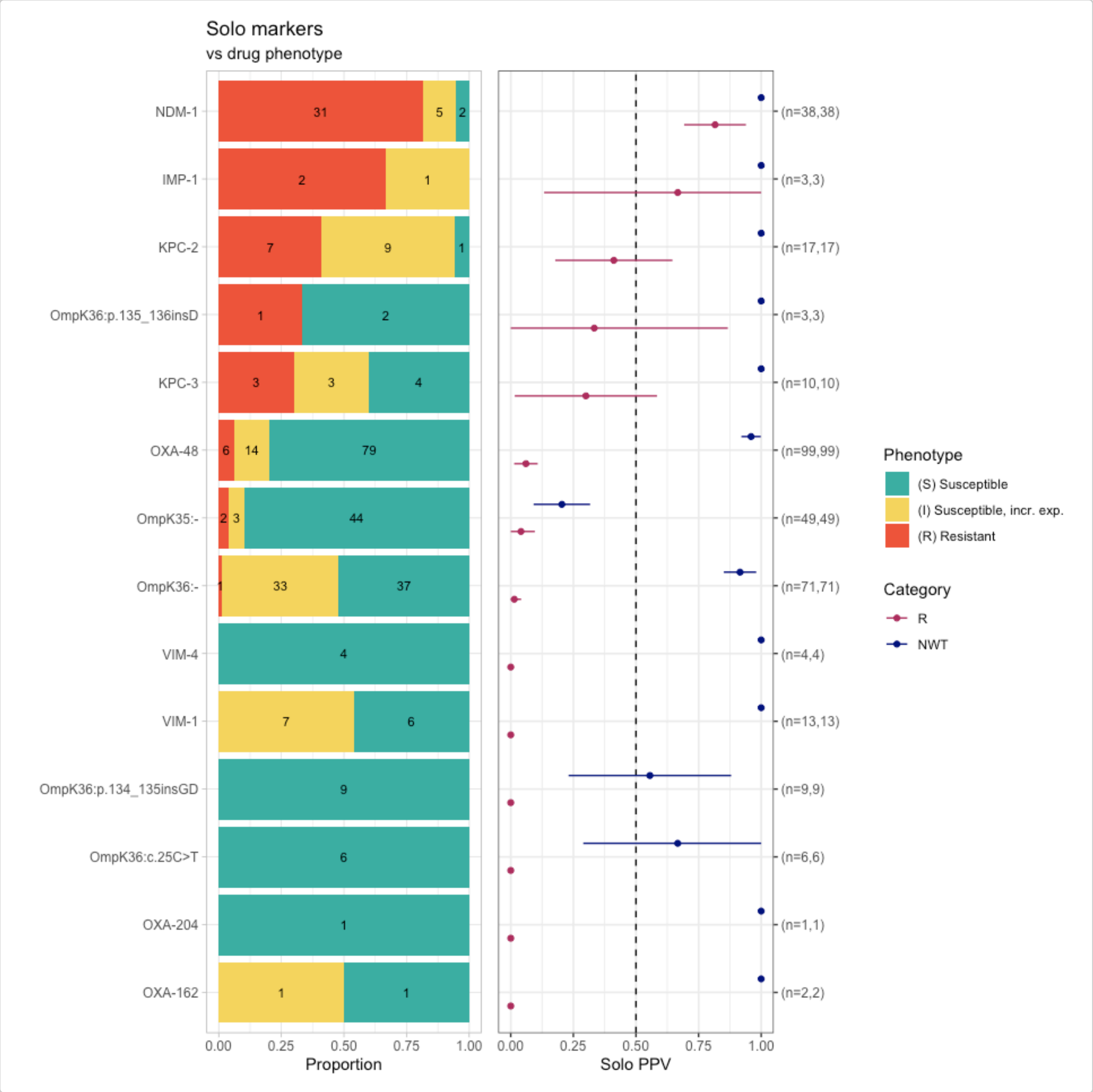


```
#>
#> $solo_binary
#> # A tibble: 325 × 8
#>    id             pheno ecoff    mic     R   NWT marker    value
#>    <chr>          <sir> <sir>  <mic> <dbl> <dbl> <chr>     <dbl>
#>  1 SAMEA3498967 I     NWT     4.00     0     1 OmpK36:-      1
#>  2 SAMEA3498970 I     NWT     4.00     0     1 OmpK36:-      1
#>  3 SAMEA3498975 S     NWT     2.00     0     1 OmpK36:-      1
#>  4 SAMEA3498992 S     NWT     0.50     0     1 OmpK36:-      1
#>  5 SAMEA3498996 S     WT     <=0.06    0     0 OmpK35:-      1
#>  6 SAMEA3498997 S     NWT     0.50     0     1 OmpK36:-      1
#>  7 SAMEA3498998 S     NWT     2.00     0     1 OmpK36:-      1
#>  8 SAMEA3499003 R     NWT   >32.00     1     1 NDM-1         1
```

```
#>   9 SAMEA3499004 R     NWT    32.00     1     1 OmpK36:-     1
#>  10 SAMEA3499010 S     WT    <=0.06     0     0 OmpK35:-     1
#> # ℹ 315 more rows
#>
#> $solo_binary_norange
#> NULL
#>
#> $amr_binary
#> # A tibble: 1,490 × 24
#>    id    pheno ecoff    mic     R   NWT `OmpK36..-` `OmpK35..-` `NDM-1` `OXA-48`
#>    <chr> <sir> <sir>  <mic> <dbl> <dbl>       <dbl>       <dbl>   <dbl>    <dbl>
#>  1 SAME… I     NWT     4.00     0     1           1           0       0        0
#>  2 SAME… S     WT    <=0.06     0     0           0           0       0        0
#>  3 SAME… S     NWT     1.00     0     1           1           1       0        0
#>  4 SAME… I     NWT     4.00     0     1           1           0       0        0
#>  5 SAME… I     NWT     4.00     0     1           1           1       0        0
#>  6 SAME… S     WT    <=0.06     0     0           0           0       0        0
#>  7 SAME… S     WT    <=0.06     0     0           0           0       0        0
#>  8 SAME… S     NWT     2.00     0     1           1           0       0        0
#>  9 SAME… S     WT    <=0.06     0     0           0           0       0        0
#> 10 SAME… S     NWT     0.50     0     1           1           0       0        0
#> # ℹ 1,480 more rows
#> # ℹ 14 more variables: `OmpK36..c.25C>T` <dbl>, `KPC-3` <dbl>,
#> #   OmpK36..p.134_135insGD <dbl>, `KPC-2` <dbl>, OmpK36..p.135_136insD <dbl>,
#> #   `OXA-204` <dbl>, `VIM-1` <dbl>, OmpK36..p.136_137insTD <dbl>,
#> #   `VIM-4` <dbl>, `KPC-12` <dbl>, `OXA-232` <dbl>, `CTX-M-33` <dbl>,
#> #   `OXA-162` <dbl>, `IMP-1` <dbl>
#>
#> $plot_order
#>                OXA-162                OXA-204        OmpK36:c.25C>T
#>              "(n=2,2)"              "(n=1,1)"             "(n=6,6)"
#> OmpK36:p.134_135insGD                   VIM-1                 VIM-4
#>              "(n=9,9)"            "(n=13,13)"             "(n=4,4)"
#>               OmpK36:-               OmpK35:-                OXA-48
#>            "(n=71,71)"            "(n=49,49)"           "(n=99,99)"
#>                  KPC-3  OmpK36:p.135_136insD                 KPC-2
#>            "(n=10,10)"              "(n=3,3)"           "(n=17,17)"
#>                  IMP-1                  NDM-1
#>              "(n=3,3)"            "(n=38,38)"
```

```
# Generate binary matrix for AMRFinderPlus
amrfp_binary_matrix <- get_binary_matrix(
  geno_table = kp_mero_amrfp,
  pheno_table = kp_mero_euscape,
  pheno_drug = "Meropenem",
  geno_class = c("Carbapenems"),
  sir_col = "pheno_eucast",
  keep_assay_values = TRUE,
  keep_assay_values_from = "mic"
)
#>  Defining NWT in binary matrix using ecoff column provided: ecoff
# Solo PPV analysis
soloPPV_amrfp_mero <- solo_ppv(binary_matrix = amrfp_binary_matrix)
```

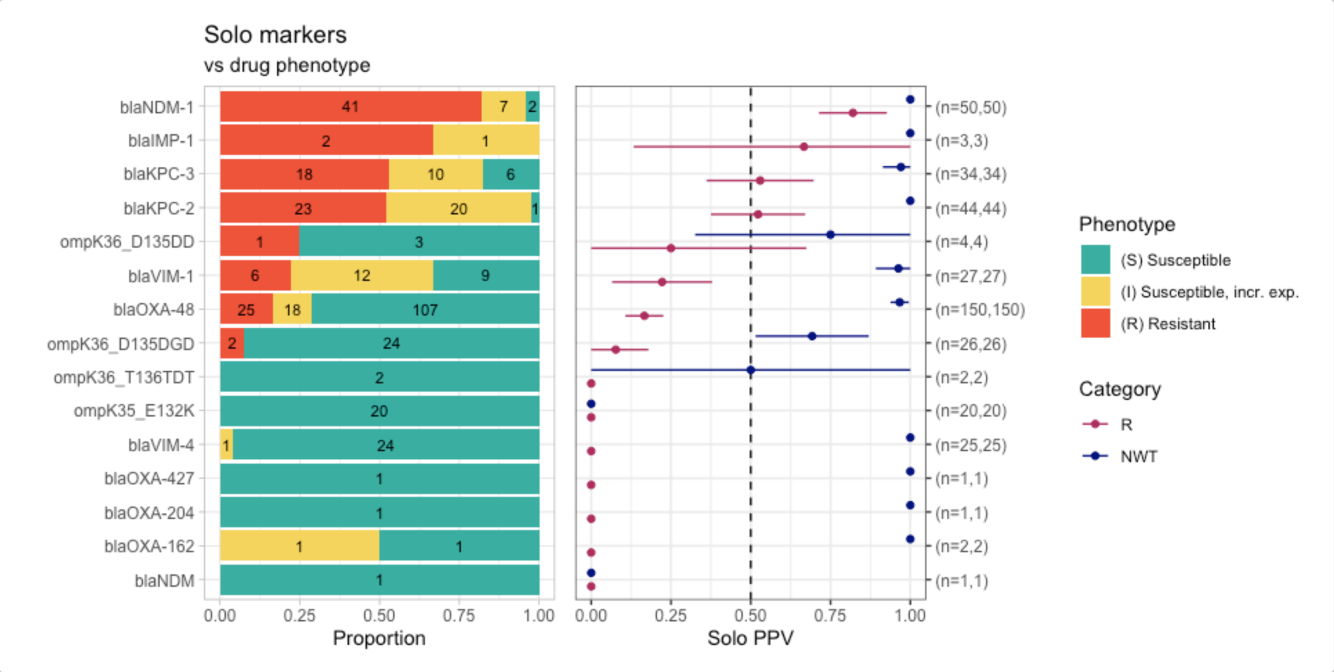


#### Combinatorial PPV Analysis for RGI AMR Markers

```
comboPPV_rgi_mero <- amr_ppv(
  binary_matrix = rgi_binary_matrix_prev80,
  order = "value",
  min_set_size = 2,
  pheno_drug = "Meropenem",
  upset_grid = TRUE,
  plot_assay = TRUE,
  assay = "mic"
)
#> Ordering markers by frequency
#> Scale for y is already present.
#> Adding another scale for y, which will replace the existing scale.
```

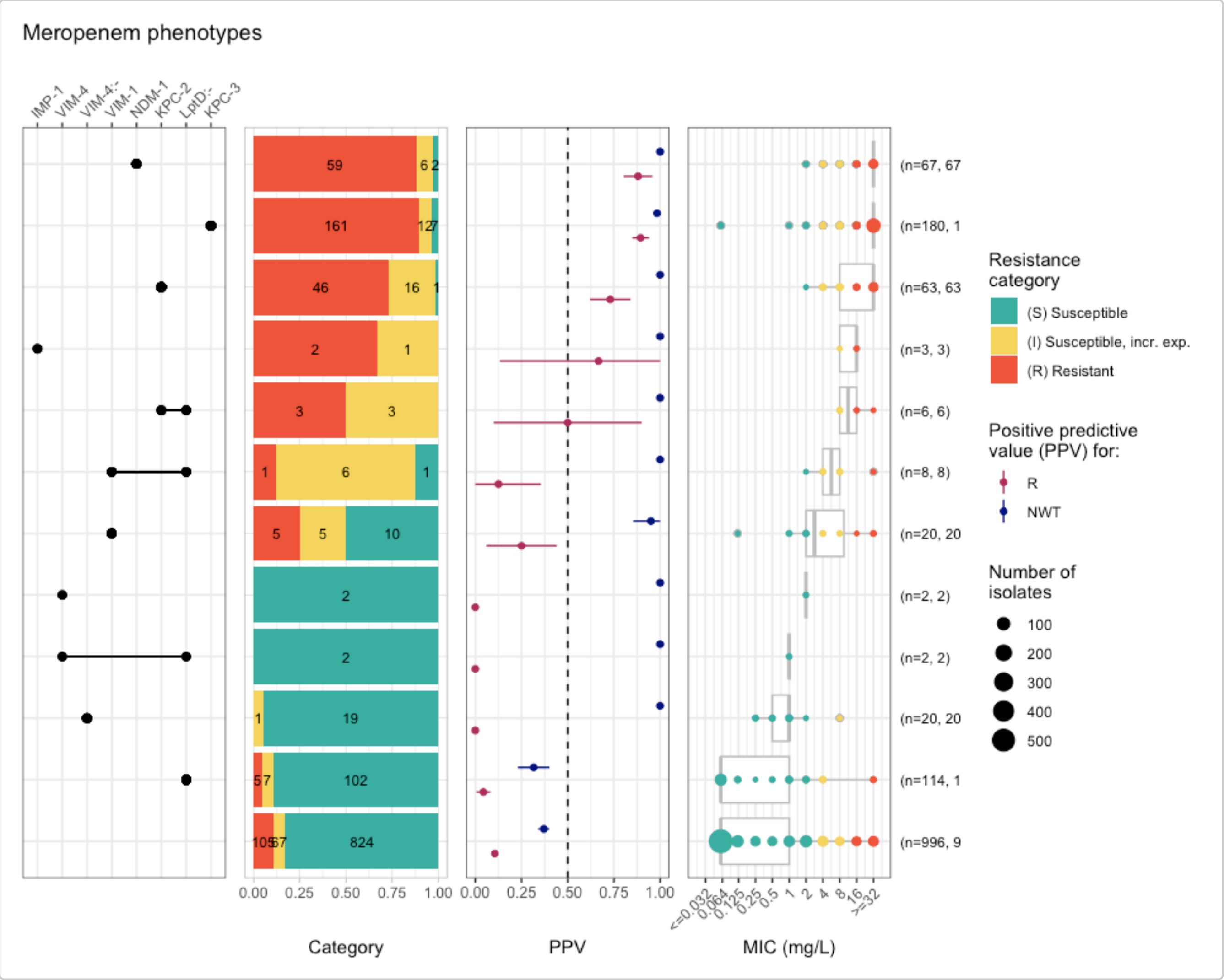


```
# Summary of combinatorial PPV
comboPPV_rgi_mero$summary
#> # A tibble: 21 × 21
#>    marker_list          marker_count     n combination_id    R.n  R.ppv R.ci_lower
#>    <chr>                       <dbl> <int> <fct>           <dbl>  <dbl>      <dbl>
#>  1 ""                              0   996 0_0_0_0_0_0_0_…   105 0.105     0.0863
#>  2 "IMP-1"                         1     3 0_0_0_0_0_0_0_…     2 0.667     0.133
#>  3 "CMY-2"                         1     1 0_0_0_0_0_0_0_…     0 0         0
#>  4 "KPC-12"                        1     1 0_0_0_0_0_0_0_…     1 1         1
#>  5 "Kpne_KpnG"                     1     1 0_0_0_0_0_0_0_…     0 0         0
#>  6 "VIM-4"                         1     2 0_0_0_0_0_0_1_…     0 0         0
#>  7 "VIM-4:-"                       1    20 0_0_0_0_0_1_0_…     0 0         0
#>  8 "VIM-1"                         1    20 0_0_0_0_1_0_0_…     5 0.25      0.0602
#>  9 "LptD:-"                        1   114 0_0_0_1_0_0_0_…     5 0.0439    0.00627
#> 10 "LptD:-, NDM-69:-"              2     1 0_0_0_1_0_0_0_…     0 0         0
#> # ℹ 11 more rows
#> # ℹ 14 more variables: R.ci_upper <dbl>, R.denom <int>, NWT.n <dbl>,
#> #   NWT.ppv <dbl>, NWT.ci_lower <dbl>, NWT.ci_upper <dbl>, NWT.denom <int>,
#> #   median_excludeRangeValues <dbl>, q25_excludeRangeValues <dbl>,
#> #   q75_excludeRangeValues <dbl>, n_excludeRangeValues <int>,
#> #   median_ignoreRanges <dbl>, q25_ignoreRanges <dbl>, q75_ignoreRanges <dbl>
```

Evidently, RGI does not detect OmpK35 or OmpK36 defects so we can only compare the carbapenemases that are detected by RGI vs. Kleborate development branch.

#### Compare RGI to Kleborate Genotype Results

```r
rgi_simplified <- rgi %>%
  filter(drug_class == "Carbapenems") %>%
  filter(`Resistance Mechanism` == "antibiotic inactivation") %>%
  select(id, marker.label) %>%
  distinct() %>%
  rename(rgi = marker.label) %>%
  group_by(id) %>%
  summarise(
    rgi_markers = rgi %>%
      sort() %>%
      str_c(collapse = ";")
  )

kleborate_simplified <- kleborate_dev %>%
  filter(Kleborate_Class == "Omp_mutations" | Kleborate_Class == "Bla_Carb_acquired") %>%
  select(id, marker.label) %>%
  rename(kleborate = marker.label) %>%
  filter(!grepl("OmpK35|OmpK36", kleborate)) %>%
  group_by(id) %>%
  summarise(
    Kleborate_markers = kleborate %>%
      sort() %>%
      str_c(collapse = ";")
  )

compare_rgi_kleborate <- full_join(
  rgi_simplified,
  kleborate_simplified
)
#> Joining with `by = join_by(id)`

compare_rgi_kleborate <- compare_rgi_kleborate %>%
  rowwise() %>%
  mutate(
    Kleborate_missing = {
      rgi_vec <- str_split(rgi_markers, ";")[[1]]
      kleb_vec <- str_split(Kleborate_markers, ";")[[1]]
      missing <- setdiff(rgi_vec, kleb_vec)
      if (length(missing) == 0) NA_character_ else str_c(missing, collapse = ";")
    },
    rgi_missing = {
      rgi_vec <- str_split(rgi_markers, ";")[[1]]
      kleb_vec <- str_split(Kleborate_markers, ";")[[1]]
      missing <- setdiff(kleb_vec, rgi_vec)
      if (length(missing) == 0) NA_character_ else str_c(missing, collapse = ";")
    }
  ) %>%
  ungroup()

compare_rgi_kleborate %>% count(Kleborate_missing, sort = TRUE)
#> # A tibble: 4 x 2
```

```
#>   Kleborate_missing     n
#>   <chr>              <int>
#> 1 <NA>                 576
#> 2 VIM-4:-               20
#> 3 CMY-2                  1
#> 4 NDM-69:-               1
compare_rgi_kleborate %>% count(rgi_missing, sort = TRUE)
#> # A tibble: 10 × 2
#>    rgi_missing      n
#>    <chr>        <int>
#>  1 <NA>           371
#>  2 OXA-48         207
#>  3 KPC-3            6
#>  4 KPC-2            4
#>  5 OXA-232          4
#>  6 OXA-162          2
#>  7 CTX-M-33         1
#>  8 NDM-1            1
#>  9 NDM-1;OXA-48     1
#> 10 OXA-204          1
```

#### Combining Kleborate, AMRFinderPlus and RGI results

To merge Kleborate (development branch), AMRFinderPlus (from EBI), and RGI results, we will combine the binary matrices generated by `get_binary_matrix()`.

Note that you will have to inspect how each of the AMR markers are named and change them so that they match and can be merged, for example AMRFinderPlus appends "bla" in front of all beta-lactamases, whereas RGI and Kleborate do not. **This assumes that the same name is referring to the same reference sequence that is used in each tool/database which is not necessarily true** (even if we wish it were true). Hypothetical example, the NDM-1 sequence in CARD/RGI is `ABCD` vs. AMRFinderPlus NDM-1 sequence is `ACCD` vs. Kleborate NDM-1 sequence is `ACDD`. All AMR databases strive to use the same reference accessions and sequences, but sometimes there can be discrepancies, which need to be kept in mind.

In the following code, unique AMR markers (i.e., only identified by one AMR genotyper) will be have a suffix to describe the AMR genotyper that it is found by (e.g., Kpne_KpnG will be Kpne_KpnG_rgi). AMR markers identified by more than one genotyper will be merged, where if it was identified by any genotyper in that sample, the binary matrix will have a `1` (present), otherwise `0` (absent).

```
# Phenotype columns to remove (that we can put back in later)
cols_to_remove <- c("pheno", "ecoff", "mic", "R", "NWT")

# Remove columns
# We will be using the RGI binary matrix where core/intrinsic genes are removed
df_rgi <- rgi_binary_matrix_prev80 %>% select(-cols_to_remove)
#> Warning: Using an external vector in selections was deprecated in tidyselect 1.1.0.
#> ℹ Please use `all_of()` or `any_of()` instead.
#>   # Was:
#>   data %>% select(cols_to_remove)
#>
#>   # Now:
#>   data %>% select(all_of(cols_to_remove))
#>
```

```r
#> See <https://tidyselect.r-lib.org/reference/faq-external-vector.html>.
#> This warning is displayed once per session.
#> Call `lifecycle::last_lifecycle_warnings()` to see where this warning was
#> generated.
df_kleborate <- kleborate_binary_matrix %>% select(-cols_to_remove)

# Massage AMRFinderPlus marker names to match RGI/Kleborate
df_amrfp <- amrfp_binary_matrix %>%
  select(-cols_to_remove) %>%
  rename("OmpK36..p.134_135insGD" = "ompK36_D135DGD") %>%
  rename("OmpK36..p.135_136insD" = "ompK36_D135DD") %>%
  rename("OmpK36..p.136_137insTD" = "ompK36_T136TDT")
colnames(df_amrfp) <- gsub("bla", "", colnames(df_amrfp))

# Sort each dataframe by id to make sure the samples are all in the same order
df_rgi <- df_rgi[order(df_rgi[[1]]), ]
df_amrfp <- df_amrfp[order(df_amrfp[[1]]), ]
df_kleborate <- df_kleborate[order(df_kleborate[[1]]), ]

# All column names (excluding id)
cols_rgi <- colnames(df_rgi)[-1]
cols_amrfp <- colnames(df_amrfp)[-1]
cols_kleb <- colnames(df_kleborate)[-1]

all_cols <- unique(c(cols_rgi, cols_amrfp, cols_kleb))

# Function to safely get column or return 0s
get_col <- function(df, col) {
  if (col %in% colnames(df)) {
    df[[col]]
  } else {
    rep(0, nrow(df))
  }
}

# Initialize final df (assuming same order of ids)
combined_binary_matrix <- data.frame(id = df_rgi[[1]])

# Merge all columns using OR
for (col in all_cols) {
  combined_binary_matrix[[col]] <- as.integer(
    get_col(df_rgi, col) |
      get_col(df_amrfp, col) |
      get_col(df_kleborate, col)
  )
}

# Identify unique columns (present in ONLY one binary matrix)
unique_rgi <- setdiff(cols_rgi, union(cols_amrfp, cols_kleb))
unique_amrfp <- setdiff(cols_amrfp, union(cols_rgi, cols_kleb))
unique_kleb <- setdiff(cols_kleb, union(cols_rgi, cols_amrfp))

# Rename unique columns with suffix
colnames(combined_binary_matrix)[colnames(combined_binary_matrix) %in% unique_rgi] <-
        paste0(unique_rgi, "_rgi")
```

```r
colnames(combined_binary_matrix)[colnames(combined_binary_matrix) %in% unique_amrfp] <-
        paste0(unique_amrfp, "_amrfp")
colnames(combined_binary_matrix)[colnames(combined_binary_matrix) %in% unique_kleb] <-
        paste0(unique_kleb, "_kleborate")

# Joining back the phenotype columns
phenotype_cols <- rgi_binary_matrix_prev80 %>%
  select(id, pheno, ecoff, mic, R, NWT)

# Merge back into your final_df
combined_binary_matrix <- combined_binary_matrix %>%
  left_join(phenotype_cols, by = "id")

# Relocate phenotype columns to the front
combined_binary_matrix <- combined_binary_matrix %>%
  relocate(pheno, ecoff, mic, R, NWT, .after = id)
```

#### Solo PPV Analysis for AMRFinderPlus, RGI, Kleborate AMR Markers

```r
combined_solo_ppv <- solo_ppv(binary_matrix = combined_binary_matrix)
```

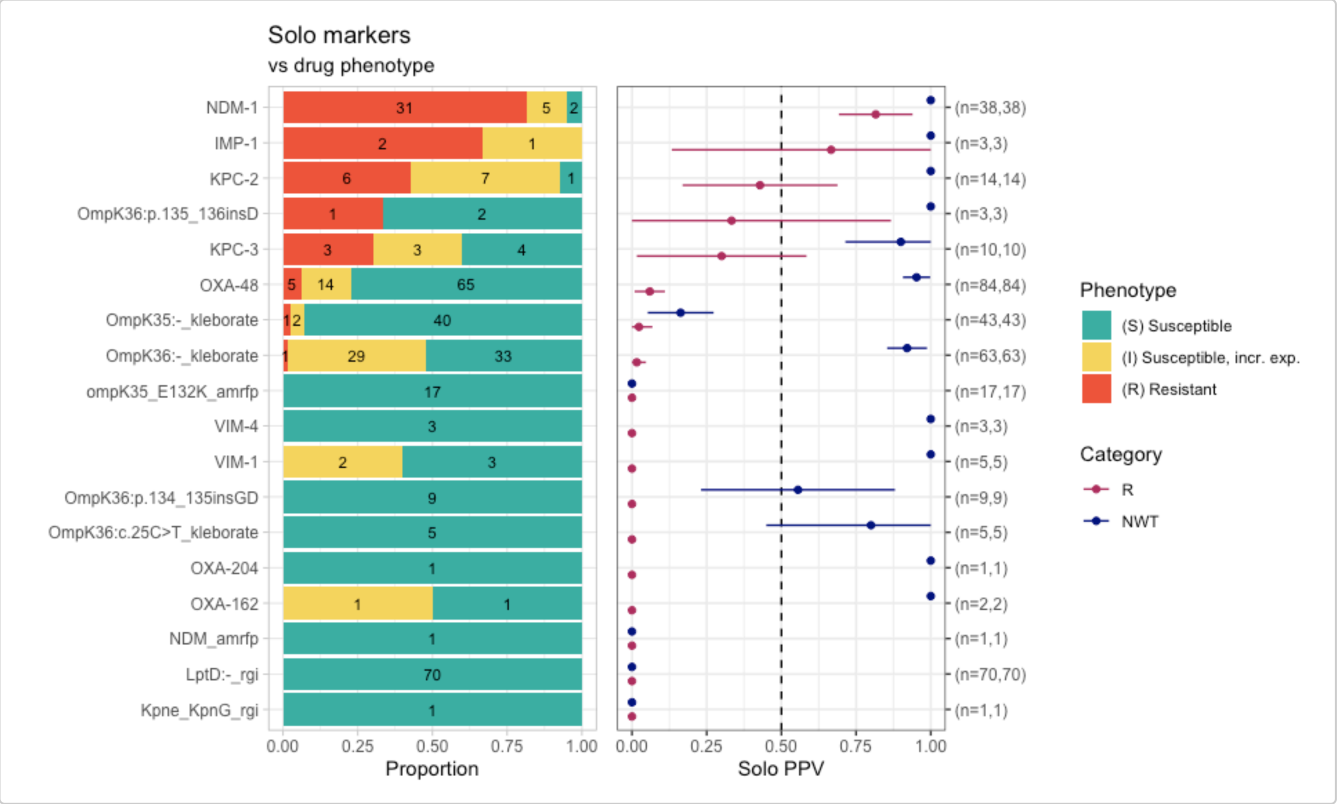


From this solo_ppv() plot, we can see that there are markers that are found alone which have strong support for a particular phenotype, e.g., NDM-1 association with meropenem resistance (n=31/38 R isolates), LptD:- identified by RGI (associated with susceptibility). Noting that LptD:- indicates a variant of LptD, so the variants need to be further investigated to see if there is a particular mutation/defect that is associated with meropenem susceptibility.

Another way to investigate the association between AMR markers and meropenem susceptibility is to use the `amr_logistic()` function to perform logistic regression to analyse the relationship between the markers and a specified antibiotic.

#### Logistic regression for AMRFinderPlus, RGI, Kleborate AMR Markers

```
combined_logist <- amr_logistic(
  binary_matrix = combined_binary_matrix,
  pheno_drug = "meropenem",
  ecoff_col = "ecoff",
  maf = 10, # filter for AMR markers in at least 10 samples
  single_plot = TRUE
)
#> ...Fitting logistic regression model to R using logistf
#>    Filtered data contains 1490 samples (391 => 1, 1099 => 0) and 14 variables.
#> ...Fitting logistic regression model to NWT using logistf
#>    Filtered data contains 1490 samples (780 => 1, 710 => 0) and 14 variables.
#> Generating plots
#> Plotting 2 models
```

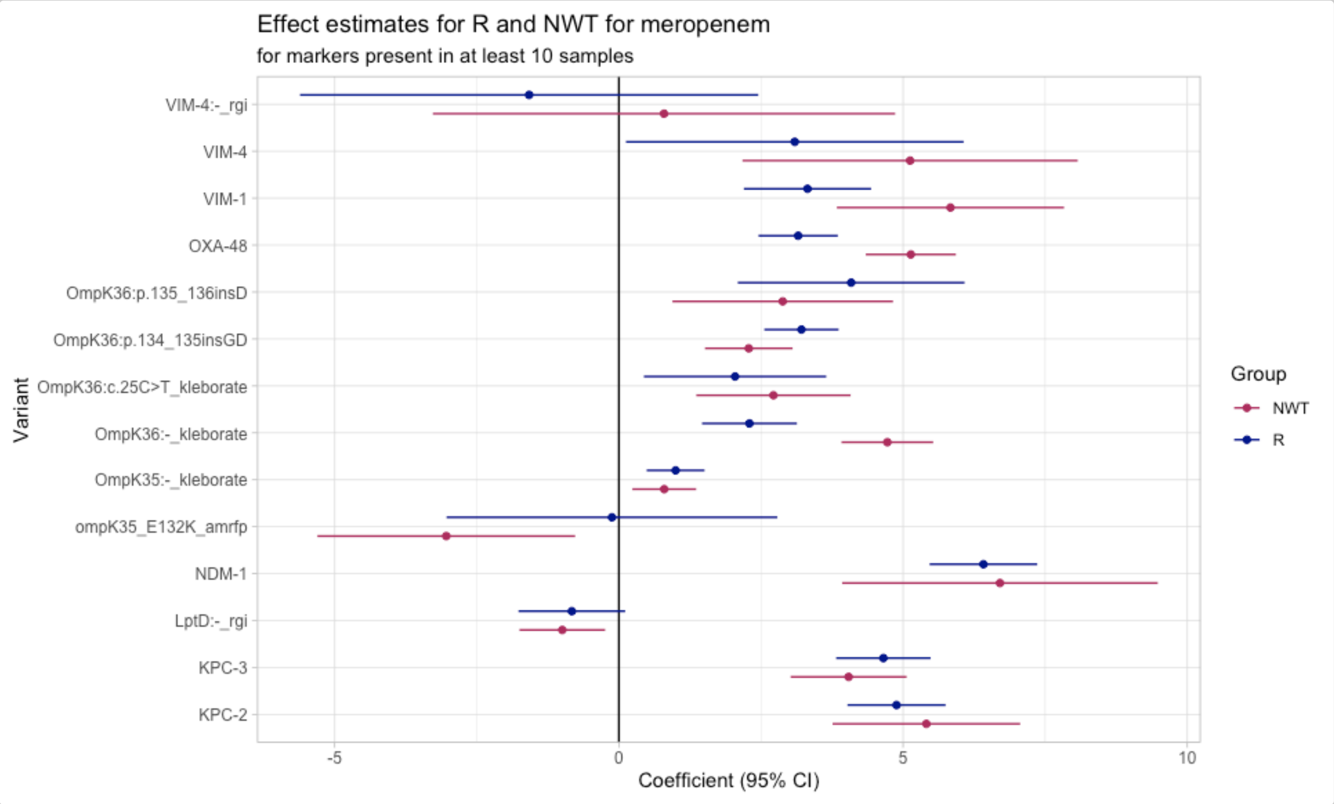


```
# model coefficients
combined_logist$modelR
#> # A tibble: 15 × 5
#>    marker                   est ci.lower ci.upper        pval
#>    <chr>                  <dbl>    <dbl>    <dbl>       <dbl>
#>  1 (Intercept)           -5.23    -5.91    -4.55  0
#>  2 NDM-1                  6.41     5.47     7.36  0
#>  3 KPC-3                  4.65     3.82     5.48  0
#>  4 KPC-2                  4.88     4.02     5.75  0
#>  5 LptD:-_rgi            -0.828   -1.77     0.111 0.0839
#>  6 VIM-1                  3.32     2.20     4.44  0.00000000623
#>  7 VIM-4:-_rgi           -1.58    -5.61     2.45  0.443
#>  8 VIM-4                  3.09     0.124    6.07  0.0411
```

```
#>  9 OXA-48                  3.15     2.46     3.85  0
#> 10 OmpK36:p.134_135insGD   3.21     2.56     3.87  0
#> 11 OmpK36:p.135_136insD    4.09     2.09     6.08  0.0000597
#> 12 ompK35_E132K_amrfp     -0.122   -3.03     2.79  0.935
#> 13 OmpK36:-_kleborate      2.30     1.46     3.13  0.0000000689
#> 14 OmpK35:-_kleborate      0.998    0.489    1.51  0.000123
#> 15 OmpK36:c.25C>T_kleborate 2.04    0.440    3.65  0.0125
#> Use ggplot2::autoplot() on this output to visualise
```

A coefficient above zero indicates that the presence of the AMR marker increases the likelihood of resistance, whereas a coefficient below zero indicates a decrease in probability of resistance. From the plot above showing AMR markers found in more than 10 isolates, majority have a positive association with resistance with the exception of LptD:- (identified by RGI), ompK35_E132K (identified by AMRFinderPlus), and VIM-4:- (identified by RGI).

#### Combinatorial PPV Analysis for AMRFinderPlus, RGI, Kleborate AMR Markers

```
comboPPV_combined_mero <- amr_ppv(
  binary_matrix = combined_binary_matrix,
  order = "value",
  min_set_size = 2,
  pheno_drug = "Meropenem",
  upset_grid = TRUE,
  plot_assay = TRUE,
  assay = "mic"
)
#> Ordering markers by frequency
#> Scale for y is already present.
#> Adding another scale for y, which will replace the existing scale.
```

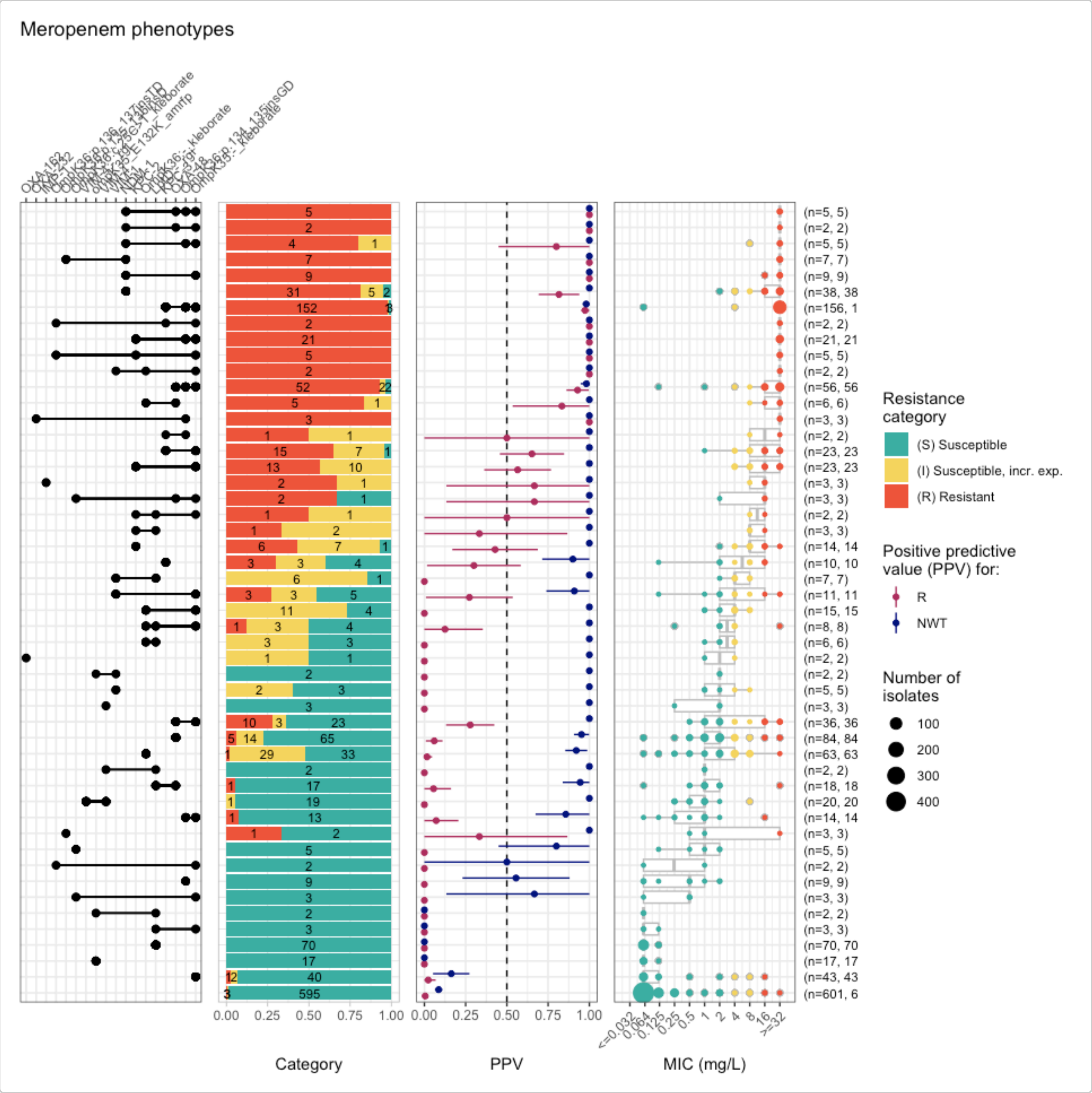


```
comboPPV_combined_mero$summary
#> # A tibble: 86 × 21
#>    marker_list         marker_count     n combination_id   R.n   R.ppv R.ci_lower
#>    <chr>                      <dbl> <int> <fct>          <dbl>   <dbl>      <dbl>
#>  1 ""                             0   601 0_0_0_0_0_0_0…     3 0.00499          0
#>  2 "OmpK36:c.25C>T_k…             1     5 0_0_0_0_0_0_0…     0 0                0
#>  3 "OmpK35:-_klebora…             1    43 0_0_0_0_0_0_0…     1 0.0233           0
#>  4 "OmpK35:-_klebora…             2     3 0_0_0_0_0_0_0…     0 0                0
#>  5 "OmpK36:-_klebora…             1    63 0_0_0_0_0_0_0…     1 0.0159           0
#>  6 "OmpK36:-_klebora…             2     1 0_0_0_0_0_0_0…     1 1                1
#>  7 "OmpK36:-_klebora…             2    15 0_0_0_0_0_0_0…     0 0                0
#>  8 "OXA-162"                      1     2 0_0_0_0_0_0_0…     0 0                0
#>  9 "OXA-427_amrfp, O…             2     1 0_0_0_0_0_0_0…     0 0                0
#> 10 "NDM_amrfp"                    1     1 0_0_0_0_0_0_0…     0 0                0
#> # ℹ 76 more rows
#> # ℹ 14 more variables: R.ci_upper <dbl>, R.denom <int>, NWT.n <dbl>,
#> #   NWT.ppv <dbl>, NWT.ci_lower <dbl>, NWT.ci_upper <dbl>, NWT.denom <int>,
```

```
#> #   median_excludeRangeValues <dbl>, q25_excludeRangeValues <dbl>,
#> #   q75_excludeRangeValues <dbl>, n_excludeRangeValues <int>,
#> #   median_ignoreRanges <dbl>, q25_ignoreRanges <dbl>, q75_ignoreRanges <dbl>
```
